# Elevational variation in heart mass and suppression of hypoxia-induced right ventricle hypertrophy in Andean leaf-eared mice (*Phyllotis*)

**DOI:** 10.1101/2025.08.07.669011

**Authors:** Naim M. Bautista, Nathanael D. Herrera, Marcial Quiroga-Carmona, Chandrasekhar Natarajan, Adriana Rico-Cernohorska, Jorge Salazar Bravo, Graham R. Scott, Guillermo D’Elía, Zachary A. Cheviron, Jay F. Storz

**Affiliations:** School of Biological Sciences, University of Nebraska, Lincoln, NE, United States; Division of Biological Sciences, University of Montana, Missoula, MT, United States; Instituto de Ciencias Ambientales y Evolutivas, Facultad de Ciencias, Universidad Austral de Chile, Valdivia, Chile; Colección de Mamíferos, Facultad de Ciencias, Universidad Austral de Chile, Campus Isla Teja, Valdivia, Chile; Colección Boliviana de Fauna, Instituto de Ecología, Universidad Mayor de San Andrés, La Paz, Bolivia; Department of Biological Sciences, Texas Tech University, Lubbock, TX, United States; Department of Biology, McMaster University, Hamilton, ON, Canada

**Keywords:** High-altitude, hypoxia, hypoxic pulmonary hypertension, junctophilin, maladaptive plasticity

## Abstract

In lowland mammals that ascend to high elevation, hypoxia-induced changes in the pulmonary circulation can give rise to hypoxic pulmonary hypertension (HPH) and associated right-ventricle (RV) hypertrophy. Some mammals that are native to high elevation have evolved a means of attenuating HPH, demonstrating how environmental adaptation may sometimes counteract the effects of ancestral acclimatization responses. Here, we examine elevational variation in heart mass and measures of RV hypertrophy in four closely related species of leaf-eared mice (genus *Phyllotis*) that are broadly co-distributed across a steep elevational gradient on the Western slope of the Andes. There was a positive relationship overall between heart mass and elevation that reflected proportional changes in both the right and left ventricles. Thus, elevation-related increases in overall heart mass were not generally attributable to RV hypertrophy, suggesting that this group of predominantly highland species have evolved a means of avoiding HPH and/or attenuating the cardiac response to HPH. To gain insight into possible transcriptional mechanisms, we examined patterns of transcriptomic variation in the right ventricles of *Phyllotis vaccarum* from two geographically distinct highland populations (both from elevations >5000 m) that exhibit strikingly different levels of RV hypertrophy. Suppression of RV hypertrophy is associated with differential expression of key regulatory genes involved in striated muscle, immune processes, and the inflammatory response. Analysis of co-expression modules identified a promising set of candidate genes for mediating the development of RV hypertrophy at extremely high elevations.

## INTRODUCTION

Theory suggests that colonization of novel environments and adaptation to marginal habitats at range edges can be facilitated directly or indirectly by adaptive phenotypic plasticity (Chevin & Lande, 2011; Yeh & Price, 2004). In some cases, by contrast, environmentally induced changes in phenotype may be costly or otherwise counterproductive in the new habitat, thereby hindering local adaptation. Examples of maladaptive plasticity have been well-documented in lowland mammals that experience environmental hypoxia upon ascent to high elevations (Dempsey & Morgan, 2015; Storz & Scott, 2019, 2021; Storz et al., 2010; West et al., 2021). An especially good example involves hypoxia-induced changes in the pulmonary circulation.

In mammals living at sea level, pulmonary arterial vessels constrict in response to low O_2_, thereby redirecting blood flow to better ventilated, non-hypoxic regions of the lungs. This dynamic matching of ventilation and perfusion enhances the efficiency of pulmonary gas exchange when there is regional variation in O_2_ levels across the lungs. Upon ascent to high elevation, however, O_2_ availability is reduced across the entirety of the lungs due to the reduced O_2_ partial pressure (*P*O_2_) of inspired air. In this situation, a global pulmonary vasoconstrictive response occurs, and within 24-48 h of hypoxia exposure the induced pulmonary vascular remodeling can result in a thickening of arterial smooth muscle (Dorrington et al., 1997; Maggiorini et al., 2001). This combination of vasoconstriction and vascular remodeling reduces the distensibility of pulmonary vessels and increases vascular resistance, which can cause a sharp increase in pulmonary arterial pressure (Shimoda & Laurie, 2014; Stenmark et al., 2006; Swenson & Bärtsch, 2012; Sylvester et al., 2012). This hypoxic pulmonary hypertension (HPH) can dangerously increase right ventricular afterload (the pressure against which the right ventricle [RV] has to contract to eject blood from the heart), which leads to RV hypertrophy and increases risk of heart failure (Soliz et al., 2005; Wang et al., 2013; Young et al., 2019). The increased hydrostatic pressure in pulmonary capillaries can also cause a well-characterized malady known as high-altitude pulmonary edema (Dempsey & Morgan, 2015; Swenson & Bärtsch, 2012).

In the globally hypoxic environment at high elevation, HPH may be exacerbated by an excessive increase in red blood cell production (erythrocytosis), a typical response to hypoxia exposure in lowland mammals. Although an increase in red cell mass increases the O_2_-carrying capacity of the blood, excessive erythrocytosis can increase blood viscosity if it is not accompanied by a concomitant expansion of blood volume and can thus lead to further increases in pulmonary vascular resistance (Naeije et al., 2013).

Some mammals that are native to high-elevation environments appear to have evolved a means of attenuating HPH (Bautista et al., 2024; Ge et al., 1998; Reyes et al., 2020; West et al., 2021), illustrating how environmental adaptation may often involve genetic compensation of maladaptive plasticity (Storz & Scott, 2019, 2021). For example, long-term acclimation experiments with deer mice (*Peromyscus maniculatus*), in which lowland and highland natives were acclimated for 6-8 weeks to chronic hypoxia (equivalent to 4300 m) (West et al., 2021), revealed that in comparison to equally well-acclimated lowland conspecifics, highland deer mice exhibit less pronounced HPH and do not exhibit RV hypertrophy or pulmonary artery thickening (West et al., 2021). The highland natives also maintain normal ventilation-perfusion matching in chronic hypoxia – unlike lowland conspecifics – and exhibit superior aerobic performance capacities in hypoxia (Bautista et al., 2024; Storz et al., 2019; Tate et al., 2020; West et al., 2021). These findings suggest that mice native to highland environments have evolved a means of attenuating certain maladaptive effects of hypoxia acclimation.

Leaf-eared mice in the genus *Phyllotis* are broadly distributed across the Andes and the adjoining Altiplano of South America, and several species have extraordinarily broad elevational distributions along the flanks of the Andean Cordillera (Jayat et al., 2021; Ojeda et al., 2021; Quiroga-Carmona et al., 2025; Steppan & Ramirez, 2015; Storz et al., 2024; Teta et al., 2022). For example, *P. vaccarum* has been documented on the summits of some of the highest peaks in the Central Andes at elevations >6700 m (Storz et al., 2023; Storz et al., 2024; Storz et al., 2020). On the western slope of the Andes, the lower elevational range limit of *P. vaccarum* extends all the way to the desert coastline of northern Chile (Quiroga-Carmona et al., 2025; Storz et al., 2024). Across this same elevational gradient, several other species of *Phyllotis* have distributions that extend from the Atacama Desert to Andean dry puna habitats at elevations between ∼3500-5500 m (Quiroga-Carmona et al., 2025; Storz et al., 2024).

Here, we examine elevational variation in body mass, heart mass, measures of RV hypertrophy, and blood hemoglobin concentration ([Hb]) in four closely related species of *Phyllotis* that are broadly co-distributed across a steep elevational gradient on the Western slope of the Andes (**Fig. 1*A***). The four species, *P. chilensis, P. limatus, P. magister*, and *P. vaccarum*, have partially overlapping elevational ranges on the western slope of the Andes, all with upper limits that exceed 3500 m (Quiroga-Carmona et al., 2025; Steppan & Ramirez, 2015; Storz et al., 2024) (**Fig. 1*B***). Our use of the species name ‘*P. chilensis* ‘(*sensu* (Pearson, 1958) follows previous work (Quiroga-Carmona et al., 2025; Storz et al., 2024) and refers to a phylogenetically distinct form within the species complex that Ojeda et al. (2021) referred to as ‘*P. posticalis*-*rupestris*’.

**Figure 1.**
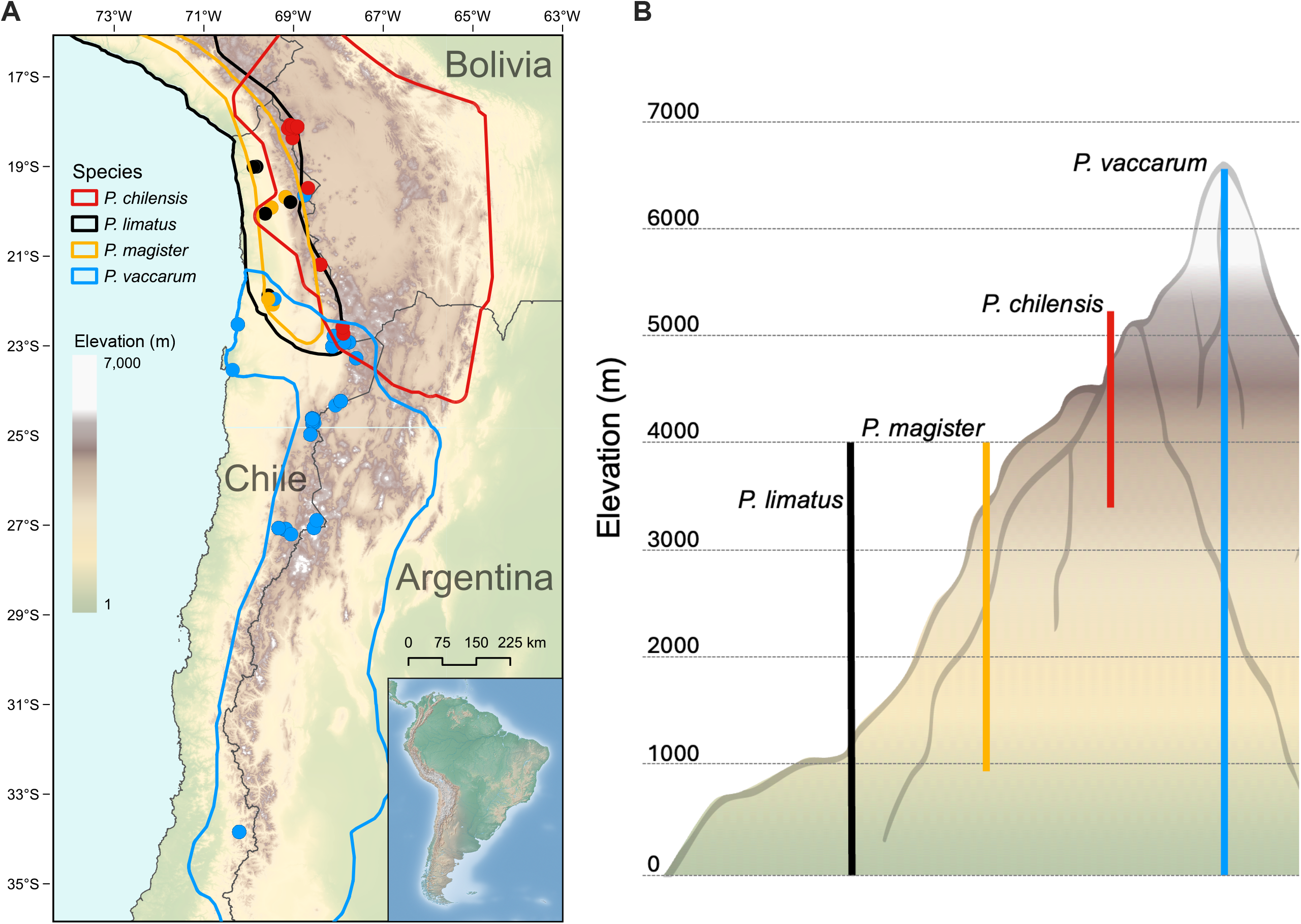
(**A**) Geographic distributions of the four focal species of *Phyllotis*, with color-coded symbols denoting specific sampling localities in Chile and Bolivia. (**B**) Elevational ranges of the four species across the western slope of the Central Andes.

We assess the extent to which elevational changes in heart mass are driven by RV hypertrophy, and we examine whether species that are native to especially high elevations such as *P. vaccarum* exhibit qualitatively or quantitatively different patterns of variation from the other species of *Phyllotis*. To gain insight into possible transcriptional mechanisms involved in the suppression of RV hypertrophy, we also examine patterns of transcriptomic variation in the right ventricles of *P. vaccarum* from two geographically distinct highland populations (both from elevations >5000 m) that exhibit strikingly different levels of RV hypertrophy.

## MATERIALS AND METHODS

During the course of small mammal surveys in the Atacama Desert and Andean dry puna of northern Chile and western Bolivia (2020-2022), we trapped *Phyllotis* mice at sites that spanned a diversity of habitats over a broad range of elevations, from near sea level to volcano summits of >6700 m (**Fig. 1*A*, Table S1**). For details regarding trapping protocols at extreme elevations, see Storz et al. (2020, 2024). We identified all specimens to the species level based on external characters (Patton et al., 2015) and we later confirmed field-based identifications with mitochondrial *cytochrome b* sequence data and low coverage whole-genome sequence data (Quiroga-Carmona et al., 2025; Storz et al., 2024).

### Live-Trapping and Specimen Preparation

We captured mice using Sherman live traps. We sampled blood for haematological measurements, then sacrificed animals in the field and prepared them as museum specimens after dissecting and preserving heart and liver in RNAlater™ (Invitrogen, Thermo Fisher Scientific, AM7021). All voucher specimens are housed in the Colección de Mamíferos of the Universidad Austral de Chile, Valdivia, Chile (UACH) or in the Colección Boliviana de Fauna (CBF), La Paz, Bolivia.

In Chile, all animals were collected in accordance with permissions to JFS, GD, and MQC from the following government agencies: Servicio Agrícola y Ganadero (SAG, Resolución exenta #’s 6633/2020, 2373/2021, 5799/2021, 3204/2022, 3565/2022, 911/2023, and 7736/2023), Corporación Nacional Forestal (CONAF, Autorización #’s 171219, 1501221, and 31362839), and Dirección Nacional de Fronteras y Límites del Estado (DIFROL, Autorización de Expedición Científica #68 and 02/22). In Bolivia, all animals were collected in accordance with permissions to ARC, JS-B, and JFS from the Ministerio de Medio Ambiente y Agua, Estado Plurinacional de Bolivia (CAR/MMAYA/VMABCCGDF/DGBAP/MEG No 6114/2021, 9 September 2021; MMAYA/VMABCCGDF/DGBAP/MEG No 0059/2023, 28 March 2023) and from National Protected Areas Service (SERNAP-CAR/DMA No 2158/2022 - HR 07755, 14 November 2022). All live-trapped animals were handled in accordance with protocols approved by the Institutional Animal Care and Use Committee of the University of Nebraska (project ID’s: 1919, 2100) and the bioethics committee of the Universidad Austral de Chile (certificate 456/2022).

### Measurements of cardiac and hematological phenotypes

In the lab, masses of the entire heart were measured using an analytical top-balance (Ohaus Corp., SC2020, Pine Brook, NJ, USA). For each mouse, the left ventricle (including the septum) and the right ventricle were dissected under a stereomicroscope and were then weighed separately. For each specimen, measured masses were used to calculate Fulton’s index (mass of right ventricle divided by the combined mass of the left ventricle and interventricular septum). We estimated blood hemoglobin ([Hb]) concentration with triplicate measurements of the blood that was freshly drawn shortly after capture using a Hemocue +201 Hemoglobin Analyzer 121721 (Hemocue America).

### Statistical analysis of morphological/physiological phenotypes

Statistical analysis and graphing were performed in R v4.4.0 (https://cran.r-project.org/), and R-studio v2024.04.0+735 (https://www.rstudio.com/). We used linear regression to assess relationships between phenotypic traits and native elevation in each species. Tests for phenotypic differences between population samples were performed with unpaired t-test.

### Sample preparation for RNA-seq

We used high-throughput RNA sequencing of the right ventricle to examine gene expression variation among individual *P. vaccarum* that we collected from a pair of high-elevation localities on the flanks of two Atacama volcanoes: 5250 m on the flanks of Ojos del Salado (*n*=6) and 5070 m on the flanks of Volcán Llullaillaco (*n*=7). Libraries were prepared and sequenced by Azenta Life Technologies. Briefly, for each sample, RNA was extracted from the right ventricle of the heart using a SMART-Seq HT kit for full-length cDNA synthesis and amplification. Sequencing libraries were prepared using an Illumina Nextera X kit, quality checked using a TapeStation and then sequenced on a shared lane of Illumina HiSeq 4000 (PE 150 bp) platform by Azenta.

### RNA-Seq data processing and transcriptomic analyses

All pipeline scripts/ command and data files are freely available via GitHub (https://github.com/NathanaeldHerrera/Phyllotis_Right_ventricle_hypertropy). Raw RNA-Seq reads were trimmed and filtered using FastP (Chen et al., 2018). Adapter sequences were automatically detected and removed, and a 5 bp sliding window was applied from the 5′ to 3′ end to trim bases with a mean quality score below 20. Reads shorter than 30 bp after trimming were discarded. Read quality was assessed using MultiQC v1.11 (Ewels et al., 2016). Cleaned reads were aligned to the *Phyllotis vaccarum* reference genome (Storz et al., 2023) using HiSat2 v2.2.1 (Zhang et al., 2021), with mismatch penalties set to a maximum and minimum of 2 (−-mp 2,2). Gene-level read counts were generated using featureCounts v2.0.3 (Liao et al., 2014), with the -O flag enabled to count reads overlapping multiple annotated features, assigning one count to each overlapping gene.

To identify outliers that may be attributable to sequencing artifacts, we conducted a principal components analysis (PCA) using the base R function prcomp on five sequencing metrics: (1) mean base quality score, (2) percent of bases with quality ≥30, (3) GC content, (4) genome alignment rate, and (5) percent of reads assigned to genes. The first two principal components were visualized using ggbiplot v0.6.2 (https://github.com/friendly/ggbiplot), and samples outside the 95% confidence ellipse were excluded from further analysis. Because these are technical quality metrics of the sequencing experiment, we do not expect substantial variation among samples. Thus, using a 95% CI provides a reasonable threshold for identifying samples with potential sequencing artifacts rather than reflecting natural biological variation. This reduced the final sample size to 10 mice (*n*=6 from Ojos del Salado, *n*=4 from Llullaillaco). To reduce measurement error from low-expression genes, we excluded those with fewer than 20 average reads across individuals, retaining 11,294 genes. Read counts were normalized for library size using the calcNormFactors function from the edgeR package v3.42.4 (Robinson et al., 2010), and log-transformed counts per million (log-CPM) were computed using the cpm function for downstream analyses.

We used a combined analytical framework to evaluate gene expression profiles of mice from Ojos del Salado and Llullaillaco and to test for associations between expression profiles and Fulton’s index. Differential expression (DE) analysis treats genes individually and identifies particular genes that exhibit significant expression differences between samples from the two populations that exhibited differences in RV hypertrophy. As a complementary approach, we used Weighted Gene Co-expression Network Analysis (WGCNA) to identify modules of co-expressed genes, which enabled us to discover coordinated regulatory responses and module-level associations with the measured phenotypes.

#### Differential expression analysis

We used edgeR to identify genes differentially expressed between populations by modeling normalized raw counts with quasi-likelihood negative binomial generalized linear models. Population of origin (Ojos del Salado or Llullaillaco) was treated as a categorical predictor in the design matrix with Llullaillaco set as the reference population. Significance was assessed using quasi-likelihood F-tests (implemented with the glmQLFit, glmQLFTest functions), and multiple testing was controlled using the Benjamini–Hochberg false discovery rate FDR (Benjamini & Hochberg, 1995). Genes were considered differentially expressed at FDR ≤ 0.1.

For genes that exhibited significant differential expression between the two population samples, we performed gene ontology (GO) and KEGG pathway enrichment analysis using the gost function in gProfiler (Raudvere et al., 2019). *Mus musculus* orthologs for each *Phyllotis* gene were used as queries, with the background consisting of all expressed genes in the right ventricle. Enrichment significance was defined at FDR < 0.05. To facilitate integration with WGCNA, we also generated a log-transformed expression matrix (logCPM) from the normalized counts using edgeR.

#### Weighted gene co-expression network analysis (WGCNA)

Co-expression network analysis was conducted using the R package WGCNA v1.72-1 (Langfelder & Horvath, 2008) on the logCPM matrix. We used a signed, scale-free network, which groups genes according to whether observed correlations are positive or negative. Networks were constructed using a soft-thresholding power of 25, selected based on scale-free topology criteria (Zhang & Horvath, 2005). Modules were identified by hierarchical clustering with dynamic tree cutting (minimum module size = 30; merge cut height = 0.25), and hub genes for each module were identified based on module membership (kME). After defining modules, we used a multistep approach to test for associations between module expression and Fulton’s index. We first summarized module expression using a PCA of gene expression profiles using the blockwiseModules function in WGCNA. Given that genes within modules are inherently correlated, we used the first principal component axis, also known as the module eigengene, to represent module expression (Langfelder & Horvath, 2008). We then used module eigengene values to test for associations between module expression and Fulton’s index (using Pearson correlation; cor function in WGCNA). We also conducted an analysis-of-variance (ANOVA) on rank-transformed module eigengene values to identify which gene co-expression modules show a significant association with the phenotype (Plachetzki et al., 2014). Module–trait associations were considered significant at a *P*-value < 0.05. Modules containing at least 10 genes were annotated using GO and KEGG enrichment via gProfiler, as described above.

## RESULTS AND DISCUSSION

We collected phenotypic data for a total of 216 *Phyllotis* mice representing four species: *P. chilensis* (*n* = 46), *P. limatus* (*n* = 21), *P. magister* (*n* = 7), and *P. vaccarum* (*n* = 142). For each species, the elevational range of sampling approximates the known elevational distribution. In the case of *P. chilensis* and *P. vaccarum*, the specimens that we collected from the highest localities (5221 and 6739 m, respectively) also represent the highest elevational records for those particular species (Storz et al., 2024).

### Elevational variation in body mass

Regression of body mass against elevation revealed no statistically significant relationship in any species other than *P. chilensis*, which exhibited a negative relationship (*R* = -0.36, *P* = 0.016) (**Fig. 2*A***). Our results for *Phyllotis* are not consistent with an elevational version of Bergmann’s rule, the empirical generalization that body mass generally increases towards more polar (colder) latitudes.

**Figure 2.**
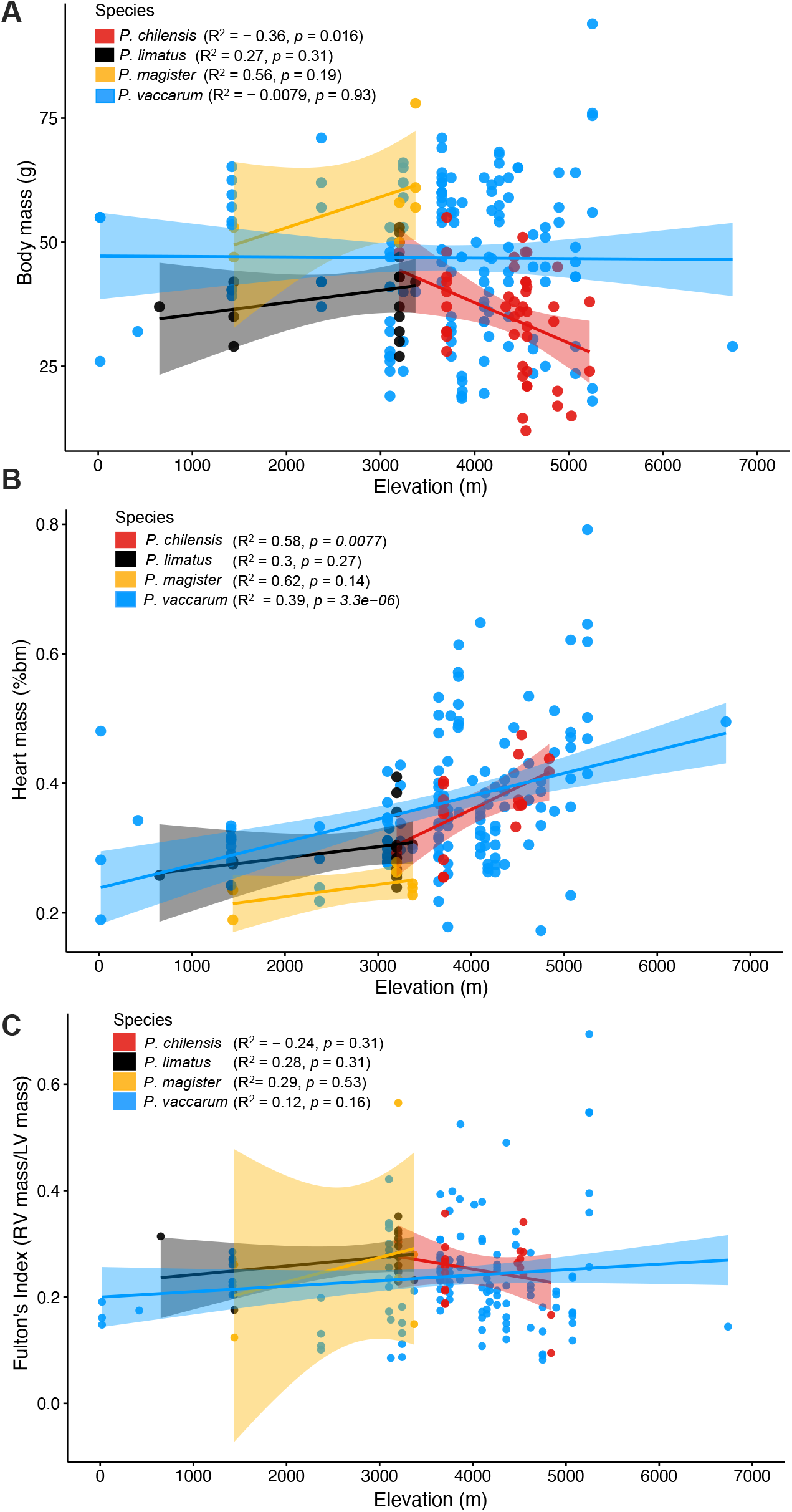
Elevational variation in body mass (**A**), heart mass (relative to body mass, bm) (**B**), and Fulton’s index (right ventricle mass relative to left ventricle and septum mass) (**C**) in four species of *Phyllotis* sampled across the western slope of the Central Andes. *R* = Pearson’s correlation coefficient.

### Elevational variation in heart mass, Fulton’s index, and [Hb]

All species exhibited a positive relationship between heart mass and elevation (*R*^2^ = 0.29 – 0.62; **Fig. 2*B***), although the relationship was only statistically significant for *P. chilensis* and *P. vaccarum* (*P* < 0.01), the two species that have the highest upper elevational range limits. Fulton’s Index did not show a significant positive association with elevation (**Fig. 2*C***), indicating that variation in heart mass largely reflects proportional changes in both the right and left ventricles. Thus, the positive relationship between heart mass and elevation in *Phyllotis* mice is not generally attributable to RV hypertrophy, suggesting that this group of predominantly highland species may have evolved a means of attenuating HPH, even at the extreme elevations inhabited by *P. chilensis* and *P. vaccarum*. Most species exhibited the expected trend of increasing [Hb] as a function of elevation, but the correlation was only significant for *P. limatus* and *P. vaccarum* (**Fig. 3**).

**Figure 3.**
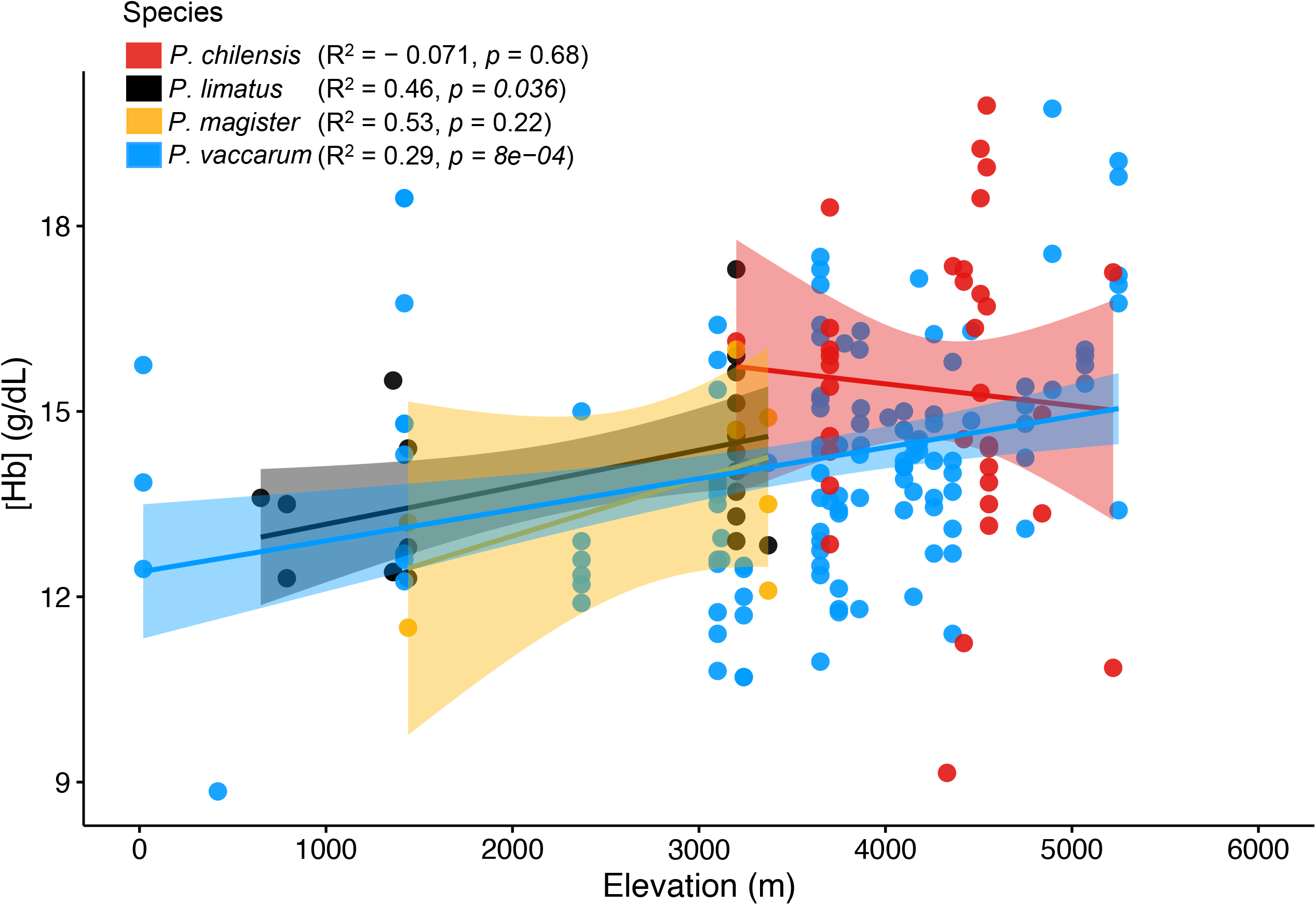
Elevational variation in hemoglobin concentration ([Hb]) and elevation in four species of *Phyllotis* sampled across the western slope of the Central Andes. *R* = Pearson’s correlation coefficient.

Within the most broadly distributed species, *P. vaccarum*, a close inspection of the data suggests some population-specific idiosyncrasies with regard to elevational trends in relative RV mass. In this species, the slight positive trend in the relationship between Fulton’s Index and elevation is largely driven by a sample of extreme high-elevation mice from the the flanks of Ojos de Salado at 5250 m, all of which show evidence for RV hypertrophy (**Fig. 4*A***). By contrast, a sample of mice from a similarly high elevation (5070 m) on the flanks of Volcán Llullaillaco show no such pattern (**Fig. 4*A***). Mice from the two localities did not show significant differences in [Hb] (**Fig. 4*B***). The observed difference in Fulton’s Index between the Ojos del Salado and Llullaillaco mice is not explained by population genetic structure, as mice from both localities form part of an exceedingly well-mixed gene pool in the Puna de Atacama (Quiroga-Carmona et al., 2025; Storz et al., 2023; Storz et al., 2024). However, there are reasons to suspect that the mice from Ojos del Salado have a shorter history of residence at extreme elevation than the mice from Llullaillaco. The mice from 5070 m on the flanks of Llullaillaco were captured in an extremely remote, undisturbed setting that approximates the elevational limits of vegetation. By contrast, the mice from 5250 m on Ojos del Salado were captured well above vegetation limits in the immediate environs of Refugio Atacama, a rustic hut that is used as a base camp shelter by mountain climbers. It is likely that the mice are present so far above vegetation limits in and around Refugio Atacama because they are attracted by food caches left by climbers. In contrast to the Llullaillaco mice that may have had many generations to adapt to the rarefied atmosphere at >5000 m, the Ojos del Salado mice may have only recently colonized the environs of Refugio Atacama from lower elevations. This provides a possible explanation for why the Ojos del Salado mice appear to present a less well-adapted phenotypic profile (RV hypertrophy) that might be expected for recent immigrants from lower elevations.

**Figure 4.**
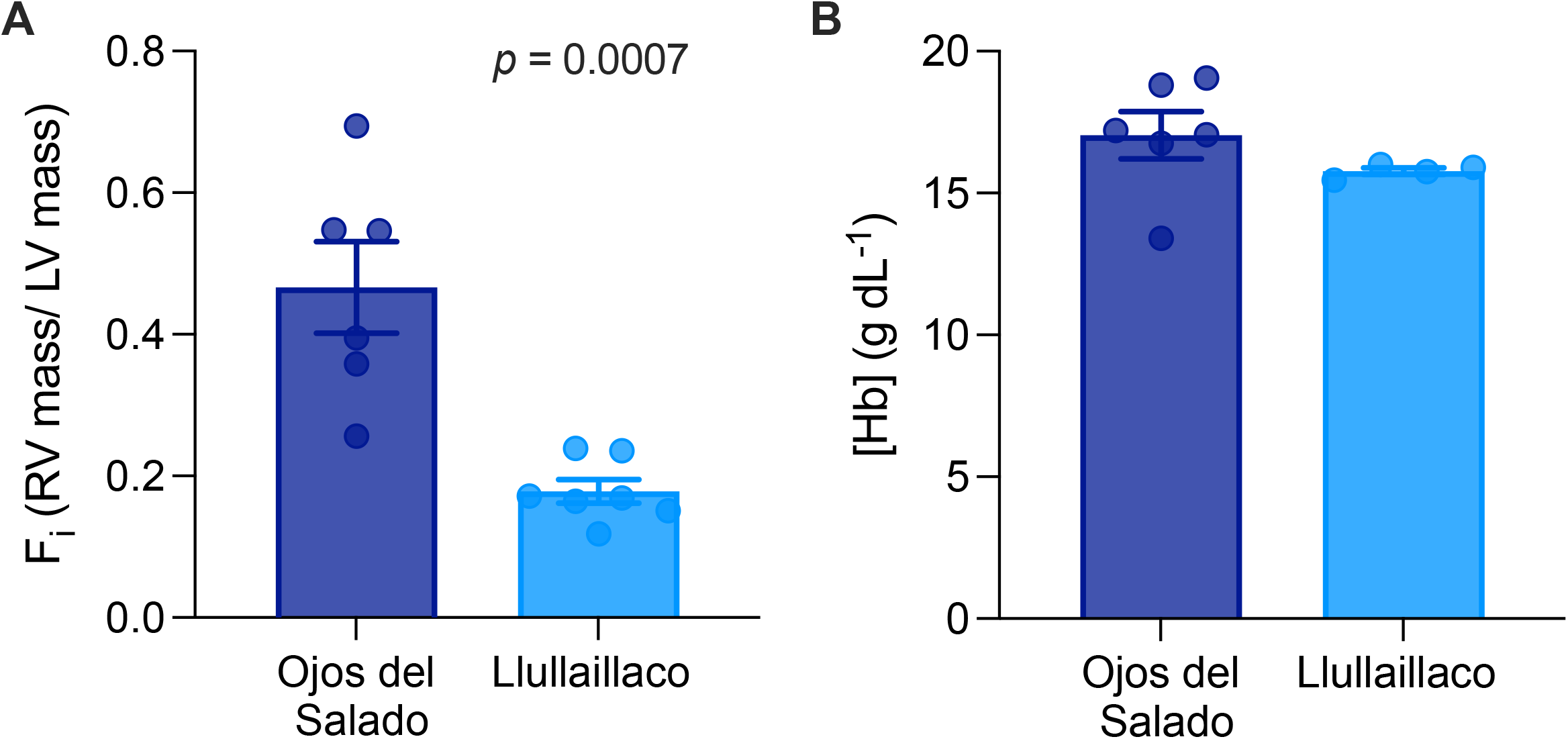
Tests for differences in Fulton’s index (F_i_, right ventricle mass relative to left ventricle and septum mass) and blood Hb concentration ([Hb]) between *P. vaccarum* from two geographically distinct high-elevation localities on the flanks of Ojos del Salado (5250 m) and Volcán Llullaillaco (5070 m).

The differences in Fulton’s index in mice from these two extreme high-elevation localities provides an opportunity to investigate regulatory mechanisms underlying hypoxia-induced changes in heart mass. Accordingly, we generated RNA-seq data for RV of all sampled mice from the two >5000 m localities. Gene expression differences between the two samples of highland mice could reveal regulatory mechanisms underlying RV hypertrophy in the Ojos del Salado mice (i.e., the transcriptional basis of maladaptive plasticity) and/or regulatory mechanisms underlying the suppression of RV hypertrophy in the Llullaillaco mice (i.e., the transcriptional basis of genetic compensation).

We took a two-pronged approach to identify the regulatory basis of population differences in RV hypertrophy. We first performed a standard differential expression analysis to identify genes that were differentially expressed between the two populations to obtain a gene-centered view of transcriptional differences that may contribute to variation in hypoxia-induced RV hypertrophy. Second, using the same RNA-seq dataset, we applied a network co-expression analysis (WGCNA; Langfelder & Horvath, 2008) to identify suites of co-regulated genes (transcriptional modules) and we then tested for associations between module expression and phenotype. This network-based analysis provided insight into regulatory networks that are specifically associated with cardiac phenotypes, and it identified key regulatory genes that may coordinate the expression of these networks.

We obtained RNA-Seq data for 10 RV samples [Ojos del Salado (*n*=6) and Llullaillaco (*n*=4); mean number of reads per individual = 45,945,408], with a mean genome alignment rate of 89.2% ± 1.6% and a mean feature counts assignment rate of 35.9% ±4.0% (**Table S2**). In the differential expression analysis, we identified a total of 278 genes that were differentially expressed (FDR < 0.1) between populations (**Fig. 5, Table S3**). Gene ontology (GO) enrichment analysis revealed that this suite of differentially expressed genes was significantly enriched for 65 GO categories, many of which are related to striated muscle structures and differentiation to immune system processes (**Table S4**). This result mirrors a pattern found in similar comparisons of highland and lowland deer mice (*Peromyscus maniculatus*) that differ in their propensity to develop hypoxia-induced RV hypertrophy. Highland deer mice sampled from the summit of Mt. Blue Sky in Colorado, USA (∼4350 m a.s.l.), exhibit reduced RV hypertrophy relative to lowland congeners when reared under the same conditions of chronic hypoxia (Velotta et al., 2018). The suppression of RV hypertrophy in highland deer mice is associated with differential expression of key regulatory genes involved in immune processes and the inflammatory response (Velotta et al., 2018). Interestingly, 68 of the 278 genes that were differentially expressed between highland populations of *P. vaccarum* overlapped with those identified in highland deer mice (**Table S5**), suggesting evolutionary conservation of regulatory mechanisms that underlie RV hypertrophy. Gene ontology enrichment analysis of this subset of overlapping genes was again significantly enriched for genes that participate in immune processes (e.g. GO:0019883 - antigen processing and presentation of endogenous antigen; **Table S6**). In both *P. vaccarum* and *P. maniculatus*, hypoxia-induced expression of genes involved in immune signaling is associated with RV hypertrophy, but it is unclear whether the observed changes in gene expression are the cause or consequence of right heart overgrowth (Velotta et al., 2018).

**Figure 5.**
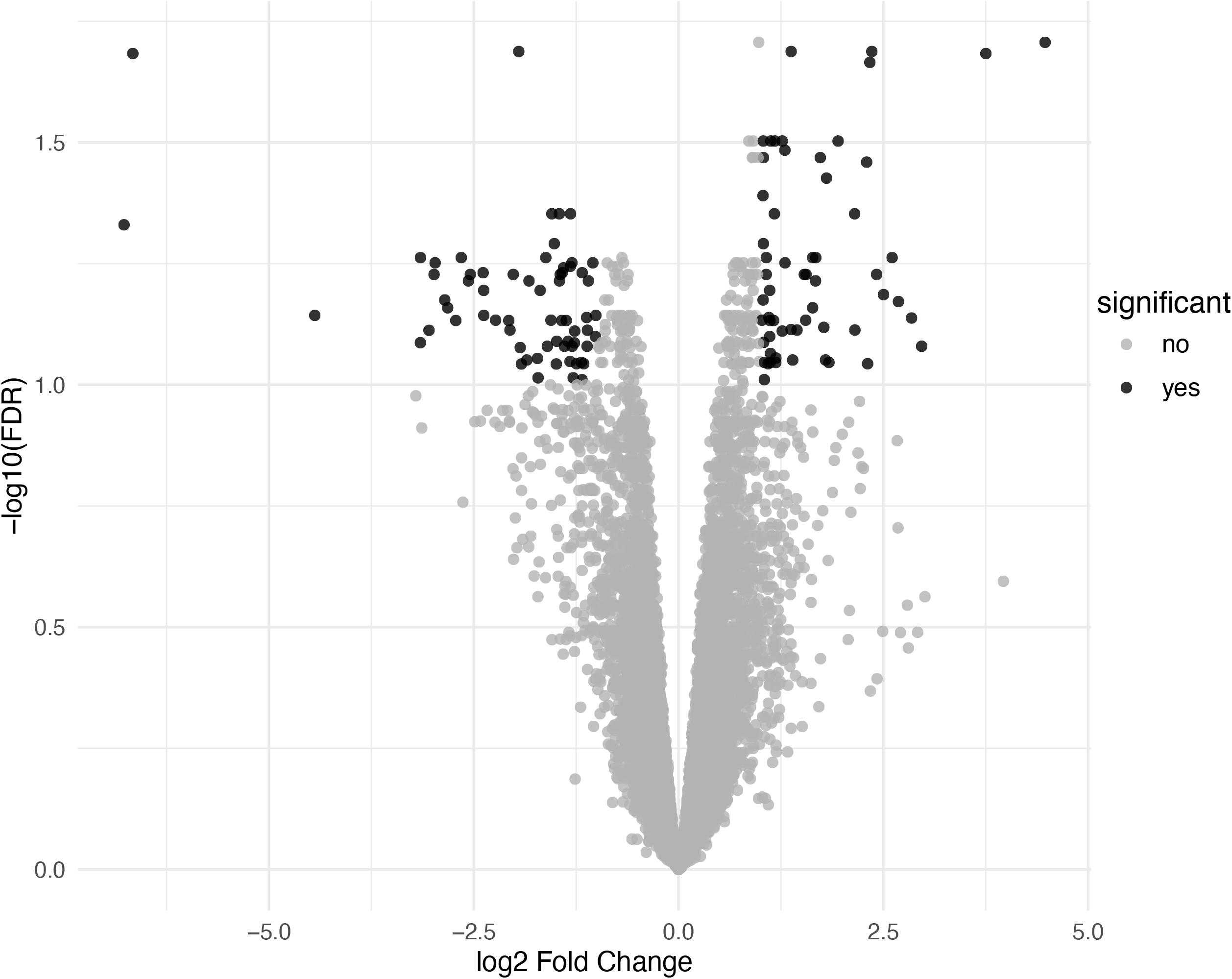
Volcano plot showing significant (FDR ≤ 0.1) DE genes between high-elevation samples of *P. vaccarum* from the flanks of two Atacama volcanoes, Ojos del Salado and Llullaillaco. The two populations exhibited pronounced differences in RV hypertrophy (Fig. 4*A*). Population was treated as a categorical predictor in the design matrix with Llullaillaco as the “reference”. We identified 148 significant DE genes showing increased expression in Ojos relative to Llullaillaco (> 0 log2 Fold Change) and 130 significant DE genes showing increased expression in Llullaillaco relative to Ojos (< 0 log2 Fold Change).

Physiological processes are regulated by suites of genes that interact in transcriptional networks whose structure can be inferred by identifying and characterizing co-expression modules. We identified a total of 103 transcriptional modules that collectively contained 11,271 genes (99.8% of the right ventricle transcriptome). Transcriptional modules ranged in size from 23-854 genes. Of these 103 modules, ten were associated with Fulton’s Index (*P* < 0.05) (**Table S7**). Although none of these associations remained significant after correcting for the false discovery rate, we characterized modules that were associated with Fulton’s index to gain insights into the regulatory networks that might underlie RV hypertrophy. Of these trait-associated modules, six exhibited positive associations between module expression and Fulton’s index, and four exhibited negative associations (**Fig. 6, Table S7**). These associations were driven primarily by pronounced differences in module expression between mice from the two different localities. Negatively-associated modules were enriched for functions related to carbohydrate metabolism (RV77), regulation of gene expression (RV19), and other biosynthetic processes (RV70), while positively associated modules were enriched for genes that participate in cardiac development and growth (module RV2), hypertrophic cardiomyopathy, blood clotting and coagulation (RV81), and motor and contractile processes (RV71) (**Table S8**). Of particular interest is module RV2 (*R*^2^ = 0.42; *P* = 0.043) which contains a total of 812 genes, dozens of which participate in processes known to impact heart growth and development **Table S8-S9)**. The hub gene of this module (**Table S10**), junctophilin 2 (*Jph2*), encodes a cardiac structural protein that is involved in the assembly of junctional membrane complexes in cardiomyocytes and plays an essential role in maintaining intracellular calcium homeostasis and normal cardiac contractility (Beavers et al., 2014; Takeshima et al., 2000). Downregulation of *Jph2* is associated with pathological cardiac remodeling (Landstrom et al., 2011; Minamisawa et al., 2004), and overexpression of the gene in models of cardiac hypertrophy has been shown to prevent the progression to heart failure (Guo et al., 2014; Reynolds et al., 2016). Variation in the expression of *Jph2* and other co-regulated genes in module RV2 may also contribute to variation in the propensity for RV hypertrophy in *P. vaccarum*. Upregulation of *Jph2* in the Ojos del Salado mice may reflect a compensatory response to HPH-related cardiac stress, perhaps attenuating RV hypertrophy and slowing the progression towards heart failure. The set of genes comprising RV2 provide a promising set of candidates for investigating transcriptional mechanisms that underlie the protective attenuation of RV hypertrophy.

**Figure 6.**
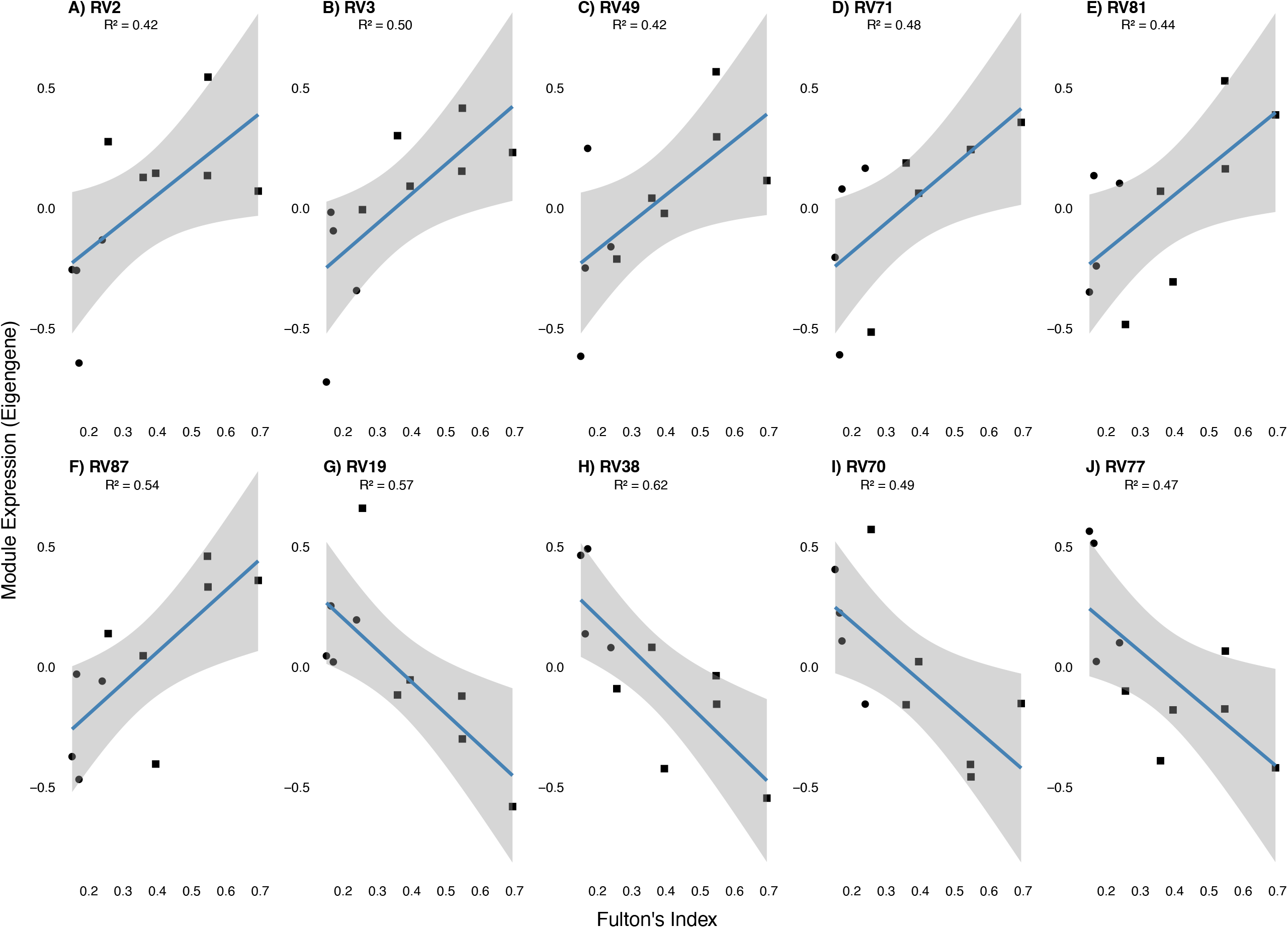
Statistically significant associations between right ventricle (RV) transcriptional modules and Fulton’s index in high-elevation *P. vaccarum* sampled from the flanks of Ojos del Salado (squares) and Volcán Llullaillaco (circles). Six modules exhibited positive associations with Fulton’s index (A-F) and four exhibited negative associations (G-J).

## Conclusion

In hypoxic conditions at high elevation, an increase in overall heart size may be physiologically beneficial to the extent that it reflects a proportional increase in the size of the ventricles and likely permits increased stroke volume and cardiac output, which may be vital for supporting the increased metabolic demands of life in the cold and hypoxic conditions at high elevation (Hayes, 1989). Indeed, relative heart size is an informative predictor of aerobic performance capacity in cold hypoxia (Bech & Østnes, 1999; Chappell et al., 1999; Hammond et al., 2000; Hammond et al., 2001; Shirkey & Hammond, 2014; Tate et al., 2020; Wearing & Scott, 2022). However, an increase in overall heart size that is attributable to disproportionate RV hypertrophy is a symptom of HPH, a clearly maladaptive response to hypoxia. Thus, at high elevation, native highlanders and recently arrived lowland immigrants may both exhibit hypoxia-induced increases in heart size, but available evidence suggests that in well-adapted, high-elevation mammals, such increases in heart size are not accompanied by disproportionate RV hypertrophy (Storz & Scott, 2019; West et al., 2021). The mice collected at elevations >5000 m on the flanks of Ojos del Salado and Llullaillaco experience levels of chronic environmental hypoxia that are more than sufficient to induce HPH in humans and other lowland mammals. However, RV hypertrophy is only observed in the Ojos del Salado mice and not in the Llullaillaco mice, suggesting that the latter represent a good model for discovering protective mechanisms of attenuating HPH and RV hypertrophy. Results of the transcriptomic analysis and observed parallels in other high-elevation rodents suggest promising hypotheses about causative mechanisms, especially with regard to *Jph2* and associated genes in the same regulatory network.

## STATEMENTS

### Ethical statement

All protocols of animal handling and sampling were performed in accordance with the

## Author contributions

**N.M.B**. conceptualization, methodology, formal analysis, investigation, writing – original draft, writing – review and editing; **N.D.H**, conceptualization, methodology, formal analysis, investigation, visualization; **M.Q.C**., investigation, visualization, writing – review and editing,; **C.N**., investigation; **A.R.C**., investigation, review and editing; **J.S.B**., investigation, review and editing; **G.R.S**., investigation, writing – original draft, writing – review and editing; **G.D**., investigation, writing – review and editing; **Z.A.C**., conceptualization, supervision; **J.F.S**. conceptualization, methodology, formal analysis, investigation, writing – original draft, writing – review and editing, project administration, supervision.

All authors have read and approved the final version of the manuscript submitted for publication and agree to be accountable for all aspects of the work in ensuring that questions related to the accuracy or integrity of any part of the work are appropriately investigated and resolved. All persons designated as authors qualify for authorship, and all those who qualify for authorship are listed.

## Data availability

Read sequence data are available for download at the SRA under BioProject accession number PRJNAXXXX. Raw phenotypic data, rna-seq QC metrics, raw featureCounts data, and associated metadata, are provided with pipeline scripts/commands via GitHub (https://github.com/NathanaeldHerrera/Phyllotis_Right_ventricle_hypertropy).

