## Supplementary Material for "Elevational variation in heart mass and suppression of hypoxia-induced right ventricle hypertrophy in Andean leaf-eared mice (*Phyllotis*)"

**This PDF file includes:**

Tables S1 to S10

**Table S1.** List of Vouchered specimens of *Phyllotis* species used in this study.

| Num_cat | Species | Locality | Elevation | Latitude_DD | Longitude_DD |
|---|---|---|---|---|---|
| GD2024 | *P. vaccarum* | Chile, Antofagasta, San Pedro de Atacama, Ruta B-241 km 117 | 2370 | -22.920033 | -67.767367 |
| GD2025 | *P. vaccarum* | Chile, Antofagasta, San Pedro de Atacama, Ruta B-241 km 117 | 2370 | -22.920033 | -67.767367 |
| GD2026 | *P. vaccarum* | Chile, Antofagasta, San Pedro de Atacama, Ruta B-241 km 117 | 2370 | -22.920033 | -67.767367 |
| GD2028 | *P. vaccarum* | Chile, Antofagasta, San Pedro de Atacama, Ruta B-241 km 117 | 2370 | -22.920033 | -67.767367 |
| GD2029 | *P. vaccarum* | Chile, Antofagasta, San Pedro de Atacama, Ruta B-241 km 117 | 2370 | -22.920033 | -67.767367 |
| GD2039 | *P. vaccarum* | Chile, Antofagasta, San Pedro de Atacama, Ruta B-241 km 117 | 2370 | -22.920033 | -67.767367 |
| GD2041 | *P. vaccarum* | Chile, Antofagasta, San Pedro de Atacama, Ruta B-245 km 20 | 3240 | -23.01825 | -68.136251 |
| GD2042 | *P. vaccarum* | Chile, Antofagasta, San Pedro de Atacama, Ruta B-245 km 20 | 3240 | -23.01825 | -68.136251 |
| GD2043 | *P. vaccarum* | Chile, Antofagasta, San Pedro de Atacama, Ruta B-245 km 20 | 3240 | -23.01825 | -68.136251 |
| GD2050 | *P. vaccarum* | Chile, Antofagasta, San Pedro de Atacama, Ruta 27 CH km 33 | 4099 | -22.77811 | -68.076983 |
| GD2052 | *P. vaccarum* | Chile, Antofagasta, San Pedro de Atacama, Ruta 27 CH km 45.5 | 4750 | -22.925717 | -67.87305 |
| GD2053 | *P. vaccarum* | Chile, Antofagasta, San Pedro de Atacama, Ruta 27 CH km 45.5 | 4750 | -22.925717 | -67.87305 |
| GD2054 | *P. vaccarum* | Chile, Antofagasta, San Pedro de Atacama, Ruta B-245 km 20 | 3240 | -23.01825 | -68.136251 |
| GD2055 | *P. vaccarum* | Chile, Antofagasta, San Pedro de Atacama, Ruta B-245 km 20 | 3240 | -23.01825 | -68.136251 |
| GD2056 | *P. vaccarum* | Chile, Antofagasta, San Pedro de Atacama, Ruta B-245 km 20 | 3240 | -23.01825 | -68.136251 |
| GD2057 | *P. vaccarum* | Chile, Antofagasta, San Pedro de Atacama, Ruta B-245 km 20 | 3240 | -23.01825 | -68.136251 |
| GD2058 | *P. vaccarum* | Chile, Antofagasta, San Pedro de Atacama, Ruta B-245 km 20 | 3240 | -23.01825 | -68.136251 |
| GD2059 | *P. vaccarum* | Chile, Antofagasta, San Pedro de Atacama, Ruta 27 CH km 45.5 | 4750 | -22.925717 | -67.87305 |
| GD2060 | *P. vaccarum* | Chile, Antofagasta, San Pedro de Atacama, Ruta 27 CH km 45.5 | 4750 | -22.925717 | -67.87305 |
| GD2061 | *P. vaccarum* | Chile, Antofagasta, San Pedro de Atacama, Ruta 27 CH km 45.5 | 4750 | -22.925717 | -67.87305 |
| GD2068 | *P. vaccarum* | Chile, Antofagasta, San Pedro de Atacama, Ruta 27 CH km 33 | 4099 | -22.77811 | -68.076983 |
| GD2069 | *P. vaccarum* | Chile, Antofagasta, San Pedro de Atacama, Ruta 27 CH km 33 | 4099 | -22.77811 | -68.076983 |
| GD2070 | *P. vaccarum* | Chile, Antofagasta, San Pedro de Atacama, Ruta 27 CH km 33 | 4099 | -22.77811 | -68.076983 |
| GD2071 | *P. vaccarum* | Chile, Antofagasta, San Pedro de Atacama, Ruta 27 CH km 33 | 4099 | -22.77811 | -68.076983 |
| GD2072 | *P. vaccarum* | Chile, Antofagasta, San Pedro de Atacama, Ruta 27 CH km 33 | 4099 | -22.77811 | -68.076983 |
| GD2077 | *P. vaccarum* | Chile, Antofagasta, Antofagasta, Parque Nacional Llullaillaco, Refugio Aguadas de Zorritas | 4150 | -24.619333 | -68.588751 |
| GD2078 | *P. vaccarum* | Chile, Antofagasta, Antofagasta, Parque Nacional Llullaillaco, Refugio Aguadas de Zorritas | 4150 | -24.619333 | -68.588751 |

| | | | | | |
|---|---|---|---|---|---|
| GD2079 | *P. vaccarum* | Chile, Antofagasta, Parque Nacional Llullaillaco, Refugio Aguadas de Zorritas | 4150 | -24.619333 | -68.588751 |
| GD2082 | *P. vaccarum* | Chile, Antofagasta, Parque Nacional Llullaillaco, 3 km al suroeste del Refugio Aguadas de Zorritas | 4360 | -24.628933 | -68.559667 |
| GD2083 | *P. vaccarum* | Chile, Antofagasta, Parque Nacional Llullaillaco, 3 km al suroeste del Refugio Aguadas de Zorritas | 4360 | -24.628933 | -68.559667 |
| GD2084 | *P. vaccarum* | Chile, Antofagasta, Parque Nacional Llullaillaco, 3 km al suroeste del Refugio Aguadas de Zorritas | 4360 | -24.628933 | -68.559667 |
| GD2085 | *P. vaccarum* | Chile, Antofagasta, Parque Nacional Llullaillaco, 3 km al suroeste del Refugio Aguadas de Zorritas | 4360 | -24.628933 | -68.559667 |
| GD2086 | *P. vaccarum* | Chile, Antofagasta, Parque Nacional Llullaillaco, 3 km al suroeste del Refugio Aguadas de Zorritas | 4360 | -24.628933 | -68.559667 |
| GD2087 | *P. vaccarum* | Chile, Antofagasta, Parque Nacional Llullaillaco, 3 km al suroeste del Refugio Aguadas de Zorritas | 4360 | -24.628933 | -68.559667 |
| GD2088 | *P. vaccarum* | Chile, Antofagasta, Parque Nacional Llullaillaco, 3 km al suroeste del Refugio Aguadas de Zorritas | 4360 | -24.628933 | -68.559667 |
| GD2091 | *P. vaccarum* | Chile, Antofagasta, Parque Nacional Llullaillaco, Refugio Aguadas de Zorritas | 4150 | -24.619333 | -68.588751 |
| GD2093 | *P. vaccarum* | Chile, Antofagasta, PN Llullaillaco, volcán Llullaillaco, campamento base Este | 5070 | -24.730183 | -68.577933 |
| GD2094 | *P. vaccarum* | Chile, Antofagasta, PN Llullaillaco, volcán Llullaillaco, campamento base Este | 5070 | -24.730183 | -68.577933 |
| GD2095 | *P. vaccarum* | Chile, Antofagasta, PN Llullaillaco, volcán Llullaillaco, campamento base Este | 5070 | -24.730183 | -68.577933 |
| GD2096 | *P. vaccarum* | Chile, Antofagasta, PN Llullaillaco, volcán Llullaillaco, campamento base Este | 5070 | -24.730183 | -68.577933 |
| GD2097 | *P. vaccarum* | Chile (Antofagasta, Antofagasta), cumbre del Volcán Llullaillaco | 6739 | -24.720085 | -68.536958 |
| GD2099 | *P. vaccarum* | Chile, Antofagasta, PN Llullaillaco, volcán Llullaillaco, campamento base Este | 5070 | -24.730183 | -68.577933 |
| GD2100 | *P. vaccarum* | Chile, Antofagasta, PN Llullaillaco, volcán Llullaillaco, campamento base Este | 5070 | -24.730183 | -68.577933 |
| GD2101 | *P. vaccarum* | Chile, Antofagasta, PN Llullaillaco, volcán Llullaillaco, campamento base Este | 5070 | -24.730183 | -68.577933 |
| GD2103 | *P. vaccarum* | Chile, Antofagasta, Antofagasta, Parque Nacional Llullaillaco, Volcán Llullaillaco, campamento base – Ruta Norte | 4620 | -24.675583 | -68.580717 |
| GD2104 | *P. vaccarum* | Chile, Antofagasta, Antofagasta, Parque Nacional Llullaillaco, Volcán Llullaillaco, campamento base – Ruta Norte | 4620 | -24.675583 | -68.580717 |
| GD2105 | *P. vaccarum* | Chile, Antofagasta, Antofagasta, Parque Nacional Llullaillaco, Volcán Llullaillaco, campamento base – Ruta Norte | 4620 | -24.675583 | -68.580717 |
| GD2106 | *P. vaccarum* | Chile, Antofagasta, Antofagasta, Parque Nacional Llullaillaco, Volcán Llullaillaco, campamento base – Ruta Norte | 4620 | -24.675583 | -68.580717 |
| GD2107 | *P. vaccarum* | Chile, Antofagasta, Antofagasta, Parque Nacional Llullaillaco, Volcán Llullaillaco, campamento base – Ruta Norte | 4620 | -24.675583 | -68.580717 |
| GD2117 | *P. limatus* | Chile, Tarapacá, Huara, Quebrada de Tarapacá, Quillahuasa | 1440 | -19.915417 | -69.496133 |
| GD2118 | *P. magister* | Chile, Tarapacá, Huara, Quebrada de Tarapacá, Quillahuasa | 1440 | -19.915417 | -69.496133 |
| GD2119 | *P. magister* | Chile, Tarapacá, Huara, Quebrada de Tarapacá, Quillahuasa | 1440 | -19.915417 | -69.496133 |
| GD2120 | *P. limatus* | Chile, Tarapacá, Huara, Quebrada de Tarapacá, Quillahuasa | 1440 | -19.915417 | -69.496133 |
| GD2121 | *P. limatus* | Chile, Tarapacá, Huara, Quebrada de Tarapacá, Quillahuasa | 1440 | -19.915417 | -69.496133 |

| | | | | | |
|---|---|---|---|---|---|
| GD2133 | *P. limatus* | Chile, Tarapacá, Huara, Quebrada de Tarapacá, Huarasiña | 1360 | -19.941331 | -69.530828 |
| GD2134 | *P. limatus* | Chile, Tarapacá, Huara, Quebrada de Tarapacá, Huarasiña | 1360 | -19.941331 | -69.530828 |
| GD2135 | *P. limatus* | Chile, Tarapacá, Huara, Quebrada de Tarapacá, Huarasiña | 1360 | -19.941331 | -69.530828 |
| GD2150 | *P. limatus* | Chile, Arica y Parinacota, Camarones, Quebrada de Camarones, Ruta A-345, km 28 | 790 | -19.002801 | -69.826975 |
| GD2155 | *P. limatus* | Chile, Arica y Parinacota, Camarones, Quebrada de Camarones, Ruta A-345, km 28 | 790 | -19.002801 | -69.826975 |
| GD2174 | *P. limatus* | Chile, Arica y Parinacota, Camarones, Quebrada de Camarones, Ruta A-345, km 20.7 | 650 | -19.009751 | -69.894184 |
| GD2175 | *P. vaccarum* | Chile, Antofagasta, Antofagasta, Reserva Natural La Chimba, Quebrada La Chimba | 420 | -23.537751 | -70.359156 |
| GD2226 | *P. limatus* | Chile, Arica y Parinacota, Huara, Chusmiza, Quebrada de Ocharaza | 3380 | -19.6832679 | -69.1799747 |
| GD2227 | *P. limatus* | Chile, Arica y Parinacota, Huara, Chusmiza, Quebrada de Ocharaza | 3380 | -19.6832679 | -69.1799747 |
| GD2228 | *P. limatus* | Chile, Arica y Parinacota, Huara, Chusmiza, Quebrada de Ocharaza | 3380 | -19.6832679 | -69.1799747 |
| GD2229 | *P. limatus* | Chile, Arica y Parinacota, Huara, Chusmiza, Quebrada de Ocharaza | 3380 | -19.6832679 | -69.1799747 |
| GD2230 | *P. chilensis* | Chile, Arica y Parinacota, Huara, Chusmiza, Quebrada de Ocharaza | 3380 | -19.6832679 | -69.1799747 |
| GD2231 | *P. limatus* | Chile, Arica y Parinacota, Huara, Chusmiza, Quebrada de Ocharaza | 3380 | -19.6832679 | -69.1799747 |
| GD2232 | *P. limatus* | Chile, Arica y Parinacota, Huara, Chusmiza, Quebrada de Ocharaza | 3380 | -19.6832679 | -69.1799747 |
| GD2233 | *P. limatus* | Chile, Arica y Parinacota, Huara, Chusmiza, Quebrada de Ocharaza | 3380 | -19.6832679 | -69.1799747 |
| GD2234 | *P. limatus* | Chile, Arica y Parinacota, Huara, Chusmiza, Quebrada de Ocharaza | 3380 | -19.6832679 | -69.1799747 |
| GD2241 | *P. limatus* | Chile, Arica y Parinacota, Huara, Chusmiza, Quebrada de Ocharaza | 3380 | -19.6832679 | -69.1799747 |
| GD2242 | *P. limatus* | Chile, Arica y Parinacota, Huara, Chusmiza, Quebrada de Ocharaza | 3380 | -19.6832679 | -69.1799747 |
| GD2243 | *P. magister* | Chile, Arica y Parinacota, Huara, Chusmiza, Quebrada de Ocharaza | 3380 | -19.6832679 | -69.1799747 |
| GD2244 | *P. magister* | Chile, Arica y Parinacota, Huara, Chusmiza, Quebrada de Ocharaza | 3380 | -19.6832679 | -69.1799747 |
| GD2245 | *P. limatus* | Chile, Arica y Parinacota, Huara, Chusmiza, Quebrada de Ocharaza | 3380 | -19.6832679 | -69.1799747 |
| GD2251 | *P. chilensis* | Chile, Arica y Parinacota, Laguna Cota Kulco | 4183 | -19.6437208 | -68.759671 |
| GD2252 | *P. chilensis* | Chile, Arica y Parinacota, Laguna Cota Kulco | 4183 | -19.6437208 | -68.759671 |

| | | | | | |
|---|---|---|---|---|---|
| GD2253 | *P. vaccarum* | Chile, Arica y Parinacota, Laguna Cota Kulco | 4183 | -19.6437208 | -68.759671 |
| GD2254 | *P. chilensis* | Chile, Arica y Parinacota, Laguna Cota Kulco | 4183 | -19.6437208 | -68.759671 |
| GD2255 | *P. chilensis* | Chile, Arica y Parinacota, Laguna Cota Kulco | 4183 | -19.6437208 | -68.759671 |
| GD2256 | *P. chilensis* | Chile, Arica y Parinacota, Laguna Cota Kulco | 4183 | -19.6437208 | -68.759671 |
| GD2257 | *P. chilensis* | Chile, Arica y Parinacota, Laguna Cota Kulco | 4183 | -19.6437208 | -68.759671 |
| GD2258 | *P. chilensis* | Chile, Arica y Parinacota, Laguna Cota Kulco | 4183 | -19.6437208 | -68.759671 |
| GD2259 | *P. chilensis* | Chile, Arica y Parinacota, Laguna Cota Kulco | 4183 | -19.6437208 | -68.759671 |
| GD2260 | *P. chilensis* | Chile, Arica y Parinacota, Laguna Cota Kulco | 4183 | -19.6437208 | -68.759671 |
| GD2261 | *P. chilensis* | Chile, Arica y Parinacota, Laguna Cota Kulco | 4183 | -19.6437208 | -68.759671 |
| GD2285 | *P. chilensis* | Chile, Arica y Parinacota, campamento Laguna Casiri | 4807 | -18.07086 | -69.08162 |
| GD2287 | *P. vaccarum* | Chile, Antofagasta, Tocopilla, Cobija | 49 | -22.52203 | -70.24418 |
| GD2289 | *P. vaccarum* | Chile, Antofagasta, Tocopilla, Cobija | 49 | -22.52203 | -70.24418 |
| GD2290 | *P. vaccarum* | Chile, Antofagasta, Tocopilla, Cobija | 49 | -22.52203 | -70.24418 |
| GD2291 | *P. vaccarum* | Chile, Antofagasta, Tocopilla, Cobija | 49 | -22.52203 | -70.24418 |
| GD2347 | *P. magister* | Chile, Antofagasta, María Elena, Rio Loa, Paso Toco (Puente Teresa) | 1100 | -21.96901 | -69.57404 |
| GD2348 | *P. magister* | Chile, Antofagasta, María Elena, Rio Loa | 974 | -21.95561 | -69.56358 |
| GD2349 | *P. magister* | Chile, Antofagasta, María Elena, Rio Loa | 974 | -21.95561 | -69.56358 |
| GD2350 | *P. limatus* | Chile, Antofagasta, María Elena, Rio Loa | 974 | -21.95561 | -69.56358 |
| GD2351 | *P. vaccarum* | Chile, Antofagasta, María Elena, Rio Loa | 974 | -21.95561 | -69.56358 |
| MCQ343 | *P. vaccarum* | Chile, Atacama, Copiapó, Parque Nacional Nevado de Tres Cruces, cerca de Laguna Santa Rosa | 3860 | -27.0961072 | -69.2035758 |
| MCQ344 | *P. vaccarum* | Chile, Atacama, Copiapó, Parque Nacional Nevado de Tres Cruces, cerca de Laguna Santa Rosa | 3861 | -27.0961072 | -69.2035758 |
| MCQ345 | *P. vaccarum* | Chile, Atacama, Copiapó, Parque Nacional Nevado de Tres Cruces, cerca de Laguna Santa Rosa | 3862 | -27.0961072 | -69.2035758 |
| MCQ346 | *P. vaccarum* | Chile, Atacama, Copiapó, Parque Nacional Nevado de Tres Cruces, cerca de Laguna Santa Rosa | 3863 | -27.0961072 | -69.2035758 |
| MCQ347 | *P. vaccarum* | Chile, Atacama, Copiapó, Parque Nacional Nevado de Tres Cruces, cerca de Laguna Santa Rosa | 3864 | -27.0961072 | -69.2035758 |

| | | | | | |
|---|---|---|---|---|---|
| MCQ350 | *P. vaccarum* | Chile, Atacama, Copiapó, Parque Nacional Nevado de Tres Cruces, cerca de Laguna Santa Rosa | 3867 | -27.0961072 | -69.2035758 |
| MCQ352 | *P. vaccarum* | Chile, Atacama, Copiapó, Parque Nacional Nevado de Tres Cruces, Refugio Maricunga | 3780 | -27.0806021 | -69.1756545 |
| MCQ353 | *P. vaccarum* | Chile, Atacama, Copiapó, Parque Nacional Nevado de Tres Cruces, cerca de Laguna Santa Rosa | 3868 | -27.0961072 | -69.2035758 |
| MCQ354 | *P. vaccarum* | Chile, Atacama, Copiapó, Parque Nacional Nevado de Tres Cruces, cerca de Laguna Santa Rosa | 3868 | -27.0961072 | -69.2035758 |
| MCQ355 | *P. vaccarum* | Chile, Atacama, Copiapó, Parque Nacional Nevado de Tres Cruces, roquerio frente a Laguna Santa Rosa | 3780 | -27.0902708 | -69.1748259 |
| MCQ357 | *P. vaccarum* | Chile, Atacama, Copiapó, Refugio Laguna Verde | 4460 | -26.8903846 | -68.4859815 |
| MCQ358 | *P. vaccarum* | Chile, Atacama, Copiapó, Refugio Laguna Verde | 4460 | -26.8903846 | -68.4859815 |
| MCQ359 | *P. vaccarum* | Chile, Atacama, Copiapó, Ojos del Salado, campamento base, Refugio Atacama | 5250 | -27.0597838 | -68.5476175 |
| MCQ360 | *P. vaccarum* | Chile, Atacama, Copiapó, Ojos del Salado, campamento base, Refugio Atacama | 5250 | -27.0597838 | -68.5476175 |
| MCQ361 | *P. vaccarum* | Chile, Atacama, Copiapó, Ojos del Salado, campamento base, Refugio Atacama | 5250 | -27.0597838 | -68.5476175 |
| MCQ362 | *P. vaccarum* | Chile, Atacama, Copiapó, Ojos del Salado, campamento base, Refugio Atacama | 5250 | -27.0597838 | -68.5476175 |
| MCQ364 | *P. vaccarum* | Chile, Atacama, Copiapó, Ojos del Salado, campamento base, Refugio Atacama | 5250 | -27.0597838 | -68.5476175 |
| MCQ365 | *P. vaccarum* | Chile, Atacama, Copiapó, Ojos del Salado, campamento base, Refugio Atacama | 5250 | -27.0597838 | -68.5476175 |
| MQC 478 | *P. chilensis* | Bolivia, Oruro, Volcán Acotango, campamento base | 5027 | -18.366389 | -69.024167 |
| MQC 482 | *P. chilensis* | Bolivia, Oruro, Sajama, mirador Monte Cielo | 4537 | -18.135556 | -68.954722 |
| MQC 483 | *P. chilensis* | Bolivia, Oruro, Sajama, mirador Monte Cielo | 4537 | -18.135556 | -68.954722 |
| MQC 484 | *P. chilensis* | Bolivia, Oruro, Sajama, mirador Monte Cielo | 4537 | -18.135556 | -68.954722 |
| MQC 485 | *P. chilensis* | Bolivia, Oruro, Sajama, mirador Monte Cielo | 4537 | -18.135556 | -68.954722 |
| MQC 486 | *P. chilensis* | Bolivia, Oruro, Sajama, mirador Monte Cielo | 4537 | -18.135556 | -68.954722 |
| MQC 487 | *P. chilensis* | Bolivia, Oruro, Sajama, mirador Monte Cielo | 4537 | -18.135556 | -68.954722 |
| MQC 488 | *P. chilensis* | Bolivia, Oruro, Sajama, mirador Monte Cielo | 4537 | -18.135556 | -68.954722 |
| MQC 496 | *P. chilensis* | Bolivia, Oruro, Sajama, mirador Monte Cielo | 4537 | -18.135556 | -68.954722 |
| MQC 497 | *P. chilensis* | Bolivia, Oruro, Sajama, mirador Monte Cielo | 4537 | -18.135556 | -68.954722 |
| MQC 499 | *P. chilensis* | Bolivia, Oruro, Geiser Sajama, Sitio 1 | 4420 | -18.0961169 | -69.03215 |

| | | | | | |
|---|---|---|---|---|---|
| MQC 500 | *P. chilensis* | Bolivia, Oruro, Geiser Sajama, Sitio 3 | 4330 | -18.110246 | -69.0111762 |
| MQC 503 | *P. chilensis* | Bolivia, Oruro, Volcán Parinacota, campamento alto | 5221 | -18.152194 | -69.122333 |
| MQC 504 | *P. chilensis* | Bolivia, Oruro, Volcán Parinacota, campamento alto | 5221 | -18.152194 | -69.122333 |
| MQC 510 | *P. chilensis* | Bolivia, Oruro, Geiser Sajama, Sitio 1 | 4420 | -18.0961169 | -69.03215 |
| MQC 512 | *P. chilensis* | Bolivia, Oruro, Geiser Sajama, Sitio 1 | 4420 | -18.0961169 | -69.03215 |
| MQC 513 | *P. chilensis* | Bolivia, Oruro, Geiser Sajama, Sitio 1 | 4420 | -18.0961169 | -69.03215 |
| MQC 514 | *P. chilensis* | Bolivia, Oruro, Geiser Sajama, Sitio 2 | 4362 | -18.1056875 | -69.0161179 |
| MQC368 | *P. vaccarum* | Chile, Antofagasta, San Pedro de Atacama, Gachi, Sector Guatín, Margen Río Puritama | 3120 | -22.78145 | -68.10543 |
| MQC369 | *P. vaccarum* | Chile, Antofagasta, San Pedro de Atacama, Gachi, Sector Guatín, Margen Río Puritama | 3120 | -22.78145 | -68.10543 |
| MQC371 | *P. chilensis* | Chile, Antofagasta, Ollagüe, Volcán Aucanquilcha, Linea 4 | 4543 | -21.18773 | -68.40397 |
| MQC372 | *P. chilensis* | Chile, Antofagasta, Ollagüe, Volcán Aucanquilcha, Linea 4 | 4543 | -21.18773 | -68.40397 |
| MQC373 | *P. chilensis* | Chile, Antofagasta, Ollagüe, Volcán Aucanquilcha, Linea 5 | 4510 | -21.18774 | -68.40162 |
| MQC374 | *P. chilensis* | Chile, Antofagasta, Ollagüe, Volcán Aucanquilcha, Linea 5 | 4510 | -21.18774 | -68.40162 |
| MQC375 | *P. chilensis* | Chile, Antofagasta, Ollagüe, Volcán Aucanquilcha, Linea 5 | 4510 | -21.18774 | -68.40162 |
| MQC376 | *P. chilensis* | Chile, Antofagasta, Ollagüe, Volcán Aucanquilcha, Linea 4 | 4543 | -21.18773 | -68.40397 |
| MQC377 | *P. chilensis* | Chile, Antofagasta, Ollagüe, Volcán Aucanquilcha, Linea 5 | 4510 | -21.18774 | -68.40162 |
| MQC378 | *P. chilensis* | Chile, Antofagasta, Ollagüe, Volcán Aucanquilcha, Linea 4 | 4543 | -21.18773 | -68.40397 |
| MQC379 | *P. chilensis* | Chile, Antofagasta, Ollagüe, Volcán Aucanquilcha, Linea 5 | 4510 | -21.18774 | -68.40162 |
| MQC382 | *P. vaccarum* | Chile, Antofagasta, San Pedro de Atacama, Volcán Acamarachi, campamento base | 4895 | -23.27625 | -67.60355 |
| MQC383 | *P. vaccarum* | Chile, Antofagasta, San Pedro de Atacama, Volcán Acamarachi, campamento base | 4895 | -23.27625 | -67.60355 |
| MQC386 | *P. vaccarum* | Chile, Antofagasta, San Pedro de Atacama, Volcán Acamarachi, campamento base | 4895 | -23.27625 | -67.60355 |
| MQC390 | *P. vaccarum* | Chile, Antofagasta, San Pedro de Atacama, Salar de Púlar, Volcán Púlar | 3651 | -24.23276 | -67.94800 |
| MQC391 | *P. vaccarum* | Chile, Antofagasta, San Pedro de Atacama, Salar de Púlar, Volcán Púlar | 3651 | -24.23276 | -67.94800 |
| MQC392 | *P. vaccarum* | Chile, Antofagasta, San Pedro de Atacama, Salar de Púlar, Volcán Púlar | 3651 | -24.23276 | -67.94800 |
| MQC393 | *P. vaccarum* | Chile, Antofagasta, San Pedro de Atacama, Salar de Púlar, Volcán Púlar | 3651 | -24.23276 | -67.94800 |
| MQC394 | *P. vaccarum* | Chile, Antofagasta, San Pedro de Atacama, Salar de Púlar, Volcán Púlar | 3651 | -24.23276 | -67.94800 |
| MQC395 | *P. vaccarum* | Chile, Antofagasta, San Pedro de Atacama, Salar de Púlar, Volcán Púlar | 3651 | -24.23276 | -67.94800 |
| MQC396 | *P. vaccarum* | Chile, Antofagasta, San Pedro de Atacama, Salar de Púlar, Volcán Púlar | 3651 | -24.23276 | -67.94800 |

| | | | | | |
|---|---|---|---|---|---|
| MQC400 | *P. vaccarum* | Chile, Antofagasta, San Pedro de Atacama, Salar de Púlar, Volcán Púlar | 3651 | -24.23276 | -67.94800 |
| MQC401 | *P. vaccarum* | Chile, Antofagasta, San Pedro de Atacama, Salar de Púlar, Volcán Púlar | 3651 | -24.23276 | -67.94800 |
| MQC402 | *P. vaccarum* | Chile, Antofagasta, San Pedro de Atacama, Salar de Púlar, Volcán Púlar | 3651 | -24.23276 | -67.94800 |
| MQC403 | *P. vaccarum* | Chile (Antofagasta, San Pedro de Atacama), cumbre de Volcán Salín | 4016 | -24.32839 | -68.06729 |
| MQC414 | *P. vaccarum* | Chile, Antofagasta, San Pedro de Atacama, Salar de Púlar, Volcán Púlar | 3651 | -24.23276 | -67.94800 |
| MQC415 | *P. vaccarum* | Chile, Antofagasta, San Pedro de Atacama, Salar de Púlar, Volcán Púlar | 3651 | -24.23276 | -67.94800 |
| MQC418 | *P. vaccarum* | Chile, Antofagasta, San Pedro de Atacama, Salar de Púlar, Volcán Púlar | 3651 | -24.23276 | -67.94800 |
| MQC419 | *P. vaccarum* | Chile, Antofagasta, San Pedro de Atacama, Salar de Púlar, Volcán Púlar | 3651 | -24.23276 | -67.94800 |
| MQC420 | *P. vaccarum* | Chile, Antofagasta, San Pedro de Atacama, Salar de Púlar, Volcán Púlar | 3651 | -24.23276 | -67.94800 |
| MQC421 | *P. vaccarum* | Chile, Antofagasta, San Pedro de Atacama, Salar de Púlar, Volcán Púlar | 3651 | -24.23276 | -67.94800 |
| MQC422 | *P. vaccarum* | Chile, Antofagasta, San Pedro de Atacama, Salar de Púlar, Volcán Púlar | 3651 | -24.23276 | -67.94800 |
| MQC423 | *P. vaccarum* | Chile, Antofagasta, San Pedro de Atacama, Salar de Púlar, Volcán Púlar | 3651 | -24.23276 | -67.94800 |
| MQC425 | *P. chilensis* | Chile, Antofagasta, San Pedro de Atacama, base del Cerro Sairecabur | 4480 | -22.72710 | -67.88941 |
| MQC426 | *P. chilensis* | Chile, Antofagasta, Ollagüe, Cerro Colorado, campamento base | 4840 | -22.58423 | -67.90534 |
| MQC427 | *P. chilensis* | Chile, Antofagasta, Ollagüe, Cerro Colorado, campamento base | 4840 | -22.58423 | -67.90534 |
| MQC429 | *P. vaccarum* | Chile, Antofagasta, Antofagasta, Salar de Aguas Calientes | 3750 | -24.9760014 | -68.625507 |
| MQC430 | *P. vaccarum* | Chile, Antofagasta, Antofagasta, Salar de Aguas Calientes | 3750 | -24.9760014 | -68.625507 |
| MQC431 | *P. vaccarum* | Chile, Antofagasta, Antofagasta, Salar de Aguas Calientes | 3750 | -24.9760014 | -68.625507 |
| MQC432 | *P. vaccarum* | Chile, Antofagasta, Antofagasta, Salar de Aguas Calientes | 3750 | -24.9760014 | -68.625507 |
| MQC433 | *P. vaccarum* | Chile, Antofagasta, Antofagasta, Salar de Aguas Calientes | 3750 | -24.9760014 | -68.625507 |
| MQC434 | *P. vaccarum* | Chile, Antofagasta, Antofagasta, Salar de Aguas Calientes | 3750 | -24.9760014 | -68.625507 |
| MQC437 | *P. vaccarum* | Chile, Antofagasta, Antofagasta, Salar de Aguas Calientes | 3750 | -24.9760014 | -68.625507 |
| MQC438 | *P. vaccarum* | Chile, Antofagasta, Antofagasta, Salar de Aguas Calientes | 3750 | -24.9760014 | -68.625507 |
| MQC439 | *P. vaccarum* | Chile, Antofagasta, Antofagasta, Salar de Aguas Calientes | 3750 | -24.9760014 | -68.625507 |
| MQC440 | *P. vaccarum* | Chile, Atacama, Copiapó, campamento base, Volcán Copiapó | 4100 | -27.2035206 | -69.0578724 |

| | | | | | |
|---|---|---|---|---|---|
| MQC441 | *P. vaccarum* | Chile, Atacama, Copiapó, campamento base, Volcán Copiapó | 4100 | -27.2035206 | -69.0578724 |
| MQC443 | *P. vaccarum* | Chile, Atacama, Copiapó, Refugio Laguna Verde | 4460 | -26.8903846 | -68.4859815 |
| MQC451 | *P. vaccarum* | Chile, Atacama, Copiapó, Vallecitos, Pajas Grandes (Colas de Zorro) | 3100 | -27.0661337 | -69.333953 |
| MQC452 | *P. vaccarum* | Chile, Atacama, Copiapó, Vallecitos, Pajas Grandes (Colas de Zorro) | 3100 | -27.0661337 | -69.333953 |
| MQC453 | *P. vaccarum* | Chile, Atacama, Copiapó, Vallecitos, Pajas Grandes (Colas de Zorro) | 3100 | -27.0661337 | -69.333953 |
| MQC454 | *P. vaccarum* | Chile, Atacama, Copiapó, Vallecitos, Pajas Grandes (Colas de Zorro) | 3100 | -27.0661337 | -69.333953 |
| MQC455 | *P. vaccarum* | Chile, Atacama, Copiapó, Vallecitos, Pajas Grandes (Colas de Zorro) | 3100 | -27.0661337 | -69.333953 |
| MQC465 | *P. vaccarum* | Chile, Atacama, Copiapó, Vallecitos, Pajas Grandes (Colas de Zorro) | 3100 | -27.0661337 | -69.333953 |
| MQC466 | *P. vaccarum* | Chile, Atacama, Copiapó, Vallecitos, Pajas Grandes (Colas de Zorro) | 3100 | -27.0661337 | -69.333953 |
| MQC467 | *P. vaccarum* | Chile, Atacama, Copiapó, Vallecitos, Pajas Grandes (Colas de Zorro) | 3100 | -27.0661337 | -69.333953 |
| MQC468 | *P. vaccarum* | Chile, Atacama, Copiapó, Vallecitos, Pajas Grandes (Colas de Zorro) | 3100 | -27.0661337 | -69.333953 |
| MQC469 | *P. vaccarum* | Chile, Atacama, Copiapó, Vallecitos, Pajas Grandes (Colas de Zorro) | 3100 | -27.0661337 | -69.333953 |
| MQC470 | *P. vaccarum* | Chile, Atacama, Copiapó, Vallecitos, Pajas Grandes (Colas de Zorro) | 3100 | -27.0661337 | -69.333953 |
| MQC471 | *P. vaccarum* | Chile, Atacama, Copiapó, Vallecitos, Pajas Grandes (Colas de Zorro) | 3100 | -27.0661337 | -69.333953 |
| MQC524 | *P. vaccarum* | Chile, Antofagasta, Antofagasta, Parque Nacional Llullaillaco, Refugio Aguadas de Zorritas | 4150 | -24.619333 | -68.588751 |
| MQC526 | *P. vaccarum* | Chile, Antofagasta, Antofagasta, Parque Nacional Llullaillaco, Refugio Aguadas de Zorritas | 4150 | -24.619333 | -68.588751 |
| MQC528 | *P. vaccarum* | Chile, Antofagasta, Antofagasta, Parque Nacional Llullaillaco, Refugio Aguadas de Zorritas | 4150 | -24.619333 | -68.588751 |
| MQC529 | *P. vaccarum* | Chile, Antofagasta, Antofagasta, Parque Nacional Llullaillaco, Refugio Aguadas de Zorritas | 4150 | -24.619333 | -68.588751 |
| MQC530 | *P. vaccarum* | Chile, Antofagasta, Antofagasta, Parque Nacional Llullaillaco, Refugio Aguadas de Zorritas | 4150 | -24.619333 | -68.588751 |
| MQC531 | *P. vaccarum* | Chile, Antofagasta, Antofagasta, Parque Nacional Llullaillaco, Refugio Aguadas de Zorritas | 4150 | -24.619333 | -68.588751 |
| MQC532 | *P. vaccarum* | Chile, Antofagasta, Antofagasta, Parque Nacional Llullaillaco, Refugio Aguadas de Zorritas | 4150 | -24.619333 | -68.588751 |
| MQC534 | *P. vaccarum* | Chile, Antofagasta, Antofagasta, Parque Nacional Llullaillaco, Refugio Aguadas de Zorritas | 4150 | -24.619333 | -68.588751 |
| MQC574 | *P. vaccarum* | Chile, Metropolitana, San Jose de Maipo, Camino Las Melosas, Sector Quebrada el Loro | 1430 | -33.8411641 | -70.2133673 |

| | | | | | |
|---|---|---|---|---|---|
| MQC576 | *P. vaccarum* | Chile, Metropolitana, San Jose de Maipo, Camino Las Melosas, Sector Quebrada el Loro | 1430 | -33.8411641 | -70.2133673 |
| MQC578 | *P. vaccarum* | Chile, Metropolitana, San Jose de Maipo, Camino Las Melosas, Sector Quebrada el Loro | 1430 | -33.8411641 | -70.2133673 |
| MQC582 | *P. vaccarum* | Chile, Metropolitana, San Jose de Maipo, Camino Las Melosas, Sector Quebrada el Loro | 1430 | -33.8411641 | -70.2133673 |
| MQC583 | *P. vaccarum* | Chile, Metropolitana, San Jose de Maipo, Camino Las Melosas, Sector Quebrada el Loro | 1430 | -33.8411641 | -70.2133673 |
| MQC585 | *P. vaccarum* | Chile, Metropolitana, San Jose de Maipo, Camino Las Melosas, Sector Quebrada el Loro | 1430 | -33.8411641 | -70.2133673 |
| MQC586 | *P. vaccarum* | Chile, Metropolitana, San Jose de Maipo, Camino Las Melosas, Sector Quebrada el Loro | 1430 | -33.8411641 | -70.2133673 |
| MQC587 | *P. vaccarum* | Chile, Metropolitana, San Jose de Maipo, Camino Las Melosas, Sector Quebrada el Loro | 1430 | -33.8411641 | -70.2133673 |
| MQC589 | *P. vaccarum* | Chile, Metropolitana, San Jose de Maipo, Camino Las Melosas, Sector Quebrada el Loro | 1430 | -33.8411641 | -70.2133673 |
| MQC645 | *P. vaccarum* | Chile, Antofagasta, Antofagasta, Parque Nacional Llullaillaco, Camino hacia el Refugio Aguadas de Zorritas | 4260 | -24.6326782 | -68.5821795 |
| MQC647 | *P. vaccarum* | Chile, Antofagasta, Antofagasta, Parque Nacional Llullaillaco, Camino hacia el Refugio Aguadas de Zorritas | 4260 | -24.6326782 | -68.5821795 |
| MQC649 | *P. vaccarum* | Chile, Antofagasta, Antofagasta, Parque Nacional Llullaillaco, Camino hacia el Refugio Aguadas de Zorritas | 4260 | -24.6326782 | -68.5821795 |
| PZ784 | *P. chilensis* | Bolivia, Oruro, Geiser Sajama | 4420 | -18.0961169 | -69.03215 |
| PZ797 | *P. chilensis* | Bolivia, Oruro, Volcán Sajama, campamento base | 4880 | -18.112778 | -68.915556 |
| PZ798 | *P. chilensis* | Bolivia, Oruro, Volcán Sajama, campamento base | 4880 | -18.112778 | -68.915556 |
| PZ802 | *P. chilensis* | Bolivia, Oruro, Volcán Sajama, campamento base | 4880 | -18.112778 | -68.915556 |

**Table S2.** Sequence quality metrics, mapping, and assignment rates for right ventricle transcriptomic data.

| ***P. vaccarum*** **right ventricle (*n* = 10)** | |
|---|---|
| **Mean Quality Score** | |
| Min | 18.2 |
| Max | 25.9 |
| mean (SD) | 22.95 ± 2.61 |
| **Percent Bases Above Q30** | |
| Min | 93.2 |
| Median | 93.45 |
| Max | 93.9 |
| mean (SD) | 93.49 ± 0.21 |
| **GC (%)** | |
| Min | 45.6 |
| Median | 47.1 |
| Max | 48 |
| mean (SD) | 46.89 ± 0.71 |
| **Reads** | |
| Min | 41571970 |
| Median | 43408610.5 |
| Max | 57039685 |
| mean (SD) | 45,945,408.10 ±5,347,994.38 |
| **Genome Alignment Rate (%)** | |
| Min | 85.9 |
| Max | 90.7 |
| mean (SD) | 89.23 ± 1.57 |
| **Assignment Rate (%)** | |
| Min | 26.1 |
| Max | 40.4 |
| mean (SD) | 35.88 ± 4.03 |

**Table S3.** Differentially expressed genes (FDR ≤ 0.1) in the RV transcriptomes of *P. vaccarum* sampled from high-elevation localities on the flanks of Ojos del Salado and Llullaillaco. For the DE analysis, we used Llullaillaco as the reference population so that a logFC > 0 is an increase in expression in Ojos mice and a log FC < 0 is an increase in expression in Llullaillaco mice.

| Gene_ID | logFC | logCPM | F | PValue | FDR | name | description |
|---|---|---|---|---|---|---|---|
| ENSMUSG00000036052 | 0.978930766 | 6.477372989 | 54.68677407 | 3.40E-06 | 0.019651834 | Dnajb5 | DnaJ heat shock protein family (Hsp40) member B5 |
| ENSMUSG00000038168 | 4.474282289 | 8.543735411 | 54.45227095 | 3.48E-06 | 0.019651834 | P3h2 | prolyl 3-hydroxylase 2 |
| ENSMUSG00000032511 | 1.373898018 | 6.260692005 | 48.55610681 | 6.60E-06 | 0.020527136 | Scn5a | sodium channel, voltage-gated, type V, alpha |
| ENSMUSG00000039956 | -1.950885373 | 6.465371731 | 46.46513073 | 8.41E-06 | 0.020527136 | Mrap | melanocortin 2 receptor accessory protein |
| ENSMUSG00000003934 | 2.35933555 | 2.891483105 | 46.15795954 | 9.09E-06 | 0.020527136 | Efnb3 | ephrin B3 |
| ENSMUSG00000074736 | 3.751551353 | 1.342065463 | 34.58767189 | 1.22E-05 | 0.020727624 | Syndig1 | synapse differentiation inducing 1 |
| ENSMUSG00000026548 | -6.661949487 | 3.722200839 | 32.43581113 | 1.28E-05 | 0.020727624 | Slamf9 | SLAM family member 9 |
| ENSMUSG00000028444 | 2.335069978 | 3.183047273 | 41.82371443 | 1.53E-05 | 0.02161139 | Cntfr | ciliary neurotrophic factor receptor |
| ENSMUSG00000024063 | 0.85729584 | 6.443066741 | 37.49748642 | 2.64E-05 | 0.031412731 | Lbh | limb-bud and heart |
| ENSMUSG00000032177 | 1.265495417 | 3.878179931 | 36.1579481 | 3.20E-05 | 0.031412731 | Pde4a | phosphodiesterase 4A, cAMP specific |
| ENSMUSG00000034872 | 1.947315054 | 3.218362501 | 35.88906135 | 3.37E-05 | 0.031412731 | Gipc3 | GIPC PDZ domain containing family, member 3 |
| ENSMUSG00000027861 | 0.911263336 | 8.989615642 | 35.69585677 | 3.41E-05 | 0.031412731 | Casq2 | calsequestrin 2 |
| ENSMUSG00000032834 | 1.174318022 | 5.64390783 | 35.28365998 | 3.62E-05 | 0.031412731 | Pwp2 | PWP2 periodic tryptophan protein homolog (yeast) |
| ENSMUSG00000017817 | 1.125877127 | 7.019003348 | 34.46672981 | 4.08E-05 | 0.031412731 | Jph2 | junctophilin 2 |
| ENSMUSG00000040147 | 1.033910886 | 6.707152058 | 34.32136499 | 4.17E-05 | 0.031412731 | Maob | monoamine oxidase B |
| ENSMUSG00000046329 | 1.297786593 | 4.493439569 | 33.60698217 | 4.65E-05 | 0.032805011 | Slc25a23 | solute carrier family 25 (mitochondrial carrier; phosphate carrier), member 23 |
| ENSMUSG00000039376 | 1.729661301 | 5.054644981 | 32.69710142 | 5.34E-05 | 0.033989129 | Synpo2l | synaptopodin 2-like |
| ENSMUSG00000060913 | 0.912092712 | 6.494773806 | 32.27978619 | 5.69E-05 | 0.033989129 | Trim55 | tripartite motif-containing 55 |
| ENSMUSG00000015605 | 1.0358789 | 5.508510302 | 32.12563284 | 5.83E-05 | 0.033989129 | Srf | serum response factor |
| ENSMUSG00000059852 | 0.900785942 | 6.78038287 | 31.60679753 | 6.32E-05 | 0.033989129 | Kcng2 | potassium voltage-gated channel, subfamily G, member 2 |
| ENSMUSG00000042828 | 0.969111004 | 6.736694168 | 31.60351462 | 6.32E-05 | 0.033989129 | Trim72 | tripartite motif-containing 72 |
| ENSMUSG00000001349 | 2.295764881 | 2.968461276 | 31.34363335 | 6.76E-05 | 0.034725999 | Cnn1 | calponin 1 |
| ENSMUSG00000038663 | 1.806030457 | 5.104975406 | 30.42713922 | 7.63E-05 | 0.037469661 | Fsd2 | fibronectin type III and SPRY domain containing 2 |
| ENSMUSG00000050953 | 1.030880392 | 7.757647264 | 29.65395359 | 8.65E-05 | 0.040709139 | Gja1 | gap junction protein, alpha 1 |
| ENSMUSG00000040345 | -1.318804127 | 3.316296186 | 28.45742534 | 0.000106258 | 0.044380504 | Arhgap9 | Rho GTPase activating protein 9 |
| ENSMUSG00000051652 | 2.149480824 | 2.880695836 | 28.48343572 | 0.00010779 | 0.044380504 | Lrrc3 | leucine rich repeat containing 3 |
| ENSMUSG00000006221 | 1.169469593 | 10.19302869 | 28.24471507 | 0.000109555 | 0.044380504 | Hspb7 | heat shock protein family, member 7 (cardiovascular) |
| ENSMUSG00000096727 | -1.54688559 | 5.790139315 | 28.1912964 | 0.000110591 | 0.044380504 | Psmb9 | proteasome (prosome, macropain) subunit, beta type 9 (large multifunctional peptidase 2) |

| | | | | | | | |
|---|---|---|---|---|---|---|---|
| ENSMUSG00000015852 | -1.4565597 | 5.048136715 | 28.01781128 | 0.000113957 | 0.044380504 | Fcrl2 | Fc receptor like 2 |
| ENSMUSG00000023078 | -6.769497736 | 5.332218266 | 26.61421874 | 0.000124242 | 0.046772796 | Cxcl13 | C-X-C motif chemokine ligand 13 |
| ENSMUSG00000070044 | 1.034967615 | 4.464525858 | 26.78312621 | 0.000141331 | 0.051147496 | Fam149a | family with sequence similarity 149, member A |
| ENSMUSG00000039005 | -1.517582755 | 2.667635532 | 26.73952319 | 0.000144919 | 0.051147496 | Tlr4 | toll-like receptor 4 |
| ENSMUSG00000030352 | 1.636531029 | 4.079217548 | 25.99904308 | 0.000162775 | 0.054658097 | Tspan9 | tetraspanin 9 |
| ENSMUSG00000051351 | 1.675339755 | 2.014597603 | 25.7949843 | 0.00017319 | 0.054658097 | Zfp46 | zinc finger protein 46 |
| ENSMUSG00000090733 | -0.69166234 | 9.318033119 | 25.5021151 | 0.000177791 | 0.054658097 | Rps27 | ribosomal protein S27 |
| ENSMUSG00000023132 | -3.150569993 | 2.50126331 | 25.40615666 | 0.000179282 | 0.054658097 | Gzma | granzyme A |
| ENSMUSG00000044788 | 2.606281719 | 3.424392387 | 25.44667406 | 0.000183064 | 0.054658097 | Fads6 | fatty acid desaturase domain family, member 6 |
| ENSMUSG00000049436 | -2.652669875 | 3.395110414 | 25.31150066 | 0.000187595 | 0.054658097 | Upk1b | uroplakin 1B |
| ENSMUSG00000046879 | -1.620980488 | 4.239282438 | 25.09996934 | 0.000191758 | 0.054658097 | Irgm1 | immunity-related GTPase family M member 1 |
| ENSMUSG00000023951 | 1.071672384 | 8.270448333 | 25.03965299 | 0.000193583 | 0.054658097 | Vegfa | vascular endothelial growth factor A |
| ENSMUSG00000031378 | 0.793154985 | 4.838417146 | 24.59661751 | 0.000210332 | 0.056024394 | Abcd1 | ATP-binding cassette, sub-family D member 1 |
| ENSMUSG00000033039 | 0.722413469 | 6.80065141 | 24.44175012 | 0.00021645 | 0.056024394 | Micall1 | microtubule associated monooxygenase, calponin and LIM domain containing -like 1 |
| ENSMUSG00000049410 | -2.972282368 | 3.289104655 | 24.47106113 | 0.000221154 | 0.056024394 | Zfp683 | zinc finger protein 683 |
| ENSMUSG00000028207 | 0.697111799 | 5.754363225 | 24.32728675 | 0.000221196 | 0.056024394 | Asph | aspartate-beta-hydroxylase |
| ENSMUSG00000027792 | 1.299241566 | 3.946743123 | 24.23226228 | 0.000225558 | 0.056024394 | Bche | butyrylcholinesterase |
| ENSMUSG00000023262 | -0.870903187 | 6.871121603 | 24.03493574 | 0.000233766 | 0.056024394 | Acy1 | aminoacylase 1 |
| ENSMUSG00000032348 | -1.046732265 | 4.785828346 | 23.83195408 | 0.000243126 | 0.056024394 | Gsta4 | glutathione S-transferase, alpha 4 |
| ENSMUSG00000018750 | 0.942636682 | 4.679902985 | 23.74054749 | 0.000247423 | 0.056024394 | Zbtb4 | zinc finger and BTB domain containing 4 |
| ENSMUSG00000052146 | -0.672423333 | 9.207494155 | 23.72526813 | 0.000248007 | 0.056024394 | Rps10 | ribosomal protein S10 |
| ENSMUSG00000055322 | 0.897250759 | 5.45386491 | 23.71023143 | 0.000248789 | 0.056024394 | Tns1 | tensin 1 |
| ENSMUSG00000000805 | -1.30156958 | 5.496199351 | 23.62333673 | 0.000252988 | 0.056024394 | Car4 | carbonic anhydrase 4 |
| ENSMUSG00000049517 | -0.740977354 | 9.824614263 | 23.23920323 | 0.000272429 | 0.056947922 | Rps23 | ribosomal protein S23 |
| ENSMUSG00000009281 | -0.809246022 | 6.244574516 | 23.23004232 | 0.000272945 | 0.056947922 | Rarres2 | retinoic acid receptor responder (tazarotene induced) 2 |
| ENSMUSG00000014956 | 0.669499369 | 7.867678207 | 23.14557999 | 0.000277444 | 0.056947922 | Ppp1cb | protein phosphatase 1 catalytic subunit beta |
| ENSMUSG00000021798 | 0.741984181 | 9.256551703 | 23.13880541 | 0.000277811 | 0.056947922 | Ldb3 | LIM domain binding 3 |
| ENSMUSG00000040061 | -1.321440595 | 3.351880171 | 23.04592108 | 0.000284111 | 0.056947922 | Plcb2 | phospholipase C, beta 2 |
| ENSMUSG00000035181 | 0.759270787 | 5.070171016 | 22.9668026 | 0.000287412 | 0.056947922 | Heatr5a | HEAT repeat containing 5A |

| | | | | | | | |
|---|---|---|---|---|---|---|---|
| ENSMUSG00000043811 | -1.403942352 | 3.205942431 | 22.86798735 | 0.000294665 | 0.057378317 | Rtn4r | reticulon 4 receptor |
| ENSMUSG00000059089 | -2.386605722 | 3.754149869 | 22.58869481 | 0.000311722 | 0.05872512 | Fcgr4 | Fc receptor, IgG, low affinity IV |
| ENSMUSG00000029020 | 0.789652901 | 7.958268187 | 22.51335645 | 0.000314212 | 0.05872512 | Mfn2 | mitofusin 2 |
| ENSMUSG00000031167 | -1.177251854 | 5.827965945 | 22.44504994 | 0.000318572 | 0.05872512 | Rbm3 | RNA binding motif (RNP1, RRM) protein 3 |
| ENSMUSG00000049751 | -1.420366348 | 5.394014256 | 22.38610005 | 0.00032238 | 0.05872512 | Rpl36al | ribosomal protein L36A-like |
| ENSMUSG00000047810 | -1.44408872 | 2.510622575 | 22.14339868 | 0.000344297 | 0.059205891 | Ccdc88b | coiled-coil domain containing 88B |
| ENSMUSG00000045838 | 1.069985055 | 4.434644382 | 22.03187183 | 0.000346244 | 0.059205891 | Ccdc9b | coiled-coil domain containing 9B |
| ENSMUSG00000025508 | -0.729397604 | 9.543961375 | 22.02436092 | 0.000346503 | 0.059205891 | Rplp2 | ribosomal protein lateral stalk subunit P2 |
| ENSMUSG00000021795 | -2.984930056 | 2.905757301 | 22.14585669 | 0.000347179 | 0.059205891 | Sftpd | surfactant associated protein D |
| ENSMUSG00000020657 | 2.417730818 | 1.402397854 | 21.18095063 | 0.000353515 | 0.059205891 | Dnajc27 | DnaJ heat shock protein family (Hsp40) member C27 |
| ENSMUSG00000090862 | -0.616712512 | 8.916771404 | 21.84755885 | 0.000359102 | 0.059205891 | Rps13 | ribosomal protein S13 |
| ENSMUSG00000024735 | 0.665788917 | 7.394312467 | 21.63876386 | 0.000374663 | 0.059205891 | Prpf19 | pre-mRNA processing factor 19 |
| ENSMUSG00000024371 | -2.017999265 | 2.891844377 | 21.63944572 | 0.000381187 | 0.059205891 | C2 | complement C2 |
| ENSMUSG00000050860 | 0.912491061 | 5.831407078 | 21.54979161 | 0.000381582 | 0.059205891 | Phospho1 | phosphatase, orphan 1 |
| ENSMUSG00000022032 | 1.556214955 | 4.006769015 | 21.48225895 | 0.000387516 | 0.059205891 | Scara5 | scavenger receptor class A, member 5 |
| ENSMUSG00000057322 | -0.779058328 | 8.142802021 | 21.46154753 | 0.000388477 | 0.059205891 | Rpl38 | ribosomal protein L38 |
| ENSMUSG00000035621 | 0.852536342 | 4.442611245 | 21.40457117 | 0.000393303 | 0.059205891 | Midn | midnolin |
| ENSMUSG00000025795 | 0.868454547 | 5.442501073 | 21.32250359 | 0.000399814 | 0.059205891 | Rassf3 | Ras association (RalGDS/AF-6) domain family member 3 |
| ENSMUSG00000071317 | 1.535092888 | 4.133158231 | 21.2693951 | 0.000404694 | 0.059205891 | Bves | blood vessel epicardial substance |
| ENSMUSG00000015093 | -2.542308727 | 1.958821895 | 21.1360467 | 0.000408257 | 0.059205891 | Clic3 | chloride intracellular channel 3 |
| ENSMUSG00000024247 | 0.959310237 | 5.906959577 | 21.21312039 | 0.000408895 | 0.059205891 | Pkdcc | protein kinase domain containing, cytoplasmic |
| ENSMUSG00000044934 | 1.672067258 | 2.412217031 | 21.08422999 | 0.00042862 | 0.061037567 | Zfp367 | zinc finger protein 367 |
| ENSMUSG00000047675 | -0.624379722 | 10.24095255 | 20.8864532 | 0.000437497 | 0.061037567 | Rps8 | ribosomal protein S8 |
| ENSMUSG00000027014 | -0.762447778 | 4.231868527 | 20.75706253 | 0.000449849 | 0.061037567 | Cwc22 | CWC22 spliceosome-associated protein |
| ENSMUSG00000005615 | 0.887625297 | 5.725174409 | 20.7086543 | 0.000454131 | 0.061037567 | Pcyt1a | phosphate cytidylyltransferase 1, choline, alpha isoform |
| ENSMUSG00000025163 | -2.565470557 | 2.679147309 | 20.77880432 | 0.00045913 | 0.061037567 | Cd7 | CD7 antigen |
| ENSMUSG00000054580 | -1.453925967 | 2.486916645 | 20.72879578 | 0.000460025 | 0.061037567 | Pla2r1 | phospholipase A2 receptor 1 |
| ENSMUSG00000021228 | -1.82581149 | 4.533488297 | 20.59422379 | 0.000465621 | 0.061037567 | Acot3 | acyl-CoA thioesterase 3 |
| ENSMUSG00000038156 | 0.813662945 | 4.904242903 | 20.58493633 | 0.000466175 | 0.061037567 | Spon1 | spondin 1, (f-spondin) extracellular matrix protein |

| | | | | | | | |
|---|---|---|---|---|---|---|---|
| ENSMUSG00000021273 | -1.101457578 | 5.485089487 | 20.54368577 | 0.000470185 | 0.061037567 | Fdft1 | farnesyl diphosphate farnesyl transferase 1 |
| ENSMUSG00000020925 | 0.681285602 | 4.619179038 | 20.48151865 | 0.000476502 | 0.061154645 | Ccdc43 | coiled-coil domain containing 43 |
| ENSMUSG00000058833 | -0.664563608 | 6.348767238 | 20.33320429 | 0.000491492 | 0.062369804 | Rex1bd | required for excision 1-B domain containing |
| ENSMUSG00000027890.1 | 0.909955378 | 6.592319229 | 20.13081509 | 0.000513097 | 0.063864183 | Gstm4 | glutathione S-transferase, mu 4 |
| ENSMUSG00000023019 | 1.111875719 | 3.981441292 | 20.07359614 | 0.000520001 | 0.063864183 | Gpd1 | glycerol-3-phosphate dehydrogenase 1 (soluble) |
| ENSMUSG00000046008 | -1.689794726 | 9.286389199 | 20.03743757 | 0.000523409 | 0.063864183 | Pnlip | pancreatic lipase |
| ENSMUSG00000024910 | -2.378965137 | 2.914395436 | 20.11080054 | 0.000525887 | 0.063864183 | Ctsw | cathepsin W |
| ENSMUSG00000109572 | 2.50221996 | 3.693595382 | 19.91379146 | 0.000542303 | 0.065157092 | Cfap99 | cilia and flagella associated protein 99 |
| ENSMUSG00000000631 | 0.637035042 | 6.809677777 | 19.80658141 | 0.000550036 | 0.065390614 | Myo18a | myosin XVIIIA |
| ENSMUSG00000028976 | 1.032474616 | 6.376949778 | 19.56240801 | 0.000579884 | 0.06683027 | Slc2a5 | solute carrier family 2 (facilitated glucose transporter), member 5 |
| ENSMUSG00000000711 | 0.584707658 | 5.907998739 | 19.51944059 | 0.000585353 | 0.06683027 | Rab5b | RAB5B, member RAS oncogene family |
| ENSMUSG00000035545 | 0.845858818 | 5.980559067 | 19.46045371 | 0.000592926 | 0.06683027 | Leng8 | leukocyte receptor cluster (LRC) member 8 |
| ENSMUSG00000037813 | -0.899242367 | 4.795342101 | 19.44540495 | 0.000595045 | 0.06683027 | D630003M21Rik | RIKEN cDNA D630003M21 gene |
| ENSMUSG00000053838 | 0.715775221 | 4.700855875 | 19.42552946 | 0.000597627 | 0.06683027 | Nudcd3 | NudC domain containing 3 |
| ENSMUSG00000042737 | -0.852105919 | 5.657817964 | 19.40931909 | 0.000599603 | 0.06683027 | Dpm3 | dolichyl-phosphate mannosyltransferase polypeptide 3 |
| ENSMUSG00000049037 | -2.853894938 | 2.089811376 | 19.21115441 | 0.000603567 | 0.06683027 | Clec4a1 | C-type lectin domain family 4, member a1 |
| ENSMUSG00000040759 | 2.683943754 | 1.372077341 | 18.2868496 | 0.000614157 | 0.067342581 | Cmtm5 | CKLF-like MARVEL transmembrane domain containing 5 |
| ENSMUSG00000028944 | 0.830873701 | 5.129243401 | 19.2000342 | 0.00062778 | 0.06817452 | Prkag2 | protein kinase, AMP-activated, gamma 2 non-catalytic subunit |
| ENSMUSG00000073375 | 1.636512251 | 4.758694587 | 19.07470554 | 0.000645589 | 0.06936359 | Lrrc30 | leucine rich repeat containing 30 |
| ENSMUSG00000059824 | -2.818596725 | 5.185713804 | 19.03889689 | 0.000651013 | 0.06936359 | Dbp | D site albumin promoter binding protein |
| ENSMUSG00000022510 | -4.440878645 | 6.599764029 | 18.82682355 | 0.000682331 | 0.071921812 | Trp63 | transformation related protein 63 |
| ENSMUSG00000043683 | 0.706736919 | 7.111837725 | 18.67945749 | 0.00070436 | 0.071921812 | Fem1a | fem 1 homolog a |
| ENSMUSG00000004730 | -2.377977174 | 4.574422723 | 18.67633302 | 0.000705925 | 0.071921812 | Adgre1 | adhesion G protein-coupled receptor E1 |
| ENSMUSG00000026207 | 0.754828019 | 7.044439655 | 18.65171001 | 0.000708744 | 0.071921812 | Speg | SPEG complex locus |
| ENSMUSG00000059291 | -0.694304623 | 9.468248591 | 18.61741239 | 0.00071419 | 0.071921812 | Rpl11 | ribosomal protein L11 |
| ENSMUSG00000021215 | 0.654740093 | 5.325047967 | 18.53047295 | 0.000728396 | 0.071921812 | Net1 | neuroepithelial cell transforming gene 1 |
| ENSMUSG00000036854 | 0.944824498 | 8.725800748 | 18.52131709 | 0.000729749 | 0.071921812 | Hspb6 | heat shock protein, alpha-crystallin-related, B6 |
| ENSMUSG00000009293 | 0.592211463 | 7.409966343 | 18.45126727 | 0.000741337 | 0.071921812 | Ube2g2 | ubiquitin-conjugating enzyme E2G 2 |
| ENSMUSG00000008683 | -0.815055642 | 9.749551807 | 18.44278832 | 0.000742752 | 0.071921812 | Rps15a | ribosomal protein S15A |

| | | | | | | | |
|---|---|---|---|---|---|---|---|
| ENSMUSG00000022043 | 0.857069209 | 5.724544403 | 18.41180966 | 0.000748056 | 0.071921812 | Trim35 | tripartite motif-containing 35 |
| ENSMUSG00000098274 | -0.603018829 | 10.20920702 | 18.32733101 | 0.000762355 | 0.071921812 | Rpl24 | ribosomal protein L24 |
| ENSMUSG00000024370 | 0.596711802 | 4.933392093 | 18.30000498 | 0.000767299 | 0.071921812 | Cdc23 | CDC23 cell division cycle 23 |
| ENSMUSG00000031389 | -1.010057083 | 4.183441783 | 18.29533563 | 0.000768553 | 0.071921812 | Arhgap4 | Rho GTPase activating protein 4 |
| ENSMUSG00000024304 | 0.561064682 | 7.95055593 | 18.25600795 | 0.000774762 | 0.071921812 | Cdh2 | cadherin 2 |
| ENSMUSG00000020462 | -0.710044791 | 5.447893786 | 18.24982897 | 0.000775989 | 0.071921812 | Cfap36 | cilia and flagella associated protein 36 |
| ENSMUSG00000033174 | -0.746988979 | 6.506434145 | 18.21000814 | 0.000782925 | 0.071921812 | Mgll | monoglyceride lipase |
| ENSMUSG00000020219 | -0.567371339 | 7.367008592 | 18.19925548 | 0.000784807 | 0.071921812 | Timm13 | translocase of inner mitochondrial membrane 13 |
| ENSMUSG00000040283 | 0.581220554 | 7.240093678 | 18.17213978 | 0.00078965 | 0.071921812 | Btnl9 | butyrophilin-like 9 |
| ENSMUSG00000037742 | -0.592639652 | 12.37007746 | 18.13368986 | 0.000796581 | 0.071972658 | Eef1a1 | eukaryotic translation elongation factor 1 alpha 1 |
| ENSMUSG00000020067 | 0.923181329 | 6.405360327 | 18.06484516 | 0.000809217 | 0.072534095 | Mypn | myopalladin |
| ENSMUSG00000054715 | 1.102714763 | 3.316432468 | 18.00619883 | 0.000822288 | 0.07264192 | Zscan22 | zinc finger and SCAN domain containing 22 |
| ENSMUSG00000038593 | -1.121929814 | 3.299307471 | 18.00245056 | 0.000823284 | 0.07264192 | Tctn1 | tectonic family member 1 |
| ENSMUSG00000048310 | 0.814050354 | 4.322199552 | 17.95036072 | 0.000831125 | 0.072765347 | Pskh1 | protein serine kinase H1 |
| ENSMUSG00000039450 | -0.755486147 | 5.095013846 | 17.89044117 | 0.000842322 | 0.072821706 | Dcxr | dicarbonyl L-xylulose reductase |
| ENSMUSG00000036510 | 2.84393152 | 1.873395358 | 17.43507684 | 0.000844665 | 0.072821706 | Cdh8 | cadherin 8 |
| ENSMUSG00000025141 | 0.766765237 | 4.26980844 | 17.74343301 | 0.000871604 | 0.07356989 | Myadml2 | myeloid-associated differentiation marker-like 2 |
| ENSMUSG00000039661 | -1.55749849 | 6.411441548 | 17.6931662 | 0.000881286 | 0.07356989 | Dusp26 | dual specificity phosphatase 26 |
| ENSMUSG00000056121 | 0.609156313 | 6.906147634 | 17.68886921 | 0.000882119 | 0.07356989 | Fez2 | fasciculation and elongation protein zeta 2 |
| ENSMUSG00000022439 | -2.233261325 | 1.73365515 | 17.6226372 | 0.00088601 | 0.07356989 | Parvg | parvin, gamma |
| ENSMUSG00000023067 | 1.018436472 | 6.680700704 | 17.61174438 | 0.000898002 | 0.07356989 | Cdkn1a | cyclin dependent kinase inhibitor 1A |
| ENSMUSG00000020325 | 1.552328307 | 2.829837853 | 17.62515911 | 0.000903478 | 0.07356989 | Fstl3 | follistatin-like 3 |
| ENSMUSG00000046841 | 0.827196496 | 4.847745532 | 17.57629266 | 0.000905664 | 0.07356989 | Ckap4 | cytoskeleton-associated protein 4 |
| ENSMUSG00000024104 | -0.589857248 | 5.522253418 | 17.55532535 | 0.000909926 | 0.07356989 | Washc2 | WASH complex subunit 2 |
| ENSMUSG00000031274 | 0.745842128 | 4.902023464 | 17.54622049 | 0.00091197 | 0.07356989 | Col4a5 | collagen, type IV, alpha 5 |
| ENSMUSG00000021944 | 0.877435473 | 5.345308989 | 17.49258656 | 0.000923324 | 0.073684257 | Gata4 | GATA binding protein 4 |
| ENSMUSG00000043122 | -1.425701626 | 3.965629002 | 17.45577134 | 0.000932398 | 0.073684257 | A530016L24Rik | RIKEN cDNA A530016L24 gene |
| ENSMUSG00000001419 | 0.629130699 | 5.105847956 | 17.42903749 | 0.000937117 | 0.073684257 | Mef2d | myocyte enhancer factor 2D |
| ENSMUSG00000027637 | -0.499731829 | 6.707584633 | 17.40357042 | 0.000942497 | 0.073684257 | Rab5if | RAB5 interacting factor |
| ENSMUSG00000040528 | -1.371341285 | 2.967313211 | 17.38291191 | 0.000952995 | 0.073684257 | Milr1 | mast cell immunoglobulin like receptor 1 |

| | | | | | | | |
|---|---|---|---|---|---|---|---|
| ENSMUSG00000029368 | -2.719539628 | 2.745733075 | 17.38237183 | 0.00096726 | 0.073684257 | Alb | albumin |
| ENSMUSG00000045659 | 1.116094397 | 3.608416572 | 17.28786905 | 0.000969928 | 0.073684257 | Plekha7 | pleckstrin homology domain containing, family A member 7 |
| ENSMUSG00000045725 | -2.072340894 | 2.632297934 | 17.35894283 | 0.000970487 | 0.073684257 | Prr15 | proline rich 15 |
| ENSMUSG00000038274 | -0.654866467 | 9.392990598 | 17.26158997 | 0.000974276 | 0.073684257 | Fau | FAU ubiquitin like and ribosomal protein S30 fusion |
| ENSMUSG00000028906 | 0.642975256 | 5.695331987 | 17.21203099 | 0.000985797 | 0.073684257 | Epb41 | erythrocyte membrane protein band 4.1 |
| ENSMUSG00000052406 | 0.671903769 | 4.497070357 | 17.18168006 | 0.000993175 | 0.073684257 | Rexo4 | REX4, 3'-5' exonuclease |
| ENSMUSG00000028898 | -0.747939933 | 4.941196734 | 17.17962983 | 0.000993495 | 0.073684257 | Trnau1ap | tRNA selenocysteine 1 associated protein 1 |
| ENSMUSG00000039994 | 0.897257561 | 4.852740278 | 17.15029881 | 0.001000388 | 0.073684257 | Timeless | timeless circadian clock 1 |
| ENSMUSG00000042686 | 1.160331993 | 3.660354086 | 17.13740933 | 0.001004726 | 0.073684257 | Jph1 | junctophilin 1 |
| ENSMUSG00000067288 | -0.747936046 | 9.295587806 | 16.97554018 | 0.001042171 | 0.075146357 | Rps28 | ribosomal protein S28 |
| ENSMUSG00000028398 | -0.646280935 | 5.756716975 | 16.97476904 | 0.001042495 | 0.075146357 | Dmac1 | distal membrane arm assembly complex 1 |
| ENSMUSG00000039221 | -0.895759772 | 5.148514147 | 16.96670274 | 0.001044624 | 0.075146357 | Rpl22l1 | ribosomal protein L22 like 1 |
| ENSMUSG00000058600 | -0.601165541 | 9.024040782 | 16.87980995 | 0.001066094 | 0.075869767 | Rpl30 | ribosomal protein L30 |
| ENSMUSG00000029669 | 0.631093012 | 5.732805698 | 16.87234106 | 0.001068115 | 0.075869767 | Tspan12 | tetraspanin 12 |
| ENSMUSG00000070287 | 1.773839338 | 2.301559721 | 16.90553588 | 0.001078242 | 0.076110374 | Slc35g2 | solute carrier family 35, member G2 |
| ENSMUSG00000061983 | -0.661027647 | 9.720935357 | 16.7682977 | 0.001094758 | 0.076796279 | Rps12 | ribosomal protein S12 |
| ENSMUSG00000068699 | 0.900880906 | 8.14596064 | 16.72285228 | 0.001106694 | 0.07694782 | Flnc | filamin C, gamma |
| ENSMUSG00000013076 | 1.373485516 | 2.213095227 | 16.77904297 | 0.001110545 | 0.07694782 | Amotl1 | angiomotin-like 1 |
| ENSMUSG00000020439 | 0.665774295 | 6.206596849 | 16.62951689 | 0.001131753 | 0.077107909 | Smtn | smoothelin |
| ENSMUSG00000036322 | -2.058209405 | 8.212087752 | 16.57727943 | 0.001145951 | 0.077107909 | H2-Ea | histocompatibility 2, class II antigen E alpha |
| ENSMUSG00000025362 | -0.6809262 | 9.688484657 | 16.56427817 | 0.001149522 | 0.077107909 | Rps26 | ribosomal protein S26 |
| ENSMUSG00000023828 | 1.445944172 | 4.224937852 | 16.55680559 | 0.001152637 | 0.077107909 | Slc22a3 | solute carrier family 22 (organic cation transporter), member 3 |
| ENSMUSG00000024743 | 0.895628027 | 3.330217843 | 16.55887327 | 0.001153404 | 0.077107909 | Syt7 | synaptotagmin VII |
| ENSMUSG00000057672 | -0.613224963 | 7.207535237 | 16.53919328 | 0.001156482 | 0.077107909 | Pkn1 | protein kinase N1 |
| ENSMUSG00000030704 | 0.773541247 | 4.352203315 | 16.4861332 | 0.00117195 | 0.077107909 | Rab6a | RAB6A, member RAS oncogene family |
| ENSMUSG00000037563 | -0.600005483 | 10.26241349 | 16.47132044 | 0.001175512 | 0.077107909 | Rps16 | ribosomal protein S16 |
| ENSMUSG00000022304 | 2.152135387 | 2.500737101 | 16.52401862 | 0.00117885 | 0.077107909 | Dpys | dihydropyrimidinase |
| ENSMUSG00000022994 | 0.808006814 | 6.10854785 | 16.45188079 | 0.001181129 | 0.077107909 | Adcy6 | adenylate cyclase 6 |

| | | | | | | | |
|---|---|---|---|---|---|---|---|
| ENSMUSG00000037321 | -1.112745234 | 5.700696298 | 16.42508418 | 0.001188865 | 0.077166935 | Tap1 | transporter 1, ATP-binding cassette, sub-family B (MDR/TAP) |
| ENSMUSG00000054226.2 | -3.046516149 | 1.693695708 | 15.81337035 | 0.001196297 | 0.077205605 | Tprkb | Tp53rk binding protein |
| ENSMUSG00000035443 | -0.608283437 | 4.983609187 | 16.35897713 | 0.001208148 | 0.077490963 | Thyn1 | thymocyte nuclear protein 1 |
| ENSMUSG00000071415 | -0.709713731 | 10.21688659 | 16.31591147 | 0.001220483 | 0.077490963 | Rpl23 | ribosomal protein L23 |
| ENSMUSG00000070394 | -0.680507935 | 7.089171015 | 16.30032083 | 0.001225125 | 0.077490963 | Tmem256 | transmembrane protein 256 |
| ENSMUSG00000022519 | 0.908413354 | 6.64379103 | 16.27310898 | 0.001233263 | 0.077490963 | Srl | sarcalumenin |
| ENSMUSG00000053192 | 1.2655276 | 3.178057278 | 16.24773441 | 0.001245288 | 0.077490963 | Mllt11 | myeloid/lymphoid or mixed-lineage leukemia; translocated to, 11 |
| ENSMUSG00000021967 | -0.896845798 | 6.988638966 | 16.22562541 | 0.001247547 | 0.077490963 | Mrpl57 | mitochondrial ribosomal protein L57 |
| ENSMUSG00000020211 | 0.780220423 | 4.610040214 | 16.21101265 | 0.001252436 | 0.077490963 | Sf3a2 | splicing factor 3a, subunit 2 |
| ENSMUSG00000036905 | -1.267583138 | 7.162029721 | 16.19913384 | 0.001255609 | 0.077490963 | C1qb | complement component 1, q subcomponent, beta polypeptide |
| ENSMUSG00000059901 | 1.112354681 | 3.375800585 | 16.07643733 | 0.001296492 | 0.079513719 | Adamts14 | ADAM metallopeptidase with thrombospondin type 1 motif 14 |
| ENSMUSG00000043505 | -1.016199862 | 3.445169219 | 16.05701817 | 0.001302465 | 0.079513719 | Gimap5 | GTPase, IMAP family member 5 |
| ENSMUSG00000018008 | -1.487182721 | 4.482919909 | 15.9149247 | 0.001346838 | 0.0813213 | Cyth4 | cytohesin 4 |
| ENSMUSG00000047417 | 0.832522908 | 4.338287325 | 15.91095418 | 0.001347959 | 0.0813213 | Rexo1 | REX1, RNA exonuclease 1 |
| ENSMUSG00000079037 | 0.612004623 | 7.333692711 | 15.89160049 | 0.001353675 | 0.0813213 | Prnp | prion protein |
| ENSMUSG00000006333 | -0.517083972 | 9.683033875 | 15.85638046 | 0.001365457 | 0.081381108 | Rps9 | ribosomal protein S9 |
| ENSMUSG00000021266 | -0.633201989 | 6.863684723 | 15.83716242 | 0.001371972 | 0.081381108 | Wars1 | tryptophanyl-tRNA synthetase1 |
| ENSMUSG00000036781 | -0.700975472 | 6.81814263 | 15.81270671 | 0.001380271 | 0.081381108 | Rps27l | ribosomal protein S27-like |
| ENSMUSG00000022575 | -1.349138607 | 3.49393395 | 15.80327581 | 0.001386709 | 0.081381108 | Gsdmd | gasdermin D |
| ENSMUSG00000000326 | 0.689069513 | 6.546776151 | 15.78226492 | 0.001390699 | 0.081381108 | Comt | catechol-O-methyltransferase |
| ENSMUSG00000004947 | -0.888402978 | 3.741039289 | 15.76523047 | 0.001398164 | 0.081396208 | Dtx2 | deltex 2, E3 ubiquitin ligase |
| ENSMUSG00000040883 | -0.933116551 | 4.92665388 | 15.70132266 | 0.001419182 | 0.081747743 | Tmem205 | transmembrane protein 205 |
| ENSMUSG00000041115 | 0.625214736 | 5.017020021 | 15.69660749 | 0.001420746 | 0.081747743 | Iqsec2 | IQ motif and Sec7 domain 2 |
| ENSMUSG00000049907 | 1.039071868 | 5.877610563 | 15.68144879 | 0.001425917 | 0.081747743 | Rasl11b | RAS-like, family 11, member B |
| ENSMUSG00000024782 | 0.729835962 | 6.127499635 | 15.64743498 | 0.001437948 | 0.081869813 | Ak3 | adenylate kinase 3 |
| ENSMUSG00000021590 | -3.152079142 | 3.915082174 | 15.67347533 | 0.001442544 | 0.081869813 | Spata9 | spermatogenesis associated 9 |

| | | | | | | | |
|---|---|---|---|---|---|---|---|
| ENSMUSG00000060802 | -1.271255051 | 9.291387979 | 15.60929878 | 0.001451529 | 0.081967832 | B2m | beta-2 microglobulin |
| ENSMUSG00000039844 | 0.585670788 | 5.349003043 | 15.56594675 | 0.001467502 | 0.082297284 | Rapgef1 | Rap guanine nucleotide exchange factor (GEF) 1 |
| ENSMUSG00000025348 | 0.976760354 | 7.479183456 | 15.52783679 | 0.001481263 | 0.082297284 | Itga7 | integrin alpha 7 |
| ENSMUSG00000017756 | 0.602379031 | 6.4130973 | 15.51993292 | 0.001484259 | 0.082297284 | Slc12a7 | solute carrier family 12, member 7 |
| ENSMUSG00000031626 | 0.884183465 | 6.777980896 | 15.51379047 | 0.00148651 | 0.082297284 | Sorbs2 | sorbin and SH3 domain containing 2 |
| ENSMUSG00000020460 | -0.632742967 | 9.604034023 | 15.49054178 | 0.001495103 | 0.08236925 | Rps27a | ribosomal protein S27A |
| ENSMUSG00000053646 | 0.789025082 | 4.968008942 | 15.44680978 | 0.001511892 | 0.082889854 | Plxnb1 | plexin B1 |
| ENSMUSG00000057388 | -0.832165204 | 7.850984082 | 15.37639617 | 0.001538409 | 0.083283756 | Mrpl18 | mitochondrial ribosomal protein L18 |
| ENSMUSG00000052934 | 0.85146913 | 5.452521871 | 15.35498661 | 0.001546927 | 0.083283756 | Fbxo31 | F-box protein 31 |
| ENSMUSG00000026117 | -1.393599779 | 2.24965794 | 15.40196735 | 0.001552766 | 0.083283756 | Zap70 | zeta-chain (TCR) associated protein kinase |
| ENSMUSG00000071001 | -0.979169098 | 4.907457335 | 15.29685145 | 0.001569901 | 0.083283756 | Hrct1 | histidine rich carboxyl terminus 1 |
| ENSMUSG00000035711 | -1.602104517 | 2.687311815 | 15.30221012 | 0.001583415 | 0.083283756 | Dok3 | docking protein 3 |
| ENSMUSG00000022129 | 2.967366508 | 1.815870443 | 14.75480889 | 0.00158354 | 0.083283756 | Dct | dopachrome tautomerase |
| ENSMUSG00000028273 | 0.595101303 | 8.716630828 | 15.25955992 | 0.001584229 | 0.083283756 | Pdlim5 | PDZ and LIM domain 5 |
| ENSMUSG00000022018 | -1.116401415 | 7.659904695 | 15.25651718 | 0.001585456 | 0.083283756 | Rgcc | regulator of cell cycle |
| ENSMUSG00000044337 | -0.725350744 | 6.791392457 | 15.22779207 | 0.001596997 | 0.083283756 | Ackr3 | atypical chemokine receptor 3 |
| ENSMUSG00000051236 | 0.627661583 | 6.147106076 | 15.22156435 | 0.001599576 | 0.083283756 | Msrb3 | methionine sulfoxide reductase B3 |
| ENSMUSG00000002668 | -1.298993556 | 2.614165497 | 15.25501306 | 0.001600193 | 0.083283756 | Dennd1c | DENN domain containing 1C |
| ENSMUSG00000022552 | -0.458865027 | 5.943523123 | 15.19352157 | 0.001610957 | 0.083459413 | Sharpin | SHANK-associated RH domain interacting protein |
| ENSMUSG00000031133 | 0.675550631 | 4.733046641 | 15.15047809 | 0.001628911 | 0.083794551 | Arhgef6 | Rac/Cdc42 guanine nucleotide exchange factor 6 |
| ENSMUSG00000091537 | -0.594221162 | 7.174840147 | 15.13186665 | 0.001636126 | 0.083794551 | Tma7 | translational machinery associated 7 |
| ENSMUSG00000033220 | -1.932946466 | 4.210162915 | 15.12119875 | 0.001642673 | 0.083794551 | Rac2 | Rac family small GTPase 2 |
| ENSMUSG00000046330.2 | -0.708186968 | 10.02009305 | 15.10540546 | 0.001647104 | 0.083794551 | Rpl37a | ribosomal protein L37a |
| ENSMUSG00000018566 | 0.491881377 | 7.877941203 | 15.07835016 | 0.001658434 | 0.083992611 | Slc2a4 | solute carrier family 2 (facilitated glucose transporter), member 4 |
| ENSMUSG00000029664 | -0.964349066 | 3.560955744 | 15.05853686 | 0.001669314 | 0.084166237 | Tfpi2 | tissue factor pathway inhibitor 2 |
| ENSMUSG00000020257 | 0.618800117 | 5.241783881 | 14.96612718 | 0.001706704 | 0.085668945 | Wdr82 | WD repeat domain containing 82 |
| ENSMUSG00000033735 | 1.121964855 | 6.620495602 | 14.90538368 | 0.00173307 | 0.086173266 | Spr | sepiapterin reductase |
| ENSMUSG00000005836 | 0.684403961 | 6.256043737 | 14.89661746 | 0.001736977 | 0.086173266 | Gata6 | GATA binding protein 6 |

| | | | | | | | |
|---|---|---|---|---|---|---|---|
| ENSMUSG00000041841 | -0.710178172 | 8.602169001 | 14.89040809 | 0.001739641 | 0.086173266 | Rpl37 | ribosomal protein L37 |
| ENSMUSG00000017404 | -0.581862025 | 9.851401906 | 14.8001204 | 0.00178027 | 0.087800754 | Rpl19 | ribosomal protein L19 |
| ENSMUSG00000026385 | -0.675797447 | 8.034420494 | 14.78279739 | 0.001788192 | 0.087807978 | Dbi | diazepam binding inhibitor |
| ENSMUSG00000004098 | 1.188167908 | 5.904392717 | 14.75498869 | 0.001801166 | 0.088062221 | Col5a3 | collagen, type V, alpha 3 |
| ENSMUSG00000062939 | -1.721864498 | 1.918007905 | 14.79771478 | 0.001812381 | 0.0882286 | Stat4 | signal transducer and activator of transcription 4 |
| ENSMUSG00000034892 | -0.713397696 | 10.45835058 | 14.67458094 | 0.001838588 | 0.088791825 | Rps29 | ribosomal protein S29 |
| ENSMUSG00000001418 | -0.491256068 | 5.608408486 | 14.67274447 | 0.001839675 | 0.088791825 | Glmp | glycosylated lysosomal membrane protein |
| ENSMUSG00000029019 | 1.793749609 | 8.820051053 | 14.65154293 | 0.001849528 | 0.088887528 | Nppb | natriuretic peptide type B |
| ENSMUSG00000096923 | 1.392915717 | 2.609625963 | 14.66524911 | 0.001859647 | 0.08891592 | Grb2 | growth factor receptor bound protein 2 |
| ENSMUSG00000024610 | -1.851886729 | 8.03190775 | 14.61745639 | 0.001865864 | 0.08891592 | Cd74 | CD74 antigen (invariant polypeptide of major histocompatibility complex, class II antigen-associated) |
| ENSMUSG00000016526 | 0.985961034 | 4.03740512 | 14.57785518 | 0.001886424 | 0.089510084 | Dyrk3 | dual-specificity tyrosine phosphorylation regulated kinase 3 |
| ENSMUSG00000031825 | 0.76097886 | 5.073663529 | 14.55726184 | 0.001895463 | 0.089510084 | Crispld2 | cysteine-rich secretory protein LCCL domain containing 2 |
| ENSMUSG00000058715 | -1.324933914 | 5.349230033 | 14.54374845 | 0.001902109 | 0.089510084 | Fcer1g | Fc receptor, IgE, high affinity I, gamma polypeptide |
| ENSMUSG00000043822 | 1.047943074 | 5.789343835 | 14.49367552 | 0.001926732 | 0.089914322 | Adamtsl5 | ADAMTS-like 5 |
| ENSMUSG00000103409 | 1.119850482 | 4.568269064 | 14.49470323 | 0.00192682 | 0.089914322 | Lsmem2 | leucine-rich single-pass membrane protein 2 |
| ENSMUSG00000059070 | -0.564992699 | 9.883631414 | 14.44692268 | 0.001950032 | 0.089914322 | Rpl18 | ribosomal protein L18 |
| ENSMUSG00000029217 | -1.186853427 | 2.153303779 | 14.50308342 | 0.001950457 | 0.089914322 | Tec | tec protein tyrosine kinase |
| ENSMUSG00000032387 | 0.859917343 | 4.302124912 | 14.44792607 | 0.001950505 | 0.089914322 | Rbpms2 | RNA binding protein with multiple splicing 2 |
| ENSMUSG00000050947 | 1.835292544 | 1.452974437 | 14.35592746 | 0.001959432 | 0.089947688 | Amigo1 | adhesion molecule with Ig like domain 1 |
| ENSMUSG00000019970 | -0.949743782 | 6.37097659 | 14.38538228 | 0.001981582 | 0.089947688 | Sgk1 | serum/glucocorticoid regulated kinase 1 |
| ENSMUSG00000015846 | 0.766034736 | 3.548574126 | 14.38000765 | 0.001986678 | 0.089947688 | Rxra | retinoid X receptor alpha |
| ENSMUSG00000042770 | -0.747952156 | 5.688333149 | 14.37534765 | 0.001986889 | 0.089947688 | Hebp1 | heme binding protein 1 |
| ENSMUSG00000006589 | -0.614571271 | 7.059741451 | 14.36055152 | 0.001994348 | 0.089947688 | Aprt | adenine phosphoribosyl transferase |
| ENSMUSG00000032816 | 0.8527166 | 4.655737575 | 14.35279001 | 0.001999014 | 0.089947688 | Igdcc4 | immunoglobulin superfamily, DCC subclass, member 4 |
| ENSMUSG00000009772 | -0.925904409 | 3.838680968 | 14.31411649 | 0.002020579 | 0.089978846 | Nuak2 | NUAK family, SNF1-like kinase, 2 |
| ENSMUSG00000038375 | 1.187054652 | 4.11874075 | 14.30988515 | 0.002022323 | 0.089978846 | Trp53inp2 | transformation related protein 53 inducible nuclear protein 2 |
| ENSMUSG00000050315 | 0.554178967 | 5.502037099 | 14.30511846 | 0.002023608 | 0.089978846 | Synpo2 | synaptopodin 2 |

| | | | | | | | |
|---|---|---|---|---|---|---|---|
| ENSMUSG00000018583.11 | 2.3076008 | 3.381028367 | 14.29147865 | 0.002045971 | 0.090486888 | G3bp1 | G3BP stress granule assembly factor 1 |
| ENSMUSG00000047514 | 0.6438148 | 5.584534226 | 14.23990733 | 0.002058378 | 0.090486888 | Tspyl1 | testis-specific protein, Y-encoded-like 1 |
| ENSMUSG00000006219 | 1.092894237 | 4.057617539 | 14.23660264 | 0.002061439 | 0.090486888 | Fblim1 | filamin binding LIM protein 1 |
| ENSMUSG00000015016 | 0.768408143 | 5.042311364 | 14.21078123 | 0.002074341 | 0.090486888 | Acsf3 | acyl-CoA synthetase family member 3 |
| ENSMUSG00000060675 | -1.245517383 | 6.733491139 | 14.20592252 | 0.002076629 | 0.090486888 | Plaat3 | phospholipase A and acyltransferase 3 |
| ENSMUSG00000036208 | -1.157431489 | 4.123711456 | 14.19681728 | 0.002083105 | 0.090486888 | Nepro | nucleolus and neural progenitor protein |
| ENSMUSG00000022148 | -1.918831552 | 2.843678473 | 14.20814399 | 0.002096656 | 0.09060075 | Fyb1 | FYN binding protein 1 |
| ENSMUSG00000029432 | 0.65300969 | 8.45401937 | 14.13962072 | 0.002112994 | 0.09060075 | Nipsnap2 | nipsnap homolog 2 |
| ENSMUSG00000041426 | 0.695257555 | 6.510854229 | 14.13268392 | 0.002116934 | 0.09060075 | Hibch | 3-hydroxyisobutyryl-Coenzyme A hydrolase |
| ENSMUSG00000003283 | -1.492452078 | 3.476356425 | 14.14004202 | 0.002117815 | 0.09060075 | Hck | hemopoietic cell kinase |
| ENSMUSG00000035776 | 0.761814755 | 4.345346831 | 14.03593589 | 0.002172427 | 0.092586387 | Cd99l2 | CD99 antigen-like 2 |
| ENSMUSG00000012848 | -0.486223331 | 9.371583501 | 13.94171167 | 0.002226222 | 0.094522378 | Rps5 | ribosomal protein S5 |
| ENSMUSG00000060032 | -0.554977364 | 6.549768749 | 13.9242882 | 0.0022366 | 0.094607359 | H2aj | H2J.A histone |
| ENSMUSG00000024780 | 0.690252222 | 6.386060447 | 13.89018586 | 0.00225694 | 0.095111513 | Cdc37l1 | cell division cycle 37-like 1 |
| ENSMUSG00000039208 | -0.665047067 | 5.795846595 | 13.86964268 | 0.002269385 | 0.095280416 | Metrnl | meteorin, glial cell differentiation regulator-like |
| ENSMUSG00000025213 | -1.71558559 | 1.787458874 | 13.84076444 | 0.002319172 | 0.096755715 | Kazald1 | Kazal-type serine peptidase inhibitor domain 1 |
| ENSMUSG00000024397 | -1.286705744 | 4.268034253 | 13.78612499 | 0.002321657 | 0.096755715 | Aif1 | allograft inflammatory factor 1 |
| ENSMUSG00000038059 | -1.17893006 | 2.557204573 | 13.76441028 | 0.00235205 | 0.097604601 | Smim3 | small integral membrane protein 3 |
| ENSMUSG00000021957 | -0.607741938 | 6.461275037 | 13.72103419 | 0.00236087 | 0.097604601 | Tkt | transketolase |
| ENSMUSG00000040511 | 1.047069617 | 4.43681785 | 13.71134614 | 0.002367953 | 0.097604601 | Pvr | poliovirus receptor |
| ENSMUSG00000028042 | 0.779491556 | 5.08205158 | 13.67091422 | 0.002393087 | 0.098018234 | Zbtb7b | zinc finger and BTB domain containing 7B |
| ENSMUSG00000024826 | 0.549683387 | 6.215039343 | 13.66688192 | 0.002395346 | 0.098018234 | Dpf2 | double PHD fingers 2 |
| ENSMUSG00000047215 | -0.546830721 | 9.592136884 | 13.64415332 | 0.002409841 | 0.098255408 | Rpl9 | ribosomal protein L9 |
| ENSMUSG00000034659 | 0.788228542 | 6.512524535 | 13.58570629 | 0.002448002 | 0.099452294 | Tmem109 | transmembrane protein 109 |

**Table S4.** Gene ontology enrichment analysis of significant (FDR ≤ 0.1) differentially expressed genes in the RV transcriptomes of high-elevation *P. vaccarum*.

| p_value | term_size | query_size | intersection_size | precision | recall | term_id | source | term_name |
|---|---|---|---|---|---|---|---|---|
| 0.00000 | 45 | 273 | 16 | 0.0586 | 0.3556 | GO:0140242 | GO:BP | translation at postsynapse |
| 0.00000 | 45 | 273 | 16 | 0.0586 | 0.3556 | GO:0140241 | GO:BP | translation at synapse |
| 0.00000 | 147 | 273 | 26 | 0.0952 | 0.1769 | GO:0002181 | GO:BP | cytoplasmic translation |
| 0.00000 | 44 | 273 | 16 | 0.0586 | 0.3636 | GO:0140236 | GO:BP | translation at presynapse |
| 0.00026 | 564 | 273 | 35 | 0.1282 | 0.0621 | GO:0006412 | GO:BP | translation |
| 0.00048 | 98 | 273 | 13 | 0.0476 | 0.1327 | GO:0042274 | GO:BP | ribosomal small subunit biogenesis |
| 0.00427 | 511 | 273 | 30 | 0.1099 | 0.0587 | GO:0098609 | GO:BP | cell-cell adhesion |
| 0.00610 | 59 | 273 | 9 | 0.0330 | 0.1525 | GO:0019882 | GO:BP | antigen processing and presentation |
| 0.01155 | 264 | 273 | 19 | 0.0696 | 0.0720 | GO:0002250 | GO:BP | adaptive immune response |
| 0.01236 | 16 | 273 | 5 | 0.0183 | 0.3125 | GO:0002478 | GO:BP | antigen processing and presentation of exogenous peptide antigen |
| 0.01839 | 28 | 273 | 6 | 0.0220 | 0.2143 | GO:0048002 | GO:BP | antigen processing and presentation of peptide antigen |
| 0.01941 | 18 | 273 | 5 | 0.0183 | 0.2778 | GO:0000028 | GO:BP | ribosomal small subunit assembly |
| 0.02672 | 77 | 273 | 9 | 0.0330 | 0.1169 | GO:0006959 | GO:BP | humoral immune response |
| 0.02672 | 11 | 273 | 4 | 0.0147 | 0.3636 | GO:0002468 | GO:BP | dendritic cell antigen processing and presentation |
| 0.02672 | 20 | 273 | 5 | 0.0183 | 0.2500 | GO:0019884 | GO:BP | antigen processing and presentation of exogenous antigen |
| 0.02672 | 11 | 273 | 4 | 0.0147 | 0.3636 | GO:0019886 | GO:BP | antigen processing and presentation of exogenous peptide antigen via MHC class II |
| 0.02723 | 828 | 273 | 38 | 0.1392 | 0.0459 | GO:0007155 | GO:BP | cell adhesion |
| 0.03511 | 140 | 273 | 12 | 0.0440 | 0.0857 | GO:0050670 | GO:BP | regulation of lymphocyte proliferation |
| 0.03511 | 967 | 273 | 42 | 0.1538 | 0.0434 | GO:0006955 | GO:BP | immune response |
| 0.03839 | 144 | 273 | 12 | 0.0440 | 0.0833 | GO:0032944 | GO:BP | regulation of mononuclear cell proliferation |
| 0.03839 | 143 | 273 | 12 | 0.0440 | 0.0839 | GO:0055002 | GO:BP | striated muscle cell development |
| 0.03839 | 13 | 273 | 4 | 0.0147 | 0.3077 | GO:0002495 | GO:BP | antigen processing and presentation of peptide antigen via MHC class II |
| 0.03839 | 187 | 273 | 14 | 0.0513 | 0.0749 | GO:0046651 | GO:BP | lymphocyte proliferation |

| | | | | | | | | |
|---|---|---|---|---|---|---|---|---|
| 0.04270 | 24 | 273 | 5 | 0.0183 | 0.2083 | GO:0044331 | GO:BP | cell-cell adhesion mediated by cadherin |
| 0.04539 | 193 | 273 | 14 | 0.0513 | 0.0725 | GO:0032943 | GO:BP | mononuclear cell proliferation |
| 0.04853 | 195 | 273 | 14 | 0.0513 | 0.0718 | GO:0033002 | GO:BP | muscle cell proliferation |
| 0.00000 | 103 | 273 | 27 | 0.0989 | 0.2621 | GO:0022626 | GO:CC | cytosolic ribosome |
| 0.00000 | 175 | 273 | 30 | 0.1099 | 0.1714 | GO:0044391 | GO:CC | ribosomal subunit |
| 0.00000 | 38 | 273 | 16 | 0.0586 | 0.4211 | GO:0022627 | GO:CC | cytosolic small ribosomal subunit |
| 0.00000 | 210 | 273 | 31 | 0.1136 | 0.1476 | GO:0005840 | GO:CC | ribosome |
| 0.00000 | 71 | 273 | 16 | 0.0586 | 0.2254 | GO:0015935 | GO:CC | small ribosomal subunit |
| 0.00000 | 49 | 273 | 12 | 0.0440 | 0.2449 | GO:0022625 | GO:CC | cytosolic large ribosomal subunit |
| 0.00000 | 108 | 273 | 15 | 0.0549 | 0.1389 | GO:0015934 | GO:CC | large ribosomal subunit |
| 0.00001 | 70 | 273 | 12 | 0.0440 | 0.1714 | GO:0032040 | GO:CC | small-subunit processome |
| 0.00004 | 510 | 273 | 32 | 0.1172 | 0.0627 | GO:0098793 | GO:CC | presynapse |
| 0.00007 | 605 | 273 | 35 | 0.1282 | 0.0579 | GO:1990904 | GO:CC | ribonucleoprotein complex |
| 0.00011 | 64 | 273 | 10 | 0.0366 | 0.1563 | GO:0016529 | GO:CC | sarcoplasmic reticulum |
| 0.00011 | 1558 | 273 | 66 | 0.2418 | 0.0424 | GO:0030054 | GO:CC | cell junction |
| 0.00018 | 84 | 273 | 11 | 0.0403 | 0.1310 | GO:0016528 | GO:CC | sarcoplasm |
| 0.00018 | 100 | 273 | 12 | 0.0440 | 0.1200 | GO:0030684 | GO:CC | preribosome |
| 0.00180 | 927 | 273 | 42 | 0.1538 | 0.0453 | GO:0005576 | GO:CC | extracellular region |
| 0.00205 | 110 | 273 | 11 | 0.0403 | 0.1000 | GO:0030018 | GO:CC | Z disc |
| 0.00318 | 487 | 273 | 26 | 0.0952 | 0.0534 | GO:0009986 | GO:CC | cell surface |
| 0.00318 | 5 | 273 | 3 | 0.0110 | 0.6000 | GO:0042611 | GO:CC | MHC protein complex |
| 0.00318 | 5 | 273 | 3 | 0.0110 | 0.6000 | GO:0042613 | GO:CC | MHC class II protein complex |
| 0.00318 | 5 | 273 | 3 | 0.0110 | 0.6000 | GO:0030314 | GO:CC | junctional membrane complex |
| 0.00357 | 49 | 273 | 7 | 0.0256 | 0.1429 | GO:0005791 | GO:CC | rough endoplasmic reticulum |
| 0.00408 | 123 | 273 | 11 | 0.0403 | 0.0894 | GO:0031674 | GO:CC | I band |
| 0.00438 | 190 | 273 | 14 | 0.0513 | 0.0737 | GO:0009897 | GO:CC | external side of plasma membrane |
| 0.00502 | 6 | 273 | 3 | 0.0110 | 0.5000 | GO:0098556 | GO:CC | cytoplasmic side of rough endoplasmic reticulum membrane |
| 0.00584 | 1163 | 273 | 47 | 0.1722 | 0.0404 | GO:0045202 | GO:CC | synapse |

| | | | | | | | | |
|---|---|---|---|---|---|---|---|---|
| 0.00760 | 383 | 273 | 21 | 0.0769 | 0.0548 | GO:0098552 | GO:CC | side of membrane |
| 0.00909 | 650 | 273 | 30 | 0.1099 | 0.0462 | GO:0005615 | GO:CC | extracellular space |
| 0.00966 | 29 | 273 | 5 | 0.0183 | 0.1724 | GO:0033017 | GO:CC | sarcoplasmic reticulum membrane |
| 0.01242 | 2927 | 273 | 95 | 0.3480 | 0.0325 | GO:0071944 | GO:CC | cell periphery |
| 0.01323 | 639 | 273 | 29 | 0.1062 | 0.0454 | GO:0098794 | GO:CC | postsynapse |
| 0.02197 | 205 | 273 | 13 | 0.0476 | 0.0634 | GO:0043292 | GO:CC | contractile muscle fiber |
| 0.04204 | 41 | 273 | 5 | 0.0183 | 0.1220 | GO:0005771 | GO:CC | multivesicular body |
| 0.04380 | 278 | 273 | 15 | 0.0549 | 0.0540 | GO:0031012 | GO:CC | extracellular matrix |
| 0.04380 | 278 | 273 | 15 | 0.0549 | 0.0540 | GO:0030312 | GO:CC | external encapsulating structure |
| 0.04929 | 62 | 273 | 6 | 0.0220 | 0.0968 | GO:0030666 | GO:CC | endocytic vesicle membrane |
| 0.00000 | 150 | 273 | 30 | 0.1099 | 0.2000 | GO:0003735 | GO:MF | structural constituent of ribosome |
| 0.00000 | 404 | 273 | 39 | 0.1429 | 0.0965 | GO:0005198 | GO:MF | structural molecule activity |
| 0.01134 | 70 | 273 | 9 | 0.0330 | 0.1286 | GO:0019843 | GO:MF | rRNA binding |
| 0.02502 | 5 | 273 | 3 | 0.0110 | 0.6000 | GO:1990932 | GO:MF | 5.8S rRNA binding |
| 0.00000 | 125 | 273 | 29 | 0.1062 | 0.2320 | KEGG:03010 | KEGG | Ribosome |
| 0.00000 | 169 | 273 | 31 | 0.1136 | 0.1834 | KEGG:05171 | KEGG | Coronavirus disease - COVID-19 |

**Table S5.** Overlap of differentially expressed genes in the RV transcriptomes of high-elevation *P. vaccarum* (this study) and high-elevation deer mice (*Peromyscus maniculatus*; Velotta et al. 2018).

| gene_ID | name | description |
|---|---|---|
| ENSMUSG00000003934 | Efnb3 | ephrin B3 |
| ENSMUSG00000028444 | Cntfr | ciliary neurotrophic factor receptor |
| ENSMUSG00000032834 | Pwp2 | PWP2 periodic tryptophan protein homolog (yeast) |
| ENSMUSG00000040147 | Maob | monoamine oxidase B |
| ENSMUSG00000039376 | Synpo2l | synaptopodin 2-like |
| ENSMUSG00000059852 | Kcng2 | potassium voltage-gated channel, subfamily G, member 2 |
| ENSMUSG00000006221 | Hspb7 | heat shock protein family, member 7 (cardiovascular) |
| ENSMUSG00000096727 | Psmb9 | proteasome (prosome, macropain) subunit, beta type 9 (large multifunctional peptidase 2) |
| ENSMUSG00000030352 | Tspan9 | tetraspanin 9 |
| ENSMUSG00000044788 | Fads6 | fatty acid desaturase domain family, member 6 |
| ENSMUSG00000031378 | Abcd1 | ATP-binding cassette, sub-family D member 1 |
| ENSMUSG00000033039 | Micall1 | microtubule associated monooxygenase, calponin and LIM domain containing -like 1 |
| ENSMUSG00000049410 | Zfp683 | zinc finger protein 683 |
| ENSMUSG00000028207 | Asph | aspartate-beta-hydroxylase |
| ENSMUSG00000032348 | Gsta4 | glutathione S-transferase, alpha 4 |
| ENSMUSG00000055322 | Tns1 | tensin 1 |
| ENSMUSG00000031167 | Rbm3 | RNA binding motif (RNP1, RRM) protein 3 |
| ENSMUSG00000020657 | Dnajc27 | DnaJ heat shock protein family (Hsp40) member C27 |
| ENSMUSG00000090862 | Rps13 | ribosomal protein S13 |
| ENSMUSG00000024735 | Prpf19 | pre-mRNA processing factor 19 |
| ENSMUSG00000024371 | C2 | complement C2 |
| ENSMUSG00000022032 | Scara5 | scavenger receptor class A, member 5 |
| ENSMUSG00000025795 | Rassf3 | Ras association (RalGDS/AF-6) domain family member 3 |
| ENSMUSG00000044934 | Zfp367 | zinc finger protein 367 |
| ENSMUSG00000047675 | Rps8 | ribosomal protein S8 |
| ENSMUSG00000054580 | Pla2r1 | phospholipase A2 receptor 1 |
| ENSMUSG00000021273 | Fdft1 | farnesyl diphosphate farnesyl transferase 1 |
| ENSMUSG00000024910 | Ctsw | cathepsin W |
| ENSMUSG00000000631 | Myo18a | myosin XVIIIA |
| ENSMUSG00000000711 | Rab5b | RAB5B, member RAS oncogene family |
| ENSMUSG00000024370 | Cdc23 | CDC23 cell division cycle 23 |
| ENSMUSG00000031389 | Arhgap4 | Rho GTPase activating protein 4 |
| ENSMUSG00000038593 | Tctn1 | tectonic family member 1 |
| ENSMUSG00000022439 | Parvg | parvin, gamma |

| | | |
|---|---|---|
| ENSMUSG00000023067 | Cdkn1a | cyclin dependent kinase inhibitor 1A |
| ENSMUSG00000046841 | Ckap4 | cytoskeleton-associated protein 4 |
| ENSMUSG00000031274 | Col4a5 | collagen, type IV, alpha 5 |
| ENSMUSG00000052406 | Rexo4 | REX4, 3'-5' exonuclease |
| ENSMUSG00000028898 | Trnau1ap | tRNA selenocysteine 1 associated protein 1 |
| ENSMUSG00000039221 | Rpl22l1 | ribosomal protein L22 like 1 |
| ENSMUSG00000029669 | Tspan12 | tetraspanin 12 |
| ENSMUSG00000070287 | Slc35g2 | solute carrier family 35, member G2 |
| ENSMUSG00000013076 | Amotl1 | angiomotin-like 1 |
| ENSMUSG00000036322 | H2-Ea | histocompatibility 2, class II antigen E alpha |
| ENSMUSG00000024743 | Syt7 | synaptotagmin VII |
| ENSMUSG00000037321 | Tap1 | transporter 1, ATP-binding cassette, sub-family B (MDR/TAP) |
| ENSMUSG00000036905 | C1qb | complement component 1, q subcomponent, beta polypeptide |
| ENSMUSG00000021266 | Wars1 | tryptophanyl-tRNA synthetase1 |
| ENSMUSG00000040883 | Tmem20 5 | transmembrane protein 205 |
| ENSMUSG00000041115 | Iqsec2 | IQ motif and Sec7 domain 2 |
| ENSMUSG00000021590 | Spata9 | spermatogenesis associated 9 |
| ENSMUSG00000060802 | B2m | beta-2 microglobulin |
| ENSMUSG00000020460 | Rps27a | ribosomal protein S27A |
| ENSMUSG00000057388 | Mrpl18 | mitochondrial ribosomal protein L18 |
| ENSMUSG00000026117 | Zap70 | zeta-chain (TCR) associated protein kinase |
| ENSMUSG00000071001 | Hrct1 | histidine rich carboxyl terminus 1 |
| ENSMUSG00000035711 | Dok3 | docking protein 3 |
| ENSMUSG00000022552 | Sharpin | SHANK-associated RH domain interacting protein |
| ENSMUSG00000033220 | Rac2 | Rac family small GTPase 2 |
| ENSMUSG00000029664 | Tfpi2 | tissue factor pathway inhibitor 2 |
| ENSMUSG00000005836 | Gata6 | GATA binding protein 6 |
| ENSMUSG00000062939 | Stat4 | signal transducer and activator of transcription 4 |
| ENSMUSG00000103409 | Lsmem2 | leucine-rich single-pass membrane protein 2 |
| ENSMUSG00000059070 | Rpl18 | ribosomal protein L18 |
| ENSMUSG00000032387 | Rbpms2 | RNA binding protein with multiple splicing 2 |
| ENSMUSG00000006589 | Aprt | adenine phosphoribosyl transferase |
| ENSMUSG00000041426 | Hibch | 3-hydroxyisobutyryl-Coenzyme A hydrolase |

**Table S6.** Gene ontology enrichment analysis of differentially expressed genes in the RV transcriptome of *P. vaccarum* that overlap with those in the RV transcriptome of high-elevation deer mice (*Peromyscus maniculatus*; Velotta et al., (2018).

| p_value | term_size | query_size | intersection_size | precision | recall | term_id | source | term_name |
|---|---|---|---|---|---|---|---|---|
| 0.01482 | 8 | 67 | 3 | 0.04478 | 0.37500 | GO:0002483 | GO:BP | antigen processing and presentation of endogenous peptide antigen |
| 0.01482 | 9 | 67 | 3 | 0.04478 | 0.33333 | GO:0019883 | GO:BP | antigen processing and presentation of endogenous antigen |
| 0.02793 | 5 | 67 | 2 | 0.02985 | 0.40000 | GO:0042613 | GO:CC | MHC class II protein complex |
| 0.02793 | 210 | 67 | 7 | 0.10448 | 0.03333 | GO:0005840 | GO:CC | ribosome |
| 0.02793 | 171 | 67 | 6 | 0.08955 | 0.03509 | GO:0030139 | GO:CC | endocytic vesicle |
| 0.02793 | 62 | 67 | 4 | 0.05970 | 0.06452 | GO:0030666 | GO:CC | endocytic vesicle membrane |
| 0.02793 | 70 | 67 | 4 | 0.05970 | 0.05714 | GO:0032040 | GO:CC | small-subunit processome |
| 0.02793 | 5 | 67 | 2 | 0.02985 | 0.40000 | GO:0042611 | GO:CC | MHC protein complex |
| 0.02793 | 7 | 67 | 2 | 0.02985 | 0.28571 | GO:0042824 | GO:CC | MHC class I peptide loading complex |
| 0.02793 | 175 | 67 | 6 | 0.08955 | 0.03429 | GO:0044391 | GO:CC | ribosomal subunit |
| 0.04631 | 38 | 67 | 3 | 0.04478 | 0.07895 | GO:0022627 | GO:CC | cytosolic small ribosomal subunit |
| 0.00699 | 10 | 67 | 3 | 0.04478 | 0.30000 | GO:0042605 | GO:MF | peptide antigen binding |
| 0.03894 | 150 | 67 | 6 | 0.08955 | 0.04000 | GO:0003735 | GO:MF | structural constituent of ribosome |
| 0.04715 | 26 | 67 | 3 | 0.04478 | 0.11538 | GO:0003823 | GO:MF | antigen binding |
| 0.00762 | 125 | 67 | 6 | 0.08955 | 0.04800 | KEGG:03010 | KEGG | Ribosome |
| 0.00762 | 169 | 67 | 7 | 0.10448 | 0.04142 | KEGG:05171 | KEGG | Coronavirus disease - COVID-19 |
| 0.02803 | 27 | 67 | 3 | 0.04478 | 0.11111 | KEGG:05150 | KEGG | Staphylococcus aureus infection |
| 0.04280 | 35 | 67 | 3 | 0.04478 | 0.08571 | KEGG:04612 | KEGG | Antigen processing and presentation |
| 0.04280 | 37 | 67 | 3 | 0.04478 | 0.08108 | KEGG:05322 | KEGG | Systemic lupus erythematosus |

**Table S7.** Expression levels of RV transcriptional modules that exhibited significant associations with Fulton's index ($P < 0.05$). (B) ANOVA between module expression and Fulton's index.

**A**

| module_ID | corr | p.val | FDRi |
|---|---|---|---|
| RV38 | -0.7897605 | 0.00657424 | 0.42188189 |
| RV19 | -0.7541901 | 0.01172674 | 0.42188189 |
| RV87 | 0.73514304 | 0.01541301 | 0.42188189 |
| RV3 | 0.70396915 | 0.02307332 | 0.42188189 |
| RV70 | -0.7029236 | 0.02336815 | 0.42188189 |
| RV71 | 0.68938098 | 0.02742437 | 0.42188189 |
| RV77 | -0.6854951 | 0.02867158 | 0.42188189 |
| RV81 | 0.66235596 | 0.0369073 | 0.44141146 |
| RV49 | 0.65059618 | 0.04164783 | 0.44141146 |
| RV2 | 0.64774578 | 0.04285548 | 0.44141146 |

**B**

| Fulton's index (FI) | | |
|---|---|---|
| **module** | **raw_FI** | **FDR_FI** |
| RV19 | 0.0011 | 0.1091 |
| RV71 | 0.0021 | 0.1091 |
| RV38 | 0.005 | 0.1347 |
| RV3 | 0.0053 | 0.1347 |
| RV87 | 0.0138 | 0.282 |
| RV49 | 0.0274 | 0.3542 |
| RV77 | 0.0284 | 0.3542 |
| RV81 | 0.0309 | 0.3542 |
| RV54 | 0.0337 | 0.3542 |
| RV70 | 0.0369 | 0.3542 |
| RV1 | 0.0382 | 0.3542 |

**Table S8.** Gene ontology enrichment analysis of RV transcriptional modules that exhibited significant associations with Fulton's index. RV49 and RV87 yielded no significant GO terms.

| query | p_value | term_size | query_size | intersection_size | precision | recall | term_id | source | term_name |
|---|---|---|---|---|---|---|---|---|---|
| RV19 | 0.01173 | 3976 | 134 | 73 | 0.54478 | 0.01836 | GO:0010467 | GO:BP | gene expression |
| RV19 | 0.01173 | 4266 | 134 | 76 | 0.56716 | 0.01782 | GO:0009059 | GO:BP | macromolecule biosynthetic process |
| RV19 | 0.01439 | 2739 | 134 | 55 | 0.41045 | 0.02008 | GO:0141187 | GO:BP | nucleic acid biosynthetic process |
| RV19 | 0.01621 | 3213 | 134 | 61 | 0.45522 | 0.01899 | GO:0090304 | GO:BP | nucleic acid metabolic process |
| RV19 | 0.01690 | 2657 | 134 | 53 | 0.39552 | 0.01995 | GO:0032774 | GO:BP | RNA biosynthetic process |
| RV19 | 0.01970 | 17 | 134 | 4 | 0.02985 | 0.23529 | GO:2000232 | GO:BP | regulation of rRNA processing |
| RV19 | 0.02348 | 2802 | 134 | 54 | 0.40299 | 0.01927 | GO:0016070 | GO:BP | RNA metabolic process |
| RV19 | 0.02348 | 2961 | 134 | 56 | 0.41791 | 0.01891 | GO:0034654 | GO:BP | nucleobase-containing compound biosynthetic process |
| RV19 | 0.02348 | 5777 | 134 | 91 | 0.67910 | 0.01575 | GO:0043170 | GO:BP | macromolecule metabolic process |
| RV19 | 0.02348 | 805 | 134 | 23 | 0.17164 | 0.02857 | GO:0045944 | GO:BP | positive regulation of transcription by RNA polymerase II |
| RV19 | 0.02746 | 408 | 134 | 15 | 0.11194 | 0.03676 | GO:0022613 | GO:BP | ribonucleoprotein complex biogenesis |
| RV19 | 0.02907 | 2 | 134 | 2 | 0.01493 | 1.00000 | GO:1905382 | GO:BP | positive regulation of snRNA transcription by RNA polymerase II |
| RV19 | 0.02907 | 2 | 134 | 2 | 0.01493 | 1.00000 | GO:1904869 | GO:BP | regulation of protein localization to Cajal body |
| RV19 | 0.02907 | 2 | 134 | 2 | 0.01493 | 1.00000 | GO:1904871 | GO:BP | positive regulation of protein localization to Cajal body |
| RV19 | 0.03124 | 3647 | 134 | 64 | 0.47761 | 0.01755 | GO:0006139 | GO:BP | nucleobase-containing compound metabolic process |
| RV19 | 0.03138 | 795 | 134 | 22 | 0.16418 | 0.02767 | GO:0006396 | GO:BP | RNA processing |
| RV19 | 0.04139 | 4977 | 134 | 80 | 0.59701 | 0.01607 | GO:0009058 | GO:BP | biosynthetic process |
| RV19 | 0.04871 | 2104 | 134 | 42 | 0.31343 | 0.01996 | GO:0051252 | GO:BP | regulation of RNA metabolic process |
| RV19 | 0.00895 | 4949 | 134 | 83 | 0.61940 | 0.01677 | GO:0005634 | GO:CC | nucleus |
| RV19 | 0.04542 | 1046 | 134 | 26 | 0.19403 | 0.02486 | GO:0140513 | GO:CC | nuclear protein-containing complex |
| RV19 | 0.00051 | 920 | 134 | 29 | 0.21642 | 0.03152 | GO:0003723 | GO:MF | RNA binding |
| RV2 | 0.00117 | 28 | 809 | 12 | 0.01483 | 0.42857 | GO:0006099 | GO:BP | tricarboxylic acid cycle |
| RV2 | 0.00723 | 370 | 809 | 52 | 0.06428 | 0.14054 | GO:0006091 | GO:BP | generation of precursor metabolites and energy |
| RV2 | 0.02685 | 34 | 809 | 11 | 0.01360 | 0.32353 | GO:0086004 | GO:BP | regulation of cardiac muscle cell contraction |

| | | | | | | | | | |
|---|---|---|---|---|---|---|---|---|---|
| RV2 | 0.02685 | 529 | 809 | 64 | 0.07911 | 0.12098 | GO:0061061 | GO:BP | muscle structure development |
| RV2 | 0.02685 | 107 | 809 | 21 | 0.02596 | 0.19626 | GO:0060048 | GO:BP | cardiac muscle contraction |
| RV2 | 0.02685 | 62 | 809 | 15 | 0.01854 | 0.24194 | GO:0055117 | GO:BP | regulation of cardiac muscle contraction |
| RV2 | 0.02837 | 274 | 809 | 39 | 0.04821 | 0.14234 | GO:0015980 | GO:BP | energy derivation by oxidation of organic compounds |
| RV2 | 0.03257 | 174 | 809 | 28 | 0.03461 | 0.16092 | GO:0060047 | GO:BP | heart contraction |
| RV2 | 0.03257 | 58 | 809 | 14 | 0.01731 | 0.24138 | GO:0086003 | GO:BP | cardiac muscle cell contraction |
| RV2 | 0.03257 | 185 | 809 | 29 | 0.03585 | 0.15676 | GO:0003015 | GO:BP | heart process |
| RV2 | 0.03257 | 242 | 809 | 35 | 0.04326 | 0.14463 | GO:0007517 | GO:BP | muscle organ development |
| RV2 | 0.03274 | 132 | 809 | 23 | 0.02843 | 0.17424 | GO:0006941 | GO:BP | striated muscle contraction |
| RV2 | 0.04057 | 40 | 809 | 11 | 0.01360 | 0.27500 | GO:1903115 | GO:BP | regulation of actin filament-based movement |
| RV2 | 0.04074 | 144 | 809 | 24 | 0.02967 | 0.16667 | GO:0008016 | GO:BP | regulation of heart contraction |
| RV2 | 0.04464 | 102 | 809 | 19 | 0.02349 | 0.18627 | GO:0008286 | GO:BP | insulin receptor signaling pathway |
| RV2 | 0.01729 | 14 | 809 | 7 | 0.00865 | 0.50000 | GO:0045239 | GO:CC | tricarboxylic acid cycle heteromeric enzyme complex |
| RV2 | 0.02694 | 17 | 809 | 7 | 0.00865 | 0.41176 | GO:0045240 | GO:CC | alpha-ketoacid dehydrogenase complex |
| RV2 | 0.02694 | 12 | 809 | 6 | 0.00742 | 0.50000 | GO:0071565 | GO:CC | nBAF complex |
| RV2 | 0.02960 | 64 | 809 | 14 | 0.01731 | 0.21875 | GO:0016529 | GO:CC | sarcoplasmic reticulum |
| RV2 | 0.01159 | 917 | 809 | 101 | 0.12485 | 0.11014 | GO:0140110 | GO:MF | transcription regulator activity |
| RV2 | 0.02266 | 744 | 809 | 83 | 0.10260 | 0.11156 | GO:0003690 | GO:MF | double-stranded DNA binding |
| RV2 | 0.02266 | 648 | 809 | 73 | 0.09023 | 0.11265 | GO:0001067 | GO:MF | transcription regulatory region nucleic acid binding |
| RV2 | 0.02266 | 522 | 809 | 61 | 0.07540 | 0.11686 | GO:0000981 | GO:MF | DNA-binding transcription factor activity, RNA polymerase II-specific |
| RV2 | 0.02266 | 644 | 809 | 72 | 0.08900 | 0.11180 | GO:0000976 | GO:MF | transcription cis-regulatory region binding |
| RV2 | 0.02266 | 554 | 809 | 65 | 0.08035 | 0.11733 | GO:0003700 | GO:MF | DNA-binding transcription factor activity |
| RV2 | 0.02266 | 757 | 809 | 82 | 0.10136 | 0.10832 | GO:0043565 | GO:MF | sequence-specific DNA binding |
| RV2 | 0.03434 | 683 | 809 | 74 | 0.09147 | 0.10835 | GO:1990837 | GO:MF | sequence-specific double-stranded DNA binding |
| RV2 | 0.03434 | 31 | 809 | 9 | 0.01112 | 0.29032 | GO:0016620 | GO:MF | oxidoreductase activity, acting on the aldehyde or oxo group of donors, NAD or NADP as acceptor |
| RV2 | 0.03434 | 570 | 809 | 64 | 0.07911 | 0.11228 | GO:0000977 | GO:MF | RNA polymerase II transcription regulatory region sequence-specific DNA binding |
| RV2 | 0.04263 | 511 | 809 | 58 | 0.07169 | 0.11350 | GO:0000987 | GO:MF | cis-regulatory region sequence-specific DNA binding |

| | | | | | | | | | |
|---|---|---|---|---|---|---|---|---|---|
| RV2 | 0.04263 | 10 | 809 | 5 | 0.00618 | 0.50000 | GO:0098847 | GO:MF | sequence-specific single stranded DNA binding |
| RV2 | 0.04780 | 493 | 809 | 56 | 0.06922 | 0.11359 | GO:0000978 | GO:MF | RNA polymerase II cis-regulatory region sequence-specific DNA binding |
| RV2 | 0.00003 | 28 | 809 | 12 | 0.01483 | 0.42857 | KEGG:00020 | KEGG | Citrate cycle (TCA cycle) |
| RV2 | 0.00003 | 27 | 809 | 12 | 0.01483 | 0.44444 | KEGG:01210 | KEGG | 2-Oxocarboxylic acid metabolism |
| RV2 | 0.00097 | 17 | 809 | 8 | 0.00989 | 0.47059 | KEGG:00785 | KEGG | Lipoic acid metabolism |
| RV2 | 0.00728 | 118 | 809 | 21 | 0.02596 | 0.17797 | KEGG:04910 | KEGG | Insulin signaling pathway |
| RV2 | 0.01196 | 98 | 809 | 18 | 0.02225 | 0.18367 | KEGG:01200 | KEGG | Carbon metabolism |
| RV3 | 0.00547 | 417 | 530 | 38 | 0.07170 | 0.09113 | GO:0000139 | GO:CC | Golgi membrane |
| RV3 | 0.00547 | 502 | 530 | 44 | 0.08302 | 0.08765 | GO:0070161 | GO:CC | anchoring junction |
| RV3 | 0.00547 | 895 | 530 | 68 | 0.12830 | 0.07598 | GO:0043005 | GO:CC | neuron projection |
| RV3 | 0.00547 | 1520 | 530 | 106 | 0.20000 | 0.06974 | GO:0042995 | GO:CC | cell projection |
| RV3 | 0.00547 | 1450 | 530 | 99 | 0.18679 | 0.06828 | GO:0120025 | GO:CC | plasma membrane bounded cell projection |
| RV3 | 0.00547 | 169 | 530 | 21 | 0.03962 | 0.12426 | GO:0030055 | GO:CC | cell-substrate junction |
| RV3 | 0.00547 | 1558 | 530 | 107 | 0.20189 | 0.06868 | GO:0030054 | GO:CC | cell junction |
| RV3 | 0.00547 | 40 | 530 | 9 | 0.01698 | 0.22500 | GO:1903293 | GO:CC | phosphatase complex |
| RV3 | 0.00547 | 40 | 530 | 9 | 0.01698 | 0.22500 | GO:0008287 | GO:CC | protein serine/threonine phosphatase complex |
| RV3 | 0.00547 | 162 | 530 | 21 | 0.03962 | 0.12963 | GO:0005925 | GO:CC | focal adhesion |
| RV3 | 0.00547 | 59 | 530 | 11 | 0.02075 | 0.18644 | GO:0005905 | GO:CC | clathrin-coated pit |
| RV3 | 0.00778 | 397 | 530 | 36 | 0.06792 | 0.09068 | GO:0015629 | GO:CC | actin cytoskeleton |
| RV3 | 0.01570 | 91 | 530 | 13 | 0.02453 | 0.14286 | GO:0042641 | GO:CC | actomyosin |
| RV3 | 0.01570 | 1038 | 530 | 73 | 0.13774 | 0.07033 | GO:0005794 | GO:CC | Golgi apparatus |
| RV3 | 0.01570 | 263 | 530 | 26 | 0.04906 | 0.09886 | GO:0098791 | GO:CC | Golgi apparatus subcompartment |
| RV3 | 0.01967 | 978 | 530 | 69 | 0.13019 | 0.07055 | GO:0031984 | GO:CC | organelle subcompartment |
| RV3 | 0.01967 | 83 | 530 | 12 | 0.02264 | 0.14458 | GO:0001725 | GO:CC | stress fiber |
| RV3 | 0.01967 | 83 | 530 | 12 | 0.02264 | 0.14458 | GO:0097517 | GO:CC | contractile actin filament bundle |
| RV3 | 0.02169 | 2895 | 530 | 170 | 0.32075 | 0.05872 | GO:0012505 | GO:CC | endomembrane system |
| RV3 | 0.02571 | 1302 | 530 | 86 | 0.16226 | 0.06605 | GO:0098588 | GO:CC | bounding membrane of organelle |
| RV3 | 0.03900 | 91 | 530 | 12 | 0.02264 | 0.13187 | GO:0032432 | GO:CC | actin filament bundle |

| | | | | | | | | | |
|---|---|---|---|---|---|---|---|---|---|
| RV3 | 0.04182 | 666 | 530 | 49 | 0.09245 | 0.07357 | GO:0036477 | GO:CC | somatodendritic compartment |
| RV3 | 0.04260 | 27 | 530 | 6 | 0.01132 | 0.22222 | GO:0051233 | GO:CC | spindle midzone |
| RV3 | 0.04542 | 603 | 530 | 45 | 0.08491 | 0.07463 | GO:0048471 | GO:CC | perinuclear region of cytoplasm |
| RV38 | 0.00300 | 27 | 75 | 4 | 0.05333 | 0.14815 | KEGG:05150 | KEGG | Staphylococcus aureus infection |
| RV70 | 0.00367 | 1450 | 51 | 19 | 0.37255 | 0.01310 | GO:0010558 | GO:BP | negative regulation of macromolecule biosynthetic process |
| RV70 | 0.00367 | 3213 | 51 | 30 | 0.58824 | 0.00934 | GO:0090304 | GO:BP | nucleic acid metabolic process |
| RV70 | 0.00367 | 579 | 51 | 12 | 0.23529 | 0.02073 | GO:0006325 | GO:BP | chromatin organization |
| RV70 | 0.00367 | 484 | 51 | 11 | 0.21569 | 0.02273 | GO:0006338 | GO:BP | chromatin remodeling |
| RV70 | 0.00376 | 250 | 51 | 8 | 0.15686 | 0.03200 | GO:0141188 | GO:BP | nucleic acid catabolic process |
| RV70 | 0.00376 | 1508 | 51 | 19 | 0.37255 | 0.01260 | GO:0009890 | GO:BP | negative regulation of biosynthetic process |
| RV70 | 0.00376 | 2802 | 51 | 27 | 0.52941 | 0.00964 | GO:0016070 | GO:BP | RNA metabolic process |
| RV70 | 0.00546 | 197 | 51 | 7 | 0.13725 | 0.03553 | GO:0006402 | GO:BP | mRNA catabolic process |
| RV70 | 0.01016 | 595 | 51 | 11 | 0.21569 | 0.01849 | GO:0000122 | GO:BP | negative regulation of transcription by RNA polymerase II |
| RV70 | 0.01017 | 602 | 51 | 11 | 0.21569 | 0.01827 | GO:0016071 | GO:BP | mRNA metabolic process |
| RV70 | 0.01035 | 229 | 51 | 7 | 0.13725 | 0.03057 | GO:0006401 | GO:BP | RNA catabolic process |
| RV70 | 0.01253 | 104 | 51 | 5 | 0.09804 | 0.04808 | GO:0000956 | GO:BP | nuclear-transcribed mRNA catabolic process |
| RV70 | 0.01253 | 3647 | 51 | 30 | 0.58824 | 0.00823 | GO:0006139 | GO:BP | nucleobase-containing compound metabolic process |
| RV70 | 0.01347 | 1736 | 51 | 19 | 0.37255 | 0.01094 | GO:0010605 | GO:BP | negative regulation of macromolecule metabolic process |
| RV70 | 0.01347 | 1902 | 51 | 20 | 0.39216 | 0.01052 | GO:0009892 | GO:BP | negative regulation of metabolic process |
| RV70 | 0.01383 | 900 | 51 | 13 | 0.25490 | 0.01444 | GO:0051253 | GO:BP | negative regulation of RNA metabolic process |
| RV70 | 0.01490 | 182 | 51 | 6 | 0.11765 | 0.03297 | GO:0040029 | GO:BP | epigenetic regulation of gene expression |
| RV70 | 0.01490 | 2104 | 51 | 21 | 0.41176 | 0.00998 | GO:0051252 | GO:BP | regulation of RNA metabolic process |
| RV70 | 0.01625 | 3004 | 51 | 26 | 0.50980 | 0.00866 | GO:0010556 | GO:BP | regulation of macromolecule biosynthetic process |
| RV70 | 0.01756 | 823 | 51 | 12 | 0.23529 | 0.01458 | GO:1902679 | GO:BP | negative regulation of RNA biosynthetic process |
| RV70 | 0.01756 | 2164 | 51 | 21 | 0.41176 | 0.00970 | GO:0010604 | GO:BP | positive regulation of macromolecule metabolic process |
| RV70 | 0.01756 | 2332 | 51 | 22 | 0.43137 | 0.00943 | GO:0019219 | GO:BP | regulation of nucleobase-containing compound metabolic process |
| RV70 | 0.01756 | 816 | 51 | 12 | 0.23529 | 0.01471 | GO:0045892 | GO:BP | negative regulation of DNA-templated transcription |
| RV70 | 0.01942 | 4266 | 51 | 32 | 0.62745 | 0.00750 | GO:0009059 | GO:BP | macromolecule biosynthetic process |

| | | | | | | | | | |
|---|---|---|---|---|---|---|---|---|---|
| RV70 | 0.02196 | 2919 | 51 | 25 | 0.49020 | 0.00856 | GO:0010468 | GO:BP | regulation of gene expression |
| RV70 | 0.02267 | 996 | 51 | 13 | 0.25490 | 0.01305 | GO:0045934 | GO:BP | negative regulation of nucleobase-containing compound metabolic process |
| RV70 | 0.02267 | 3138 | 51 | 26 | 0.50980 | 0.00829 | GO:0009889 | GO:BP | regulation of biosynthetic process |
| RV70 | 0.02267 | 7 | 51 | 2 | 0.03922 | 0.28571 | GO:0097368 | GO:BP | establishment of Sertoli cell barrier |
| RV70 | 0.02267 | 400 | 51 | 8 | 0.15686 | 0.02000 | GO:0034655 | GO:BP | nucleobase-containing compound catabolic process |
| RV70 | 0.02339 | 1443 | 51 | 16 | 0.31373 | 0.01109 | GO:0006357 | GO:BP | regulation of transcription by RNA polymerase II |
| RV70 | 0.02749 | 36 | 51 | 3 | 0.05882 | 0.08333 | GO:0000184 | GO:BP | nuclear-transcribed mRNA catabolic process, nonsense-mediated decay |
| RV70 | 0.02749 | 3976 | 51 | 30 | 0.58824 | 0.00755 | GO:0010467 | GO:BP | gene expression |
| RV70 | 0.02749 | 230 | 51 | 6 | 0.11765 | 0.02609 | GO:1903311 | GO:BP | regulation of mRNA metabolic process |
| RV70 | 0.03169 | 789 | 51 | 11 | 0.21569 | 0.01394 | GO:0010629 | GO:BP | negative regulation of gene expression |
| RV70 | 0.03376 | 1371 | 51 | 15 | 0.29412 | 0.01094 | GO:0045935 | GO:BP | positive regulation of nucleobase-containing compound metabolic process |
| RV70 | 0.03376 | 95 | 51 | 4 | 0.07843 | 0.04211 | GO:0046546 | GO:BP | development of primary male sexual characteristics |
| RV70 | 0.03376 | 94 | 51 | 4 | 0.07843 | 0.04255 | GO:0008584 | GO:BP | male gonad development |
| RV70 | 0.03376 | 1215 | 51 | 14 | 0.27451 | 0.01152 | GO:0051254 | GO:BP | positive regulation of RNA metabolic process |
| RV70 | 0.03376 | 10 | 51 | 2 | 0.03922 | 0.20000 | GO:0051451 | GO:BP | myoblast migration |
| RV70 | 0.03376 | 3285 | 51 | 26 | 0.50980 | 0.00791 | GO:0080090 | GO:BP | regulation of primary metabolic process |
| RV70 | 0.03376 | 1690 | 51 | 17 | 0.33333 | 0.01006 | GO:0010557 | GO:BP | positive regulation of macromolecule biosynthetic process |
| RV70 | 0.03376 | 1516 | 51 | 16 | 0.31373 | 0.01055 | GO:0006366 | GO:BP | transcription by RNA polymerase II |
| RV70 | 0.03578 | 2384 | 51 | 21 | 0.41176 | 0.00881 | GO:0009893 | GO:BP | positive regulation of metabolic process |
| RV70 | 0.03578 | 2739 | 51 | 23 | 0.45098 | 0.00840 | GO:0141187 | GO:BP | nucleic acid biosynthetic process |
| RV70 | 0.04312 | 3544 | 51 | 27 | 0.52941 | 0.00762 | GO:0060255 | GO:BP | regulation of macromolecule metabolic process |
| RV70 | 0.04758 | 183 | 51 | 5 | 0.09804 | 0.02732 | GO:0007548 | GO:BP | sex differentiation |
| RV70 | 0.04769 | 108 | 51 | 4 | 0.07843 | 0.03704 | GO:0046661 | GO:BP | male sex differentiation |
| RV70 | 0.01657 | 558 | 51 | 10 | 0.19608 | 0.01792 | GO:0000785 | GO:CC | chromatin |
| RV70 | 0.01657 | 991 | 51 | 14 | 0.27451 | 0.01413 | GO:0005694 | GO:CC | chromosome |
| RV70 | 0.02829 | 79 | 51 | 4 | 0.07843 | 0.05063 | GO:0010494 | GO:CC | cytoplasmic stress granule |
| RV71 | 0.00127 | 124 | 51 | 6 | 0.11765 | 0.04839 | KEGG:04814 | KEGG | Motor proteins |

| | | | | | | | | | |
|---|---|---|---|---|---|---|---|---|---|
| RV77 | 0.02398 | 5 | 45 | 2 | 0.04444 | 0.40000 | GO:0042840 | GO:BP | D-glucuronate catabolic process |
| RV77 | 0.02398 | 5 | 45 | 2 | 0.04444 | 0.40000 | GO:0042839 | GO:BP | D-glucuronate metabolic process |
| RV77 | 0.02398 | 4 | 45 | 2 | 0.04444 | 0.50000 | GO:0042732 | GO:BP | D-xylose metabolic process |
| RV77 | 0.02398 | 5 | 45 | 2 | 0.04444 | 0.40000 | GO:0031424 | GO:BP | keratinization |
| RV77 | 0.02398 | 5 | 45 | 2 | 0.04444 | 0.40000 | GO:0019640 | GO:BP | D-glucuronate catabolic process to D-xylulose 5-phosphate |
| RV77 | 0.02398 | 5 | 45 | 2 | 0.04444 | 0.40000 | GO:0019585 | GO:BP | glucuronate metabolic process |
| RV77 | 0.02398 | 5 | 45 | 2 | 0.04444 | 0.40000 | GO:0015811 | GO:BP | L-cystine transport |
| RV77 | 0.02398 | 2 | 45 | 2 | 0.04444 | 1.00000 | GO:0005997 | GO:BP | xylulose metabolic process |
| RV77 | 0.02398 | 5 | 45 | 2 | 0.04444 | 0.40000 | GO:0006063 | GO:BP | uronic acid metabolic process |
| RV77 | 0.02398 | 5 | 45 | 2 | 0.04444 | 0.40000 | GO:0006064 | GO:BP | glucuronate catabolic process |
| RV77 | 0.02990 | 6 | 45 | 2 | 0.04444 | 0.33333 | GO:1901159 | GO:BP | xylulose 5-phosphate biosynthetic process |
| RV77 | 0.02990 | 6 | 45 | 2 | 0.04444 | 0.33333 | GO:0051167 | GO:BP | xylulose 5-phosphate metabolic process |
| RV77 | 0.03688 | 233 | 45 | 6 | 0.13333 | 0.02575 | GO:0006790 | GO:BP | sulfur compound metabolic process |
| RV77 | 0.03962 | 35 | 45 | 3 | 0.06667 | 0.08571 | GO:0007622 | GO:BP | rhythmic behavior |
| RV77 | 0.04952 | 1 | 45 | 1 | 0.02222 | 1.00000 | GO:1903981 | GO:MF | enterobactin binding |
| RV77 | 0.04952 | 1 | 45 | 1 | 0.02222 | 1.00000 | GO:0180009 | GO:MF | broad specificity neutral L-amino acid:basic L-amino acid antiporter activity |
| RV77 | 0.04952 | 1 | 45 | 1 | 0.02222 | 1.00000 | GO:0098606 | GO:MF | selenocystathionine gamma-lyase activity |
| RV77 | 0.04952 | 1 | 45 | 1 | 0.02222 | 1.00000 | GO:0080146 | GO:MF | L-cysteine desulfhydrase activity |
| RV77 | 0.04952 | 43 | 45 | 3 | 0.06667 | 0.06977 | GO:0070279 | GO:MF | vitamin B6 binding |
| RV77 | 0.04952 | 1 | 45 | 1 | 0.02222 | 1.00000 | GO:0004123 | GO:MF | cystathionine gamma-lyase activity |
| RV77 | 0.04952 | 1 | 45 | 1 | 0.02222 | 1.00000 | GO:0051717 | GO:MF | inositol-1,3,4,5-tetrakisphosphate 3-phosphatase activity |
| RV77 | 0.04952 | 12 | 45 | 2 | 0.04444 | 0.16667 | GO:0004438 | GO:MF | phosphatidylinositol-3-phosphate phosphatase activity |
| RV77 | 0.04952 | 1 | 45 | 1 | 0.02222 | 1.00000 | GO:0004617 | GO:MF | phosphoglycerate dehydrogenase activity |
| RV77 | 0.04952 | 1 | 45 | 1 | 0.02222 | 1.00000 | GO:0004856 | GO:MF | D-xylulokinase activity |
| RV77 | 0.04952 | 8 | 45 | 2 | 0.04444 | 0.25000 | GO:0015643 | GO:MF | toxic substance binding |
| RV77 | 0.04952 | 1 | 45 | 1 | 0.02222 | 1.00000 | GO:0016286 | GO:MF | small conductance calcium-activated potassium channel activity |
| RV77 | 0.04952 | 23 | 45 | 2 | 0.04444 | 0.08696 | GO:0016289 | GO:MF | acyl-CoA hydrolase activity |
| RV77 | 0.04952 | 42 | 45 | 3 | 0.06667 | 0.07143 | GO:0030170 | GO:MF | pyridoxal phosphate binding |

| | | | | | | | | | |
|---|---|---|---|---|---|---|---|---|---|
| RV77 | 0.04952 | 23 | 45 | 2 | 0.04444 | 0.08696 | GO:0034593 | GO:MF | phosphatidylinositol bisphosphate phosphatase activity |
| RV77 | 0.04952 | 1 | 45 | 1 | 0.02222 | 1.00000 | GO:0044540 | GO:MF | L-cystine L-cysteine-lyase (deaminating) |
| RV77 | 0.04952 | 1 | 45 | 1 | 0.02222 | 1.00000 | GO:0047536 | GO:MF | 2-aminoadipate transaminase activity |
| RV77 | 0.04952 | 23 | 45 | 2 | 0.04444 | 0.08696 | GO:0047617 | GO:MF | fatty acyl-CoA hydrolase activity |
| RV77 | 0.04952 | 1 | 45 | 1 | 0.02222 | 1.00000 | GO:0047958 | GO:MF | glycine:2-oxoglutarate aminotransferase activity |
| RV77 | 0.04952 | 1 | 45 | 1 | 0.02222 | 1.00000 | GO:0047982 | GO:MF | homocysteine desulfhydrase activity |
| RV77 | 0.04952 | 1 | 45 | 1 | 0.02222 | 1.00000 | GO:0050038 | GO:MF | L-xylulose reductase (NADPH) activity |
| RV77 | 0.04952 | 1 | 45 | 1 | 0.02222 | 1.00000 | GO:0050094 | GO:MF | methionine-glyoxylate transaminase activity |
| RV77 | 0.04952 | 1 | 45 | 1 | 0.02222 | 1.00000 | GO:0051800 | GO:MF | phosphatidylinositol-3,4-bisphosphate 3-phosphatase activity |
| RV77 | 0.04952 | 14 | 45 | 2 | 0.04444 | 0.14286 | GO:0052744 | GO:MF | phosphatidylinositol monophosphate phosphatase activity |
| RV81 | 0.00000 | 121 | 43 | 10 | 0.23256 | 0.08264 | GO:0007596 | GO:BP | blood coagulation |
| RV81 | 0.00000 | 124 | 43 | 10 | 0.23256 | 0.08065 | GO:0007599 | GO:BP | hemostasis |
| RV81 | 0.00000 | 124 | 43 | 10 | 0.23256 | 0.08065 | GO:0050817 | GO:BP | coagulation |
| RV81 | 0.00000 | 71 | 43 | 8 | 0.18605 | 0.11268 | GO:0030168 | GO:BP | platelet activation |
| RV81 | 0.00000 | 211 | 43 | 10 | 0.23256 | 0.04739 | GO:0050878 | GO:BP | regulation of body fluid levels |
| RV81 | 0.00001 | 9 | 43 | 4 | 0.09302 | 0.44444 | GO:0072378 | GO:BP | blood coagulation, fibrin clot formation |
| RV81 | 0.00001 | 264 | 43 | 10 | 0.23256 | 0.03788 | GO:0042060 | GO:BP | wound healing |
| RV81 | 0.00001 | 12 | 43 | 4 | 0.09302 | 0.33333 | GO:0072376 | GO:BP | protein activation cascade |
| RV81 | 0.00011 | 359 | 43 | 10 | 0.23256 | 0.02786 | GO:0009611 | GO:BP | response to wounding |
| RV81 | 0.00111 | 11 | 43 | 3 | 0.06977 | 0.27273 | GO:0010572 | GO:BP | positive regulation of platelet activation |
| RV81 | 0.00167 | 39 | 43 | 4 | 0.09302 | 0.10256 | GO:0070527 | GO:BP | platelet aggregation |
| RV81 | 0.00747 | 58 | 43 | 4 | 0.09302 | 0.06897 | GO:0034109 | GO:BP | homotypic cell-cell adhesion |
| RV81 | 0.01995 | 77 | 43 | 4 | 0.09302 | 0.05195 | GO:0007229 | GO:BP | integrin-mediated signaling pathway |
| RV81 | 0.01995 | 6 | 43 | 2 | 0.04651 | 0.33333 | GO:0007597 | GO:BP | blood coagulation, intrinsic pathway |
| RV81 | 0.03254 | 37 | 43 | 3 | 0.06977 | 0.08108 | GO:0010543 | GO:BP | regulation of platelet activation |
| RV81 | 0.00282 | 15 | 43 | 3 | 0.06977 | 0.20000 | GO:0031091 | GO:CC | platelet alpha granule |
| RV81 | 0.00282 | 2 | 43 | 2 | 0.04651 | 1.00000 | GO:0070442 | GO:CC | integrin alphaIIb-beta3 complex |
| RV81 | 0.00695 | 4 | 43 | 2 | 0.04651 | 0.50000 | GO:1990779 | GO:CC | glycoprotein Ib-IX-V complex |

| | | | | | | | | | |
|---|---|---|---|---|---|---|---|---|---|
| RV81 | 0.00721 | 2927 | 43 | 23 | 0.53488 | 0.00786 | GO:0071944 | GO:CC | cell periphery |
| RV81 | 0.00906 | 222 | 43 | 6 | 0.13953 | 0.02703 | GO:0030141 | GO:CC | secretory granule |
| RV81 | 0.00913 | 232 | 43 | 6 | 0.13953 | 0.02586 | GO:0005770 | GO:CC | late endosome |
| RV81 | 0.00913 | 2644 | 43 | 21 | 0.48837 | 0.00794 | GO:0005886 | GO:CC | plasma membrane |
| RV81 | 0.01305 | 365 | 43 | 7 | 0.16279 | 0.01918 | GO:0099503 | GO:CC | secretory vesicle |
| RV81 | 0.01465 | 99 | 43 | 4 | 0.09302 | 0.04040 | GO:0098802 | GO:CC | plasma membrane signaling receptor complex |
| RV81 | 0.01773 | 1456 | 43 | 14 | 0.32558 | 0.00962 | GO:0031982 | GO:CC | vesicle |
| RV81 | 0.02872 | 125 | 43 | 4 | 0.09302 | 0.03200 | GO:0031902 | GO:CC | late endosome membrane |
| RV81 | 0.03604 | 63 | 43 | 3 | 0.06977 | 0.04762 | GO:0070062 | GO:CC | extracellular exosome |
| RV81 | 0.03879 | 19 | 43 | 2 | 0.04651 | 0.10526 | GO:0090665 | GO:CC | glycoprotein complex |
| RV81 | 0.03879 | 73 | 43 | 3 | 0.06977 | 0.04110 | GO:0065010 | GO:CC | extracellular membrane-bounded organelle |
| RV81 | 0.03879 | 73 | 43 | 3 | 0.06977 | 0.04110 | GO:0043230 | GO:CC | extracellular organelle |
| RV81 | 0.03879 | 71 | 43 | 3 | 0.06977 | 0.04225 | GO:1903561 | GO:CC | extracellular vesicle |
| RV81 | 0.03879 | 369 | 43 | 6 | 0.13953 | 0.01626 | GO:0010008 | GO:CC | endosome membrane |
| RV81 | 0.04115 | 24 | 43 | 2 | 0.04651 | 0.08333 | GO:0008305 | GO:CC | integrin complex |
| RV81 | 0.04115 | 78 | 43 | 3 | 0.06977 | 0.03846 | GO:0032587 | GO:CC | ruffle membrane |
| RV81 | 0.04115 | 1 | 43 | 1 | 0.02326 | 1.00000 | GO:0035866 | GO:CC | alphav-beta3 integrin-PKCalpha complex |
| RV81 | 0.04115 | 1 | 43 | 1 | 0.02326 | 1.00000 | GO:0061742 | GO:CC | chaperone-mediated autophagy translocation complex |
| RV81 | 0.04115 | 1 | 43 | 1 | 0.02326 | 1.00000 | GO:0071133 | GO:CC | alpha9-beta1 integrin-ADAM8 complex |
| RV81 | 0.04115 | 169 | 43 | 4 | 0.09302 | 0.02367 | GO:0030055 | GO:CC | cell-substrate junction |
| RV81 | 0.04509 | 1378 | 43 | 12 | 0.27907 | 0.00871 | GO:0031410 | GO:CC | cytoplasmic vesicle |
| RV81 | 0.04509 | 1382 | 43 | 12 | 0.27907 | 0.00868 | GO:0097708 | GO:CC | intracellular vesicle |
| RV81 | 0.02905 | 28 | 43 | 3 | 0.06977 | 0.10714 | GO:0005484 | GO:MF | SNAP receptor activity |
| RV81 | 0.02905 | 6 | 43 | 2 | 0.04651 | 0.33333 | GO:0070051 | GO:MF | fibrinogen binding |
| RV81 | 0.03597 | 8 | 43 | 2 | 0.04651 | 0.25000 | GO:0045236 | GO:MF | CXCR chemokine receptor binding |
| RV81 | 0.00009 | 94 | 43 | 6 | 0.13953 | 0.06383 | KEGG:04611 | KEGG | Platelet activation |
| RV81 | 0.00012 | 62 | 43 | 5 | 0.11628 | 0.08065 | KEGG:04512 | KEGG | ECM-receptor interaction |
| RV81 | 0.00053 | 45 | 43 | 4 | 0.09302 | 0.08889 | KEGG:04640 | KEGG | Hematopoietic cell lineage |

| | | | | | | | | | |
|---|---|---|---|---|---|---|---|---|---|
| RV81 | 0.00318 | 30 | 43 | 3 | 0.06977 | 0.10000 | KEGG:04130 | KEGG | SNARE interactions in vesicular transport |
| RV81 | 0.01537 | 55 | 43 | 3 | 0.06977 | 0.05455 | KEGG:05412 | KEGG | Arrhythmogenic right ventricular cardiomyopathy |
| RV81 | 0.02170 | 66 | 43 | 3 | 0.06977 | 0.04545 | KEGG:05410 | KEGG | Hypertrophic cardiomyopathy |
| RV81 | 0.02292 | 71 | 43 | 3 | 0.06977 | 0.04225 | KEGG:05414 | KEGG | Dilated cardiomyopathy |

**Table S9.** Genes comprising each of the RV transcriptional modules that exhibited a significant association with Fulton's index.

| Module | Gene_ID | Gene_name | description |
|---|---|---|---|
| RV19 | ENSMUSG00000071757 | Zhx2 | zinc fingers and homeoboxes 2 |
| RV19 | ENSMUSG00000037085 | Trmt12 | tRNA methyltransferase 12 |
| RV19 | ENSMUSG00000022378 | Cyrib | CYFIP related Rac1 interactor B |
| RV19 | ENSMUSG00000022620 | Arsa | arylsulfatase A |
| RV19 | ENSMUSG00000022635 | Zcrb1 | zinc finger CCHC-type and RNA binding motif 1 |
| RV19 | ENSMUSG00000033228 | Scaf11 | SR-related CTD-associated factor 11 |
| RV19 | ENSMUSG00000013495 | Tmem175 | transmembrane protein 175 |
| RV19 | ENSMUSG00000029313 | Aff1 | AF4/FMR2 family, member 1 |
| RV19 | ENSMUSG00000027304 | Rtf1 | RTF1, Paf1/RNA polymerase II complex component |
| RV19 | ENSMUSG00000027236 | Eif3j1 | eukaryotic translation initiation factor 3, subunit J1 |
| RV19 | ENSMUSG00000037110 | Ralgapa2 | Ral GTPase activating protein, alpha subunit 2 (catalytic) |
| RV19 | ENSMUSG00000027655 | Dhx35 | DEAH-box helicase 35 |
| RV19 | ENSMUSG00000090077 | Lime1 | Lck interacting transmembrane adaptor 1 |
| RV19 | ENSMUSG00000042133 | Ppig | peptidyl-prolyl isomerase G (cyclophilin G) |
| RV19 | ENSMUSG00000026944 | Abca2 | ATP-binding cassette, sub-family A member 2 |
| RV19 | ENSMUSG00000028577 | Plaa | phospholipase A2, activating protein |
| RV19 | ENSMUSG00000031545 | Gpat4 | glycerol-3-phosphate acyltransferase 4 |
| RV19 | ENSMUSG00000038291 | Snx25 | sorting nexin 25 |
| RV19 | ENSMUSG00000035790 | Cep19 | centrosomal protein 19 |
| RV19 | ENSMUSG00000050675 | Gp1ba | glycoprotein 1b, alpha polypeptide |
| RV19 | ENSMUSG00000040938 | Slc16a11 | solute carrier family 16 (monocarboxylic acid transporters), member 11 |
| RV19 | ENSMUSG00000020334 | Slc22a4 | solute carrier family 22 (organic cation transporter), member 4 |
| RV19 | ENSMUSG00000018899 | Irf1 | interferon regulatory factor 1 |
| RV19 | ENSMUSG00000042677 | Zc3h12a | zinc finger CCCH type containing 12A |
| RV19 | ENSMUSG00000042558 | Adprs | ADP-ribosylserine hydrolase |
| RV19 | ENSMUSG00000028826 | Maco1 | macoilin 1 |
| RV19 | ENSMUSG00000028943 | Espn | espin |
| RV19 | ENSMUSG00000029062 | Cdk11b | cyclin dependent kinase 11B |
| RV19 | ENSMUSG00000043241 | Upf2 | UPF2 regulator of nonsense transcripts homolog (yeast) |
| RV19 | ENSMUSG00000026743 | Mllt10 | myeloid/lymphoid or mixed-lineage leukemia; translocated to, 10 |
| RV19 | ENSMUSG00000035569 | Ankrd11 | ankyrin repeat domain 11 |
| RV19 | ENSMUSG00000024081 | Cebpz | CCAAT/enhancer binding protein zeta |
| RV19 | ENSMUSG00000024991 | Eif3a | eukaryotic translation initiation factor 3, subunit A |
| RV19 | ENSMUSG00000024976 | Shoc2 | Shoc2, leucine rich repeat scaffold protein |
| RV19 | ENSMUSG00000025047 | Pdcd11 | programmed cell death 11 |

| | | | |
|---|---|---|---|
| RV19 | ENSMUSG00000025049 | Taf5 | TATA-box binding protein associated factor 5 |
| RV19 | ENSMUSG00000069833 | Ahnak | AHNAK nucleoprotein |
| RV19 | ENSMUSG00000094936 | Rbm4 | RNA binding motif protein 4 |
| RV19 | ENSMUSG00000024831 | Ighmbp2 | immunoglobulin mu DNA binding protein 2 |
| RV19 | ENSMUSG00000025484 | Bet1l | Bet1 golgi vesicular membrane trafficking protein like |
| RV19 | ENSMUSG00000038708 | Golga4 | golgin A4 |
| RV19 | ENSMUSG00000032582 | Rbm6 | RNA binding motif protein 6 |
| RV19 | ENSMUSG00000040813 | Tex264 | testis expressed gene 264 ER-phagy receptor |
| RV19 | ENSMUSG00000032567 | Aste1 | asteroid homolog 1 |
| RV19 | ENSMUSG00000079469 | Pigb | phosphatidylinositol glycan anchor biosynthesis, class B |
| RV19 | ENSMUSG00000022119 | Rbm26 | RNA binding motif protein 26 |
| RV19 | ENSMUSG00000022099 | Dmtn | dematin actin binding protein |
| RV19 | ENSMUSG00000079184 | Mphosph8 | M-phase phosphoprotein 8 |
| RV19 | ENSMUSG00000054509 | Parp4 | poly (ADP-ribose) polymerase family, member 4 |
| RV19 | ENSMUSG00000006281 | Tep1 | telomerase associated protein 1 |
| RV19 | ENSMUSG00000021830 | Txndc16 | thioredoxin domain containing 16 |
| RV19 | ENSMUSG00000028333 | Anp32b | acidic nuclear phosphoprotein 32 family member B |
| RV19 | ENSMUSG00000027506 | Tpd52 | tumor protein D52 |
| RV19 | ENSMUSG00000067851 | Arfgef1 | ADP ribosylation factor guanine nucleotide exchange factor 1 |
| RV19 | ENSMUSG00000027599 | Armc1 | armadillo repeat containing 1 |
| RV19 | ENSMUSG00000029771 | Irf5 | interferon regulatory factor 5 |
| RV19 | ENSMUSG00000023089 | Ndufa5 | NADH:ubiquinone oxidoreductase subunit A5 |
| RV19 | ENSMUSG00000029014 | Dnajc2 | DnaJ heat shock protein family (Hsp40) member C2 |
| RV19 | ENSMUSG00000018707 | Dync1h1 | dynein cytoplasmic 1 heavy chain 1 |
| RV19 | ENSMUSG00000021244 | Ylpm1 | YLP motif containing 1 |
| RV19 | ENSMUSG00000021113 | Snapc1 | small nuclear RNA activating complex, polypeptide 1 |
| RV19 | ENSMUSG00000019996 | Map7 | microtubule-associated protein 7 |
| RV19 | ENSMUSG00000026102 | Inpp1 | inositol polyphosphate-1-phosphatase |
| RV19 | ENSMUSG00000073664 | Nbeal1 | neurobeachin like 1 |
| RV19 | ENSMUSG00000025959 | Klf7 | Kruppel-like transcription factor 7 (ubiquitous) |
| RV19 | ENSMUSG00000026205 | Slc23a3 | solute carrier family 23 (nucleobase transporters), member 3 |
| RV19 | ENSMUSG00000026197 | Zfand2b | zinc finger, AN1 type domain 2B |
| RV19 | ENSMUSG00000004364 | Cul3 | cullin 3 |
| RV19 | ENSMUSG00000047022 | Mipol1 | mirror-image polydactyly 1 |
| RV19 | ENSMUSG00000015748 | Prpf3 | pre-mRNA processing factor 3 |
| RV19 | ENSMUSG00000034427 | Myo15b | myosin XVB |
| RV19 | ENSMUSG00000078651 | Aoc2 | amine oxidase copper containing 2 |
| RV19 | ENSMUSG00000018160 | Med1 | mediator complex subunit 1 |
| RV19 | ENSMUSG00000018543 | Spmap1 | sperm microtubule associated protein 1 |

| | | | |
|---|---|---|---|
| RV19 | ENSMUSG00000020877 | Scrn2 | secernin 2 |
| RV19 | ENSMUSG00000018377 | Vezf1 | vascular endothelial zinc finger 1 |
| RV19 | ENSMUSG00000023912 | Slc25a27 | solute carrier family 25, member 27 |
| RV19 | ENSMUSG00000023932 | Cdc5l | cell division cycle 5-like |
| RV19 | ENSMUSG00000023944 | Hsp90ab1 | heat shock protein 90 alpha (cytosolic), class B member 1 |
| RV19 | ENSMUSG00000047986 | Palm3 | paralemmin 3 |
| RV19 | ENSMUSG00000056608 | Chd9 | chromodomain helicase DNA binding protein 9 |
| RV19 | ENSMUSG00000046707 | Csnk2a2 | casein kinase 2, alpha prime polypeptide |
| RV19 | ENSMUSG00000041438 | Utp4 | UTP4 small subunit processome component |
| RV19 | ENSMUSG00000047388 | Atmin | ATM interactor |
| RV19 | ENSMUSG00000024483 | Ankhd1 | ankyrin repeat and KH domain containing 1 |
| RV19 | ENSMUSG00000024350 | Dnajc18 | DnaJ heat shock protein family (Hsp40) member C18 |
| RV19 | ENSMUSG00000063281 | Zfp35 | zinc finger protein 35 |
| RV19 | ENSMUSG00000025326 | Ube3a | ubiquitin protein ligase E3A |
| RV19 | ENSMUSG00000027706 | Sec62 | SEC62 homolog, preprotein translocation |
| RV19 | ENSMUSG00000022973 | Synj1 | synaptojanin 1 |
| RV19 | ENSMUSG00000022865 | Cxadr | coxsackie virus and adenovirus receptor |
| RV19 | ENSMUSG00000026083 | Eif5b | eukaryotic translation initiation factor 5B |
| RV19 | ENSMUSG00000003226 | Ranbp2 | RAN binding protein 2 |
| RV19 | ENSMUSG00000033427 | Upb1 | ureidopropionase, beta |
| RV19 | ENSMUSG00000020265 | Sumo3 | small ubiquitin-like modifier 3 |
| RV19 | ENSMUSG00000039233 | Tbce | tubulin-specific chaperone E |
| RV19 | ENSMUSG00000046573 | Lyrm4 | LYR motif containing 4 |
| RV19 | ENSMUSG00000021428 | Riok1 | RIO kinase 1 |
| RV19 | ENSMUSG00000021494 | Ddx41 | DEAD box helicase 41 |
| RV19 | ENSMUSG00000034648 | Lrrn1 | leucine rich repeat protein 1, neuronal |
| RV19 | ENSMUSG00000035378 | Shq1 | SHQ1 homolog (S. cerevisiae) |
| RV19 | ENSMUSG00000032745 | Gpbp1 | GC-rich promoter binding protein 1 |
| RV19 | ENSMUSG00000022191 | Drosha | drosha, ribonuclease type III |
| RV19 | ENSMUSG00000002028 | Kmt2a | lysine (K)-specific methyltransferase 2A |
| RV19 | ENSMUSG00000032098 | Treh | trehalase (brush-border membrane glycoprotein) |
| RV19 | ENSMUSG00000032044 | Rpusd4 | RNA pseudouridylate synthase domain containing 4 |
| RV19 | ENSMUSG00000031939 | Taf1d | TATA-box binding protein associated factor, RNA polymerase I, D |
| RV19 | ENSMUSG00000002015 | Bcap31 | B cell receptor associated protein 31 |
| RV19 | ENSMUSG00000025347 | Tmt1b | thiol methyltransferase 1B |
| RV19 | ENSMUSG00000020024 | Cep83 | centrosomal protein 83 |
| RV19 | ENSMUSG00000036112 | Metap2 | methionine aminopeptidase 2 |
| RV19 | ENSMUSG00000059430 | Actg2 | actin, gamma 2, smooth muscle, enteric |
| RV19 | ENSMUSG00000027968 | Larp7 | La ribonucleoprotein 7, transcriptional regulator |

| | | | |
|---|---|---|---|
| RV19 | ENSMUSG00000048109 | Rbm15 | RNA binding motif protein 15 |
| RV19 | ENSMUSG00000020122 | Egfr | epidermal growth factor receptor |
| RV19 | ENSMUSG00000006005 | Tpr | translocated promoter region, nuclear basket protein |
| RV19 | ENSMUSG00000026353 | Ubxn4 | UBX domain protein 4 |
| RV19 | ENSMUSG00000049792 | Bag5 | BCL2-associated athanogene 5 |
| RV19 | ENSMUSG00000041396 | Mettl18 | methyltransferase like 18 |
| RV19 | ENSMUSG00000038095 | Sbno1 | strawberry notch 1 |
| RV19 | ENSMUSG00000047635 | Mtrfr | mitochondrial translation release factor in rescue |
| RV19 | ENSMUSG00000023106 | Denr | density-regulated protein |
| RV19 | ENSMUSG00000029726 | Mepce | methylphosphate capping enzyme |
| RV19 | ENSMUSG00000039623 | Ap5z1 | adaptor-related protein complex 5, zeta 1 subunit |
| RV19 | ENSMUSG00000029625 | Cpsf4 | cleavage and polyadenylation specific factor 4 |
| RV19 | ENSMUSG00000041420 | Meis3 | Meis homeobox 3 |
| RV19 | ENSMUSG00000001249 | Hpn | hepsin |
| RV19 | ENSMUSG00000026229 | Psmd1 | proteasome (prosome, macropain) 26S subunit, non-ATPase, 1 |
| RV19 | ENSMUSG00000026234 | Ncl | nucleolin |
| RV19 | ENSMUSG00000059263 | Usp47 | ubiquitin specific peptidase 47 |
| RV19 | ENSMUSG00000035623 | Rsf1 | remodeling and spacing factor 1 |
| RV19 | ENSMUSG00000035642 | Aamdc | adipogenesis associated Mth938 domain containing |
| RV19 | ENSMUSG00000040957 | Cables1 | CDK5 and Abl enzyme substrate 1 |
| RV19 | ENSMUSG00000022812 | Gsk3b | glycogen synthase kinase 3 beta |
| RV2 | ENSMUSG00000062397 | Zfp706 | zinc finger protein 706 |
| RV2 | ENSMUSG00000022304 | Dpys | dihydropyrimidinase |
| RV2 | ENSMUSG00000022358 | Fbxo32 | F-box protein 32 |
| RV2 | ENSMUSG00000032501 | Trib1 | tribbles pseudokinase 1 |
| RV2 | ENSMUSG00000034730 | Adgrb1 | adhesion G protein-coupled receptor B1 |
| RV2 | ENSMUSG00000075590 | Nrbp2 | nuclear receptor binding protein 2 |
| RV2 | ENSMUSG00000022558 | Mroh1 | maestro heat-like repeat family member 1 |
| RV2 | ENSMUSG00000071711 | Mpst | mercaptopyruvate sulfurtransferase |
| RV2 | ENSMUSG00000033287 | Kctd17 | potassium channel tetramerisation domain containing 17 |
| RV2 | ENSMUSG00000033128 | Gga1 | golgi associated, gamma adaptin ear containing, ARF binding protein 1 |
| RV2 | ENSMUSG00000033039 | Micall1 | microtubule associated monooxygenase, calponin and LIM domain containing -like 1 |
| RV2 | ENSMUSG00000042622 | Maff | v-maf musculoaponeurotic fibrosarcoma oncogene family, protein F (avian) |
| RV2 | ENSMUSG00000055065 | Ddx17 | DEAD box helicase 17 |
| RV2 | ENSMUSG00000042524 | Sun2 | Sad1 and UNC84 domain containing 2 |
| RV2 | ENSMUSG00000048546 | Tob2 | transducer of ERBB2, 2 |
| RV2 | ENSMUSG00000061740.2 | Cyp2d22 | cytochrome P450, family 2, subfamily d, polypeptide 22 |
| RV2 | ENSMUSG00000041852 | Tcf20 | transcription factor 20 |

| | | | |
|---|---|---|---|
| RV2 | ENSMUSG00000022383 | Ppara | peroxisome proliferator activated receptor alpha |
| RV2 | ENSMUSG00000016028 | Celsr1 | cadherin, EGF LAG seven-pass G-type receptor 1 |
| RV2 | ENSMUSG00000051864 | Tbc1d22a | TBC1 domain family, member 22a |
| RV2 | ENSMUSG00000036529 | Sbf1 | SET binding factor 1 |
| RV2 | ENSMUSG00000052369 | Tmem106c | transmembrane protein 106C |
| RV2 | ENSMUSG00000033065 | Pfkm | phosphofructokinase, muscle |
| RV2 | ENSMUSG00000022987 | Zfp641 | zinc finger protein 641 |
| RV2 | ENSMUSG00000022994 | Adcy6 | adenylate cyclase 6 |
| RV2 | ENSMUSG00000003354 | Ccdc65 | coiled-coil domain containing 65 |
| RV2 | ENSMUSG00000023018 | Smarcd1 | SWI/SNF related, matrix associated, actin dependent regulator of chromatin, subfamily d, member 1 |
| RV2 | ENSMUSG00000023019 | Gpd1 | glycerol-3-phosphate dehydrogenase 1 (soluble) |
| RV2 | ENSMUSG00000037353 | Letmd1 | LETM1 domain containing 1 |
| RV2 | ENSMUSG00000000532 | Acvr1b | activin A receptor, type 1B |
| RV2 | ENSMUSG00000036966 | Spryd3 | SPRY domain containing 3 |
| RV2 | ENSMUSG00000046897 | Zfp740 | zinc finger protein 740 |
| RV2 | ENSMUSG00000036061 | Smug1 | single-strand selective monofunctional uracil DNA glycosylase |
| RV2 | ENSMUSG00000056531 | Ccdc18 | coiled-coil domain containing 18 |
| RV2 | ENSMUSG00000034842 | Art3 | ADP-ribosyltransferase 3 |
| RV2 | ENSMUSG00000082361 | Btc | betacellulin, epidermal growth factor family member |
| RV2 | ENSMUSG00000044221 | Grsf1 | G-rich RNA sequence binding factor 1 |
| RV2 | ENSMUSG00000036285 | Noa1 | nitric oxide associated 1 |
| RV2 | ENSMUSG00000036087 | Slain2 | SLAIN motif family, member 2 |
| RV2 | ENSMUSG00000005220 | Corin | corin, serine peptidase |
| RV2 | ENSMUSG00000032724 | Abtb2 | ankyrin repeat and BTB domain containing 2 |
| RV2 | ENSMUSG00000027162 | Lin7c | lin-7 homolog C, crumbs cell polarity complex component |
| RV2 | ENSMUSG00000068614 | Actc1 | actin, alpha, cardiac muscle 1 |
| RV2 | ENSMUSG00000045838 | Ccdc9b | coiled-coil domain containing 9B |
| RV2 | ENSMUSG00000034216 | Vps18 | VPS18 CORVET/HOPS core subunit |
| RV2 | ENSMUSG00000027291 | Vps39 | VPS39 HOPS complex subunit |
| RV2 | ENSMUSG00000027246 | Ell3 | elongation factor RNA polymerase II-like 3 |
| RV2 | ENSMUSG00000091337 | Eid1 | EP300 interacting inhibitor of differentiation 1 |
| RV2 | ENSMUSG00000027394 | Ttl | tubulin tyrosine ligase |
| RV2 | ENSMUSG00000027406 | Idh3b | isocitrate dehydrogenase 3 (NAD+) beta |
| RV2 | ENSMUSG00000027300 | Ubox5 | U box domain containing 5 |
| RV2 | ENSMUSG00000027309 | Dnaaf9 | dynein axonemal assembly factor 9 |
| RV2 | ENSMUSG00000027333 | Smox | spermine oxidase |
| RV2 | ENSMUSG00000027424 | Mgme1 | mitochondrial genome maintenance exonuclease 1 |
| RV2 | ENSMUSG00000074736 | Syndig1 | synapse differentiation inducing 1 |

| | | | |
|---|---|---|---|
| RV2 | ENSMUSG00000027452 | Acss1 | acyl-CoA synthetase short-chain family member 1 |
| RV2 | ENSMUSG00000032046 | Abhd12 | abhydrolase domain containing 12 |
| RV2 | ENSMUSG00000059842 | Zfp341 | zinc finger protein 341 |
| RV2 | ENSMUSG00000074652 | Myh7b | myosin, heavy chain 7B, cardiac muscle, beta |
| RV2 | ENSMUSG00000027651 | Rprd1b | regulation of nuclear pre-mRNA domain containing 1B |
| RV2 | ENSMUSG00000017817 | Jph2 | junctophilin 2 |
| RV2 | ENSMUSG00000039804 | Ncoa5 | nuclear receptor coactivator 5 |
| RV2 | ENSMUSG00000039536 | Stau1 | staufen double-stranded RNA binding protein 1 |
| RV2 | ENSMUSG00000051149 | Adnp | activity-dependent neuroprotective protein |
| RV2 | ENSMUSG00000049999 | Ppp1r3d | protein phosphatase 1, regulatory subunit 3D |
| RV2 | ENSMUSG00000039108 | Lsm14b | LSM family member 14B |
| RV2 | ENSMUSG00000038914 | Dido1 | death inducer-obliterator 1 |
| RV2 | ENSMUSG00000016344 | Ppdpf | pancreatic progenitor cell differentiation and proliferation factor |
| RV2 | ENSMUSG00000027582 | Zgpat | zinc finger, CCCH-type with G patch domain |
| RV2 | ENSMUSG00000034075 | Zdhhc5 | zinc finger, DHHC domain containing 5 |
| RV2 | ENSMUSG00000042155 | Klhl23 | kelch-like 23 |
| RV2 | ENSMUSG00000026836 | Acvr1 | activin A receptor, type 1 |
| RV2 | ENSMUSG00000087679 | Tmem250 | transmembrane protein 250 |
| RV2 | ENSMUSG00000036281 | Snapc4 | small nuclear RNA activating complex, polypeptide 4 |
| RV2 | ENSMUSG00000026926 | Pmpca | peptidase (mitochondrial processing) alpha |
| RV2 | ENSMUSG00000052406 | Rexo4 | REX4, 3'-5' exonuclease |
| RV2 | ENSMUSG00000026917 | Wdr5 | WD repeat domain 5 |
| RV2 | ENSMUSG00000015846 | Rxra | retinoid X receptor alpha |
| RV2 | ENSMUSG00000026816 | Gtf3c5 | general transcription factor IIIC, polypeptide 5 |
| RV2 | ENSMUSG00000026858 | Miga2 | mitoguardin 2 |
| RV2 | ENSMUSG00000026848 | Tor1b | torsin family 1, member B |
| RV2 | ENSMUSG00000026820 | Ptges2 | prostaglandin E synthase 2 |
| RV2 | ENSMUSG00000026792 | Lrsam1 | leucine rich repeat and sterile alpha motif containing 1 |
| RV2 | ENSMUSG00000048351 | Coa7 | cytochrome c oxidase assembly factor 7 |
| RV2 | ENSMUSG00000043572 | Pars2 | prolyl-tRNA synthetase (mitochondrial)(putative) |
| RV2 | ENSMUSG00000028538 | St3gal3 | ST3 beta-galactoside alpha-2,3-sialyltransferase 3 |
| RV2 | ENSMUSG00000034035 | Ccdc17 | coiled-coil domain containing 17 |
| RV2 | ENSMUSG00000028700 | Pomgnt1 | protein O-linked mannose beta 1,2-N-acetylglucosaminyltransferase |
| RV2 | ENSMUSG00000039298 | Cdk5rap2 | CDK5 regulatory subunit associated protein 2 |
| RV2 | ENSMUSG00000038894 | Irs2 | insulin receptor substrate 2 |
| RV2 | ENSMUSG00000031444 | F10 | coagulation factor X |
| RV2 | ENSMUSG00000031446 | Cul4a | cullin 4A |
| RV2 | ENSMUSG00000037234 | Hook3 | hook microtubule tethering protein 3 |

| | | | |
|---|---|---|---|
| RV2 | ENSMUSG00000070044 | Fam149a | family with sequence similarity 149, member A |
| RV2 | ENSMUSG00000031636 | Pdlim3 | PDZ and LIM domain 3 |
| RV2 | ENSMUSG00000031647 | Mfap3l | microfibrillar-associated protein 3-like |
| RV2 | ENSMUSG00000004319 | Clcn3 | chloride channel, voltage-sensitive 3 |
| RV2 | ENSMUSG00000058056 | Palld | palladin, cytoskeletal associated protein |
| RV2 | ENSMUSG00000031791 | Tmem38a | transmembrane protein 38A |
| RV2 | ENSMUSG00000002393 | Nr2f6 | nuclear receptor subfamily 2, group F, member 6 |
| RV2 | ENSMUSG00000034863 | Ano8 | anoctamin 8 |
| RV2 | ENSMUSG00000031813 | Mvb12a | multivesicular body subunit 12A |
| RV2 | ENSMUSG00000071076 | Jund | jun D proto-oncogene |
| RV2 | ENSMUSG00000000325 | Brca1 | BRCA1, DNA repair associated |
| RV2 | ENSMUSG00000022768 | Ccdc116 | coiled-coil domain containing 116 |
| RV2 | ENSMUSG00000045983 | Eif4g1 | eukaryotic translation initiation factor 4, gamma 1 |
| RV2 | ENSMUSG00000033581 | Igf2bp2 | insulin-like growth factor 2 mRNA binding protein 2 |
| RV2 | ENSMUSG00000004462 | Tbccd1 | TBCC domain containing 1 |
| RV2 | ENSMUSG00000035376 | Hacd2 | 3-hydroxyacyl-CoA dehydratase 2 |
| RV2 | ENSMUSG00000022840 | Adcy5 | adenylate cyclase 5 |
| RV2 | ENSMUSG00000022905 | Kpna1 | karyopherin subunit alpha 1 |
| RV2 | ENSMUSG00000000631 | Myo18a | myosin XVIIIA |
| RV2 | ENSMUSG00000038195 | Rilp | Rab interacting lysosomal protein |
| RV2 | ENSMUSG00000000751 | Rpa1 | replication protein A1 |
| RV2 | ENSMUSG00000020741 | Cluh | clustered mitochondria homolog |
| RV2 | ENSMUSG00000020785 | Camkk1 | calcium/calmodulin-dependent protein kinase kinase 1, alpha |
| RV2 | ENSMUSG00000057778 | Cyb5d2 | cytochrome b5 domain containing 2 |
| RV2 | ENSMUSG00000040483 | Xaf1 | XIAP associated factor 1 |
| RV2 | ENSMUSG00000053574 | 4930563E22Rik | RIKEN cDNA 4930563E22 gene |
| RV2 | ENSMUSG00000040712 | Camta2 | calmodulin binding transcription activator 2 |
| RV2 | ENSMUSG00000093989 | Rnasek | ribonuclease, RNase K |
| RV2 | ENSMUSG00000018566 | Slc2a4 | solute carrier family 2 (facilitated glucose transporter), member 4 |
| RV2 | ENSMUSG00000047284 | Neurl4 | neuralized E3 ubiquitin protein ligase 4 |
| RV2 | ENSMUSG00000019461 | Plscr3 | phospholipid scramblase 3 |
| RV2 | ENSMUSG00000018765 | Fxr2 | FMR1 autosomal homolog 2 |
| RV2 | ENSMUSG00000003934 | Efnb3 | ephrin B3 |
| RV2 | ENSMUSG00000020894 | Vamp2 | vesicle-associated membrane protein 2 |
| RV2 | ENSMUSG00000045176 | Borcs6 | BLOC-1 related complex subunit 6 |
| RV2 | ENSMUSG00000018736 | Ndel1 | nudE neurodevelopment protein 1 like 1 |
| RV2 | ENSMUSG00000020902 | Ntn1 | netrin 1 |
| RV2 | ENSMUSG00000042148 | Cox10 | heme A:farnesyltransferase cytochrome c oxidase assembly factor 10 |

| | | | |
|---|---|---|---|
| RV2 | ENSMUSG00000042709 | Atpaf2 | ATP synthase mitochondrial F1 complex assembly factor 2 |
| RV2 | ENSMUSG00000020496 | Rnf187 | ring finger protein 187 |
| RV2 | ENSMUSG00000020491 | 2810021J22Rik | RIKEN cDNA 2810021J22 gene |
| RV2 | ENSMUSG00000086962 | Faxdc2 | fatty acid hydroxylase domain containing 2 |
| RV2 | ENSMUSG00000020361 | Hspa4 | heat shock protein 4 |
| RV2 | ENSMUSG00000020392 | Cdkn2aipnl | CDKN2A interacting protein N-terminal like |
| RV2 | ENSMUSG00000040283 | Btnl9 | butyrophilin-like 9 |
| RV2 | ENSMUSG00000040365 | Trim41 | tripartite motif-containing 41 |
| RV2 | ENSMUSG00000000085 | Scmh1 | sex comb on midleg homolog 1 |
| RV2 | ENSMUSG00000043207 | Zmpste24 | zinc metallopeptidase, STE24 |
| RV2 | ENSMUSG00000043962 | Thrap3 | thyroid hormone receptor associated protein 3 |
| RV2 | ENSMUSG00000028849 | Map7d1 | MAP7 domain containing 1 |
| RV2 | ENSMUSG00000003731 | Kpna6 | karyopherin subunit alpha 6 |
| RV2 | ENSMUSG00000028784 | Spocd1 | SPOC domain containing 1 |
| RV2 | ENSMUSG00000028901 | Gmeb1 | glucocorticoid modulatory element binding protein 1 |
| RV2 | ENSMUSG00000037443 | Cep85 | centrosomal protein 85 |
| RV2 | ENSMUSG00000028803 | Nipal3 | NIPA-like domain containing 3 |
| RV2 | ENSMUSG00000062157 | Ifnlr1 | interferon lambda receptor 1 |
| RV2 | ENSMUSG00000028672 | Hmgcl | 3-hydroxy-3-methylglutaryl-Coenzyme A lyase |
| RV2 | ENSMUSG00000028668 | Eloa | elongin A |
| RV2 | ENSMUSG00000051351 | Zfp46 | zinc finger protein 46 |
| RV2 | ENSMUSG00000028756 | Pink1 | PTEN induced putative kinase 1 |
| RV2 | ENSMUSG00000041143 | Tmco4 | transmembrane and coiled-coil domains 4 |
| RV2 | ENSMUSG00000028737 | Aldh4a1 | aldehyde dehydrogenase 4 family, member A1 |
| RV2 | ENSMUSG00000029020 | Mfn2 | mitofusin 2 |
| RV2 | ENSMUSG00000029019 | Nppb | natriuretic peptide type B |
| RV2 | ENSMUSG00000029016 | Clcn6 | chloride channel, voltage-sensitive 6 |
| RV2 | ENSMUSG00000028977 | Casz1 | castor zinc finger 1 |
| RV2 | ENSMUSG00000063077 | Kif1b | kinesin family member 1B |
| RV2 | ENSMUSG00000044700 | Tmem201 | transmembrane protein 201 |
| RV2 | ENSMUSG00000028976 | Slc2a5 | solute carrier family 2 (facilitated glucose transporter), member 5 |
| RV2 | ENSMUSG00000073700 | Klhl21 | kelch-like 21 |
| RV2 | ENSMUSG00000028937 | Acot7 | acyl-CoA thioesterase 7 |
| RV2 | ENSMUSG00000058498 | Rnf207 | ring finger protein 207 |
| RV2 | ENSMUSG00000039523 | Cep104 | centrosomal protein 104 |
| RV2 | ENSMUSG00000029056 | Pank4 | pantothenate kinase 4 |
| RV2 | ENSMUSG00000029073 | Cptp | ceramide-1-phosphate transfer protein |
| RV2 | ENSMUSG00000043415 | Otud1 | OTU domain containing 1 |
| RV2 | ENSMUSG00000024236 | Svil | supervillin |

| | | | |
|---|---|---|---|
| RV2 | ENSMUSG00000024283 | Wac | WW domain containing adaptor with coiled-coil |
| RV2 | ENSMUSG00000036904 | Fzd8 | frizzled class receptor 8 |
| RV2 | ENSMUSG00000031984 | 2810004N23Rik | RIKEN cDNA 2810004N23 gene |
| RV2 | ENSMUSG00000031986 | Sprtn | SprT-like N-terminal domain |
| RV2 | ENSMUSG00000000738 | Spg7 | SPG7, paraplegin matrix AAA peptidase subunit |
| RV2 | ENSMUSG00000017478 | Zc3h18 | zinc finger CCCH-type containing 18 |
| RV2 | ENSMUSG00000031816 | Mthfsd | methenyltetrahydrofolate synthetase domain containing |
| RV2 | ENSMUSG00000002228 | Ppm1j | protein phosphatase 1J |
| RV2 | ENSMUSG00000032902 | Slc16a1 | solute carrier family 16 (monocarboxylic acid transporters), member 1 |
| RV2 | ENSMUSG00000052539 | Magi3 | membrane associated guanylate kinase, WW and PDZ domain containing 3 |
| RV2 | ENSMUSG00000007379 | Dennd2c | DENN domain containing 2C |
| RV2 | ENSMUSG00000027861 | Casq2 | calsequestrin 2 |
| RV2 | ENSMUSG00000020638 | Cmpk2 | cytidine/uridine monophosphate kinase 2 |
| RV2 | ENSMUSG00000020634 | Ubxn2a | UBX domain protein 2A |
| RV2 | ENSMUSG00000020668 | Kif3c | kinesin family member 3C |
| RV2 | ENSMUSG00000029166 | Mapre3 | microtubule-associated protein, RP/EB family, member 3 |
| RV2 | ENSMUSG00000006641 | Slc5a6 | solute carrier family 5 (sodium-dependent vitamin transporter), member 6 |
| RV2 | ENSMUSG00000029134 | Plb1 | phospholipase B1 |
| RV2 | ENSMUSG00000024077 | Strn | striatin, calmodulin binding protein |
| RV2 | ENSMUSG00000035473 | Galm | galactose mutarotase |
| RV2 | ENSMUSG00000024140 | Epas1 | endothelial PAS domain protein 1 |
| RV2 | ENSMUSG00000024143 | Rhoq | ras homolog family member Q |
| RV2 | ENSMUSG00000077450 | Rab11b | RAB11B, member RAS oncogene family |
| RV2 | ENSMUSG00000024050 | Wiz | widely-interspaced zinc finger motifs |
| RV2 | ENSMUSG00000035863 | Palm | paralemmin |
| RV2 | ENSMUSG00000013833 | Med16 | mediator complex subunit 16 |
| RV2 | ENSMUSG00000013858 | Tmem259 | transmembrane protein 259 |
| RV2 | ENSMUSG00000047417 | Rexo1 | REX1, RNA exonuclease 1 |
| RV2 | ENSMUSG00000003345 | Csnk1g2 | casein kinase 1, gamma 2 |
| RV2 | ENSMUSG00000003344 | Btbd2 | BTB domain containing 2 |
| RV2 | ENSMUSG00000020190 | Mknk2 | MAP kinase-interacting serine/threonine kinase 2 |
| RV2 | ENSMUSG00000020198 | Ap3d1 | adaptor-related protein complex 3, delta 1 subunit |
| RV2 | ENSMUSG00000035278 | Plekhj1 | pleckstrin homology domain containing, family J member 1 |
| RV2 | ENSMUSG00000020211 | Sf3a2 | splicing factor 3a, subunit 2 |
| RV2 | ENSMUSG00000004937 | Sgta | small glutamine-rich tetratricopeptide repeat (TPR)-containing, alpha |
| RV2 | ENSMUSG00000034872 | Gipc3 | GIPC PDZ domain containing family, member 3 |
| RV2 | ENSMUSG00000020235 | Fzr1 | fizzy and cell division cycle 20 related 1 |
| RV2 | ENSMUSG00000043683 | Fem1a | fem 1 homolog a |

| | | | |
|---|---|---|---|
| RV2 | ENSMUSG00000041168 | Lonp1 | lon peptidase 1, mitochondrial |
| RV2 | ENSMUSG00000054723 | Vmac | vimentin-type intermediate filament associated coiled-coil protein |
| RV2 | ENSMUSG00000024206 | Rfx2 | regulatory factor X, 2 (influences HLA class II expression) |
| RV2 | ENSMUSG00000024212 | Mllt1 | myeloid/lymphoid or mixed-lineage leukemia; translocated to, 1 |
| RV2 | ENSMUSG00000046329 | Slc25a23 | solute carrier family 25 (mitochondrial carrier; phosphate carrier), member 23 |
| RV2 | ENSMUSG00000035283 | Adrb1 | adrenergic receptor, beta 1 |
| RV2 | ENSMUSG00000024978 | Gpam | glycerol-3-phosphate acyltransferase, mitochondrial |
| RV2 | ENSMUSG00000025209 | Twnk | twinkle mtDNA helicase |
| RV2 | ENSMUSG00000051984 | Sec31b | SEC31 homolog B, COPII coat complex component |
| RV2 | ENSMUSG00000040018 | Cox15 | cytochrome c oxidase assembly protein 15 |
| RV2 | ENSMUSG00000044345 | Marveld1 | MARVEL (membrane-associating) domain containing 1 |
| RV2 | ENSMUSG00000025006 | Sorbs1 | sorbin and SH3 domain containing 1 |
| RV2 | ENSMUSG00000055044 | Pdlim1 | PDZ and LIM domain 1 (elfin) |
| RV2 | ENSMUSG00000067279 | Ppp1r3c | protein phosphatase 1, regulatory subunit 3C |
| RV2 | ENSMUSG00000034459 | Ifit1 | interferon-induced protein with tetratricopeptide repeats 1 |
| RV2 | ENSMUSG00000053536 | Cstf2t | cleavage stimulation factor, 3' pre-RNA subunit 2, tau |
| RV2 | ENSMUSG00000024782 | Ak3 | adenylate kinase 3 |
| RV2 | ENSMUSG00000032702 | Kank1 | KN motif and ankyrin repeat domains 1 |
| RV2 | ENSMUSG00000024878 | Zng1 | Zn regulated GTPase metalloprotein activator 1 |
| RV2 | ENSMUSG00000074922 | Pabir1 | PP2A A alpha (PPP2R1A) and B55A (PPP2R2A) interacting phosphatase regulator 1 |
| RV2 | ENSMUSG00000024687 | Osbp | oxysterol binding protein |
| RV2 | ENSMUSG00000024735 | Prpf19 | pre-mRNA processing factor 19 |
| RV2 | ENSMUSG00000034659 | Tmem109 | transmembrane protein 109 |
| RV2 | ENSMUSG00000048832 | Vps37c | vacuolar protein sorting 37C |
| RV2 | ENSMUSG00000024969 | Mark2 | MAP/microtubule affinity regulating kinase 2 |
| RV2 | ENSMUSG00000024764 | Naa40 | N(alpha)-acetyltransferase 40, NatD catalytic subunit |
| RV2 | ENSMUSG00000024962 | Vegfb | vascular endothelial growth factor B |
| RV2 | ENSMUSG00000032648 | Pygm | muscle glycogen phosphorylase |
| RV2 | ENSMUSG00000024949 | Sf1 | splicing factor 1 |
| RV2 | ENSMUSG00000024787 | Snx15 | sorting nexin 15 |
| RV2 | ENSMUSG00000024826 | Dpf2 | double PHD fingers 2 |
| RV2 | ENSMUSG00000047658 | Gal3st3 | galactose-3-O-sulfotransferase 3 |
| RV2 | ENSMUSG00000024870 | Rab1b | RAB1B, member RAS oncogene family |
| RV2 | ENSMUSG00000024824 | Rad9a | RAD9 checkpoint clamp component A |
| RV2 | ENSMUSG00000075289 | Carns1 | carnosine synthase 1 |
| RV2 | ENSMUSG00000024913 | Lrp5 | low density lipoprotein receptor-related protein 5 |
| RV2 | ENSMUSG00000043795 | Prr33 | proline rich 33 |
| RV2 | ENSMUSG00000037887 | Dusp8 | dual specificity phosphatase 8 |

| | | | |
|---|---|---|---|
| RV2 | ENSMUSG00000025139 | Tollip | toll interacting protein |
| RV2 | ENSMUSG00000025505 | Tmem80 | transmembrane protein 80 |
| RV2 | ENSMUSG00000025480 | Syce1 | synaptonemal complex central element protein 1 |
| RV2 | ENSMUSG00000025477 | Inpp5a | inositol polyphosphate-5-phosphatase A |
| RV2 | ENSMUSG00000042828 | Trim72 | tripartite motif-containing 72 |
| RV2 | ENSMUSG00000053877 | Srcap | Snf2-related CREBBP activator protein |
| RV2 | ENSMUSG00000032637 | Atxn2l | ataxin 2-like |
| RV2 | ENSMUSG00000030889 | Vwa3a | von Willebrand factor A domain containing 3A |
| RV2 | ENSMUSG00000025239 | Limd1 | LIM domains containing 1 |
| RV2 | ENSMUSG00000032551 | 1110059G10Rik | RIKEN cDNA 1110059G10 gene |
| RV2 | ENSMUSG00000032523 | Hhatl | hedgehog acyltransferase-like |
| RV2 | ENSMUSG00000013419 | Zbtb47 | zinc finger and BTB domain containing 47 |
| RV2 | ENSMUSG00000061536 | Sec22c | SEC22 homolog C, vesicle trafficking protein |
| RV2 | ENSMUSG00000032536 | Trak1 | trafficking protein, kinesin binding 1 |
| RV2 | ENSMUSG00000032513 | Gorasp1 | golgi reassembly stacking protein 1 |
| RV2 | ENSMUSG00000032511 | Scn5a | sodium channel, voltage-gated, type V, alpha |
| RV2 | ENSMUSG00000056167 | Cnot10 | CCR4-NOT transcription complex, subunit 10 |
| RV2 | ENSMUSG00000033392 | Clasp2 | CLIP associating protein 2 |
| RV2 | ENSMUSG00000032481 | Smarcc1 | SWI/SNF related, matrix associated, actin dependent regulator of chromatin, subfamily c, member 1 |
| RV2 | ENSMUSG00000032479 | Map4 | microtubule-associated protein 4 |
| RV2 | ENSMUSG00000053646 | Plxnb1 | plexin B1 |
| RV2 | ENSMUSG00000066357 | Wdr6 | WD repeat domain 6 |
| RV2 | ENSMUSG00000006676 | Usp19 | ubiquitin specific peptidase 19 |
| RV2 | ENSMUSG00000032607 | Amt | aminomethyltransferase |
| RV2 | ENSMUSG00000032606 | Nicn1 | nicolin 1 |
| RV2 | ENSMUSG00000041528 | Rnf123 | ring finger protein 123 |
| RV2 | ENSMUSG00000103409 | Lsmem2 | leucine-rich single-pass membrane protein 2 |
| RV2 | ENSMUSG00000032579 | Hemk1 | HemK methyltransferase family member 1 |
| RV2 | ENSMUSG00000023495 | Pcbp4 | poly(rC) binding protein 4 |
| RV2 | ENSMUSG00000053716 | Dusp7 | dual specificity phosphatase 7 |
| RV2 | ENSMUSG00000032565 | Nudt16 | nudix hydrolase 16 |
| RV2 | ENSMUSG00000032531 | Amotl2 | angiomotin-like 2 |
| RV2 | ENSMUSG00000032527 | Pccb | propionyl Coenzyme A carboxylase, beta polypeptide |
| RV2 | ENSMUSG00000070287 | Slc35g2 | solute carrier family 35, member G2 |
| RV2 | ENSMUSG00000032469 | Dbr1 | debranching RNA lariats 1 |
| RV2 | ENSMUSG00000046997 | Spsb4 | splA/ryanodine receptor domain and SOCS box containing 4 |
| RV2 | ENSMUSG00000032263 | Bckdhb | branched chain ketoacid dehydrogenase E1, beta polypeptide |
| RV2 | ENSMUSG00000032352 | Lrrc1 | leucine rich repeat containing 1 |

| | | | |
|---|---|---|---|
| RV2 | ENSMUSG00000032221 | Mns1 | meiosis-specific nuclear structural protein 1 |
| RV2 | ENSMUSG00000032199 | Polr2m | polymerase (RNA) II (DNA directed) polypeptide M |
| RV2 | ENSMUSG00000032387 | Rbpms2 | RNA binding protein with multiple splicing 2 |
| RV2 | ENSMUSG00000025545 | Clybl | citrate lyase beta like |
| RV2 | ENSMUSG00000041765 | Ubac2 | ubiquitin associated domain containing 2 |
| RV2 | ENSMUSG00000022129 | Dct | dopachrome tautomerase |
| RV2 | ENSMUSG00000033060 | Lmo7 | LIM domain only 7 |
| RV2 | ENSMUSG00000035566 | Pcdh17 | protocadherin 17 |
| RV2 | ENSMUSG00000022110 | Sucla2 | succinate-Coenzyme A ligase, ADP-forming, beta subunit |
| RV2 | ENSMUSG00000022053 | Ebf2 | early B cell factor 2 |
| RV2 | ENSMUSG00000022043 | Trim35 | tripartite motif-containing 35 |
| RV2 | ENSMUSG00000022031 | Elp3 | elongator acetyltransferase complex subunit 3 |
| RV2 | ENSMUSG00000021944 | Gata4 | GATA binding protein 4 |
| RV2 | ENSMUSG00000021993 | Mipep | mitochondrial intermediate peptidase |
| RV2 | ENSMUSG00000021987 | Mtmr6 | myotubularin related protein 6 |
| RV2 | ENSMUSG00000022215 | Fitm1 | fat storage-inducing transmembrane protein 1 |
| RV2 | ENSMUSG00000040759 | Cmtm5 | CKLF-like MARVEL transmembrane domain containing 5 |
| RV2 | ENSMUSG00000089682 | Bcl2l2 | BCL2-like 2 |
| RV2 | ENSMUSG00000022179 | 4931414P19Rik | RIKEN cDNA 4931414P19 gene |
| RV2 | ENSMUSG00000016831 | Tox4 | TOX high mobility group box family member 4 |
| RV2 | ENSMUSG00000072571 | Tmem253 | transmembrane protein 253 |
| RV2 | ENSMUSG00000004558 | Ndrg2 | N-myc downstream regulated gene 2 |
| RV2 | ENSMUSG00000021831 | Ero1a | endoplasmic reticulum oxidoreductase 1 alpha |
| RV2 | ENSMUSG00000039376 | Synpo2l | synaptopodin 2-like |
| RV2 | ENSMUSG00000034235 | Usp54 | ubiquitin specific peptidase 54 |
| RV2 | ENSMUSG00000028458 | Tesk1 | testis specific protein kinase 1 |
| RV2 | ENSMUSG00000028461 | Ccdc107 | coiled-coil domain containing 107 |
| RV2 | ENSMUSG00000078716 | Tmem8b | transmembrane protein 8B |
| RV2 | ENSMUSG00000048232 | Fbxo10 | F-box protein 10 |
| RV2 | ENSMUSG00000028322 | Exosc3 | exosome component 3 |
| RV2 | ENSMUSG00000028328 | Tmod1 | tropomodulin 1 |
| RV2 | ENSMUSG00000033845 | Mrpl15 | mitochondrial ribosomal protein L15 |
| RV2 | ENSMUSG00000043542 | Zc2hc1a | zinc finger, C2HC-type containing 1A |
| RV2 | ENSMUSG00000042686 | Jph1 | junctophilin 1 |
| RV2 | ENSMUSG00000025911 | Adhfe1 | alcohol dehydrogenase, iron containing, 1 |
| RV2 | ENSMUSG00000060913 | Trim55 | tripartite motif-containing 55 |
| RV2 | ENSMUSG00000028278 | Rragd | Ras-related GTP binding D |
| RV2 | ENSMUSG00000028300 | C9orf72 | C9orf72, member of C9orf72-SMCR8 complex |
| RV2 | ENSMUSG00000028411 | Aptx | aprataxin |

| | | | |
|---|---|---|---|
| RV2 | ENSMUSG00000028444 | Cntfr | ciliary neurotrophic factor receptor |
| RV2 | ENSMUSG00000036052 | Dnajb5 | DnaJ heat shock protein family (Hsp40) member B5 |
| RV2 | ENSMUSG00000029708 | Gcc1 | golgi coiled coil 1 |
| RV2 | ENSMUSG00000029669 | Tspan12 | tetraspanin 12 |
| RV2 | ENSMUSG00000015112 | Slc25a13 | solute carrier family 25 (mitochondrial carrier, adenine nucleotide translocator), member 13 |
| RV2 | ENSMUSG00000039987 | Phtf2 | putative homeodomain transcription factor 2 |
| RV2 | ENSMUSG00000044674 | Fzd1 | frizzled class receptor 1 |
| RV2 | ENSMUSG00000037957 | Wdr20 | WD repeat domain 20 |
| RV2 | ENSMUSG00000058070 | Eml1 | echinoderm microtubule associated protein like 1 |
| RV2 | ENSMUSG00000021200 | Asb2 | ankyrin repeat and SOCS box-containing 2 |
| RV2 | ENSMUSG00000033530 | Ttc7b | tetratricopeptide repeat domain 7B |
| RV2 | ENSMUSG00000021177 | Tdp1 | tyrosyl-DNA phosphodiesterase 1 |
| RV2 | ENSMUSG00000021009 | Ptpn21 | protein tyrosine phosphatase, non-receptor type 21 |
| RV2 | ENSMUSG00000021038 | Vipas39 | VPS33B interacting protein, apical-basolateral polarity regulator, spe-39 homolog |
| RV2 | ENSMUSG00000021255 | Esrrb | estrogen related receptor, beta |
| RV2 | ENSMUSG00000021245 | Mlh3 | mutL homolog 3 |
| RV2 | ENSMUSG00000004789 | Dlst | dihydrolipoamide S-succinyltransferase |
| RV2 | ENSMUSG00000021221 | Dpf3 | double PHD fingers 3 |
| RV2 | ENSMUSG00000090935 | Synj2bp | synaptojanin 2 binding protein |
| RV2 | ENSMUSG00000032705 | Exd2 | exonuclease 3'-5' domain containing 2 |
| RV2 | ENSMUSG00000021123 | Rdh12 | retinol dehydrogenase 12 |
| RV2 | ENSMUSG00000059436 | Max | Max protein |
| RV2 | ENSMUSG00000021061 | Sptb | spectrin beta, erythrocytic |
| RV2 | ENSMUSG00000053110 | Yap1 | yes-associated protein 1 |
| RV2 | ENSMUSG00000023828 | Slc22a3 | solute carrier family 22 (organic cation transporter), member 3 |
| RV2 | ENSMUSG00000006818 | Sod2 | superoxide dismutase 2, mitochondrial |
| RV2 | ENSMUSG00000061759 | Armt1 | acidic residue methyltransferase 1 |
| RV2 | ENSMUSG00000062866 | Phactr2 | phosphatase and actin regulator 2 |
| RV2 | ENSMUSG00000039910 | Cited2 | Cbp/p300-interacting transactivator, with Glu/Asp-rich carboxy-terminal domain, 2 |
| RV2 | ENSMUSG00000019851 | Perp | PERP, TP53 apoptosis effector |
| RV2 | ENSMUSG00000019990 | Pde7b | phosphodiesterase 7B |
| RV2 | ENSMUSG00000019878 | Hsf2 | heat shock factor 2 |
| RV2 | ENSMUSG00000047514 | Tspyl1 | testis-specific protein, Y-encoded-like 1 |
| RV2 | ENSMUSG00000047139 | Cd24a | CD24a antigen |
| RV2 | ENSMUSG00000019864 | Rtn4ip1 | reticulon 4 interacting protein 1 |
| RV2 | ENSMUSG00000019848 | Popdc3 | popeye domain containing 3 |
| RV2 | ENSMUSG00000045954 | Cavin2 | caveolae associated 2 |
| RV2 | ENSMUSG00000046994 | Mars2 | methionine-tRNA synthetase 2 (mitochondrial) |

| | | | |
|---|---|---|---|
| RV2 | ENSMUSG00000025968 | Ndufs1 | NADH:ubiquinone oxidoreductase core subunit S1 |
| RV2 | ENSMUSG00000026004 | Kansl1l | KAT8 regulatory NSL complex subunit 1-like |
| RV2 | ENSMUSG00000061816 | Myl1 | myosin, light polypeptide 1 |
| RV2 | ENSMUSG00000055322 | Tns1 | tensin 1 |
| RV2 | ENSMUSG00000026180 | Cxcr2 | C-X-C motif chemokine receptor 2 |
| RV2 | ENSMUSG00000026203 | Dnajb2 | DnaJ heat shock protein family (Hsp40) member B2 |
| RV2 | ENSMUSG00000026207 | Speg | SPEG complex locus |
| RV2 | ENSMUSG00000033021 | Gmppa | GDP-mannose pyrophosphorylase A |
| RV2 | ENSMUSG00000026211 | Obsl1 | obscurin-like 1 |
| RV2 | ENSMUSG00000032883 | Acsl3 | acyl-CoA synthetase long-chain family member 3 |
| RV2 | ENSMUSG00000035181 | Heatr5a | HEAT repeat containing 5A |
| RV2 | ENSMUSG00000020954 | Strn3 | striatin, calmodulin binding protein 3 |
| RV2 | ENSMUSG00000036188 | Ankmy2 | ankyrin repeat and MYND domain containing 2 |
| RV2 | ENSMUSG00000020664.2 | Dld | dihydrolipoamide dehydrogenase |
| RV2 | ENSMUSG00000027792 | Bche | butyrylcholinesterase |
| RV2 | ENSMUSG00000034009 | Rxfp1 | relaxin/insulin-like family peptide receptor 1 |
| RV2 | ENSMUSG00000033752 | Mnd1 | meiotic nuclear divisions 1 |
| RV2 | ENSMUSG00000004897 | Hdgf | heparin binding growth factor |
| RV2 | ENSMUSG00000028070 | Naxe | NAD(P)HX epimerase |
| RV2 | ENSMUSG00000001419 | Mef2d | myocyte enhancer factor 2D |
| RV2 | ENSMUSG00000050144 | Slc25a44 | solute carrier family 25, member 44 |
| RV2 | ENSMUSG00000008604 | Ubqln4 | ubiquilin 4 |
| RV2 | ENSMUSG00000068921 | Dap3 | death associated protein 3 |
| RV2 | ENSMUSG00000041263 | Rusc1 | RUN and SH3 domain containing 1 |
| RV2 | ENSMUSG00000042784 | Muc1 | mucin 1, transmembrane |
| RV2 | ENSMUSG00000027954 | Efna1 | ephrin A1 |
| RV2 | ENSMUSG00000028042 | Zbtb7b | zinc finger and BTB domain containing 7B |
| RV2 | ENSMUSG00000038777 | Sema6c | sema domain, transmembrane domain (TM), and cytoplasmic domain, (semaphorin) 6C |
| RV2 | ENSMUSG00000053192 | Mllt11 | myeloid/lymphoid or mixed-lineage leukemia; translocated to, 11 |
| RV2 | ENSMUSG00000038526 | Car14 | carbonic anhydrase 14 |
| RV2 | ENSMUSG00000091405 | H4c14 | H4 clustered histone 14 |
| RV2 | ENSMUSG00000090210 | Itga10 | integrin, alpha 10 |
| RV2 | ENSMUSG00000028101 | Pias3 | protein inhibitor of activated STAT 3 |
| RV2 | ENSMUSG00000028093 | Acp6 | acid phosphatase 6, lysophosphatidic |
| RV2 | ENSMUSG00000038170 | Pde4dip | phosphodiesterase 4D interacting protein (myomegalin) |
| RV2 | ENSMUSG00000025175 | Fn3k | fructosamine 3 kinase |
| RV2 | ENSMUSG00000025141 | Myadml2 | myeloid-associated differentiation marker-like 2 |
| RV2 | ENSMUSG00000061111 | Mcrip1 | MAPK regulated corepressor interacting protein 1 |

| | | | |
|---|---|---|---|
| RV2 | ENSMUSG00000025583 | Rptor | regulatory associated protein of MTOR, complex 1 |
| RV2 | ENSMUSG00000039976 | Tbc1d16 | TBC1 domain family, member 16 |
| RV2 | ENSMUSG00000048277 | Syngr2 | synaptogyrin 2 |
| RV2 | ENSMUSG00000020776 | Fbf1 | Fas binding factor 1 |
| RV2 | ENSMUSG00000034341 | Wbp2 | WW domain binding protein 2 |
| RV2 | ENSMUSG00000020758 | Itgb4 | integrin beta 4 |
| RV2 | ENSMUSG00000048442 | Smim5 | small integral membrane protein 5 |
| RV2 | ENSMUSG00000034471 | Caskin2 | CASK-interacting protein 2 |
| RV2 | ENSMUSG00000044788 | Fads6 | fatty acid desaturase domain family, member 6 |
| RV2 | ENSMUSG00000041623 | Mtnap1 | mitochondrial nucleoid associated protein 1 |
| RV2 | ENSMUSG00000020719 | Ddx5 | DEAD box helicase 5 |
| RV2 | ENSMUSG00000040548 | Tex2 | testis expressed gene 2 |
| RV2 | ENSMUSG00000078622 | Ccdc47 | coiled-coil domain containing 47 |
| RV2 | ENSMUSG00000020925 | Ccdc43 | coiled-coil domain containing 43 |
| RV2 | ENSMUSG00000020923 | Ubtf | upstream binding transcription factor, RNA polymerase I |
| RV2 | ENSMUSG00000017314 | Mpp2 | membrane protein, palmitoylated 2 (MAGUK p55 subfamily member 2) |
| RV2 | ENSMUSG00000003518 | Dusp3 | dual specificity phosphatase 3 (vaccinia virus phosphatase VH1-related) |
| RV2 | ENSMUSG00000035007 | Rundc1 | RUN domain containing 1 |
| RV2 | ENSMUSG00000006920 | Ezh1 | enhancer of zeste 1 polycomb repressive complex 2 subunit |
| RV2 | ENSMUSG00000035172 | Plekhh3 | pleckstrin homology domain containing, family H (with MyTH4 domain) member 3 |
| RV2 | ENSMUSG00000019302 | Atp6v0a1 | ATPase, H+ transporting, lysosomal V0 subunit A1 |
| RV2 | ENSMUSG00000001552 | Jup | junction plakoglobin |
| RV2 | ENSMUSG00000052915 | Msl1 | male specific lethal 1 |
| RV2 | ENSMUSG00000058756 | Thra | thyroid hormone receptor alpha |
| RV2 | ENSMUSG00000038208 | Pgap3 | post-GPI attachment to proteins 3 |
| RV2 | ENSMUSG00000038216 | Pnmt | phenylethanolamine-N-methyltransferase |
| RV2 | ENSMUSG00000038485 | Socs7 | suppressor of cytokine signaling 7 |
| RV2 | ENSMUSG00000050860 | Phospho1 | phosphatase, orphan 1 |
| RV2 | ENSMUSG00000038909 | Kat7 | K(lysine) acetyltransferase 7 |
| RV2 | ENSMUSG00000001508 | Sgca | sarcoglycan, alpha (dystrophin-associated glycoprotein) |
| RV2 | ENSMUSG00000076435 | Acsf2 | acyl-CoA synthetase family member 2 |
| RV2 | ENSMUSG00000020866 | Cacna1g | calcium channel, voltage-dependent, T type, alpha 1G subunit |
| RV2 | ENSMUSG00000037573 | Tob1 | transducer of ErbB-2.1 |
| RV2 | ENSMUSG00000033983 | Coil | coilin |
| RV2 | ENSMUSG00000018428 | Akap1 | A kinase anchor protein 1 |
| RV2 | ENSMUSG00000069769 | Msi2 | musashi RNA-binding protein 2 |
| RV2 | ENSMUSG00000018378 | Cuedc1 | CUE domain containing 1 |
| RV2 | ENSMUSG00000020527 | Myo19 | myosin XIX |

| | | | |
|---|---|---|---|
| RV2 | ENSMUSG00000020684 | Rasl10b | RAS-like, family 10, member B |
| RV2 | ENSMUSG00000001143 | Lman2l | lectin, mannose-binding 2-like |
| RV2 | ENSMUSG00000010453 | Kansl3 | KAT8 regulatory NSL complex subunit 3 |
| RV2 | ENSMUSG00000023951 | Vegfa | vascular endothelial growth factor A |
| RV2 | ENSMUSG00000015605 | Srf | serum response factor |
| RV2 | ENSMUSG00000023991 | Foxp4 | forkhead box P4 |
| RV2 | ENSMUSG00000038250 | Usp38 | ubiquitin specific peptidase 38 |
| RV2 | ENSMUSG00000001909 | Trmt1 | tRNA methyltransferase 1 |
| RV2 | ENSMUSG00000004994 | Yju2b | YJU2 splicing factor homolog B |
| RV2 | ENSMUSG00000046408 | 1700067K01Rik | RIKEN cDNA 1700067K01 gene |
| RV2 | ENSMUSG00000005469 | Prkaca | protein kinase, cAMP dependent, catalytic, alpha |
| RV2 | ENSMUSG00000031703 | Itfg1 | integrin alpha FG-GAP repeat containing 1 |
| RV2 | ENSMUSG00000031778 | Cx3cl1 | C-X3-C motif chemokine ligand 1 |
| RV2 | ENSMUSG00000046556 | Zfp319 | zinc finger protein 319 |
| RV2 | ENSMUSG00000031672 | Got2 | glutamatic-oxaloacetic transaminase 2, mitochondrial |
| RV2 | ENSMUSG00000036510 | Cdh8 | cadherin 8 |
| RV2 | ENSMUSG00000060560 | Ces4a | carboxylesterase 4A |
| RV2 | ENSMUSG00000037415 | Ranbp10 | RAN binding protein 10 |
| RV2 | ENSMUSG00000036442 | Thap11 | THAP domain containing 11 |
| RV2 | ENSMUSG00000017765 | Slc12a4 | solute carrier family 12, member 4 |
| RV2 | ENSMUSG00000031921 | Terf2 | telomeric repeat binding factor 2 |
| RV2 | ENSMUSG00000033430 | Terf2ip | telomeric repeat binding factor 2, interacting protein |
| RV2 | ENSMUSG00000049090 | Ptgr3 | prostaglandin reductase 3 |
| RV2 | ENSMUSG00000059852 | Kcng2 | potassium voltage-gated channel, subfamily G, member 2 |
| RV2 | ENSMUSG00000043079 | Synpo | synaptopodin |
| RV2 | ENSMUSG00000034653 | Ythdc2 | YTH domain containing 2 |
| RV2 | ENSMUSG00000060450 | Rnf14 | ring finger protein 14 |
| RV2 | ENSMUSG00000024442 | Dele1 | DAP3 binding cell death enhancer 1 |
| RV2 | ENSMUSG00000046668 | Cxxc5 | CXXC finger 5 |
| RV2 | ENSMUSG00000073600 | Prob1 | proline rich basic protein 1 |
| RV2 | ENSMUSG00000024357 | Sil1 | SIL1 nucleotide exchange factor |
| RV2 | ENSMUSG00000024370 | Cdc23 | CDC23 cell division cycle 23 |
| RV2 | ENSMUSG00000042680 | Garem1 | GRB2 associated regulator of MAPK1 subtype 1 |
| RV2 | ENSMUSG00000036225 | Kctd1 | potassium channel tetramerisation domain containing 1 |
| RV2 | ENSMUSG00000029838 | Ptn | pleiotrophin |
| RV2 | ENSMUSG00000061758 | Akr1b10 | aldo-keto reductase family 1, member B10 |
| RV2 | ENSMUSG00000021890 | Eaf1 | ELL associated factor 1 |
| RV2 | ENSMUSG00000014496 | Ankrd28 | ankyrin repeat domain 28 |
| RV2 | ENSMUSG00000021943 | Gdf10 | growth differentiation factor 10 |

| | | | |
|---|---|---|---|
| RV2 | ENSMUSG00000021796 | Bmpr1a | bone morphogenetic protein receptor, type 1A |
| RV2 | ENSMUSG00000021779 | Thrb | thyroid hormone receptor beta |
| RV2 | ENSMUSG00000021748 | Pdhb | pyruvate dehydrogenase (lipoamide) beta |
| RV2 | ENSMUSG00000021901 | Bap1 | Brca1 associated protein 1 |
| RV2 | ENSMUSG00000021870 | Slmap | sarcolemma associated protein |
| RV2 | ENSMUSG00000001687 | Ubl3 | ubiquitin-like 3 |
| RV2 | ENSMUSG00000016520 | Lnx2 | ligand of numb-protein X 2 |
| RV2 | ENSMUSG00000005534 | Insr | insulin receptor |
| RV2 | ENSMUSG00000002949 | Timm44 | translocase of inner mitochondrial membrane 44 |
| RV2 | ENSMUSG00000038623 | Tm6sf1 | transmembrane 6 superfamily member 1 |
| RV2 | ENSMUSG00000025103 | Btbd1 | BTB domain containing 1 |
| RV2 | ENSMUSG00000038663 | Fsd2 | fibronectin type III and SPRY domain containing 2 |
| RV2 | ENSMUSG00000038930 | Rccd1 | RCC1 domain containing 1 |
| RV2 | ENSMUSG00000050973 | Gdpgp1 | GDP-D-glucose phosphorylase 1 |
| RV2 | ENSMUSG00000030539 | Sema4b | sema domain, immunoglobulin domain (Ig), transmembrane domain (TM) and short cytoplasmic domain, (semaphorin) 4B |
| RV2 | ENSMUSG00000030605 | Mfge8 | milk fat globule EGF and factor V/VIII domain containing |
| RV2 | ENSMUSG00000027796 | Smad9 | SMAD family member 9 |
| RV2 | ENSMUSG00000049504 | Proser1 | proline and serine rich 1 |
| RV2 | ENSMUSG00000049940 | Pgrmc2 | progesterone receptor membrane component 2 |
| RV2 | ENSMUSG00000037818 | Abhd18 | abhydrolase domain containing 18 |
| RV2 | ENSMUSG00000027709 | Mccc1 | methylcrotonoyl-Coenzyme A carboxylase 1 (alpha) |
| RV2 | ENSMUSG00000027668 | Mfn1 | mitofusin 1 |
| RV2 | ENSMUSG00000037730 | Mynn | myoneurin |
| RV2 | ENSMUSG00000022913 | Psmg1 | proteasome (prosome, macropain) assembly chaperone 1 |
| RV2 | ENSMUSG00000022897 | Dyrk1a | dual-specificity tyrosine phosphorylation regulated kinase 1a |
| RV2 | ENSMUSG00000040820 | Hlcs | holocarboxylase synthetase (biotin- [propriony-Coenzyme A-carboxylase (ATP-hydrolysing)] ligase) |
| RV2 | ENSMUSG00000052299 | Ltn1 | listerin E3 ubiquitin protein ligase 1 |
| RV2 | ENSMUSG00000009647 | Mcu | mitochondrial calcium uniporter |
| RV2 | ENSMUSG00000020109 | Dnajb12 | DnaJ heat shock protein family (Hsp40) member B12 |
| RV2 | ENSMUSG00000004207 | Psap | prosaposin |
| RV2 | ENSMUSG00000001665 | Gstt3 | glutathione S-transferase, theta 3 |
| RV2 | ENSMUSG00000020230 | Prmt2 | protein arginine N-methyltransferase 2 |
| RV2 | ENSMUSG00000020262 | Adarb1 | adenosine deaminase, RNA-specific, B1 |
| RV2 | ENSMUSG00000051652 | Lrrc3 | leucine rich repeat containing 3 |
| RV2 | ENSMUSG00000032834 | Pwp2 | PWP2 periodic tryptophan protein homolog (yeast) |
| RV2 | ENSMUSG00000118646 | Gm56820 | predicted gene, 56820 |
| RV2 | ENSMUSG00000024036 | Slc37a1 | solute carrier family 37 (glycerol-3-phosphate transporter), member 1 |
| RV2 | ENSMUSG00000056692 | Ilrun | inflammation and lipid regulator with UBA-like and NBR1-like domains |

| | | | |
|---|---|---|---|
| RV2 | ENSMUSG00000079605 | Zbtb9 | zinc finger and BTB domain containing 9 |
| RV2 | ENSMUSG00000007036 | Abhd16a | abhydrolase domain containing 16A |
| RV2 | ENSMUSG00000007030 | Vwa7 | von Willebrand factor A domain containing 7 |
| RV2 | ENSMUSG00000015474 | Ppt2 | palmitoyl-protein thioesterase 2 |
| RV2 | ENSMUSG00000034254 | Agpat1 | 1-acylglycerol-3-phosphate O-acyltransferase 1 |
| RV2 | ENSMUSG00000015478 | Rnf5 | ring finger protein 5 |
| RV2 | ENSMUSG00000034673 | Pbx2 | pre B cell leukemia homeobox 2 |
| RV2 | ENSMUSG00000039656 | Rxrb | retinoid X receptor beta |
| RV2 | ENSMUSG00000050705 | 2310061I04Rik | RIKEN cDNA 2310061I04 gene |
| RV2 | ENSMUSG00000003534 | Ddr1 | discoidin domain receptor family, member 1 |
| RV2 | ENSMUSG00000033799 | Tasor2 | transcription activation suppressor family member 2 |
| RV2 | ENSMUSG00000033781 | Asb13 | ankyrin repeat and SOCS box-containing 13 |
| RV2 | ENSMUSG00000021215 | Net1 | neuroepithelial cell transforming gene 1 |
| RV2 | ENSMUSG00000052565 | H1f3 | H1.3 linker histone, cluster member |
| RV2 | ENSMUSG00000021338 | Carmil1 | capping protein regulator and myosin 1 linker 1 |
| RV2 | ENSMUSG00000076431 | Sox4 | SRY (sex determining region Y)-box 4 |
| RV2 | ENSMUSG00000054889 | Dsp | desmoplakin |
| RV2 | ENSMUSG00000091264 | Smim13 | small integral membrane protein 13 |
| RV2 | ENSMUSG00000021366 | Hivep1 | human immunodeficiency virus type I enhancer binding protein 1 |
| RV2 | ENSMUSG00000038132 | Rbm24 | RNA binding motif protein 24 |
| RV2 | ENSMUSG00000038025 | Phf2 | PHD finger protein 2 |
| RV2 | ENSMUSG00000056749 | Nfil3 | nuclear factor, interleukin 3, regulated |
| RV2 | ENSMUSG00000021474 | Sfxn1 | sideroflexin 1 |
| RV2 | ENSMUSG00000025869 | Nop16 | NOP16 nucleolar protein |
| RV2 | ENSMUSG00000034928 | Rnf44 | ring finger protein 44 |
| RV2 | ENSMUSG00000007836 | Hnrnpa0 | heterogeneous nuclear ribonucleoprotein A0 |
| RV2 | ENSMUSG00000044934 | Zfp367 | zinc finger protein 367 |
| RV2 | ENSMUSG00000021483 | Cdk20 | cyclin dependent kinase 20 |
| RV2 | ENSMUSG00000016487 | Ppfibp1 | PTPRF interacting protein, binding protein 1 (liprin beta 1) |
| RV2 | ENSMUSG00000030279 | C2cd5 | C2 calcium-dependent domain containing 5 |
| RV2 | ENSMUSG00000030216 | Wbp11 | WW domain binding protein 11 |
| RV2 | ENSMUSG00000032652 | Crebl2 | cAMP responsive element binding protein-like 2 |
| RV2 | ENSMUSG00000030203 | Dusp16 | dual specificity phosphatase 16 |
| RV2 | ENSMUSG00000072704 | Smim10l1 | small integral membrane protein 10 like 1 |
| RV2 | ENSMUSG00000079304 | Tex52 | testis expressed 52 |
| RV2 | ENSMUSG00000030120 | Mlf2 | myeloid leukemia factor 2 |
| RV2 | ENSMUSG00000005069 | Pex5 | peroxisomal biogenesis factor 5 |
| RV2 | ENSMUSG00000051586 | Mical3 | microtubule associated monooxygenase, calponin and LIM domain containing 3 |

| | | | |
|---|---|---|---|
| RV2 | ENSMUSG00000058979 | Hdhd5 | haloacid dehalogenase like hydrolase domain containing 5 |
| RV2 | ENSMUSG00000030019 | Fbxl14 | F-box and leucine-rich repeat protein 14 |
| RV2 | ENSMUSG00000000441 | Raf1 | v-raf-leukemia viral oncogene 1 |
| RV2 | ENSMUSG00000093661 | Eif4e3 | eukaryotic translation initiation factor 4E member 3 |
| RV2 | ENSMUSG00000061838 | Suclg2 | succinate-Coenzyme A ligase, GDP-forming, beta subunit |
| RV2 | ENSMUSG00000028207 | Asph | aspartate-beta-hydroxylase |
| RV2 | ENSMUSG00000045205 | Dpy19l4 | dpy-19 like 4 |
| RV2 | ENSMUSG00000043252 | Tmem64 | transmembrane protein 64 |
| RV2 | ENSMUSG00000034525 | Ice1 | interactor of little elongation complex ELL subunit 1 |
| RV2 | ENSMUSG00000021604 | Irx4 | Iroquois homeobox 4 |
| RV2 | ENSMUSG00000021650 | Ptcd2 | pentatricopeptide repeat domain 2 |
| RV2 | ENSMUSG00000021643 | Serf1 | small EDRK-rich factor 1 |
| RV2 | ENSMUSG00000021645 | Smn1 | survival motor neuron 1 |
| RV2 | ENSMUSG00000055737 | Ghr | growth hormone receptor |
| RV2 | ENSMUSG00000022186 | Oxct1 | 3-oxoacid CoA transferase 1 |
| RV2 | ENSMUSG00000022201 | Zfr | zinc finger RNA binding protein |
| RV2 | ENSMUSG00000032249 | Anp32a | acidic nuclear phosphoprotein 32 family member A |
| RV2 | ENSMUSG00000032289 | Thsd4 | thrombospondin, type I, domain containing 4 |
| RV2 | ENSMUSG00000032338 | Hcn4 | hyperpolarization-activated, cyclic nucleotide-gated K+ 4 |
| RV2 | ENSMUSG00000032306 | Mpi | mannose phosphate isomerase |
| RV2 | ENSMUSG00000037493 | Cib2 | calcium and integrin binding family member 2 |
| RV2 | ENSMUSG00000041268 | Dmxl2 | Dmx-like 2 |
| RV2 | ENSMUSG00000032050 | Rdx | radixin |
| RV2 | ENSMUSG00000037112 | Sik2 | salt inducible kinase 2 |
| RV2 | ENSMUSG00000000168 | Dlat | dihydrolipoamide S-acetyltransferase |
| RV2 | ENSMUSG00000039542 | Ncam1 | neural cell adhesion molecule 1 |
| RV2 | ENSMUSG00000032264 | Zw10 | zw10 kinetochore protein |
| RV2 | ENSMUSG00000032267 | Usp28 | ubiquitin specific peptidase 28 |
| RV2 | ENSMUSG00000048537 | Phldb1 | pleckstrin homology like domain, family B, member 1 |
| RV2 | ENSMUSG00000032126 | Hmbs | hydroxymethylbilane synthase |
| RV2 | ENSMUSG00000032119 | Hinfp | histone H4 transcription factor |
| RV2 | ENSMUSG00000032010 | Usp2 | ubiquitin specific peptidase 2 |
| RV2 | ENSMUSG00000031993 | Snx19 | sorting nexin 19 |
| RV2 | ENSMUSG00000032194 | Kank2 | KN motif and ankyrin repeat domains 2 |
| RV2 | ENSMUSG00000048429 | Timm29 | translocase of inner mitochondrial membrane 29 |
| RV2 | ENSMUSG00000032185 | Carm1 | coactivator-associated arginine methyltransferase 1 |
| RV2 | ENSMUSG00000032177 | Pde4a | phosphodiesterase 4A, cAMP specific |
| RV2 | ENSMUSG00000003299 | Mrpl4 | mitochondrial ribosomal protein L4 |
| RV2 | ENSMUSG00000004100 | Ppan | peter pan homolog |

| | | | |
|---|---|---|---|
| RV2 | ENSMUSG00000079084 | Ccdc82 | coiled-coil domain containing 82 |
| RV2 | ENSMUSG00000009995 | Tafazzin | tafazzin, phospholipid-lysophospholipid transacylase |
| RV2 | ENSMUSG00000002010 | Idh3g | isocitrate dehydrogenase 3 (NAD+), gamma |
| RV2 | ENSMUSG00000002007 | Srpk3 | serine/arginine-rich protein specific kinase 3 |
| RV2 | ENSMUSG00000045237 | Eola1 | endothelium and lymphocyte associated ASCH domain 1 |
| RV2 | ENSMUSG00000031137 | Fgf13 | fibroblast growth factor 13 |
| RV2 | ENSMUSG00000060681 | Slc9a6 | solute carrier family 9 (sodium/hydrogen exchanger), member 6 |
| RV2 | ENSMUSG00000031111 | Igsf1 | immunoglobulin superfamily, member 1 |
| RV2 | ENSMUSG00000036959 | Bcorl1 | BCL6 co-repressor-like 1 |
| RV2 | ENSMUSG00000001986 | Gria3 | glutamate receptor, ionotropic, AMPA3 (alpha 3) |
| RV2 | ENSMUSG00000042903 | Foxo4 | forkhead box O4 |
| RV2 | ENSMUSG00000060890 | Arr3 | arrestin 3, retinal |
| RV2 | ENSMUSG00000031299 | Pdha1 | pyruvate dehydrogenase E1 alpha 1 |
| RV2 | ENSMUSG00000031438 | Rnf128 | ring finger protein 128 |
| RV2 | ENSMUSG00000031274 | Col4a5 | collagen, type IV, alpha 5 |
| RV2 | ENSMUSG00000006423 | Steep1 | STING1 ER exit protein 1 |
| RV2 | ENSMUSG00000025266 | Gnl3l | guanine nucleotide binding protein nucleolar 3 like |
| RV2 | ENSMUSG00000039556 | Ppp1r3f | protein phosphatase 1, regulatory subunit 3F |
| RV2 | ENSMUSG00000031161 | Hdac6 | histone deacetylase 6 |
| RV2 | ENSMUSG00000031166 | Wdr13 | WD repeat domain 13 |
| RV2 | ENSMUSG00000058254 | Tspan7 | tetraspanin 7 |
| RV2 | ENSMUSG00000025037 | Maoa | monoamine oxidase A |
| RV2 | ENSMUSG00000031065 | Cdk16 | cyclin dependent kinase 16 |
| RV2 | ENSMUSG00000001127 | Araf | Araf proto-oncogene, serine/threonine kinase |
| RV2 | ENSMUSG00000009406 | Elk1 | ELK1, member of ETS oncogene family |
| RV2 | ENSMUSG00000025348 | Itga7 | integrin alpha 7 |
| RV2 | ENSMUSG00000000711 | Rab5b | RAB5B, member RAS oncogene family |
| RV2 | ENSMUSG00000025364 | Pa2g4 | proliferation-associated 2G4 |
| RV2 | ENSMUSG00000025369 | Smarcc2 | SWI/SNF related, matrix associated, actin dependent regulator of chromatin, subfamily c, member 2 |
| RV2 | ENSMUSG00000005683 | Cs | citrate synthase |
| RV2 | ENSMUSG00000039994 | Timeless | timeless circadian clock 1 |
| RV2 | ENSMUSG00000025393 | Atp5f1b | ATP synthase F1 subunit beta |
| RV2 | ENSMUSG00000025404 | R3hdm2 | R3H domain containing 2 |
| RV2 | ENSMUSG00000116429 | Ddit3 | DNA-damage inducible transcript 3 |
| RV2 | ENSMUSG00000025795 | Rassf3 | Ras association (RalGDS/AF-6) domain family member 3 |
| RV2 | ENSMUSG00000020205 | Phlda1 | pleckstrin homology like domain, family A, member 1 |
| RV2 | ENSMUSG00000046567 | Rlig1 | RNA 5'-phosphate and 3'-OH ligase 1 |
| RV2 | ENSMUSG00000019948 | Actr6 | ARP6 actin-related protein 6 |

| | | | |
|---|---|---|---|
| RV2 | ENSMUSG00000020032 | Nuak1 | NUAK family, SNF1-like kinase, 1 |
| RV2 | ENSMUSG00000035620 | Ric8b | RIC8 guanine nucleotide exchange factor B |
| RV2 | ENSMUSG00000020044 | Timp3 | tissue inhibitor of metalloproteinase 3 |
| RV2 | ENSMUSG00000074794 | Arrdc3 | arrestin domain containing 3 |
| RV2 | ENSMUSG00000017756 | Slc12a7 | solute carrier family 12, member 7 |
| RV2 | ENSMUSG00000021577 | Sdha | succinate dehydrogenase complex, subunit A, flavoprotein (Fp) |
| RV2 | ENSMUSG00000021579 | Lrrc14b | leucine rich repeat containing 14B |
| RV2 | ENSMUSG00000051695 | Pcbp1 | poly(rC) binding protein 1 |
| RV2 | ENSMUSG00000030008 | Pradc1 | protease-associated domain containing 1 |
| RV2 | ENSMUSG00000068323 | Slc4a5 | solute carrier family 4, sodium bicarbonate cotransporter, member 5 |
| RV2 | ENSMUSG00000031865 | Dctn1 | dynactin 1 |
| RV2 | ENSMUSG00000038319 | Kcnh2 | potassium voltage-gated channel, subfamily H (eag-related), member 2 |
| RV2 | ENSMUSG00000028973 | Abcb8 | ATP-binding cassette, sub-family B member 8 |
| RV2 | ENSMUSG00000028949 | Smarcd3 | SWI/SNF related, matrix associated, actin dependent regulator of chromatin, subfamily d, member 3 |
| RV2 | ENSMUSG00000039000 | Ube3c | ubiquitin protein ligase E3C |
| RV2 | ENSMUSG00000028986 | Klhl7 | kelch-like 7 |
| RV2 | ENSMUSG00000004980 | Hnrnpa2b1 | heterogeneous nuclear ribonucleoprotein A2/B1 |
| RV2 | ENSMUSG00000005225 | Plekha8 | pleckstrin homology domain containing, family A (phosphoinositide binding specific) member 8 |
| RV2 | ENSMUSG00000029910 | Mad2l1 | MAD2 mitotic arrest deficient-like 1 |
| RV2 | ENSMUSG00000053460 | Ggcx | gamma-glutamyl carboxylase |
| RV2 | ENSMUSG00000028199 | Cryz | crystallin, zeta |
| RV2 | ENSMUSG00000027984 | Hadh | hydroxyacyl-Coenzyme A dehydrogenase |
| RV2 | ENSMUSG00000033400 | Agl | amylo-1,6-glucosidase, 4-alpha-glucanotransferase |
| RV2 | ENSMUSG00000000340 | Dbt | dihydrolipoamide branched chain transacylase E2 |
| RV2 | ENSMUSG00000027981 | Rnpc3 | RNA-binding region (RNP1, RRM) containing 3 |
| RV2 | ENSMUSG00000058135 | Gstm1 | glutathione S-transferase, mu 1 |
| RV2 | ENSMUSG00000027893 | Ahcyl1 | S-adenosylhomocysteine hydrolase-like 1 |
| RV2 | ENSMUSG00000020305 | Asb3 | ankyrin repeat and SOCS box-containing 3 |
| RV2 | ENSMUSG00000020315 | Sptbn1 | spectrin beta, non-erythrocytic 1 |
| RV2 | ENSMUSG00000004018 | Fancl | Fanconi anemia, complementation group L |
| RV2 | ENSMUSG00000078970 | Dnaaf10 | dynein axonemal assembly factor 10 |
| RV2 | ENSMUSG00000040978 | Gm11992 | predicted gene 11992 |
| RV2 | ENSMUSG00000020456 | Ogdh | oxoglutarate (alpha-ketoglutarate) dehydrogenase (lipoamide) |
| RV2 | ENSMUSG00000004394 | Tmed4 | transmembrane p24 trafficking protein 4 |
| RV2 | ENSMUSG00000004393 | Ddx56 | DEAD box helicase 56 |
| RV2 | ENSMUSG00000096923 | Gm3764 | predicted gene 3764 |
| RV2 | ENSMUSG00000053838 | Nudcd3 | NudC domain containing 3 |
| RV2 | ENSMUSG00000020475 | Pgam2 | phosphoglycerate mutase 2 |

| | | | |
|---|---|---|---|
| RV2 | ENSMUSG00000049680 | Urgcp | upregulator of cell proliferation |
| RV2 | ENSMUSG00000020482 | Ccdc117 | coiled-coil domain containing 117 |
| RV2 | ENSMUSG00000009090 | Ap1b1 | adaptor protein complex AP-1, beta 1 subunit |
| RV2 | ENSMUSG00000020412 | Ascc2 | activating signal cointegrator 1 complex subunit 2 |
| RV2 | ENSMUSG00000034354 | Mtmr3 | myotubularin related protein 3 |
| RV2 | ENSMUSG00000020435 | Osbp2 | oxysterol binding protein 2 |
| RV2 | ENSMUSG00000020439 | Smtn | smoothelin |
| RV2 | ENSMUSG00000019295 | Tmem129 | transmembrane protein 129 |
| RV2 | ENSMUSG00000109572 | Cfap99 | cilia and flagella associated protein 99 |
| RV2 | ENSMUSG00000044716 | Dok7 | docking protein 7 |
| RV2 | ENSMUSG00000029190 | D5Ertd579e | DNA segment, Chr 5, ERATO Doi 579, expressed |
| RV2 | ENSMUSG00000039474 | Wfs1 | wolframin ER transmembrane glycoprotein |
| RV2 | ENSMUSG00000029123 | Stk32b | serine/threonine kinase 32B |
| RV2 | ENSMUSG00000033722 | BC034090 | cDNA sequence BC034090 |
| RV2 | ENSMUSG00000073557 | Ppp1r12b | protein phosphatase 1, regulatory subunit 12B |
| RV2 | ENSMUSG00000042429 | Adora1 | adenosine A1 receptor |
| RV2 | ENSMUSG00000042046 | Dstyk | dual serine/threonine and tyrosine protein kinase |
| RV2 | ENSMUSG00000016528 | Mapkapk2 | MAP kinase-activated protein kinase 2 |
| RV2 | ENSMUSG00000026409 | Pfkfb2 | 6-phosphofructo-2-kinase/fructose-2,6-biphosphatase 2 |
| RV2 | ENSMUSG00000026342 | Slc35f5 | solute carrier family 35, member F5 |
| RV2 | ENSMUSG00000064302 | Clasp1 | CLIP associating protein 1 |
| RV2 | ENSMUSG00000009905 | Kdsr | 3-ketodihydrosphingosine reductase |
| RV2 | ENSMUSG00000015579 | Nkx2-5 | NK2 homeobox 5 |
| RV2 | ENSMUSG00000037098 | Rab11fip3 | RAB11 family interacting protein 3 (class II) |
| RV2 | ENSMUSG00000073434 | Wdr90 | WD repeat domain 90 |
| RV2 | ENSMUSG00000025733 | Rhot2 | ras homolog family member T2 |
| RV2 | ENSMUSG00000002280 | Ciao3 | cytosolic iron-sulfur assembly component 3 |
| RV2 | ENSMUSG00000002279 | Lmf1 | lipase maturation factor 1 |
| RV2 | ENSMUSG00000024142 | Mlst8 | MTOR associated protein, LST8 homolog (S. cerevisiae) |
| RV2 | ENSMUSG00000004069 | Dnaja3 | DnaJ heat shock protein family (Hsp40) member A3 |
| RV2 | ENSMUSG00000004071 | Cdip1 | cell death inducing Trp53 target 1 |
| RV2 | ENSMUSG00000022517 | Mgrn1 | mahogunin, ring finger 1 |
| RV2 | ENSMUSG00000068663 | Clec16a | C-type lectin domain family 16, member A |
| RV2 | ENSMUSG00000037972 | Snn | stannin |
| RV2 | ENSMUSG00000022677 | Cep20 | centrosomal protein 20 |
| RV2 | ENSMUSG00000006356 | Crip2 | cysteine rich protein 2 |
| RV2 | ENSMUSG00000021144 | Mta1 | metastasis associated 1 |
| RV2 | ENSMUSG00000072825 | Cep170b | centrosomal protein 170B |
| RV2 | ENSMUSG00000037686 | Aspg | asparaginase |

| | | | |
|---|---|---|---|
| RV2 | ENSMUSG00000001270 | Ckb | creatine kinase, brain |
| RV2 | ENSMUSG00000021281 | Tnfaip2 | tumor necrosis factor, alpha-induced protein 2 |
| RV2 | ENSMUSG00000021279 | Cdc42bpb | CDC42 binding protein kinase beta |
| RV2 | ENSMUSG00000056050 | Mia3 | MIA SH3 domain ER export factor 3 |
| RV2 | ENSMUSG00000026520 | Pycr2 | pyrroline-5-carboxylate reductase family, member 2 |
| RV2 | ENSMUSG00000010609 | Psen2 | presenilin 2 |
| RV2 | ENSMUSG00000003464 | Pex19 | peroxisomal biogenesis factor 19 |
| RV2 | ENSMUSG00000005674 | Tomm40l | translocase of outer mitochondrial membrane 40-like |
| RV2 | ENSMUSG00000058076 | Sdhc | succinate dehydrogenase complex, subunit C, integral membrane protein |
| RV2 | ENSMUSG00000040723 | Rcsd1 | RCSD domain containing 1 |
| RV2 | ENSMUSG00000029432 | Nipsnap2 | nipsnap homolog 2 |
| RV2 | ENSMUSG00000029392 | Rilpl1 | Rab interacting lysosomal protein-like 1 |
| RV2 | ENSMUSG00000029402 | Snrnp35 | small nuclear ribonucleoprotein 35 (U11/U12) |
| RV2 | ENSMUSG00000049327 | Kmt5a | lysine methyltransferase 5A |
| RV2 | ENSMUSG00000038342 | Mlxip | MLX interacting protein |
| RV2 | ENSMUSG00000029467 | Atp2a2 | ATPase, Ca++ transporting, cardiac muscle, slow twitch 2 |
| RV2 | ENSMUSG00000042605 | Atxn2 | ataxin 2 |
| RV2 | ENSMUSG00000043733 | Ptpn11 | protein tyrosine phosphatase, non-receptor type 11 |
| RV2 | ENSMUSG00000018263 | Tbx5 | T-box 5 |
| RV2 | ENSMUSG00000002486 | Tchp | trichoplein, keratin filament binding |
| RV2 | ENSMUSG00000042010 | Acacb | acetyl-Coenzyme A carboxylase beta |
| RV2 | ENSMUSG00000029591 | Ung | uracil DNA glycosylase |
| RV2 | ENSMUSG00000042216 | Sgsm1 | small G protein signaling modulator 1 |
| RV2 | ENSMUSG00000029345 | Tfip11 | tuftelin interacting protein 11 |
| RV2 | ENSMUSG00000061979 | Rcc1l | reculator of chromosome condensation 1 like |
| RV2 | ENSMUSG00000060261 | Gtf2i | general transcription factor II I |
| RV2 | ENSMUSG00000004951 | Hspb1 | heat shock protein 1 |
| RV2 | ENSMUSG00000023328 | Ache | acetylcholinesterase |
| RV2 | ENSMUSG00000051502 | Ufsp1 | UFM1-specific peptidase 1 |
| RV2 | ENSMUSG00000036980 | Taf6 | TATA-box binding protein associated factor 6 |
| RV2 | ENSMUSG00000037017 | Zscan21 | zinc finger and SCAN domain containing 21 |
| RV2 | ENSMUSG00000056014 | A430033K04Rik | RIKEN cDNA A430033K04 gene |
| RV2 | ENSMUSG00000018143 | Mafk | v-maf musculoaponeurotic fibrosarcoma oncogene family, protein K (avian) |
| RV2 | ENSMUSG00000046658 | Zfp316 | zinc finger protein 316 |
| RV2 | ENSMUSG00000038970 | Lmtk2 | lemur tyrosine kinase 2 |
| RV2 | ENSMUSG00000007564 | Ppp2r1a | protein phosphatase 2, regulatory subunit A, alpha |
| RV2 | ENSMUSG00000035545 | Leng8 | leukocyte receptor cluster (LRC) member 8 |
| RV2 | ENSMUSG00000006154 | Eps8l1 | EPS8-like 1 |

| | | | |
|---|---|---|---|
| RV2 | ENSMUSG00000019254 | Ppp1r12c | protein phosphatase 1, regulatory subunit 12C |
| RV2 | ENSMUSG00000061374 | Fiz1 | Flt3 interacting zinc finger protein 1 |
| RV2 | ENSMUSG00000043290 | Zfp784 | zinc finger protein 784 |
| RV2 | ENSMUSG00000030435 | U2af2 | U2 small nuclear ribonucleoprotein auxiliary factor (U2AF) 2 |
| RV2 | ENSMUSG00000044876 | Zfp444 | zinc finger protein 444 |
| RV2 | ENSMUSG00000054715 | Zscan22 | zinc finger and SCAN domain containing 22 |
| RV2 | ENSMUSG00000049600 | Zbtb45 | zinc finger and BTB domain containing 45 |
| RV2 | ENSMUSG00000005566 | Trim28 | tripartite motif-containing 28 |
| RV2 | ENSMUSG00000070814 | Zswim9 | zinc finger SWIM-type containing 9 |
| RV2 | ENSMUSG00000002083 | Bbc3 | BCL2 binding component 3 |
| RV2 | ENSMUSG00000059273 | Zc3h4 | zinc finger CCCH-type containing 4 |
| RV2 | ENSMUSG00000048920 | Fkrp | fukutin related protein |
| RV2 | ENSMUSG00000003099 | Ppp5c | protein phosphatase 5, catalytic subunit |
| RV2 | ENSMUSG00000040841 | Six5 | sine oculis-related homeobox 5 |
| RV2 | ENSMUSG00000052214 | Opa3 | optic atrophy 3 |
| RV2 | ENSMUSG00000051403 | Ppp1r37 | protein phosphatase 1, regulatory subunit 37 |
| RV2 | ENSMUSG00000061028 | Clasrp | CLK4-associating serine/arginine rich protein |
| RV2 | ENSMUSG00000054499 | Dedd2 | death effector domain-containing DNA binding protein 2 |
| RV2 | ENSMUSG00000057177 | Gsk3a | glycogen synthase kinase 3 alpha |
| RV2 | ENSMUSG00000040857 | Erf | Ets2 repressor factor |
| RV2 | ENSMUSG00000045039 | Megf8 | multiple EGF-like-domains 8 |
| RV2 | ENSMUSG00000057229 | Dmac2 | distal membrane arm assembly complex 2 |
| RV2 | ENSMUSG00000002608 | Ccdc97 | coiled-coil domain containing 97 |
| RV2 | ENSMUSG00000004056 | Akt2 | thymoma viral proto-oncogene 2 |
| RV2 | ENSMUSG00000040390 | Map3k10 | mitogen-activated protein kinase kinase kinase 10 |
| RV2 | ENSMUSG00000002409 | Dyrk1b | dual-specificity tyrosine phosphorylation regulated kinase 1b |
| RV2 | ENSMUSG00000046058 | Eid2 | EP300 interacting inhibitor of differentiation 2 |
| RV2 | ENSMUSG00000070705 | Eid2b | EP300 interacting inhibitor of differentiation 2B |
| RV2 | ENSMUSG00000109336 | Samd4b | sterile alpha motif domain containing 4B |
| RV2 | ENSMUSG00000070699 | Sars2 | seryl-aminoacyl-tRNA synthetase 2 |
| RV2 | ENSMUSG00000047473 | Zfp30 | zinc finger protein 30 |
| RV2 | ENSMUSG00000011427 | Zfp790 | zinc finger protein 790 |
| RV2 | ENSMUSG00000036826 | Igflr1 | IGF-like family receptor 1 |
| RV2 | ENSMUSG00000036733 | Rbm42 | RNA binding motif protein 42 |
| RV2 | ENSMUSG00000061099 | Gapdhs | glyceraldehyde-3-phosphate dehydrogenase, spermatogenic |
| RV2 | ENSMUSG00000058239 | Usf2 | upstream transcription factor 2 |
| RV2 | ENSMUSG00000054676 | 1600014C10Rik | RIKEN cDNA 1600014C10 gene |
| RV2 | ENSMUSG00000096146 | Kcnj11 | potassium inwardly rectifying channel, subfamily J, member 11 |
| RV2 | ENSMUSG00000002781 | Tmem143 | transmembrane protein 143 |

| | | | |
|---|---|---|---|
| RV2 | ENSMUSG00000057342 | Sphk2 | sphingosine kinase 2 |
| RV2 | ENSMUSG00000030826 | Bcat2 | branched chain aminotransferase 2, mitochondrial |
| RV2 | ENSMUSG00000003865 | Gys1 | glycogen synthase 1, muscle |
| RV2 | ENSMUSG00000063511 | Snrnp70 | small nuclear ribonucleoprotein 70 (U1) |
| RV2 | ENSMUSG00000038239 | Hrc | histidine rich calcium binding protein |
| RV2 | ENSMUSG00000038406 | Scaf1 | SR-related CTD-associated factor 1 |
| RV2 | ENSMUSG00000002968 | Med25 | mediator complex subunit 25 |
| RV2 | ENSMUSG00000011096 | Akt1s1 | AKT1 substrate 1 |
| RV2 | ENSMUSG00000008140 | Emc10 | ER membrane protein complex subunit 10 |
| RV2 | ENSMUSG00000026254 | Eif4e2 | eukaryotic translation initiation factor 4E member 2 |
| RV2 | ENSMUSG00000026255 | Efhd1 | EF hand domain containing 1 |
| RV2 | ENSMUSG00000034432 | Cops8 | COP9 signalosome subunit 8 |
| RV2 | ENSMUSG00000034220 | Gpc1 | glypican 1 |
| RV2 | ENSMUSG00000047067 | Dusp28 | dual specificity phosphatase 28 |
| RV2 | ENSMUSG00000024228 | Nudt12 | nudix hydrolase 12 |
| RV2 | ENSMUSG00000073375 | Lrrc30 | leucine rich repeat containing 30 |
| RV2 | ENSMUSG00000030942 | Thumpd1 | THUMP domain containing 1 |
| RV2 | ENSMUSG00000035901 | Dennd5a | DENN domain containing 5A |
| RV2 | ENSMUSG00000066279 | Chrna10 | cholinergic receptor, nicotinic, alpha polypeptide 10 |
| RV2 | ENSMUSG00000030996 | Art1 | ADP-ribosyltransferase 1 |
| RV2 | ENSMUSG00000066306 | Numa1 | nuclear mitotic apparatus protein 1 |
| RV2 | ENSMUSG00000032737 | Inppl1 | inositol polyphosphate phosphatase-like 1 |
| RV2 | ENSMUSG00000030701 | Plekhb1 | pleckstrin homology domain containing, family B (evectins) member 1 |
| RV2 | ENSMUSG00000030706 | Mrpl48 | mitochondrial ribosomal protein L48 |
| RV2 | ENSMUSG00000051515 | Fam181b | family with sequence similarity 181, member B |
| RV2 | ENSMUSG00000039428 | Tmem135 | transmembrane protein 135 |
| RV2 | ENSMUSG00000022911 | Arl13b | ADP-ribosylation factor-like 13B |
| RV2 | ENSMUSG00000034206 | Polq | polymerase (DNA directed), theta |
| RV2 | ENSMUSG00000046961 | Gpr156 | G protein-coupled receptor 156 |
| RV2 | ENSMUSG00000022790 | Igsf11 | immunoglobulin superfamily, member 11 |
| RV3 | ENSMUSG00000022309 | Angpt1 | angiopoietin 1 |
| RV3 | ENSMUSG00000037343 | Taf2 | TATA-box binding protein associated factor 2 |
| RV3 | ENSMUSG00000022365 | Derl1 | Der1-like domain family, member 1 |
| RV3 | ENSMUSG00000047921 | Trappc9 | trafficking protein particle complex 9 |
| RV3 | ENSMUSG00000022565 | Plec | plectin |
| RV3 | ENSMUSG00000022561 | Gpaa1 | GPI anchor attachment protein 1 |
| RV3 | ENSMUSG00000033055 | Ankrd54 | ankyrin repeat domain 54 |
| RV3 | ENSMUSG00000022390 | Zc3h7b | zinc finger CCCH type containing 7B |

| | | | |
|---|---|---|---|
| RV3 | ENSMUSG00000034333 | Zbed4 | zinc finger, BED type containing 4 |
| RV3 | ENSMUSG00000015363 | Trabd | TraB domain containing |
| RV3 | ENSMUSG00000052560 | Cpne8 | copine VIII |
| RV3 | ENSMUSG00000022992 | Kansl2 | KAT8 regulatory NSL complex subunit 2 |
| RV3 | ENSMUSG00000023010 | Tmbim6 | transmembrane BAX inhibitor motif containing 6 |
| RV3 | ENSMUSG00000037525 | Bcdin3d | BCDIN3 domain containing |
| RV3 | ENSMUSG00000023021 | Cers5 | ceramide synthase 5 |
| RV3 | ENSMUSG00000023030 | Slc11a2 | solute carrier family 11 (proton-coupled divalent metal ion transporters), member 2 |
| RV3 | ENSMUSG00000029265 | Dr1 | down-regulator of transcription 1 |
| RV3 | ENSMUSG00000029328 | Hnrnpdl | heterogeneous nuclear ribonucleoprotein D-like |
| RV3 | ENSMUSG00000029426 | Scarb2 | scavenger receptor class B, member 2 |
| RV3 | ENSMUSG00000034981 | Parm1 | prostate androgen-regulated mucin-like protein 1 |
| RV3 | ENSMUSG00000029247 | Paics | phosphoribosylaminoimidazole carboxylase, phosphoribosylaminoribosylaminoimidazole, succinocarboxamide synthetase |
| RV3 | ENSMUSG00000037653 | Kctd8 | potassium channel tetramerisation domain containing 8 |
| RV3 | ENSMUSG00000029179 | Zcchc4 | zinc finger, CCHC domain containing 4 |
| RV3 | ENSMUSG00000045106 | Ccdc73 | coiled-coil domain containing 73 |
| RV3 | ENSMUSG00000027134 | Lpcat4 | lysophosphatidylcholine acyltransferase 4 |
| RV3 | ENSMUSG00000027349 | Fam98b | family with sequence similarity 98, member B |
| RV3 | ENSMUSG00000074918 | Inafm2 | InaF motif containing 2 |
| RV3 | ENSMUSG00000027324 | Rpusd2 | RNA pseudouridylate synthase domain containing 2 |
| RV3 | ENSMUSG00000050619 | Zscan29 | zinc finger SCAN domains 29 |
| RV3 | ENSMUSG00000000308 | Ckmt1 | creatine kinase, mitochondrial 1, ubiquitous |
| RV3 | ENSMUSG00000027224 | Duoxa1 | dual oxidase maturation factor 1 |
| RV3 | ENSMUSG00000033268 | Duox1 | dual oxidase 1 |
| RV3 | ENSMUSG00000037703 | Lzts3 | leucine zipper, putative tumor suppressor family member 3 |
| RV3 | ENSMUSG00000037523 | Mavs | mitochondrial antiviral signaling protein |
| RV3 | ENSMUSG00000015932 | Dstn | destrin |
| RV3 | ENSMUSG00000033059 | Pygb | brain glycogen phosphorylase |
| RV3 | ENSMUSG00000027459 | Fam110a | family with sequence similarity 110, member A |
| RV3 | ENSMUSG00000038375 | Trp53inp2 | transformation related protein 53 inducible nuclear protein 2 |
| RV3 | ENSMUSG00000027610 | Gss | glutathione synthetase |
| RV3 | ENSMUSG00000027628 | Aar2 | AAR2 splicing factor homolog |
| RV3 | ENSMUSG00000061689 | Dlgap4 | DLG associated protein 4 |
| RV3 | ENSMUSG00000037820 | Tgm2 | transglutaminase 2, C polypeptide |
| RV3 | ENSMUSG00000050373 | Snx21 | sorting nexin family member 21 |
| RV3 | ENSMUSG00000017670 | Elmo2 | engulfment and cell motility 2 |
| RV3 | ENSMUSG00000039671 | Zmynd8 | zinc finger, MYND-type containing 8 |
| RV3 | ENSMUSG00000078923 | Ube2v1 | ubiquitin-conjugating enzyme E2 variant 1 |

| | | | |
|---|---|---|---|
| RV3 | ENSMUSG00000090213 | Peds1 | plasmanylethanolamine desaturase 1 |
| RV3 | ENSMUSG00000027551 | Zfp64 | zinc finger protein 64 |
| RV3 | ENSMUSG00000027496 | Aurka | aurora kinase A |
| RV3 | ENSMUSG00000054455 | Vapb | vesicle-associated membrane protein, associated protein B and C |
| RV3 | ENSMUSG00000016253 | Nelfcd | negative elongation factor complex member C/D, Th1l |
| RV3 | ENSMUSG00000027080 | Med19 | mediator complex subunit 19 |
| RV3 | ENSMUSG00000013465 | Nelfb | negative elongation factor complex member B |
| RV3 | ENSMUSG00000036752 | Tubb4b | tubulin, beta 4B class IVB |
| RV3 | ENSMUSG00000026930 | Gpsm1 | G-protein signalling modulator 1 (AGS3-like, C. elegans) |
| RV3 | ENSMUSG00000039844 | Rapgef1 | Rap guanine nucleotide exchange factor (GEF) 1 |
| RV3 | ENSMUSG00000059316 | Slc27a4 | solute carrier family 27 (fatty acid transporter), member 4 |
| RV3 | ENSMUSG00000039648 | Kyat1 | kynurenine aminotransferase 1 |
| RV3 | ENSMUSG00000007476 | Lrrc8a | leucine rich repeat containing 8A VRAC subunit A |
| RV3 | ENSMUSG00000026856 | Dolpp1 | dolichyl pyrophosphate phosphatase 1 |
| RV3 | ENSMUSG00000039515 | Ptpa | protein phosphatase 2 protein activator |
| RV3 | ENSMUSG00000039254 | Pomt1 | protein-O-mannosyltransferase 1 |
| RV3 | ENSMUSG00000009566 | Fpgs | folylpolyglutamyl synthetase |
| RV3 | ENSMUSG00000039021 | Ttc16 | tetratricopeptide repeat domain 16 |
| RV3 | ENSMUSG00000028514 | Usp24 | ubiquitin specific peptidase 24 |
| RV3 | ENSMUSG00000028517 | Plpp3 | phospholipid phosphatase 3 |
| RV3 | ENSMUSG00000062937 | Mtap | methylthioadenosine phosphorylase |
| RV3 | ENSMUSG00000070923 | Klhl9 | kelch-like 9 |
| RV3 | ENSMUSG00000028540 | Dph2 | DPH2 homolog |
| RV3 | ENSMUSG00000028709 | Mob3c | MOB kinase activator 3C |
| RV3 | ENSMUSG00000066798 | Zbtb6 | zinc finger and BTB domain containing 6 |
| RV3 | ENSMUSG00000038007 | Acer2 | alkaline ceramidase 2 |
| RV3 | ENSMUSG00000038506 | Dcun1d2 | defective in cullin neddylation 1 domain containing 2 |
| RV3 | ENSMUSG00000085795 | Zfp703 | zinc finger protein 703 |
| RV3 | ENSMUSG00000031626 | Sorbs2 | sorbin and SH3 domain containing 2 |
| RV3 | ENSMUSG00000038193 | Hand2 | heart and neural crest derivatives expressed 2 |
| RV3 | ENSMUSG00000002396 | Ocel1 | occludin/ELL domain containing 1 |
| RV3 | ENSMUSG00000043243 | Niban3 | niban apoptosis regulator 3 |
| RV3 | ENSMUSG00000019261 | Map1s | microtubule-associated protein 1S |
| RV3 | ENSMUSG00000031840 | Rab3a | RAB3A, member RAS oncogene family |
| RV3 | ENSMUSG00000070002 | Ell | elongation factor RNA polymerase II |
| RV3 | ENSMUSG00000058301 | Upf1 | UPF1 RNA helicase and ATPase |
| RV3 | ENSMUSG00000036054 | Sugp2 | SURP and G patch domain containing 2 |
| RV3 | ENSMUSG00000022789 | Dnm1l | dynamin 1-like |
| RV3 | ENSMUSG00000022841 | Ap2m1 | adaptor-related protein complex 2, mu 1 subunit |

| | | | |
|---|---|---|---|
| RV3 | ENSMUSG00000051065 | Mb21d2 | Mab-21 domain containing 2 |
| RV3 | ENSMUSG00000005615 | Pcyt1a | phosphate cytidylyltransferase 1, choline, alpha isoform |
| RV3 | ENSMUSG00000022797 | Tfrc | transferrin receptor |
| RV3 | ENSMUSG00000011877 | Git1 | GIT ArfGAP 1 |
| RV3 | ENSMUSG00000020841 | Cpd | carboxypeptidase D |
| RV3 | ENSMUSG00000010392 | Gosr1 | golgi SNAP receptor complex member 1 |
| RV3 | ENSMUSG00000020849 | Ywhae | tyrosine 3-monooxygenase/tryptophan 5-monooxygenase activation protein, epsilon polypeptide |
| RV3 | ENSMUSG00000017774 | Myo1c | myosin IC |
| RV3 | ENSMUSG00000055670 | Zzef1 | zinc finger, ZZ-type with EF hand domain 1 |
| RV3 | ENSMUSG00000020794 | Ube2g1 | ubiquitin-conjugating enzyme E2G 1 |
| RV3 | ENSMUSG00000014609 | Chrne | cholinergic receptor, nicotinic, epsilon polypeptide |
| RV3 | ENSMUSG00000087279 | 4930544D05Rik | RIKEN cDNA 4930544D05 gene |
| RV3 | ENSMUSG00000078812 | Eif5a | eukaryotic translation initiation factor 5A |
| RV3 | ENSMUSG00000018750 | Zbtb4 | zinc finger and BTB domain containing 4 |
| RV3 | ENSMUSG00000044795 | Cyb5d1 | cytochrome b5 domain containing 1 |
| RV3 | ENSMUSG00000042331 | Specc1 | sperm antigen with calponin homology and coiled-coil domains 1 |
| RV3 | ENSMUSG00000042506 | Usp22 | ubiquitin specific peptidase 22 |
| RV3 | ENSMUSG00000042650 | Alkbh5 | alkB homolog 5, RNA demethylase |
| RV3 | ENSMUSG00000013646 | Sh3bp5l | SH3 binding domain protein 5 like |
| RV3 | ENSMUSG00000018583.11 | G3bp1 | G3BP stress granule assembly factor 1 |
| RV3 | ENSMUSG00000036275 | 9530068E07Rik | RIKEN cDNA 9530068E07 gene |
| RV3 | ENSMUSG00000020362 | Cnot6 | CCR4-NOT transcription complex, subunit 6 |
| RV3 | ENSMUSG00000028645 | Slc2a1 | solute carrier family 2 (facilitated glucose transporter), member 1 |
| RV3 | ENSMUSG00000032998 | Foxj3 | forkhead box J3 |
| RV3 | ENSMUSG00000032870 | Smap2 | small ArfGAP 2 |
| RV3 | ENSMUSG00000028833 | Ncdn | neurochondrin |
| RV3 | ENSMUSG00000028811 | Yars1 | tyrosyl-tRNA synthetase 1 |
| RV3 | ENSMUSG00000053841 | Txlna | taxilin alpha |
| RV3 | ENSMUSG00000028790 | Khdrbs1 | KH domain containing, RNA binding, signal transduction associated 1 |
| RV3 | ENSMUSG00000028580 | Pum1 | pumilio RNA-binding family member 1 |
| RV3 | ENSMUSG00000028906 | Epb41 | erythrocyte membrane protein band 4.1 |
| RV3 | ENSMUSG00000012117 | Dhdds | dehydrodolichyl diphosphate synthase |
| RV3 | ENSMUSG00000037242 | Clic4 | chloride intracellular channel 4 |
| RV3 | ENSMUSG00000057530 | Ece1 | endothelin converting enzyme 1 |
| RV3 | ENSMUSG00000041120 | Nbl1 | NBL1, DAN family BMP antagonist |
| RV3 | ENSMUSG00000028745 | Capzb | capping actin protein of muscle Z-line subunit beta |
| RV3 | ENSMUSG00000078517 | Emc1 | ER membrane protein complex subunit 1 |
| RV3 | ENSMUSG00000040761 | Spen | spen family transcription repressor |

| | | | |
|---|---|---|---|
| RV3 | ENSMUSG00000006219 | Fblim1 | filamin binding LIM protein 1 |
| RV3 | ENSMUSG00000047719 | Ubiad1 | UbiA prenyltransferase domain containing 1 |
| RV3 | ENSMUSG00000041459 | Tardbp | TAR DNA binding protein |
| RV3 | ENSMUSG00000028992 | Nmnat1 | nicotinamide nucleotide adenylyltransferase 1 |
| RV3 | ENSMUSG00000029027 | Dffb | DNA fragmentation factor, beta subunit |
| RV3 | ENSMUSG00000029036 | Atad3a | ATPase family, AAA domain containing 3A |
| RV3 | ENSMUSG00000050796 | B3galt6 | UDP-Gal:betaGal beta 1,3-galactosyltransferase, polypeptide 6 |
| RV3 | ENSMUSG00000026718 | Stam | signal transducing adaptor molecule (SH3 domain and ITAM motif) 1 |
| RV3 | ENSMUSG00000033960 | Jcad | junctional cadherin 5 associated |
| RV3 | ENSMUSG00000006362 | Cbfa2t3 | CBFA2/RUNX1 translocation partner 3 |
| RV3 | ENSMUSG00000006585 | Cdt1 | chromatin licensing and DNA replication factor 1 |
| RV3 | ENSMUSG00000034390 | Cmip | c-Maf inducing protein |
| RV3 | ENSMUSG00000000561 | Wdr77 | WD repeat domain 77 |
| RV3 | ENSMUSG00000036333 | Kidins220 | kinase D-interacting substrate 220 |
| RV3 | ENSMUSG00000052593 | Adam17 | a disintegrin and metallopeptidase domain 17 |
| RV3 | ENSMUSG00000020657 | Dnajc27 | DnaJ heat shock protein family (Hsp40) member C27 |
| RV3 | ENSMUSG00000020671 | Rab10 | RAB10, member RAS oncogene family |
| RV3 | ENSMUSG00000062761 | Zfp512 | zinc finger protein 512 |
| RV3 | ENSMUSG00000014956 | Ppp1cb | protein phosphatase 1 catalytic subunit beta |
| RV3 | ENSMUSG00000024063 | Lbh | limb-bud and heart |
| RV3 | ENSMUSG00000024097 | Srsf7 | serine and arginine-rich splicing factor 7 |
| RV3 | ENSMUSG00000024247 | Pkdcc | protein kinase domain containing, cytoplasmic |
| RV3 | ENSMUSG00000037104 | Socs5 | suppressor of cytokine signaling 5 |
| RV3 | ENSMUSG00000024045 | Akap8 | A kinase anchor protein 8 |
| RV3 | ENSMUSG00000020325 | Fstl3 | follistatin-like 3 |
| RV3 | ENSMUSG00000035621 | Midn | midnolin |
| RV3 | ENSMUSG00000048696 | Mex3d | mex3 RNA binding family member D |
| RV3 | ENSMUSG00000001229 | Dpp9 | dipeptidylpeptidase 9 |
| RV3 | ENSMUSG00000025231 | Sufu | SUFU negative regulator of hedgehog signaling |
| RV3 | ENSMUSG00000011752 | Pgam1 | phosphoglycerate mutase 1 |
| RV3 | ENSMUSG00000025007 | Aldh18a1 | aldehyde dehydrogenase 18 family, member A1 |
| RV3 | ENSMUSG00000024999 | Noc3l | NOC3 like DNA replication regulator |
| RV3 | ENSMUSG00000024795 | Kif20b | kinesin family member 20B |
| RV3 | ENSMUSG00000013662 | Atad1 | ATPase family, AAA domain containing 1 |
| RV3 | ENSMUSG00000024810 | Il33 | interleukin 33 |
| RV3 | ENSMUSG00000074925 | Ptar1 | protein prenyltransferase alpha subunit repeat containing 1 |
| RV3 | ENSMUSG00000033207 | Mamdc2 | MAM domain containing 2 |
| RV3 | ENSMUSG00000046139 | Patl1 | protein associated with topoisomerase II homolog 1 (yeast) |

| | | | |
|---|---|---|---|
| RV3 | ENSMUSG00000024743 | Syt7 | synaptotagmin VII |
| RV3 | ENSMUSG00000035735 | Dagla | diacylglycerol lipase, alpha |
| RV3 | ENSMUSG00000024970 | Spindoc | spindlin interactor and repressor of chromatin binding |
| RV3 | ENSMUSG00000024952 | Rps6ka4 | ribosomal protein S6 kinase, polypeptide 4 |
| RV3 | ENSMUSG00000024777 | Ppp2r5b | protein phosphatase 2, regulatory subunit B', beta |
| RV3 | ENSMUSG00000024939 | Fam89b | family with sequence similarity 89, member B |
| RV3 | ENSMUSG00000024855 | Pacs1 | phosphofurin acidic cluster sorting protein 1 |
| RV3 | ENSMUSG00000056481 | Cd248 | CD248 antigen, endosialin |
| RV3 | ENSMUSG00000031078 | Cttn | cortactin |
| RV3 | ENSMUSG00000059119 | Nap1l4 | nucleosome assembly protein 1-like 4 |
| RV3 | ENSMUSG00000030862 | Cpxm2 | carboxypeptidase X, M14 family member 2 |
| RV3 | ENSMUSG00000006205 | Htra1 | HtrA serine peptidase 1 |
| RV3 | ENSMUSG00000030850 | Ate1 | arginyltransferase 1 |
| RV3 | ENSMUSG00000042423 | Fbrs | fibrosin |
| RV3 | ENSMUSG00000047371 | Zfp768 | zinc finger protein 768 |
| RV3 | ENSMUSG00000030682 | Cdipt | CDP-diacylglycerol--inositol 3-phosphatidyltransferase |
| RV3 | ENSMUSG00000030689 | Ino80e | INO80 complex subunit E |
| RV3 | ENSMUSG00000030722 | Nfatc2ip | nuclear factor of activated T cells, cytoplasmic, calcineurin dependent 2 interacting protein |
| RV3 | ENSMUSG00000009741 | Ubp1 | upstream binding protein 1 |
| RV3 | ENSMUSG00000032504 | Pdcd6ip | programmed cell death 6 interacting protein |
| RV3 | ENSMUSG00000039952 | Dag1 | dystroglycan 1 |
| RV3 | ENSMUSG00000032583 | Mon1a | MON1 homolog A, secretory traffciking associated |
| RV3 | ENSMUSG00000010064 | Slc38a3 | solute carrier family 38, member 3 |
| RV3 | ENSMUSG00000020257 | Wdr82 | WD repeat domain containing 82 |
| RV3 | ENSMUSG00000032839 | Trpc1 | transient receptor potential cation channel, subfamily C, member 1 |
| RV3 | ENSMUSG00000045414 | Dipk2a | divergent protein kinase domain 2A |
| RV3 | ENSMUSG00000032420 | Nt5e | 5' nucleotidase, ecto |
| RV3 | ENSMUSG00000035941 | Ibtk | inhibitor of Bruton agammaglobulinemia tyrosine kinase |
| RV3 | ENSMUSG00000032328 | Tmem30a | transmembrane protein 30A |
| RV3 | ENSMUSG00000032216 | Nedd4 | neural precursor cell expressed, developmentally down-regulated 4 |
| RV3 | ENSMUSG00000040524 | Zfp609 | zinc finger protein 609 |
| RV3 | ENSMUSG00000032816 | Igdcc4 | immunoglobulin superfamily, DCC subclass, member 4 |
| RV3 | ENSMUSG00000022120 | Obi1 | ORC ubiquitin ligase 1 |
| RV3 | ENSMUSG00000022003 | Slc25a30 | solute carrier family 25, member 30 |
| RV3 | ENSMUSG00000022100 | Xpo7 | exportin 7 |
| RV3 | ENSMUSG00000022094 | Slc39a14 | solute carrier family 39 (zinc transporter), member 14 |
| RV3 | ENSMUSG00000022048 | Dpysl2 | dihydropyrimidinase-like 2 |
| RV3 | ENSMUSG00000021978 | Extl3 | exostosin-like glycosyltransferase 3 |

| | | | |
|---|---|---|---|
| RV3 | ENSMUSG00000021983 | Atp8a2 | ATPase, aminophospholipid transporter-like, class I, type 8A, member 2 |
| RV3 | ENSMUSG00000021969 | Zdhhc20 | zinc finger, DHHC domain containing 20 |
| RV3 | ENSMUSG00000048582 | Gja3 | gap junction protein, alpha 3 |
| RV3 | ENSMUSG00000060373 | Hnrnpc | heterogeneous nuclear ribonucleoprotein C |
| RV3 | ENSMUSG00000021840 | Mapk1ip1l | mitogen-activated protein kinase 1 interacting protein 1-like |
| RV3 | ENSMUSG00000015759 | Cnih1 | cornichon family AMPA receptor auxiliary protein 1 |
| RV3 | ENSMUSG00000021823 | Vcl | vinculin |
| RV3 | ENSMUSG00000039357 | Fut11 | fucosyltransferase 11 |
| RV3 | ENSMUSG00000039740 | Alg2 | ALG2 alpha-1,3/1,6-mannosyltransferase |
| RV3 | ENSMUSG00000028378 | Ptgr1 | prostaglandin reductase 1 |
| RV3 | ENSMUSG00000027499 | Pkia | protein kinase inhibitor, alpha |
| RV3 | ENSMUSG00000025776 | Crispld1 | cysteine-rich secretory protein LCCL domain containing 1 |
| RV3 | ENSMUSG00000025921 | Rdh10 | retinol dehydrogenase 10 (all-trans) |
| RV3 | ENSMUSG00000028261 | Ndufaf4 | NADH:ubiquinone oxidoreductase complex assembly factor 4 |
| RV3 | ENSMUSG00000028433 | Ubap2 | ubiquitin-associated protein 2 |
| RV3 | ENSMUSG00000029772 | Ahcyl2 | S-adenosylhomocysteine hydrolase-like 2 |
| RV3 | ENSMUSG00000021264 | Yy1 | YY1 transcription factor |
| RV3 | ENSMUSG00000034126 | Pomt2 | protein-O-mannosyltransferase 2 |
| RV3 | ENSMUSG00000034290 | Nek9 | NIMA (never in mitosis gene a)-related expressed kinase 9 |
| RV3 | ENSMUSG00000042350 | Arel1 | apoptosis resistant E3 ubiquitin protein ligase 1 |
| RV3 | ENSMUSG00000021219 | Rgs6 | regulator of G-protein signaling 6 |
| RV3 | ENSMUSG00000021133 | Susd6 | sushi domain containing 6 |
| RV3 | ENSMUSG00000021127 | Zfp36l1 | zinc finger protein 36, C3H type-like 1 |
| RV3 | ENSMUSG00000047454 | Gphn | gephyrin |
| RV3 | ENSMUSG00000045690 | Wdr89 | WD repeat domain 89 |
| RV3 | ENSMUSG00000034574 | Daam1 | dishevelled associated activator of morphogenesis 1 |
| RV3 | ENSMUSG00000052397 | Ezr | ezrin |
| RV3 | ENSMUSG00000034377 | Tulp4 | TUB like protein 4 |
| RV3 | ENSMUSG00000055493 | Epm2a | epilepsy, progressive myoclonic epilepsy, type 2 gene alpha |
| RV3 | ENSMUSG00000019877 | Serinc1 | serine incorporator 1 |
| RV3 | ENSMUSG00000050953 | Gja1 | gap junction protein, alpha 1 |
| RV3 | ENSMUSG00000003746 | Man1a | mannosidase 1, alpha |
| RV3 | ENSMUSG00000019818 | Cd164 | CD164 antigen |
| RV3 | ENSMUSG00000071317 | Bves | blood vessel epicardial substance |
| RV3 | ENSMUSG00000056870 | Gulp1 | GULP, engulfment adaptor PTB domain containing 1 |
| RV3 | ENSMUSG00000117809 | Asdurf | Asnsd1 upstream reading frame |
| RV3 | ENSMUSG00000033124 | Atg9a | autophagy related 9A |
| RV3 | ENSMUSG00000026202 | Tuba4a | tubulin, alpha 4A |

| | | | |
|---|---|---|---|
| RV3 | ENSMUSG00000026249 | Serpine2 | serine (or cysteine) peptidase inhibitor, clade E, member 2 |
| RV3 | ENSMUSG00000034109 | Golim4 | golgi integral membrane protein 4 |
| RV3 | ENSMUSG00000033910 | Gucy1a1 | guanylate cyclase 1, soluble, alpha 1 |
| RV3 | ENSMUSG00000028086 | Fbxw7 | F-box and WD-40 domain protein 7 |
| RV3 | ENSMUSG00000001416 | Cct3 | chaperonin containing TCP1 subunit 3 |
| RV3 | ENSMUSG00000001415 | Smg5 | SMG5 nonsense mediated mRNA decay factor |
| RV3 | ENSMUSG00000028126 | Pip5k1a | phosphatidylinositol-4-phosphate 5-kinase, type 1 alpha |
| RV3 | ENSMUSG00000046519 | Golph3l | golgi phosphoprotein 3-like |
| RV3 | ENSMUSG00000027879 | Sec22b | SEC22 homolog B, vesicle trafficking protein |
| RV3 | ENSMUSG00000042035 | Igsf3 | immunoglobulin superfamily, member 3 |
| RV3 | ENSMUSG00000033161 | Atp1a1 | ATPase, Na+/K+ transporting, alpha 1 polypeptide |
| RV3 | ENSMUSG00000039275 | Foxk2 | forkhead box K2 |
| RV3 | ENSMUSG00000039307 | Hexd | hexosaminidase D |
| RV3 | ENSMUSG00000025132 | Arhgdia | Rho GDP dissociation inhibitor alpha |
| RV3 | ENSMUSG00000039850 | Endov | endonuclease V |
| RV3 | ENSMUSG00000059248 | Septin9 | septin 9 |
| RV3 | ENSMUSG00000034120 | Srsf2 | serine and arginine-rich splicing factor 2 |
| RV3 | ENSMUSG00000020792 | Exoc7 | exocyst complex component 7 |
| RV3 | ENSMUSG00000020740 | Gga3 | golgi associated, gamma adaptin ear containing, ARF binding protein 3 |
| RV3 | ENSMUSG00000034586 | Hid1 | HID1 domain containing |
| RV3 | ENSMUSG00000034714 | Ttyh2 | tweety family member 2 |
| RV3 | ENSMUSG00000000567 | Sox9 | SRY (sex determining region Y)-box 9 |
| RV3 | ENSMUSG00000040592 | Cd79b | CD79B antigen |
| RV3 | ENSMUSG00000049354 | Dcaf7 | DDB1 and CUL4 associated factor 7 |
| RV3 | ENSMUSG00000020941 | Map3k14 | mitogen-activated protein kinase kinase kinase 14 |
| RV3 | ENSMUSG00000050288 | Fzd2 | frizzled class receptor 2 |
| RV3 | ENSMUSG00000035198 | Tubg1 | tubulin, gamma 1 |
| RV3 | ENSMUSG00000004044 | Cavin1 | caveolae associated 1 |
| RV3 | ENSMUSG00000004040 | Stat3 | signal transducer and activator of transcription 3 |
| RV3 | ENSMUSG00000017837 | Nkiras2 | NFKB inhibitor interacting Ras-like protein 2 |
| RV3 | ENSMUSG00000078676 | Casc3 | exon junction complex subunit |
| RV3 | ENSMUSG00000017210 | Med24 | mediator complex subunit 24 |
| RV3 | ENSMUSG00000003119 | Cdk12 | cyclin dependent kinase 12 |
| RV3 | ENSMUSG00000018666.1 | Cbx1 | chromobox 1 |
| RV3 | ENSMUSG00000020864 | Ankrd40 | ankyrin repeat domain 40 |
| RV3 | ENSMUSG00000020544 | Cox11 | cytochrome c oxidase assembly protein 11, copper chaperone |
| RV3 | ENSMUSG00000069763 | Tmem100 | transmembrane protein 100 |
| RV3 | ENSMUSG00000020516 | Rps6kb1 | ribosomal protein S6 kinase, polypeptide 1 |

| | | | |
|---|---|---|---|
| RV3 | ENSMUSG00000018651 | Tada2a | transcriptional adaptor 2A |
| RV3 | ENSMUSG00000037351 | Actr1b | ARP1 actin-related protein 1B, centractin beta |
| RV3 | ENSMUSG00000037470 | Uggt1 | UDP-glucose glycoprotein glucosyltransferase 1 |
| RV3 | ENSMUSG00000073725 | Lmbrd1 | LMBR1 domain containing 1 |
| RV3 | ENSMUSG00000034509 | Mad2l1bp | MAD2L1 binding protein |
| RV3 | ENSMUSG00000059409 | Ppp2r5d | protein phosphatase 2, regulatory subunit B', delta |
| RV3 | ENSMUSG00000036568 | Bicral | BRD4 interacting chromatin remodeling complex associated protein like |
| RV3 | ENSMUSG00000063253 | Scoc | short coiled-coil protein |
| RV3 | ENSMUSG00000058355 | Abce1 | ATP-binding cassette, sub-family E member 1 |
| RV3 | ENSMUSG00000005483 | Dnajb1 | DnaJ heat shock protein family (Hsp40) member B1 |
| RV3 | ENSMUSG00000031696 | Vps35 | VPS35 retromer complex component |
| RV3 | ENSMUSG00000031751 | Amfr | autocrine motility factor receptor |
| RV3 | ENSMUSG00000050079 | Rspry1 | ring finger and SPRY domain containing 1 |
| RV3 | ENSMUSG00000036534 | Slc38a7 | solute carrier family 38, member 7 |
| RV3 | ENSMUSG00000005698 | Ctcf | CCCTC-binding factor |
| RV3 | ENSMUSG00000048310 | Pskh1 | protein serine kinase H1 |
| RV3 | ENSMUSG00000045538 | Ddx28 | DEAD box helicase 28 |
| RV3 | ENSMUSG00000060098 | Prmt7 | protein arginine N-methyltransferase 7 |
| RV3 | ENSMUSG00000031749 | St3gal2 | ST3 beta-galactoside alpha-2,3-sialyltransferase 2 |
| RV3 | ENSMUSG00000003316 | Glg1 | golgi apparatus protein 1 |
| RV3 | ENSMUSG00000031959 | Wdr59 | WD repeat domain 59 |
| RV3 | ENSMUSG00000024563 | Smad2 | SMAD family member 2 |
| RV3 | ENSMUSG00000024558 | Mapk4 | mitogen-activated protein kinase 4 |
| RV3 | ENSMUSG00000038121 | Fam210a | family with sequence similarity 210, member A |
| RV3 | ENSMUSG00000032688 | Malt1 | MALT1 paracaspase |
| RV3 | ENSMUSG00000024576 | Csnk1a1 | casein kinase 1, alpha 1 |
| RV3 | ENSMUSG00000024535 | Snx24 | sorting nexing 24 |
| RV3 | ENSMUSG00000024487 | Yipf5 | Yip1 domain family, member 5 |
| RV3 | ENSMUSG00000033272 | Slc35a4 | solute carrier family 35, member A4 |
| RV3 | ENSMUSG00000024486 | Hbegf | heparin-binding EGF-like growth factor |
| RV3 | ENSMUSG00000091896 | Ube2d2a | ubiquitin-conjugating enzyme E2D 2A |
| RV3 | ENSMUSG00000092124 | B930094E09Rik | RIKEN cDNA B930094E09 gene |
| RV3 | ENSMUSG00000024277 | Mapre2 | microtubule-associated protein, RP/EB family, member 2 |
| RV3 | ENSMUSG00000029860 | Zyx | zyxin |
| RV3 | ENSMUSG00000058486 | Wdr91 | WD repeat domain 91 |
| RV3 | ENSMUSG00000030029 | Lrig1 | leucine-rich repeats and immunoglobulin-like domains 1 |
| RV3 | ENSMUSG00000056234 | Ncoa4 | nuclear receptor coactivator 4 |
| RV3 | ENSMUSG00000050666 | Vstm4 | V-set and transmembrane domain containing 4 |

| | | | |
|---|---|---|---|
| RV3 | ENSMUSG00000021794 | Glud1 | glutamate dehydrogenase 1 |
| RV3 | ENSMUSG00000021798 | Ldb3 | LIM domain binding 3 |
| RV3 | ENSMUSG00000021994 | Wnt5a | wingless-type MMTV integration site family, member 5A |
| RV3 | ENSMUSG00000043702 | Pde12 | phosphodiesterase 12 |
| RV3 | ENSMUSG00000030505 | Prmt3 | protein arginine N-methyltransferase 3 |
| RV3 | ENSMUSG00000016128 | Stard13 | StAR related lipid transfer domain containing 13 |
| RV3 | ENSMUSG00000019470 | Xab2 | XPA binding protein 2 |
| RV3 | ENSMUSG00000038797 | Zscan2 | zinc finger and SCAN domain containing 2 |
| RV3 | ENSMUSG00000039043 | Arpin | actin-related protein 2/3 complex inhibitor |
| RV3 | ENSMUSG00000055652 | Klhl25 | kelch-like 25 |
| RV3 | ENSMUSG00000025790 | Slco3a1 | solute carrier organic anion transporter family, member 3a1 |
| RV3 | ENSMUSG00000027803 | Wwtr1 | WW domain containing transcription regulator 1 |
| RV3 | ENSMUSG00000036615 | Rfxap | regulatory factor X-associated protein |
| RV3 | ENSMUSG00000023087 | Noct | nocturnin |
| RV3 | ENSMUSG00000025757 | Hspa4l | heat shock protein 4 like |
| RV3 | ENSMUSG00000037400 | Atp11b | ATPase, class VI, type 11B |
| RV3 | ENSMUSG00000027669 | Gnb4 | guanine nucleotide binding protein (G protein), beta 4 |
| RV3 | ENSMUSG00000022898 | Vps26c | VPS26 endosomal protein sorting factor C |
| RV3 | ENSMUSG00000022892 | App | amyloid beta precursor protein |
| RV3 | ENSMUSG00000019916 | P4ha1 | procollagen-proline, 2-oxoglutarate 4-dioxygenase (proline 4-hydroxylase), alpha 1 polypeptide |
| RV3 | ENSMUSG00000020099 | Unc5b | unc-5 netrin receptor B |
| RV3 | ENSMUSG00000059901 | Adamts14 | ADAM metallopeptidase with thrombospondin type 1 motif 14 |
| RV3 | ENSMUSG00000033416 | Gucd1 | guanylyl cyclase domain containing 1 |
| RV3 | ENSMUSG00000001436 | Slc19a1 | solute carrier family 19 (folate transporter), member 1 |
| RV3 | ENSMUSG00000023067 | Cdkn1a | cyclin dependent kinase inhibitor 1A |
| RV3 | ENSMUSG00000024006 | Stk38 | serine/threonine kinase 38 |
| RV3 | ENSMUSG00000057789 | Bak1 | BCL2-antagonist/killer 1 |
| RV3 | ENSMUSG00000067629 | Syngap1 | synaptic Ras GTPase activating protein 1 homolog (rat) |
| RV3 | ENSMUSG00000007050 | Lsm2 | LSM2 homolog, U6 small nuclear RNA and mRNA degradation associated |
| RV3 | ENSMUSG00000058672 | Tubb2a | tubulin, beta 2A class IIA |
| RV3 | ENSMUSG00000021360 | Gcnt2 | glucosaminyl (N-acetyl) transferase 2 (I blood group) |
| RV3 | ENSMUSG00000021375 | Kif13a | kinesin family member 13A |
| RV3 | ENSMUSG00000038014 | Fam120a | family with sequence similarity 120, member A |
| RV3 | ENSMUSG00000037933 | Bicd2 | BICD cargo adaptor 2 |
| RV3 | ENSMUSG00000021493 | Pdlim7 | PDZ and LIM domain 7 |
| RV3 | ENSMUSG00000030301 | Ccdc91 | coiled-coil domain containing 91 |
| RV3 | ENSMUSG00000048776 | Pthlh | parathyroid hormone-like peptide |
| RV3 | ENSMUSG00000030268 | Bcat1 | branched chain aminotransferase 1, cytosolic |

| | | | |
|---|---|---|---|
| RV3 | ENSMUSG00000041741 | Pde3a | phosphodiesterase 3A, cGMP inhibited |
| RV3 | ENSMUSG00000030223 | Ptpro | protein tyrosine phosphatase receptor type O |
| RV3 | ENSMUSG00000030208 | Emp1 | epithelial membrane protein 1 |
| RV3 | ENSMUSG00000001521 | Tulp3 | TUB like protein 3 |
| RV3 | ENSMUSG00000030352 | Tspan9 | tetraspanin 9 |
| RV3 | ENSMUSG00000030339 | Ltbr | lymphotoxin B receptor |
| RV3 | ENSMUSG00000038252 | Ncapd2 | non-SMC condensin I complex, subunit D2 |
| RV3 | ENSMUSG00000030330 | Ing4 | inhibitor of growth family, member 4 |
| RV3 | ENSMUSG00000038346 | Zfp384 | zinc finger protein 384 |
| RV3 | ENSMUSG00000030127 | Cops7a | COP9 signalosome subunit 7A |
| RV3 | ENSMUSG00000030327 | Necap1 | NECAP endocytosis associated 1 |
| RV3 | ENSMUSG00000003154 | Foxj2 | forkhead box J2 |
| RV3 | ENSMUSG00000030177 | Ccdc77 | coiled-coil domain containing 77 |
| RV3 | ENSMUSG00000033933 | Vhl | von Hippel-Lindau tumor suppressor |
| RV3 | ENSMUSG00000030105 | Arl8b | ADP-ribosylation factor-like 8B |
| RV3 | ENSMUSG00000052144 | Ppp4r2 | protein phosphatase 4, regulatory subunit 2 |
| RV3 | ENSMUSG00000040550 | Otud6b | OTU domain containing 6B |
| RV3 | ENSMUSG00000034575 | Tent4a | terminal nucleotidyltransferase 4A |
| RV3 | ENSMUSG00000021594 | Srd5a1 | steroid 5 alpha-reductase 1 |
| RV3 | ENSMUSG00000041747 | Utp15 | UTP15 small subunit processome component |
| RV3 | ENSMUSG00000021756 | Il6st | interleukin 6 signal transducer |
| RV3 | ENSMUSG00000022142 | Nup155 | nucleoporin 155 |
| RV3 | ENSMUSG00000022206 | Npr3 | natriuretic peptide receptor 3 |
| RV3 | ENSMUSG00000022195 | 6030458C11Rik | RIKEN cDNA 6030458C11 gene |
| RV3 | ENSMUSG00000032243 | Itga11 | integrin alpha 11 |
| RV3 | ENSMUSG00000041729 | Coro2b | coronin, actin binding protein, 2B |
| RV3 | ENSMUSG00000032290 | Ptpn9 | protein tyrosine phosphatase, non-receptor type 9 |
| RV3 | ENSMUSG00000032076 | Cadm1 | cell adhesion molecule 1 |
| RV3 | ENSMUSG00000035382 | Pcsk7 | proprotein convertase subtilisin/kexin type 7 |
| RV3 | ENSMUSG00000042138 | Msantd2 | Myb/SANT-like DNA-binding domain containing 2 |
| RV3 | ENSMUSG00000038119 | Cdon | cell adhesion molecule-related/down-regulated by oncogenes |
| RV3 | ENSMUSG00000031988 | Vps26b | VPS26 retromer complex component B |
| RV3 | ENSMUSG00000031969 | Acad8 | acyl-Coenzyme A dehydrogenase family, member 8 |
| RV3 | ENSMUSG00000031965 | Tbx20 | T-box 20 |
| RV3 | ENSMUSG00000001349 | Cnn1 | calponin 1 |
| RV3 | ENSMUSG00000033335 | Dnm2 | dynamin 2 |
| RV3 | ENSMUSG00000004098 | Col5a3 | collagen, type V, alpha 3 |
| RV3 | ENSMUSG00000031931 | Ankrd49 | ankyrin repeat domain 49 |
| RV3 | ENSMUSG00000013076 | Amotl1 | angiomotin-like 1 |

| | | | |
|---|---|---|---|
| RV3 | ENSMUSG00000031196 | F8 | coagulation factor VIII |
| RV3 | ENSMUSG00000015291 | Gdi1 | GDP dissociation inhibitor 1 |
| RV3 | ENSMUSG00000019087 | Atp6ap1 | ATPase, H+ transporting, lysosomal accessory protein 1 |
| RV3 | ENSMUSG00000031391 | L1cam | L1 cell adhesion molecule |
| RV3 | ENSMUSG00000035776 | Cd99l2 | CD99 antigen-like 2 |
| RV3 | ENSMUSG00000036985 | Zdhhc9 | zinc finger, DHHC domain containing 9 |
| RV3 | ENSMUSG00000001173 | Ocrl | OCRL, inositol polyphosphate-5-phosphatase |
| RV3 | ENSMUSG00000055733 | Nap1l3 | nucleosome assembly protein 1-like 3 |
| RV3 | ENSMUSG00000025059 | Gk | glycerol kinase |
| RV3 | ENSMUSG00000025899 | Alkbh8 | alkB homolog 8, tRNA methyltransferase |
| RV3 | ENSMUSG00000050379 | Septin6 | septin 6 |
| RV3 | ENSMUSG00000038344 | Txlng | taxilin gamma |
| RV3 | ENSMUSG00000031377 | Bmx | BMX non-receptor tyrosine kinase |
| RV3 | ENSMUSG00000031342 | Gpm6b | glycoprotein m6b |
| RV3 | ENSMUSG00000000605 | Clcn4 | chloride channel, voltage-sensitive 4 |
| RV3 | ENSMUSG00000041115 | Iqsec2 | IQ motif and Sec7 domain 2 |
| RV3 | ENSMUSG00000025269 | Apex2 | apurinic/apyrimidinic endonuclease 2 |
| RV3 | ENSMUSG00000050148 | Ubqln2 | ubiquilin 2 |
| RV3 | ENSMUSG00000031169 | Porcn | porcupine O-acyltransferase |
| RV3 | ENSMUSG00000064127.1 | Med14 | mediator complex subunit 14 |
| RV3 | ENSMUSG00000040147 | Maob | monoamine oxidase B |
| RV3 | ENSMUSG00000025432 | Avil | advillin |
| RV3 | ENSMUSG00000046841 | Ckap4 | cytoskeleton-associated protein 4 |
| RV3 | ENSMUSG00000060935 | Tmem263 | transmembrane protein 263 |
| RV3 | ENSMUSG00000020038 | Cry1 | cryptochrome circadian regulator 1 |
| RV3 | ENSMUSG00000014850 | Msh3 | mutS homolog 3 |
| RV3 | ENSMUSG00000079477 | Rab7 | RAB7, member RAS oncogene family |
| RV3 | ENSMUSG00000028944 | Prkag2 | protein kinase, AMP-activated, gamma 2 non-catalytic subunit |
| RV3 | ENSMUSG00000054116 | E130116L18Rik | RIKEN cDNA E130116L18 gene |
| RV3 | ENSMUSG00000029836 | Cbx3 | chromobox 3 |
| RV3 | ENSMUSG00000086040 | Wipf3 | WAS/WASL interacting protein family, member 3 |
| RV3 | ENSMUSG00000037788 | Vopp1 | vesicular, overexpressed in cancer, prosurvival protein 1 |
| RV3 | ENSMUSG00000062190 | Lancl2 | LanC (bacterial lantibiotic synthetase component C)-like 2 |
| RV3 | ENSMUSG00000030045 | Mrpl19 | mitochondrial ribosomal protein L19 |
| RV3 | ENSMUSG00000052852 | Reep1 | receptor accessory protein 1 |
| RV3 | ENSMUSG00000053907 | Mat2a | methionine adenosyltransferase 2A |
| RV3 | ENSMUSG00000028173 | Wls | wntless WNT ligand secretion mediator |
| RV3 | ENSMUSG00000028034 | Fubp1 | far upstream element (FUSE) binding protein 1 |
| RV3 | ENSMUSG00000028266 | Lmo4 | LIM domain only 4 |

| | | | |
|---|---|---|---|
| RV3 | ENSMUSG00000028273 | Pdlim5 | PDZ and LIM domain 5 |
| RV3 | ENSMUSG00000028149 | Rap1gds1 | RAP1, GTP-GDP dissociation stimulator 1 |
| RV3 | ENSMUSG00000028159 | Dapp1 | dual adaptor for phosphotyrosine and 3-phosphoinositides 1 |
| RV3 | ENSMUSG00000028161 | Ppp3ca | protein phosphatase 3, catalytic subunit, alpha isoform |
| RV3 | ENSMUSG00000050315 | Synpo2 | synaptopodin 2 |
| RV3 | ENSMUSG00000028125 | Abca4 | ATP-binding cassette, sub-family A member 4 |
| RV3 | ENSMUSG00000027887 | Sypl2 | synaptophysin like 2 |
| RV3 | ENSMUSG00000050947 | Amigo1 | adhesion molecule with Ig like domain 1 |
| RV3 | ENSMUSG00000020463 | Ppp4r3b | protein phosphatase 4 regulatory subunit 3B |
| RV3 | ENSMUSG00000020176 | Grb10 | growth factor receptor bound protein 10 |
| RV3 | ENSMUSG00000020422 | Tns3 | tensin 3 |
| RV3 | ENSMUSG00000002741 | Ykt6 | YKT6 v-SNARE homolog (S. cerevisiae) |
| RV3 | ENSMUSG00000020471 | Pold2 | polymerase (DNA directed), delta 2, regulatory subunit |
| RV3 | ENSMUSG00000009073 | Nf2 | neurofibromin 2 |
| RV3 | ENSMUSG00000020448 | Rnf185 | ring finger protein 185 |
| RV3 | ENSMUSG00000029110 | Rnf4 | ring finger protein 4 |
| RV3 | ENSMUSG00000029106 | Add1 | adducin 1 |
| RV3 | ENSMUSG00000029094 | Afap1 | actin filament associated protein 1 |
| RV3 | ENSMUSG00000055302 | Mrfap1 | Morf4 family associated protein 1 |
| RV3 | ENSMUSG00000005103 | Wdr1 | WD repeat domain 1 |
| RV3 | ENSMUSG00000029089 | 5730480H06Rik | RIKEN cDNA 5730480H06 gene |
| RV3 | ENSMUSG00000033557 | Fam20b | FAM20B, glycosaminoglycan xylosylkinase |
| RV3 | ENSMUSG00000056708 | Ier5 | immediate early response 5 |
| RV3 | ENSMUSG00000042772 | Smg7 | SMG7 nonsense mediated mRNA decay factor |
| RV3 | ENSMUSG00000041801 | Phlda3 | pleckstrin homology like domain, family A, member 3 |
| RV3 | ENSMUSG00000042305 | Tmem183a | transmembrane protein 183A |
| RV3 | ENSMUSG00000041757 | Plekha6 | pleckstrin homology domain containing, family A member 6 |
| RV3 | ENSMUSG00000059149 | Mfsd4a | major facilitator superfamily domain containing 4A |
| RV3 | ENSMUSG00000036155 | Mgat5 | mannoside acetylglucosaminyltransferase 5 |
| RV3 | ENSMUSG00000057967 | Fgf18 | fibroblast growth factor 18 |
| RV3 | ENSMUSG00000020271 | Fbxw11 | F-box and WD-40 domain protein 11 |
| RV3 | ENSMUSG00000024165 | Jpt2 | Jupiter microtubule associated homolog 2 |
| RV3 | ENSMUSG00000034681 | Rnps1 | RNA binding protein with serine rich domain 1 |
| RV3 | ENSMUSG00000024121 | Atp6v0c | ATPase, H+ transporting, lysosomal V0 subunit C |
| RV3 | ENSMUSG00000040097 | Flywch1 | FLYWCH-type zinc finger 1 |
| RV3 | ENSMUSG00000023909 | Paqr4 | progestin and adipoQ receptor family member IV |
| RV3 | ENSMUSG00000022519 | Srl | sarcalumenin |
| RV3 | ENSMUSG00000022505 | Emp2 | epithelial membrane protein 2 |
| RV3 | ENSMUSG00000062203 | Gspt1 | G1 to S phase transition 1 |

| | | | |
|---|---|---|---|
| RV3 | ENSMUSG00000071669 | Snx29 | sorting nexin 29 |
| RV3 | ENSMUSG00000065979 | Cpped1 | calcineurin-like phosphoesterase domain containing 1 |
| RV3 | ENSMUSG00000022545 | Ercc4 | excision repair cross-complementing rodent repair deficiency, complementation group 4 |
| RV3 | ENSMUSG00000037904 | Ankrd9 | ankyrin repeat domain 9 |
| RV3 | ENSMUSG00000037395 | Rcor3 | REST corepressor 3 |
| RV3 | ENSMUSG00000046836 | Brox | BRO1 domain and CAAX motif containing |
| RV3 | ENSMUSG00000062169 | Cnih4 | cornichon family AMPA receptor auxiliary protein 4 |
| RV3 | ENSMUSG00000063659 | Zbtb18 | zinc finger and BTB domain containing 18 |
| RV3 | ENSMUSG00000052423 | B4galt3 | UDP-Gal:betaGlcNAc beta 1,4-galactosyltransferase, polypeptide 3 |
| RV3 | ENSMUSG00000026576 | Atp1b1 | ATPase, Na+/K+ transporting, beta 1 polypeptide |
| RV3 | ENSMUSG00000025340 | Rabgef1 | RAB guanine nucleotide exchange factor (GEF) 1 |
| RV3 | ENSMUSG00000025532 | Crcp | calcitonin gene-related peptide-receptor component protein |
| RV3 | ENSMUSG00000066735 | Vkorc1l1 | vitamin K epoxide reductase complex, subunit 1-like 1 |
| RV3 | ENSMUSG00000029482 | Aacs | acetoacetyl-CoA synthetase |
| RV3 | ENSMUSG00000079215 | Zfp664 | zinc finger protein 664 |
| RV3 | ENSMUSG00000038582 | Pptc7 | PTC7 protein phosphatase homolog |
| RV3 | ENSMUSG00000018604 | Tbx3 | T-box 3 |
| RV3 | ENSMUSG00000032867 | Fbxw8 | F-box and WD-40 domain protein 8 |
| RV3 | ENSMUSG00000029518 | Rab35 | RAB35, member RAS oncogene family |
| RV3 | ENSMUSG00000009013 | Dynll1 | dynein light chain LC8-type 1 |
| RV3 | ENSMUSG00000048578 | Mlec | malectin |
| RV3 | ENSMUSG00000029575 | Mmab | methylmalonic aciduria (cobalamin deficiency) cblB type homolog (human) |
| RV3 | ENSMUSG00000040731 | Eif4h | eukaryotic translation initiation factor 4H |
| RV3 | ENSMUSG00000029675 | Eln | elastin |
| RV3 | ENSMUSG00000051391 | Ywhag | tyrosine 3-monooxygenase/tryptophan 5-monooxygenase activation protein, gamma polypeptide |
| RV3 | ENSMUSG00000029705 | Cux1 | cut-like homeobox 1 |
| RV3 | ENSMUSG00000004846 | Plod3 | procollagen-lysine, 2-oxoglutarate 5-dioxygenase 3 |
| RV3 | ENSMUSG00000023348 | Trip6 | thyroid hormone receptor interactor 6 |
| RV3 | ENSMUSG00000075593 | Gal3st4 | galactose-3-O-sulfotransferase 4 |
| RV3 | ENSMUSG00000036968 | Cnpy4 | canopy FGF signaling regulator 4 |
| RV3 | ENSMUSG00000000149 | Gna12 | guanine nucleotide binding protein, alpha 12 |
| RV3 | ENSMUSG00000038780 | Smurf1 | SMAD specific E3 ubiquitin protein ligase 1 |
| RV3 | ENSMUSG00000068566 | Myadm | myeloid-associated differentiation marker |
| RV3 | ENSMUSG00000063802 | Hspbp1 | HSPA (heat shock 70kDa) binding protein, cytoplasmic cochaperone 1 |
| RV3 | ENSMUSG00000035203 | Epn1 | epsin 1 |
| RV3 | ENSMUSG00000033961 | Zfp446 | zinc finger protein 446 |
| RV3 | ENSMUSG00000058230 | Arhgap35 | Rho GTPase activating protein 35 |

| | | | |
|---|---|---|---|
| RV3 | ENSMUSG00000040511 | Pvr | poliovirus receptor |
| RV3 | ENSMUSG00000058402 | Zfp420 | zinc finger protein 420 |
| RV3 | ENSMUSG00000036864 | Proser3 | proline and serine rich 3 |
| RV3 | ENSMUSG00000009687 | Fxyd5 | FXYD domain-containing ion transport regulator 5 |
| RV3 | ENSMUSG00000066568 | Lsm14a | LSM14A mRNA processing body assembly factor |
| RV3 | ENSMUSG00000006763 | Saal1 | serum amyloid A-like 1 |
| RV3 | ENSMUSG00000030824 | Nucb1 | nucleobindin 1 |
| RV3 | ENSMUSG00000030796 | Tead2 | TEA domain family member 2 |
| RV3 | ENSMUSG00000003184 | Irf3 | interferon regulatory factor 3 |
| RV3 | ENSMUSG00000109511 | Nup62 | nucleoporin 62 |
| RV3 | ENSMUSG00000070738 | Dgkd | diacylglycerol kinase, delta |
| RV3 | ENSMUSG00000026308 | Klhl30 | kelch-like 30 |
| RV3 | ENSMUSG00000007805 | Twist2 | twist basic helix-loop-helix transcription factor 2 |
| RV3 | ENSMUSG00000026335 | Pam | peptidylglycine alpha-amidating monooxygenase |
| RV3 | ENSMUSG00000030655 | Smg1 | SMG1 nonsense mediated mRNA decay associated PI3K related kinase |
| RV3 | ENSMUSG00000045659 | Plekha7 | pleckstrin homology domain containing, family A member 7 |
| RV3 | ENSMUSG00000038156 | Spon1 | spondin 1, (f-spondin) extracellular matrix protein |
| RV3 | ENSMUSG00000038244 | Mical2 | microtubule associated monooxygenase, calponin and LIM domain containing 2 |
| RV3 | ENSMUSG00000034825 | Nrip3 | nuclear receptor interacting protein 3 |
| RV3 | ENSMUSG00000030704 | Rab6a | RAB6A, member RAS oncogene family |
| RV3 | ENSMUSG00000035227 | Spcs2 | signal peptidase complex subunit 2 homolog (S. cerevisiae) |
| RV3 | ENSMUSG00000070436 | Serpinh1 | serine (or cysteine) peptidase inhibitor, clade H, member 1 |
| RV3 | ENSMUSG00000090958 | Lrrc32 | leucine rich repeat containing 32 |
| RV3 | ENSMUSG00000035704 | Alg8 | ALG8 alpha-1,3-glucosyltransferase |
| RV3 | ENSMUSG00000035713 | Usp35 | ubiquitin specific peptidase 35 |
| RV3 | ENSMUSG00000005836 | Gata6 | GATA binding protein 6 |
| RV3 | ENSMUSG00000037013 | Ss18 | SS18, subunit of BAF chromatin remodeling complex |
| RV3 | ENSMUSG00000071533 | Pcnp | PEST proteolytic signal containing nuclear protein |
| RV3 | ENSMUSG00000035258 | Abi3bp | ABI family member 3 binding protein |
| RV38 | ENSMUSG00000063268 | Parp10 | poly (ADP-ribose) polymerase family, member 10 |
| RV38 | ENSMUSG00000091780 | Sco2 | SCO2 cytochrome c oxidase assembly protein |
| RV38 | ENSMUSG00000049382 | Krt8 | keratin 8 |
| RV38 | ENSMUSG00000023043 | Krt18 | keratin 18 |
| RV38 | ENSMUSG00000036323 | Srp72 | signal recognition particle 72 |
| RV38 | ENSMUSG00000051674 | Dcun1d4 | defective in cullin neddylation 1 domain containing 4 |
| RV38 | ENSMUSG00000037795 | N4bp2 | NEDD4 binding protein 2 |
| RV38 | ENSMUSG00000029173 | Sepsecs | Sep (O-phosphoserine) tRNA:Sec (selenocysteine) tRNA synthase |
| RV38 | ENSMUSG00000027397 | Slc20a1 | solute carrier family 20, member 1 |

| | | | |
|---|---|---|---|
| RV38 | ENSMUSG00000043110 | Lrrn4 | leucine rich repeat neuronal 4 |
| RV38 | ENSMUSG00000005881 | Ergic3 | ERGIC and golgi 3 |
| RV38 | ENSMUSG00000027637 | Rab5if | RAB5 interacting factor |
| RV38 | ENSMUSG00000017734 | Dbndd2 | dysbindin domain containing 2 |
| RV38 | ENSMUSG00000047907 | Tshz2 | teashirt zinc finger family member 2 |
| RV38 | ENSMUSG00000015093 | Clic3 | chloride intracellular channel 3 |
| RV38 | ENSMUSG00000035949 | Fbxw2 | F-box and WD-40 domain protein 2 |
| RV38 | ENSMUSG00000022763 | Aifm3 | apoptosis-inducing factor, mitochondrion-associated 3 |
| RV38 | ENSMUSG00000003235 | Eif2b5 | eukaryotic translation initiation factor 2B, subunit 5 epsilon |
| RV38 | ENSMUSG00000017291 | Taok1 | TAO kinase 1 |
| RV38 | ENSMUSG00000018559 | Ctdnep1 | CTD nuclear envelope phosphatase 1 |
| RV38 | ENSMUSG00000020904 | Cfap52 | cilia and flagella associated protein 52 |
| RV38 | ENSMUSG00000028860 | Sytl1 | synaptotagmin-like 1 |
| RV38 | ENSMUSG00000037306 | Man1c1 | mannosidase, alpha, class 1C, member 1 |
| RV38 | ENSMUSG00000029003 | Mad2l2 | MAD2 mitotic arrest deficient-like 2 |
| RV38 | ENSMUSG00000031970 | Dbndd1 | dysbindin domain containing 1 |
| RV38 | ENSMUSG00000002227 | Mov10 | Mov10 RISC complex RNA helicase |
| RV38 | ENSMUSG00000005370 | Msh6 | mutS homolog 6 |
| RV38 | ENSMUSG00000024841 | Eif1ad | eukaryotic translation initiation factor 1A domain containing |
| RV38 | ENSMUSG00000032381 | Ciao2a | cytosolic iron-sulfur assembly component 2A |
| RV38 | ENSMUSG00000027559 | Car3 | carbonic anhydrase 3 |
| RV38 | ENSMUSG00000021194 | Chga | chromogranin A |
| RV38 | ENSMUSG00000021236 | Entpd5 | ectonucleoside triphosphate diphosphohydrolase 5 |
| RV38 | ENSMUSG00000041831 | Sytl3 | synaptotagmin-like 3 |
| RV38 | ENSMUSG00000015659 | Serac1 | serine active site containing 1 |
| RV38 | ENSMUSG00000041763 | Tpp2 | tripeptidyl peptidase II |
| RV38 | ENSMUSG00000028082 | Sh3d19 | SH3 domain protein D19 |
| RV38 | ENSMUSG00000020911 | Krt19 | keratin 19 |
| RV38 | ENSMUSG00000002871 | Tpra1 | transmembrane protein, adipocyte asscociated 1 |
| RV38 | ENSMUSG00000060038 | Dhps | deoxyhypusine synthase |
| RV38 | ENSMUSG00000052926 | Rnaseh2a | ribonuclease H2, large subunit |
| RV38 | ENSMUSG00000004996 | Mri1 | methylthioribose-1-phosphate isomerase 1 |
| RV38 | ENSMUSG00000109941 | Exosc6 | exosome component 6 |
| RV38 | ENSMUSG00000012519 | Mlkl | mixed lineage kinase domain-like |
| RV38 | ENSMUSG00000041840 | Haus1 | HAUS augmin-like complex, subunit 1 |
| RV38 | ENSMUSG00000021795 | Sftpd | surfactant associated protein D |
| RV38 | ENSMUSG00000021775 | Nr1d2 | nuclear receptor subfamily 1, group D, member 2 |
| RV38 | ENSMUSG00000040717 | Il17rd | interleukin 17 receptor D |
| RV38 | ENSMUSG00000030551 | Nr2f2 | nuclear receptor subfamily 2, group F, member 2 |

| | | | |
|---|---|---|---|
| RV38 | ENSMUSG00000022867 | Usp25 | ubiquitin specific peptidase 25 |
| RV38 | ENSMUSG00000013787 | Ehmt2 | euchromatic histone lysine N-methyltransferase 2 |
| RV38 | ENSMUSG00000024371 | C2 | complement C2 |
| RV38 | ENSMUSG00000090231 | Cfb | complement factor B |
| RV38 | ENSMUSG00000061353 | Cxcl12 | C-X-C motif chemokine ligand 12 |
| RV38 | ENSMUSG00000024104 | Washc2 | WASH complex subunit 2 |
| RV38 | ENSMUSG00000064177 | Ghrl | ghrelin |
| RV38 | ENSMUSG00000030281 | Il17rc | interleukin 17 receptor C |
| RV38 | ENSMUSG00000037845 | Fdxacb1 | ferredoxin-fold anticodon binding domain containing 1 |
| RV38 | ENSMUSG00000067873 | Htatsf1 | HIV TAT specific factor 1 |
| RV38 | ENSMUSG00000025351 | Cd63 | CD63 antigen |
| RV38 | ENSMUSG00000034602 | Mon2 | MON2 homolog, regulator of endosome to Golgi trafficking |
| RV38 | ENSMUSG00000027901.1 | Dennd2d | DENN domain containing 2D |
| RV38 | ENSMUSG00000036693 | Nop14 | NOP14 nucleolar protein |
| RV38 | ENSMUSG00000063011 | Msln | mesothelin |
| RV38 | ENSMUSG00000035521 | Gnptg | N-acetylglucosamine-1-phosphotransferase, gamma subunit |
| RV38 | ENSMUSG00000024168 | Tmem204 | transmembrane protein 204 |
| RV38 | ENSMUSG00000008393 | Carhsp1 | calcium regulated heat stable protein 1 |
| RV38 | ENSMUSG00000041609 | Bicdl1 | BICD family like cargo adaptor 1 |
| RV38 | ENSMUSG00000041939 | Mvk | mevalonate kinase |
| RV38 | ENSMUSG00000042985 | Upk3b | uroplakin 3B |
| RV38 | ENSMUSG00000001739 | Cldn15 | claudin 15 |
| RV38 | ENSMUSG00000002635 | Pdcd2l | programmed cell death 2-like |
| RV38 | ENSMUSG00000003873 | Bax | BCL2-associated X protein |
| RV38 | ENSMUSG00000045411 | 2410002F23Rik | RIKEN cDNA 2410002F23 gene |
| RV38 | ENSMUSG00000026278 | Bok | BCL2-related ovarian killer |
| RV38 | ENSMUSG00000049436 | Upk1b | uroplakin 1B |
| RV38 | ENSMUSG00000022663 | Atg3 | autophagy related 3 |
| RV49 | ENSMUSG00000062373 | Tmem65 | transmembrane protein 65 |
| RV49 | ENSMUSG00000060992 | Copz1 | coatomer protein complex, subunit zeta 1 |
| RV49 | ENSMUSG00000027194 | Ttc17 | tetratricopeptide repeat domain 17 |
| RV49 | ENSMUSG00000027180 | Fbxo3 | F-box protein 3 |
| RV49 | ENSMUSG00000027287 | Snap23 | synaptosomal-associated protein 23 |
| RV49 | ENSMUSG00000068040 | Tm9sf4 | transmembrane 9 superfamily member 4 |
| RV49 | ENSMUSG00000050043 | Tmx2 | thioredoxin-related transmembrane protein 2 |
| RV49 | ENSMUSG00000042369 | Rbm45 | RNA binding motif protein 45 |
| RV49 | ENSMUSG00000015083 | C8g | complement component 8, gamma polypeptide |
| RV49 | ENSMUSG00000069020 | Urm1 | ubiquitin related modifier 1 |
| RV49 | ENSMUSG00000026880 | Stom | stomatin |

| | | | |
|---|---|---|---|
| RV49 | ENSMUSG00000015994 | Fnta | farnesyltransferase, CAAX box, alpha |
| RV49 | ENSMUSG00000017686 | Rhot1 | ras homolog family member T1 |
| RV49 | ENSMUSG00000046731 | Kctd11 | potassium channel tetramerisation domain containing 11 |
| RV49 | ENSMUSG00000020898 | Ctc1 | CTS telomere maintenance complex component 1 |
| RV49 | ENSMUSG00000042298 | Ttc19 | tetratricopeptide repeat domain 19 |
| RV49 | ENSMUSG00000036860 | Mrpl55 | mitochondrial ribosomal protein L55 |
| RV49 | ENSMUSG00000028583 | Pdpn | podoplanin |
| RV49 | ENSMUSG00000044496 | 2510039O18Rik | RIKEN cDNA 2510039O18 gene |
| RV49 | ENSMUSG00000039768 | Dnajc11 | DnaJ heat shock protein family (Hsp40) member C11 |
| RV49 | ENSMUSG00000006517 | Mvd | mevalonate (diphospho) decarboxylase |
| RV49 | ENSMUSG00000020641 | Rsad2 | radical S-adenosyl methionine domain containing 2 |
| RV49 | ENSMUSG00000054469 | Lclat1 | lysocardiolipin acyltransferase 1 |
| RV49 | ENSMUSG00000024069 | Slc30a6 | solute carrier family 30 (zinc transporter), member 6 |
| RV49 | ENSMUSG00000002017 | Fam98a | family with sequence similarity 98, member A |
| RV49 | ENSMUSG00000024732 | Ccdc86 | coiled-coil domain containing 86 |
| RV49 | ENSMUSG00000038801 | Scgb1c1 | secretoglobin, family 1C, member 1 |
| RV49 | ENSMUSG00000038775 | Vill | villin-like |
| RV49 | ENSMUSG00000042487 | Leo1 | Leo1, Paf1/RNA polymerase II complex component |
| RV49 | ENSMUSG00000032192 | Gnb5 | guanine nucleotide binding protein (G protein), beta 5 |
| RV49 | ENSMUSG00000022044 | Stmn4 | stathmin-like 4 |
| RV49 | ENSMUSG00000021975 | Ints9 | integrator complex subunit 9 |
| RV49 | ENSMUSG00000021939 | Ctsb | cathepsin B |
| RV49 | ENSMUSG00000021824 | Ap3m1 | adaptor-related protein complex 3, mu 1 subunit |
| RV49 | ENSMUSG00000015733 | Capza2 | capping actin protein of muscle Z-line subunit alpha 2 |
| RV49 | ENSMUSG00000038305 | Spats2l | spermatogenesis associated, serine-rich 2-like |
| RV49 | ENSMUSG00000039703 | Nploc4 | NPL4 homolog, ubiquitin recognition factor |
| RV49 | ENSMUSG00000001493 | Meox1 | mesenchyme homeobox 1 |
| RV49 | ENSMUSG00000018882 | Mrpl45 | mitochondrial ribosomal protein L45 |
| RV49 | ENSMUSG00000018171 | Vmp1 | vacuole membrane protein 1 |
| RV49 | ENSMUSG00000023988 | Bysl | bystin-like |
| RV49 | ENSMUSG00000033545 | Znrf1 | zinc and ring finger 1 |
| RV49 | ENSMUSG00000024583 | Txnl1 | thioredoxin-like 1 |
| RV49 | ENSMUSG00000053644 | Aldh7a1 | aldehyde dehydrogenase family 7, member A1 |
| RV49 | ENSMUSG00000024528 | Srfbp1 | serum response factor binding protein 1 |
| RV49 | ENSMUSG00000030095 | Tmem43 | transmembrane protein 43 |
| RV49 | ENSMUSG00000027698 | Nceh1 | neutral cholesterol ester hydrolase 1 |
| RV49 | ENSMUSG00000033444 | Specc1l | sperm antigen with calponin homology and coiled-coil domains 1-like |
| RV49 | ENSMUSG00000021427 | Ssr1 | signal sequence receptor, alpha |

| | | | |
|---|---|---|---|
| RV49 | ENSMUSG00000021484 | Lman2 | lectin, mannose-binding 2 |
| RV49 | ENSMUSG00000030110 | Ret | ret proto-oncogene |
| RV49 | ENSMUSG00000028214 | Gem | GTP binding protein overexpressed in skeletal muscle |
| RV49 | ENSMUSG00000041632 | Mrps27 | mitochondrial ribosomal protein S27 |
| RV49 | ENSMUSG00000021711 | Trappc13 | trafficking protein particle complex 13 |
| RV49 | ENSMUSG00000020029 | Nudt4 | nudix hydrolase 4 |
| RV49 | ENSMUSG00000035840 | Lysmd3 | LysM, putative peptidoglycan-binding, domain containing 3 |
| RV49 | ENSMUSG00000029804 | Herc3 | hect domain and RLD 3 |
| RV49 | ENSMUSG00000020134 | Peli1 | pellino 1 |
| RV49 | ENSMUSG00000044066 | Cep68 | centrosomal protein 68 |
| RV49 | ENSMUSG00000060519 | Tor3a | torsin family 3, member A |
| RV49 | ENSMUSG00000026475 | Rgs16 | regulator of G-protein signaling 16 |
| RV49 | ENSMUSG00000032649 | Colgalt2 | collagen beta(1-O)galactosyltransferase 2 |
| RV49 | ENSMUSG00000026634 | Angel2 | angel homolog 2 |
| RV49 | ENSMUSG00000109324 | Prmt1 | protein arginine N-methyltransferase 1 |
| RV49 | ENSMUSG00000022742 | Cpox | coproporphyrinogen oxidase |
| RV49 | ENSMUSG00000022744 | Cldnd1 | claudin domain containing 1 |
| RV63 | ENSMUSG00000035505 | Cox18 | cytochrome c oxidase assembly protein 18 |
| RV63 | ENSMUSG00000027605 | Acss2 | acyl-CoA synthetase short-chain family member 2 |
| RV63 | ENSMUSG00000039501 | Znfx1 | zinc finger, NFX1-type containing 1 |
| RV63 | ENSMUSG00000026922 | Agpat2 | 1-acylglycerol-3-phosphate O-acyltransferase 2 |
| RV63 | ENSMUSG00000026853 | Crat | carnitine acetyltransferase |
| RV63 | ENSMUSG00000028563 | Tm2d1 | TM2 domain containing 1 |
| RV63 | ENSMUSG00000037958 | Nsrp1 | nuclear speckle regulatory protein 1 |
| RV63 | ENSMUSG00000002812 | Flii | flightless I actin binding protein |
| RV63 | ENSMUSG00000040740 | Slc25a34 | solute carrier family 25, member 34 |
| RV63 | ENSMUSG00000020628 | Trappc12 | trafficking protein particle complex 12 |
| RV63 | ENSMUSG00000059447 | Hadhb | hydroxyacyl-CoA dehydrogenase trifunctional multienzyme complex subunit beta |
| RV63 | ENSMUSG00000024867 | Pip5k1b | phosphatidylinositol-4-phosphate 5-kinase, type 1 beta |
| RV63 | ENSMUSG00000024797 | Vps51 | VPS51 GARP complex subunit |
| RV63 | ENSMUSG00000024851 | Pitpnm1 | phosphatidylinositol transfer protein, membrane-associated 1 |
| RV63 | ENSMUSG00000020253 | Ppm1m | protein phosphatase 1M |
| RV63 | ENSMUSG00000022014 | Epsti1 | epithelial stromal interaction 1 |
| RV63 | ENSMUSG00000000776 | Polr3d | polymerase (RNA) III (DNA directed) polypeptide D |
| RV63 | ENSMUSG00000029695 | Aass | aminoadipate-semialdehyde synthase |
| RV63 | ENSMUSG00000064127 | Med14 | mediator complex subunit 14 |
| RV63 | ENSMUSG00000021259 | Cyp46a1 | cytochrome P450, family 46, subfamily a, polypeptide 1 |
| RV63 | ENSMUSG00000019854 | Reps1 | RalBP1 associated Eps domain containing protein |

| | | | |
|---|---|---|---|
| RV63 | ENSMUSG00000048756 | Foxo3 | forkhead box O3 |
| RV63 | ENSMUSG00000004151 | Etv1 | ets variant 1 |
| RV63 | ENSMUSG00000025155 | Dus1l | dihydrouridine synthase 1 like |
| RV63 | ENSMUSG00000025142 | Aspscr1 | ASPSCR1 tether for SLC2A4, UBX domain containing |
| RV63 | ENSMUSG00000020777 | Acox1 | acyl-Coenzyme A oxidase 1, palmitoyl |
| RV63 | ENSMUSG00000075528 | Aarsd1 | alanyl-tRNA synthetase domain containing 1 |
| RV63 | ENSMUSG00000002763 | Pex6 | peroxisomal biogenesis factor 6 |
| RV63 | ENSMUSG00000034467 | Dynlrb2 | dynein light chain roadblock-type 2 |
| RV63 | ENSMUSG00000035765 | Dym | dymeclin |
| RV63 | ENSMUSG00000057606 | Colq | collagen like tail subunit of asymmetric acetylcholinesterase |
| RV63 | ENSMUSG00000021918 | Nek4 | NIMA (never in mitosis gene a)-related expressed kinase 4 |
| RV63 | ENSMUSG00000039187 | Fanci | Fanconi anemia, complementation group I |
| RV63 | ENSMUSG00000048234 | Rnf149 | ring finger protein 149 |
| RV63 | ENSMUSG00000049422 | Chchd10 | coiled-coil-helix-coiled-coil-helix domain containing 10 |
| RV63 | ENSMUSG00000054021 | Sirt5 | sirtuin 5 |
| RV63 | ENSMUSG00000052253 | Zfp622 | zinc finger protein 622 |
| RV63 | ENSMUSG00000034617 | Mtrr | 5-methyltetrahydrofolate-homocysteine methyltransferase reductase |
| RV63 | ENSMUSG00000050697 | Prkaa1 | protein kinase, AMP-activated, alpha 1 catalytic subunit |
| RV63 | ENSMUSG00000036411 | Matcap2 | microtubule associated tyrosine carboxypeptidase 2 |
| RV63 | ENSMUSG00000002820 | Atg4d | autophagy related 4D, cysteine peptidase |
| RV63 | ENSMUSG00000031066 | Usp11 | ubiquitin specific peptidase 11 |
| RV63 | ENSMUSG00000069539 | Scyl2 | SCY1-like 2 (S. cerevisiae) |
| RV63 | ENSMUSG00000090066 | 1110002E22Rik | RIKEN cDNA 1110002E22 gene |
| RV63 | ENSMUSG00000020309 | Chac2 | ChaC, cation transport regulator 2 |
| RV63 | ENSMUSG00000049811 | Fam161a | family with sequence similarity 161, member A |
| RV63 | ENSMUSG00000002504 | Nherf2 | NHERF family PDZ scaffold protein 2 |
| RV63 | ENSMUSG00000002496 | Tsc2 | TSC complex subunit 2 |
| RV63 | ENSMUSG00000024132 | Eci1 | enoyl-Coenzyme A delta isomerase 1 |
| RV63 | ENSMUSG00000022515 | Anks3 | ankyrin repeat and sterile alpha motif domain containing 3 |
| RV63 | ENSMUSG00000062729 | Ppox | protoporphyrinogen oxidase |
| RV63 | ENSMUSG00000005514 | Por | cytochrome p450 oxidoreductase |
| RV63 | ENSMUSG00000040811 | Eml2 | echinoderm microtubule associated protein like 2 |
| RV63 | ENSMUSG00000040428 | Plekha4 | pleckstrin homology domain containing, family A (phosphoinositide binding specific) member 4 |
| RV63 | ENSMUSG00000036989 | Trim3 | tripartite motif-containing 3 |
| RV70 | ENSMUSG00000023052 | Npff | neuropeptide FF-amide peptide precursor |
| RV70 | ENSMUSG00000097392 | Thoc2l | THO complex subunit 2-like |
| RV70 | ENSMUSG00000042745 | Id1 | inhibitor of DNA binding 1, HLH protein |
| RV70 | ENSMUSG00000057133 | Chd6 | chromodomain helicase DNA binding protein 6 |

| | | | |
|---|---|---|---|
| RV70 | ENSMUSG00000000826 | Dnajc5 | DnaJ heat shock protein family (Hsp40) member C5 |
| RV70 | ENSMUSG00000027007 | Itprid2 | ITPR interacting domain containing 2 |
| RV70 | ENSMUSG00000068882 | Ssb | small RNA binding exonuclease protection factor La |
| RV70 | ENSMUSG00000036202 | Rif1 | replication timing regulatory factor 1 |
| RV70 | ENSMUSG00000039356 | Exosc2 | exosome component 2 |
| RV70 | ENSMUSG00000034210 | Efcab14 | EF-hand calcium binding domain 14 |
| RV70 | ENSMUSG00000037857 | Nufip2 | nuclear FMR1 interacting protein 2 |
| RV70 | ENSMUSG00000020359 | Phykpl | 5-phosphohydroxy-L-lysine phospholyase |
| RV70 | ENSMUSG00000057572 | Zbtb8os | zinc finger and BTB domain containing 8 opposite strand |
| RV70 | ENSMUSG00000025200 | Cwf19l1 | CWF19 like cell cycle control factor 1 |
| RV70 | ENSMUSG00000024695 | Zfp91 | zinc finger protein 91 |
| RV70 | ENSMUSG00000049734 | Trex1 | three prime repair exonuclease 1 |
| RV70 | ENSMUSG00000032212 | Sltm | SAFB-like, transcription modulator |
| RV70 | ENSMUSG00000021981 | Cab39l | calcium binding protein 39-like |
| RV70 | ENSMUSG00000056458 | Mok | MOK protein kinase |
| RV70 | ENSMUSG00000010608 | Rbm25 | RNA binding motif protein 25 |
| RV70 | ENSMUSG00000048118 | Arid4a | AT-rich interaction domain 4A |
| RV70 | ENSMUSG00000004698 | Hdac9 | histone deacetylase 9 |
| RV70 | ENSMUSG00000008763 | Man1a2 | mannosidase, alpha, class 1A, member 2 |
| RV70 | ENSMUSG00000040481 | Bptf | bromodomain PHD finger transcription factor |
| RV70 | ENSMUSG00000020541 | Tom1l1 | target of myb1-like 1 (chicken) |
| RV70 | ENSMUSG00000031660 | Brd7 | bromodomain containing 7 |
| RV70 | ENSMUSG00000003847 | Nfat5 | nuclear factor of activated T cells 5 |
| RV70 | ENSMUSG00000038538 | Ubn2 | ubinuclein 2 |
| RV70 | ENSMUSG00000078671 | Chd2 | chromodomain helicase DNA binding protein 2 |
| RV70 | ENSMUSG00000036513 | Commd2 | COMM domain containing 2 |
| RV70 | ENSMUSG00000027722 | Afg2a | AFG2 AAA ATPase homolog A |
| RV70 | ENSMUSG00000040785 | Ttc3 | tetratricopeptide repeat domain 3 |
| RV70 | ENSMUSG00000019947 | Arid5b | AT-rich interaction domain 5B |
| RV70 | ENSMUSG00000039219 | Arid4b | AT-rich interaction domain 4B |
| RV70 | ENSMUSG00000035367 | Rmi1 | RecQ mediated genome instability 1 |
| RV70 | ENSMUSG00000008540 | Mgst1 | microsomal glutathione S-transferase 1 |
| RV70 | ENSMUSG00000030180 | Kdm5a | lysine demethylase 5A |
| RV70 | ENSMUSG00000022141 | Nipbl | NIPBL cohesin loading factor |
| RV70 | ENSMUSG00000039704 | Lmbrd2 | LMBR1 domain containing 2 |
| RV70 | ENSMUSG00000032041 | Tirap | toll-interleukin 1 receptor (TIR) domain-containing adaptor protein |
| RV70 | ENSMUSG00000061315 | Naca | nascent polypeptide-associated complex alpha polypeptide |
| RV70 | ENSMUSG00000033991 | Skic3 | SKI3 subunit of superkiller complex |
| RV70 | ENSMUSG00000032740 | Ccdc88a | coiled coil domain containing 88A |

| | | | |
|---|---|---|---|
| RV70 | ENSMUSG00000029127 | Zbtb49 | zinc finger and BTB domain containing 49 |
| RV70 | ENSMUSG00000061755 | Bod1l | biorientation of chromosomes in cell division 1-like |
| RV70 | ENSMUSG00000026600 | Soat1 | sterol O-acyltransferase 1 |
| RV70 | ENSMUSG00000040181 | Fmo1 | flavin containing monooxygenase 1 |
| RV70 | ENSMUSG00000040423 | Rc3h1 | RING CCCH (C3H) domains 1 |
| RV70 | ENSMUSG00000002210 | Smg9 | SMG9 nonsense mediated mRNA decay factor |
| RV70 | ENSMUSG00000034647 | Ankrd12 | ankyrin repeat domain 12 |
| RV70 | ENSMUSG00000024290 | Rock1 | Rho-associated coiled-coil containing protein kinase 1 |
| RV71 | ENSMUSG00000022339 | Ebag9 | estrogen receptor-binding fragment-associated gene 9 |
| RV71 | ENSMUSG00000046380 | Jrk | jerky |
| RV71 | ENSMUSG00000022401 | Xpnpep3 | X-prolyl aminopeptidase 3, mitochondrial |
| RV71 | ENSMUSG00000055782 | Abcd2 | ATP-binding cassette, sub-family D member 2 |
| RV71 | ENSMUSG00000063296 | Tmem117 | transmembrane protein 117 |
| RV71 | ENSMUSG00000027223 | Mapk8ip1 | mitogen-activated protein kinase 8 interacting protein 1 |
| RV71 | ENSMUSG00000067818 | Myl9 | myosin, light polypeptide 9, regulatory |
| RV71 | ENSMUSG00000039476 | Prrx2 | paired related homeobox 2 |
| RV71 | ENSMUSG00000073792 | Alg6 | ALG6 alpha-1,3-glucosyltransferase |
| RV71 | ENSMUSG00000038457 | Tmem255b | transmembrane protein 255B |
| RV71 | ENSMUSG00000020520 | Galnt10 | polypeptide N-acetylgalactosaminyltransferase 10 |
| RV71 | ENSMUSG00000028759 | Hp1bp3 | heterochromatin protein 1, binding protein 3 |
| RV71 | ENSMUSG00000031972 | Acta1 | actin alpha 1, skeletal muscle |
| RV71 | ENSMUSG00000031825 | Crispld2 | cysteine-rich secretory protein LCCL domain containing 2 |
| RV71 | ENSMUSG00000027858 | Tspan2 | tetraspanin 2 |
| RV71 | ENSMUSG00000025470 | Zfp511 | zinc finger protein 511 |
| RV71 | ENSMUSG00000032215 | Rsl24d1 | ribosomal L24 domain containing 1 |
| RV71 | ENSMUSG00000022035 | Ccdc25 | coiled-coil domain containing 25 |
| RV71 | ENSMUSG00000034532 | Fbxo16 | F-box protein 16 |
| RV71 | ENSMUSG00000039367 | Sec24c | SEC24 homolog C, COPII coat complex component |
| RV71 | ENSMUSG00000028464 | Tpm2 | tropomyosin 2, beta |
| RV71 | ENSMUSG00000029757 | Dync1i1 | dynein cytoplasmic 1 intermediate chain 1 |
| RV71 | ENSMUSG00000021103 | Mnat1 | menage a trois 1 |
| RV71 | ENSMUSG00000014771 | Pdcd2 | programmed cell death 2 |
| RV71 | ENSMUSG00000038240 | Pdss2 | prenyl (solanesyl) diphosphate synthase, subunit 2 |
| RV71 | ENSMUSG00000026163 | Sphkap | SPHK1 interactor, AKAP domain containing |
| RV71 | ENSMUSG00000039253 | Fn3krp | fructosamine 3 kinase related protein |
| RV71 | ENSMUSG00000042215 | Bag2 | BCL2-associated athanogene 2 |
| RV71 | ENSMUSG00000024542 | Cep192 | centrosomal protein 192 |
| RV71 | ENSMUSG00000018999 | Slc35b4 | solute carrier family 35, member B4 |
| RV71 | ENSMUSG00000041445 | Mmrn2 | multimerin 2 |

| | | | |
|---|---|---|---|
| RV71 | ENSMUSG00000045795 | Whamm | WAS protein homolog associated with actin, golgi membranes and microtubules |
| RV71 | ENSMUSG00000070520 | Nsmce3 | NSE3 homolog, SMC5-SMC6 complex component |
| RV71 | ENSMUSG00000022947 | Cbr3 | carbonyl reductase 3 |
| RV71 | ENSMUSG00000039046 | Usp6nl | USP6 N-terminal like |
| RV71 | ENSMUSG00000021196 | Pfkp | phosphofructokinase, platelet |
| RV71 | ENSMUSG00000032085 | Tagln | transgelin |
| RV71 | ENSMUSG00000037419 | Endod1 | endonuclease domain containing 1 |
| RV71 | ENSMUSG00000025374 | Nabp2 | nucleic acid binding protein 2 |
| RV71 | ENSMUSG00000040043 | Rbms2 | RNA binding motif, single stranded interacting protein 2 |
| RV71 | ENSMUSG00000025410 | Dctn2 | dynactin 2 |
| RV71 | ENSMUSG00000029111 | Nelfa | negative elongation factor complex member A, Whsc2 |
| RV71 | ENSMUSG00000018830 | Myh11 | myosin, heavy polypeptide 11, smooth muscle |
| RV71 | ENSMUSG00000029500 | Pgam5 | phosphoglycerate mutase family member 5 |
| RV71 | ENSMUSG00000052833 | Sae1 | SUMO1 activating enzyme subunit 1 |
| RV71 | ENSMUSG00000030603 | Psmc4 | proteasome (prosome, macropain) 26S subunit, ATPase, 4 |
| RV71 | ENSMUSG00000030588 | Yif1b | Yip1 interacting factor homolog B (S. cerevisiae) |
| RV71 | ENSMUSG00000036427 | Gpi1 | glucose-6-phosphate isomerase 1 |
| RV71 | ENSMUSG00000003190 | Bcl2l12 | BCL2 like 12 |
| RV71 | ENSMUSG00000004473 | Clec11a | C-type lectin domain family 11, member a |
| RV71 | ENSMUSG00000026239 | Pde6d | phosphodiesterase 6D, cGMP-specific, rod, delta |
| RV75 | ENSMUSG00000034022 | Cpsf1 | cleavage and polyadenylation specific factor 1 |
| RV75 | ENSMUSG00000022434 | Fam118a | family with sequence similarity 118, member A |
| RV75 | ENSMUSG00000035898 | Uba6 | ubiquitin-like modifier activating enzyme 6 |
| RV75 | ENSMUSG00000042548 | Asxl1 | ASXL transcriptional regulator 1 |
| RV75 | ENSMUSG00000060988 | Galnt13 | polypeptide N-acetylgalactosaminyltransferase 13 |
| RV75 | ENSMUSG00000028683 | Eif2b3 | eukaryotic translation initiation factor 2B, subunit 3 |
| RV75 | ENSMUSG00000038497 | Tmco3 | transmembrane and coiled-coil domains 3 |
| RV75 | ENSMUSG00000014074 | Rnf168 | ring finger protein 168 |
| RV75 | ENSMUSG00000020850 | Prpf8 | pre-mRNA processing factor 8 |
| RV75 | ENSMUSG00000010122 | Slc47a1 | solute carrier family 47, member 1 |
| RV75 | ENSMUSG00000056895 | H2bc27 | H2B clustered histone 27 |
| RV75 | ENSMUSG00000028863 | Meaf6 | MYST/Esa1-associated factor 6 |
| RV75 | ENSMUSG00000040025 | Ythdf2 | YTH N6-methyladenosine RNA binding protein 2 |
| RV75 | ENSMUSG00000024164 | C3 | complement component 3 |
| RV75 | ENSMUSG00000024990 | Rbp4 | retinol binding protein 4, plasma |
| RV75 | ENSMUSG00000024856 | Cdk2ap2 | cyclin dependent kinase 2 associated protein 2 |
| RV75 | ENSMUSG00000030674 | Qprt | quinolinate phosphoribosyltransferase |
| RV75 | ENSMUSG00000040325 | Dcaf1 | DDB1 and CUL4 associated factor 1 |

| | | | |
|---|---|---|---|
| RV75 | ENSMUSG00000020258 | Glyctk | glycerate kinase |
| RV75 | ENSMUSG00000032410 | Xrn1 | 5'-3' exoribonuclease 1 |
| RV75 | ENSMUSG00000044447 | Dock5 | dedicator of cytokinesis 5 |
| RV75 | ENSMUSG00000041341 | Atg2b | autophagy related 2B |
| RV75 | ENSMUSG00000038160 | Atg5 | autophagy related 5 |
| RV75 | ENSMUSG00000026031 | Cflar | CASP8 and FADD-like apoptosis regulator |
| RV75 | ENSMUSG00000033285 | Wdr3 | WD repeat domain 3 |
| RV75 | ENSMUSG00000025792 | Slc25a10 | solute carrier family 25 (mitochondrial carrier, dicarboxylate transporter), member 10 |
| RV75 | ENSMUSG00000041920 | Slc16a6 | solute carrier family 16 (monocarboxylic acid transporters), member 6 |
| RV75 | ENSMUSG00000078653 | Cntd1 | cyclin N-terminal domain containing 1 |
| RV75 | ENSMUSG00000001755 | Coasy | Coenzyme A synthase |
| RV75 | ENSMUSG00000031709 | Tbc1d9 | TBC1 domain family, member 9 |
| RV75 | ENSMUSG00000047264 | Zfp358 | zinc finger protein 358 |
| RV75 | ENSMUSG00000037211 | Spry1 | sprouty RTK signaling antagonist 1 |
| RV75 | ENSMUSG00000090877 | Hspa1b | heat shock protein 1B |
| RV75 | ENSMUSG00000044566 | Cage1 | cancer antigen 1 |
| RV75 | ENSMUSG00000037946 | Fgd3 | FYVE, RhoGEF and PH domain containing 3 |
| RV75 | ENSMUSG00000060477 | Irak2 | interleukin-1 receptor-associated kinase 2 |
| RV75 | ENSMUSG00000030270 | Cpne9 | copine family member IX |
| RV75 | ENSMUSG00000089678 | Agxt2 | alanine-glyoxylate aminotransferase 2 |
| RV75 | ENSMUSG00000028630 | Dyrk2 | dual-specificity tyrosine phosphorylation regulated kinase 2 |
| RV75 | ENSMUSG00000048096 | Lmod1 | leiomodin 1 (smooth muscle) |
| RV75 | ENSMUSG00000029547 | Ints1 | integrator complex subunit 1 |
| RV75 | ENSMUSG00000056394 | Lig1 | ligase I, DNA, ATP-dependent |
| RV75 | ENSMUSG00000006019 | Dhx34 | DExH-box helicase 34 |
| RV75 | ENSMUSG00000037463 | Fbxo27 | F-box protein 27 |
| RV75 | ENSMUSG00000030613 | Ccdc90b | coiled-coil domain containing 90B |
| RV75 | ENSMUSG00000047879 | Usp14 | ubiquitin specific peptidase 14 |
| RV75 | ENSMUSG00000035356 | Nfkbiz | nuclear factor of kappa light polypeptide gene enhancer in B cells inhibitor, zeta |
| RV76 | ENSMUSG00000022377 | Asap1 | ArfGAP with SH3 domain, ankyrin repeat and PH domain1 |
| RV76 | ENSMUSG00000048175 | Asb8 | ankyrin repeat and SOCS box-containing 8 |
| RV76 | ENSMUSG00000006471 | Ndor1 | NADPH dependent diflavin oxidoreductase 1 |
| RV76 | ENSMUSG00000037295 | Ldlrap1 | low density lipoprotein receptor adaptor protein 1 |
| RV76 | ENSMUSG00000029009 | Mthfr | methylenetetrahydrofolate reductase |
| RV76 | ENSMUSG00000026723 | Trdmt1 | tRNA aspartic acid methyltransferase 1 |
| RV76 | ENSMUSG00000031819 | Emc8 | ER membrane protein complex subunit 8 |
| RV76 | ENSMUSG00000048120 | Entpd1 | ectonucleoside triphosphate diphosphohydrolase 1 |
| RV76 | ENSMUSG00000025485 | Ric8a | RIC8 guanine nucleotide exchange factor A |

| | | | |
|---|---|---|---|
| RV76 | ENSMUSG00000007989 | Fzd3 | frizzled class receptor 3 |
| RV76 | ENSMUSG00000019843 | Fyn | Fyn proto-oncogene |
| RV76 | ENSMUSG00000038446 | Cdc40 | cell division cycle 40 |
| RV76 | ENSMUSG00000026036 | Nif3l1 | Ngg1 interacting factor 3-like 1 (S. pombe) |
| RV76 | ENSMUSG00000026179 | Pnkd | paroxysmal nonkinesiogenic dyskinesia |
| RV76 | ENSMUSG00000028044 | Cks1b | CDC28 protein kinase 1b |
| RV76 | ENSMUSG00000105827 | H2bc18 | H2B clustered histone 18 |
| RV76 | ENSMUSG00000015947 | Fcgr1 | Fc receptor, IgG, high affinity I |
| RV76 | ENSMUSG00000087610 | Gm16253 | predicted gene 16253 |
| RV76 | ENSMUSG00000000056 | Narf | nuclear prelamin A recognition factor |
| RV76 | ENSMUSG00000038811 | Gngt2 | guanine nucleotide binding protein (G protein), gamma transducing activity polypeptide 2 |
| RV76 | ENSMUSG00000036353 | P2ry12 | purinergic receptor P2Y, G-protein coupled 12 |
| RV76 | ENSMUSG00000022978 | Mis18a | MIS18 kinetochore protein A |
| RV76 | ENSMUSG00000044442 | N6amt1 | N-6 adenine-specific DNA methyltransferase 1 (putative) |
| RV76 | ENSMUSG00000067586 | S1pr3 | sphingosine-1-phosphate receptor 3 |
| RV76 | ENSMUSG00000030122 | Ptms | parathymosin |
| RV76 | ENSMUSG00000053289 | Ddx10 | DEAD box helicase 10 |
| RV76 | ENSMUSG00000045519 | Zfp560 | zinc finger protein 560 |
| RV76 | ENSMUSG00000031119 | Gpc4 | glypican 4 |
| RV76 | ENSMUSG00000025262 | Fam120c | family with sequence similarity 120, member C |
| RV76 | ENSMUSG00000031060 | Rbm10 | RNA binding motif protein 10 |
| RV76 | ENSMUSG00000019975 | Ikbip | IKBKB interacting protein |
| RV76 | ENSMUSG00000030032 | Wdr54 | WD repeat domain 54 |
| RV76 | ENSMUSG00000029817 | Tra2a | transformer 2 alpha |
| RV76 | ENSMUSG00000039701 | Usp53 | ubiquitin specific peptidase 53 |
| RV76 | ENSMUSG00000056536 | Pign | phosphatidylinositol glycan anchor biosynthesis, class N |
| RV76 | ENSMUSG00000040713 | Creg1 | cellular repressor of E1A-stimulated genes 1 |
| RV76 | ENSMUSG00000029386 | Tctn2 | Tectonic family member 2 |
| RV76 | ENSMUSG00000029388 | Eif2b1 | eukaryotic translation initiation factor 2B, subunit alpha |
| RV76 | ENSMUSG00000000915 | Hip1r | huntingtin interacting protein 1 related |
| RV76 | ENSMUSG00000042190 | Cmklr1 | chemerin chemokine-like receptor 1 |
| RV76 | ENSMUSG00000007207 | Stx1a | syntaxin 1A (brain) |
| RV76 | ENSMUSG00000029729 | Zkscan1 | zinc finger with KRAB and SCAN domains 1 |
| RV76 | ENSMUSG00000002984 | Tomm40 | translocase of outer mitochondrial membrane 40 |
| RV76 | ENSMUSG00000005686 | Ampd3 | adenosine monophosphate deaminase 3 |
| RV76 | ENSMUSG00000030726 | Pold3 | polymerase (DNA-directed), delta 3, accessory subunit |
| RV76 | ENSMUSG00000022723 | Crybg3 | beta-gamma crystallin domain containing 3 |
| RV77 | ENSMUSG00000022552 | Sharpin | SHANK-associated RH domain interacting protein |

| | | | |
|---|---|---|---|
| RV77 | ENSMUSG00000023034 | Nr4a1 | nuclear receptor subfamily 4, group A, member 1 |
| RV77 | ENSMUSG00000029368 | Alb | albumin |
| RV77 | ENSMUSG00000014353 | Tmem87b | transmembrane protein 87B |
| RV77 | ENSMUSG00000057228 | Aadat | aminoadipate aminotransferase |
| RV77 | ENSMUSG00000002908 | Kcnn1 | potassium intermediate/small conductance calcium-activated channel, subfamily N, member 1 |
| RV77 | ENSMUSG00000045394 | Epcam | epithelial cell adhesion molecule |
| RV77 | ENSMUSG00000034709 | Ppp1r21 | protein phosphatase 1, regulatory subunit 21 |
| RV77 | ENSMUSG00000013663 | Pten | phosphatase and tensin homolog |
| RV77 | ENSMUSG00000024812 | Tjp2 | tight junction protein 2 |
| RV77 | ENSMUSG00000009545 | Kcnq1 | potassium voltage-gated channel, subfamily Q, member 1 |
| RV77 | ENSMUSG00000002957 | Ap2a2 | adaptor-related protein complex 2, alpha 2 subunit |
| RV77 | ENSMUSG00000025241 | Fyco1 | FYVE and coiled-coil domain containing 1 |
| RV77 | ENSMUSG00000035769 | Xylb | xylulokinase homolog (H. influenzae) |
| RV77 | ENSMUSG00000032369 | Plscr1 | phospholipid scramblase 1 |
| RV77 | ENSMUSG00000032348 | Gsta4 | glutathione S-transferase, alpha 4 |
| RV77 | ENSMUSG00000014547 | Wdfy2 | WD repeat and FYVE domain containing 2 |
| RV77 | ENSMUSG00000039081 | Zfp503 | zinc finger protein 503 |
| RV77 | ENSMUSG00000028412 | Slc44a1 | solute carrier family 44, member 1 |
| RV77 | ENSMUSG00000021228 | Acot3 | acyl-CoA thioesterase 3 |
| RV77 | ENSMUSG00000072949 | Acot1 | acyl-CoA thioesterase 1 |
| RV77 | ENSMUSG00000053398 | Phgdh | 3-phosphoglycerate dehydrogenase |
| RV77 | ENSMUSG00000039450 | Dcxr | dicarbonyl L-xylulose reductase |
| RV77 | ENSMUSG00000034282 | Evpl | envoplakin |
| RV77 | ENSMUSG00000052837 | Junb | jun B proto-oncogene |
| RV77 | ENSMUSG00000057672 | Pkn1 | protein kinase N1 |
| RV77 | ENSMUSG00000024352 | Spata24 | spermatogenesis associated 24 |
| RV77 | ENSMUSG00000053453 | Thoc7 | THO complex 7 |
| RV77 | ENSMUSG00000006527 | Sfmbt1 | Scm-like with four mbt domains 1 |
| RV77 | ENSMUSG00000052395 | Rft1 | RFT1 homolog |
| RV77 | ENSMUSG00000004568 | Arhgef18 | Rho/Rac guanine nucleotide exchange factor 18 |
| RV77 | ENSMUSG00000030545 | Pex11a | peroxisomal biogenesis factor 11 alpha |
| RV77 | ENSMUSG00000047141 | Zfp654 | zinc finger protein 654 |
| RV77 | ENSMUSG00000049037 | Clec4a1 | C-type lectin domain family 4, member a1 |
| RV77 | ENSMUSG00000030316 | Tamm41 | TAM41 mitochondrial translocator assembly and maintenance homolog |
| RV77 | ENSMUSG00000041052 | Slc7a13 | solute carrier family 7, (cationic amino acid transporter, y+ system) member 13 |
| RV77 | ENSMUSG00000039438 | Ttc36 | tetratricopeptide repeat domain 36 |
| RV77 | ENSMUSG00000031337 | Mtm1 | X-linked myotubular myopathy gene 1 |
| RV77 | ENSMUSG00000020102 | Slc16a7 | solute carrier family 16 (monocarboxylic acid transporters), member 7 |

| | | | |
|---|---|---|---|
| RV77 | ENSMUSG00000028179 | Cth | cystathionine gamma lyase |
| RV77 | ENSMUSG00000068523 | Gng5 | G protein subunit gamma 5 |
| RV77 | ENSMUSG00000079283 | 2310009B15Rik | RIKEN cDNA 2310009B15 gene |
| RV77 | ENSMUSG00000041130 | Zfp598 | zinc finger protein 598 |
| RV77 | ENSMUSG00000036560 | Lgi4 | leucine-rich repeat LGI family, member 4 |
| RV77 | ENSMUSG00000030492 | Slc7a9 | solute carrier family 7 (cationic amino acid transporter, y+ system), member 9 |
| RV81 | ENSMUSG00000022438 | Parvb | parvin, beta |
| RV81 | ENSMUSG00000022454 | Nell2 | NEL-like 2 |
| RV81 | ENSMUSG00000029373 | Pf4 | platelet factor 4 |
| RV81 | ENSMUSG00000029372 | Ppbp | pro-platelet basic protein |
| RV81 | ENSMUSG00000005802 | Slc30a4 | solute carrier family 30 (zinc transporter), member 4 |
| RV81 | ENSMUSG00000027366 | Sppl2a | signal peptide peptidase like 2A |
| RV81 | ENSMUSG00000074622 | Mafb | MAF bZIP transcription factor B |
| RV81 | ENSMUSG00000050761 | Gp1bb | glycoprotein Ib, beta polypeptide |
| RV81 | ENSMUSG00000033653 | Vps8 | VPS8 CORVET complex subunit |
| RV81 | ENSMUSG00000035699 | Slc51a | solute carrier family 51, alpha subunit |
| RV81 | ENSMUSG00000018334 | Ksr1 | kinase suppressor of ras 1 |
| RV81 | ENSMUSG00000020788 | Atp2a3 | ATPase, Ca++ transporting, ubiquitous |
| RV81 | ENSMUSG00000023075 | Akirin1 | akirin 1 |
| RV81 | ENSMUSG00000058908 | Pla2g2a | phospholipase A2, group IIA (platelets, synovial fluid) |
| RV81 | ENSMUSG00000024983 | Vti1a | vesicle transport through interaction with t-SNAREs 1A |
| RV81 | ENSMUSG00000024965 | Fermt3 | fermitin family member 3 |
| RV81 | ENSMUSG00000010755 | Cars1 | cysteinyl-tRNA synthetase 1 |
| RV81 | ENSMUSG00000028437 | Ubap1 | ubiquitin-associated protein 1 |
| RV81 | ENSMUSG00000001467 | Cyp51 | cytochrome P450, family 51 |
| RV81 | ENSMUSG00000039232 | Stx11 | syntaxin 11 |
| RV81 | ENSMUSG00000019998 | Stx7 | syntaxin 7 |
| RV81 | ENSMUSG00000004233 | Wars2 | tryptophanyl tRNA synthetase 2 (mitochondrial) |
| RV81 | ENSMUSG00000020689 | Itgb3 | integrin beta 3 |
| RV81 | ENSMUSG00000034664 | Itga2b | integrin alpha 2b |
| RV81 | ENSMUSG00000018659 | Pnpo | pyridoxine 5'-phosphate oxidase |
| RV81 | ENSMUSG00000023993 | Treml1 | triggering receptor expressed on myeloid cells-like 1 |
| RV81 | ENSMUSG00000025429 | Pstpip2 | proline-serine-threonine phosphatase-interacting protein 2 |
| RV81 | ENSMUSG00000037325 | Bbs7 | Bardet-Biedl syndrome 7 |
| RV81 | ENSMUSG00000022962 | Gart | phosphoribosylglycinamide formyltransferase |
| RV81 | ENSMUSG00000039109 | F13a1 | coagulation factor XIII, A1 subunit |
| RV81 | ENSMUSG00000030159 | Clec1b | C-type lectin domain family 1, member b |
| RV81 | ENSMUSG00000000838 | Fmr1 | fragile X messenger ribonucleoprotein 1 |

| | | | |
|---|---|---|---|
| RV81 | ENSMUSG00000037005 | Xpnpep2 | X-prolyl aminopeptidase (aminopeptidase P) 2, membrane-bound |
| RV81 | ENSMUSG00000016534 | Lamp2 | lysosomal-associated membrane protein 2 |
| RV81 | ENSMUSG00000030054 | Gp9 | glycoprotein 9 platelet |
| RV81 | ENSMUSG00000020120 | Plek | pleckstrin |
| RV81 | ENSMUSG00000034201 | Gas2l1 | growth arrest-specific 2 like 1 |
| RV81 | ENSMUSG00000071256 | Zfp213 | zinc finger protein 213 |
| RV81 | ENSMUSG00000026579 | F5 | coagulation factor V |
| RV81 | ENSMUSG00000040297 | Suco | SUN domain containing ossification factor |
| RV81 | ENSMUSG00000000916 | Nsun5 | NOL1/NOP2/Sun domain family, member 5 |
| RV81 | ENSMUSG00000078810 | Gp6 | glycoprotein 6 platelet |
| RV81 | ENSMUSG00000081984 | Dnajb3 | DnaJ heat shock protein family (Hsp40) member B3 |
| RV87 | ENSMUSG00000115798 | Gm55359 | predicted gene, 55359 |
| RV87 | ENSMUSG00000035891 | Cerk | ceramide kinase |
| RV87 | ENSMUSG00000058794 | Nfe2 | nuclear factor, erythroid derived 2 |
| RV87 | ENSMUSG00000067787 | Blcap | bladder cancer associated protein |
| RV87 | ENSMUSG00000061313 | Ddhd2 | DDHD domain containing 2 |
| RV87 | ENSMUSG00000003233 | Dvl3 | dishevelled segment polarity protein 3 |
| RV87 | ENSMUSG00000061981 | Flot2 | flotillin 2 |
| RV87 | ENSMUSG00000000320 | Alox12 | arachidonate 12-lipoxygenase |
| RV87 | ENSMUSG00000042826 | Fgf11 | fibroblast growth factor 11 |
| RV87 | ENSMUSG00000036819 | Jmjd4 | jumonji domain containing 4 |
| RV87 | ENSMUSG00000061894 | Zscan20 | zinc finger and SCAN domains 20 |
| RV87 | ENSMUSG00000028741 | Mrto4 | mRNA turnover 4, ribosome maturation factor |
| RV87 | ENSMUSG00000057637 | Prdm2 | PR domain containing 2, with ZNF domain |
| RV87 | ENSMUSG00000023286 | Ube2j2 | ubiquitin-conjugating enzyme E2J 2 |
| RV87 | ENSMUSG00000025195 | Dnmbp | dynamin binding protein |
| RV87 | ENSMUSG00000059363 | Fxn | frataxin |
| RV87 | ENSMUSG00000047379 | B4gat1 | beta-1,4-glucuronyltransferase 1 |
| RV87 | ENSMUSG00000025481 | Urah | urate (5-hydroxyiso-) hydrolase |
| RV87 | ENSMUSG00000045598 | Zfp553 | zinc finger protein 553 |
| RV87 | ENSMUSG00000035078 | Mtmr9 | myotubularin related protein 9 |
| RV87 | ENSMUSG00000021265 | Slc25a29 | solute carrier family 25 (mitochondrial carrier, palmitoylcarnitine transporter), member 29 |
| RV87 | ENSMUSG00000045064 | Zc2hc1c | zinc finger, C2HC-type containing 1C |
| RV87 | ENSMUSG00000028019 | Pdgfc | platelet-derived growth factor, C polypeptide |
| RV87 | ENSMUSG00000028078 | Dclk2 | doublecortin-like kinase 2 |
| RV87 | ENSMUSG00000027933 | Ints3 | integrator complex subunit 3 |
| RV87 | ENSMUSG00000031714 | Gab1 | growth factor receptor bound protein 2-associated protein 1 |
| RV87 | ENSMUSG00000041915 | Ammecr1l | AMME chromosomal region gene 1-like |

| | | | |
|---|---|---|---|
| RV87 | ENSMUSG00000038784 | Cnot4 | CCR4-NOT transcription complex, subunit 4 |
| RV87 | ENSMUSG00000025104 | Hdgfl3 | HDGF like 3 |
| RV87 | ENSMUSG00000033458 | Fan1 | FANCD2/FANCI-associated nuclease 1 |
| RV87 | ENSMUSG00000020070 | Rufy2 | RUN and FYVE domain-containing 2 |
| RV87 | ENSMUSG00000001630 | Stk38l | serine/threonine kinase 38 like |
| RV87 | ENSMUSG00000030137 | Tuba8 | tubulin, alpha 8 |
| RV87 | ENSMUSG00000097221 | 1810049J17Rik | RIKEN cDNA 1810049J17 gene |
| RV87 | ENSMUSG00000037709 | Fam13a | family with sequence similarity 13, member A |
| RV87 | ENSMUSG00000027983 | Cyp2u1 | cytochrome P450, family 2, subfamily u, polypeptide 1 |
| RV87 | ENSMUSG00000092572 | Serpinb10 | serine (or cysteine) peptidase inhibitor, clade B (ovalbumin), member 10 |
| RV87 | ENSMUSG00000026554 | Dcaf8 | DDB1 and CUL4 associated factor 8 |
| RV87 | ENSMUSG00000029627 | Zkscan14 | zinc finger with KRAB and SCAN domains 14 |
| RV87 | ENSMUSG00000057101 | Zfp180 | zinc finger protein 180 |
| RV87 | ENSMUSG00000002475 | Abhd3 | abhydrolase domain containing 3 |

**Table S10.** Hub genes of RV transcriptional modules.

| Module | ENSEMBL_ID | gene_ID | Description |
|---|---|---|---|
| RV1 | ENSMUSG00000067288 | Rps28 | ribosomal protein S28 |
| RV2 | ENSMUSG00000017817 | Jph2 | junctophilin 2 |
| RV3 | ENSMUSG00000034290 | Nek9 | NIMA (never in mitosis gene a)-related expressed kinase 9 |
| RV4 | ENSMUSG00000056666 | Retsat | retinol saturase (all trans retinol 13,14 reductase) |
| RV5 | ENSMUSG00000020869 | Lrrc59 | leucine rich repeat containing 59 |
| RV6 | ENSMUSG00000051855 | Mest | mesoderm specific transcript |
| RV7 | ENSMUSG00000030247 | Kcnj8 | potassium inwardly-rectifying channel, subfamily J, member 8 |
| RV8 | ENSMUSG00000022102 | Dok2 | docking protein 2 |
| RV9 | ENSMUSG00000068699 | Flnc | filamin C, gamma |
| RV10 | ENSMUSG00000011658 | Fuz | fuzzy planar cell polarity protein |
| RV11 | ENSMUSG00000025728 | Pigq | phosphatidylinositol glycan anchor biosynthesis, class Q |
| RV12 | ENSMUSG00000030793 | Pycard | PYD and CARD domain containing |
| RV13 | ENSMUSG00000036278 | Macrod1 | mono-ADP ribosylhydrolase 1 |
| RV14 | ENSMUSG00000071506 | Tmem139 | transmembrane protein 139 |
| RV15 | ENSMUSG00000032357 | Tinag | tubulointerstitial nephritis antigen |
| RV16 | ENSMUSG00000041842 | Fhdc1 | FH2 domain containing 1 |
| RV17 | ENSMUSG00000052906 | Ubxn8 | UBX domain protein 8 |
| RV18 | ENSMUSG00000048000 | Gigyf2 | GRB10 interacting GYF protein 2 |
| RV19 | ENSMUSG00000003226 | Ranbp2 | RAN binding protein 2 |
| RV20 | ENSMUSG00000024338 | Psmb8 | proteasome (prosome, macropain) subunit, beta type 8 (large multifunctional peptidase 7) |
| RV21 | ENSMUSG00000051346 | Spryd4 | SPRY domain containing 4 |
| RV22 | ENSMUSG00000002803 | Btbd6 | BTB domain containing 6 |
| RV23 | ENSMUSG00000003200 | Sh3gl1 | SH3-domain GRB2-like 1 |
| RV24 | ENSMUSG00000020212 | Mdm1 | MDM1 nuclear protein |
| RV25 | ENSMUSG00000054720 | Lrrc8c | leucine rich repeat containing 8 family, member C |
| RV26 | ENSMUSG00000054309 | Cpsf3 | cleavage and polyadenylation specificity factor 3 |
| RV27 | ENSMUSG00000049892 | Rasd1 | RAS, dexamethasone-induced 1 |
| RV28 | ENSMUSG00000066233 | Tmem42 | transmembrane protein 42 |
| RV29 | ENSMUSG00000022090 | Pdlim2 | PDZ and LIM domain 2 |
| RV30 | ENSMUSG00000030965 | Abraxas2 | BRISC complex subunit |
| RV31 | ENSMUSG00000044864 | Ankrd50 | ankyrin repeat domain 50 |
| RV32 | ENSMUSG00000039834 | Zfp335 | zinc finger protein 335 |
| RV33 | ENSMUSG00000059495 | Arhgef12 | Rho guanine nucleotide exchange factor 12 |
| RV34 | ENSMUSG00000078853 | Igtp | interferon gamma induced GTPase |
| RV35 | ENSMUSG00000029922 | Mkrn1 | makorin, ring finger protein, 1 |
| RV36 | ENSMUSG00000030411 | Nova2 | NOVA alternative splicing regulator 2 |

| | | | |
|---|---|---|---|
| RV37 | ENSMUSG00000019433 | Gipc1 | GIPC PDZ domain containing family, member 1 |
| RV38 | ENSMUSG00000049436 | Upk1b | uroplakin 1B |
| RV39 | ENSMUSG00000029304 | Spp1 | secreted phosphoprotein 1 |
| RV40 | ENSMUSG00000029993 | Nfu1 | NFU1 iron-sulfur cluster scaffold |
| RV41 | ENSMUSG00000029869 | Ephb6 | Eph receptor B6 |
| RV42 | ENSMUSG00000027884 | Clcc1 | chloride channel CLIC-like 1 |
| RV43 | ENSMUSG00000049922 | Slc35c1 | solute carrier family 35, member C1 |
| RV44 | ENSMUSG00000023307 | Marchf5 | membrane associated ring-CH-type finger 5 |
| RV45 | ENSMUSG00000050335 | Lgals3 | lectin, galactose binding, soluble 3 |
| RV46 | ENSMUSG00000021763 | Cspg4b | chondroitin sulfate proteoglycan 4B |
| RV47 | ENSMUSG00000025464 | Paox | polyamine oxidase (exo-N4-amino) |
| RV48 | ENSMUSG00000054414 | Slc30a7 | solute carrier family 30 (zinc transporter), member 7 |
| RV49 | ENSMUSG00000054469 | Lclat1 | lysocardiolipin acyltransferase 1 |
| RV50 | ENSMUSG00000021234 | Fam161b | family with sequence similarity 161, member B |
| RV51 | ENSMUSG00000020697 | Lig3 | ligase III, DNA, ATP-dependent |
| RV52 | ENSMUSG00000033186 | Mzt1 | mitotic spindle organizing protein 1 |
| RV53 | ENSMUSG00000039157 | Eeig1 | estrogen-induced osteoclastogenesis regulator 1 |
| RV54 | ENSMUSG00000050211 | Pla2g4e | phospholipase A2, group IVE |
| RV55 | ENSMUSG00000037221 | Mospd3 | motile sperm domain containing 3 |
| RV56 | ENSMUSG00000033068 | Entpd6 | ectonucleoside triphosphate diphosphohydrolase 6 |
| RV57 | ENSMUSG00000024736 | Tmem132a | transmembrane protein 132A |
| RV58 | ENSMUSG00000024966 | Stip1 | stress-induced phosphoprotein 1 |
| RV59 | ENSMUSG00000036995 | Asap3 | ArfGAP with SH3 domain, ankyrin repeat and PH domain 3 |
| RV60 | ENSMUSG00000037206 | Islr | immunoglobulin superfamily containing leucine-rich repeat |
| RV61 | ENSMUSG00000055923 | Aasdh | aminoadipate-semialdehyde dehydrogenase |
| RV62 | ENSMUSG00000056899 | Immp2l | IMP2 inner mitochondrial membrane peptidase-like (S. cerevisiae) |
| RV63 | ENSMUSG00000025142 | Aspscr1 | ASPSCR1 tether for SLC2A4, UBX domain containing |
| RV64 | ENSMUSG00000028291 | Akirin2 | akirin 2 |
| RV65 | ENSMUSG00000028832 | Stmn1 | stathmin 1 |
| RV66 | ENSMUSG00000029465 | Arpc3 | actin related protein 2/3 complex, subunit 3 |
| RV67 | ENSMUSG00000019143 | Hars2 | histidyl-tRNA synthetase 2 |
| RV68 | ENSMUSG00000028382 | Ptbp3 | polypyrimidine tract binding protein 3 |
| RV69 | ENSMUSG00000026049 | Tex30 | testis expressed 30 |
| RV70 | ENSMUSG00000034647 | Ankrd12 | ankyrin repeat domain 12 |
| RV71 | ENSMUSG00000052833 | Sae1 | SUMO1 activating enzyme subunit 1 |
| RV72 | ENSMUSG00000071847 | Apcdd1 | adenomatosis polyposis coli down-regulated 1 |
| RV73 | ENSMUSG00000039599 | Fam149b | family with sequence similarity 149, member B |
| RV74 | ENSMUSG00000025171 | Ubtd1 | ubiquitin domain containing 1 |
| RV75 | ENSMUSG00000030674 | Qprt | quinolinate phosphoribosyltransferase |

| | | | |
|---|---|---|---|
| RV76 | ENSMUSG00000007989 | Fzd3 | frizzled class receptor 3 |
| RV77 | ENSMUSG00000041052 | Slc7a13 | solute carrier family 7, (cationic amino acid transporter, y+ system) member 13 |
| RV78 | ENSMUSG00000040859 | Bsdc1 | BSD domain containing 1 |
| RV79 | ENSMUSG00000096199 | Ptrhd1 | peptidyl-tRNA hydrolase domain containing 1 |
| RV80 | ENSMUSG00000036873 | 2410004B18Rik | RIKEN cDNA 2410004B18 gene |
| RV81 | ENSMUSG00000078810 | Gp6 | glycoprotein 6 platelet |
| RV82 | ENSMUSG00000006392 | Med8 | mediator complex subunit 8 |
| RV83 | ENSMUSG00000041895 | Wipi1 | WD repeat domain, phosphoinositide interacting 1 |
| RV84 | ENSMUSG00000028971 | Cort | cortistatin |
| RV85 | ENSMUSG00000033964.3 | Zbtb41 | zinc finger and BTB domain containing 41 |
| RV86 | ENSMUSG00000045248 | Med26 | mediator complex subunit 26 |
| RV87 | ENSMUSG00000035078 | Mtmr9 | myotubularin related protein 9 |
| RV88 | ENSMUSG00000037622 | Wdtc1 | WD and tetratricopeptide repeats 1 |
| RV89 | ENSMUSG00000035235 | Trim13 | tripartite motif-containing 13 |
| RV90 | ENSMUSG00000044950 | Pwwp2a | PWWP domain containing 2A |
| RV91 | ENSMUSG00000070469 | Adamtsl3 | ADAMTS-like 3 |
| RV92 | ENSMUSG00000062604 | Srpk2 | serine/arginine-rich protein specific kinase 2 |
| RV93 | ENSMUSG00000056313 | Tcim | transcriptional and immune response regulator |
| RV94 | ENSMUSG00000063810 | Alms1 | ALMS1, centrosome and basal body associated |
| RV95 | ENSMUSG00000021482 | Prxl2c | peroxiredoxin like 2C |
| RV96 | ENSMUSG00000031548 | Sfrp1 | secreted frizzled-related protein 1 |
| RV97 | ENSMUSG00000024485 | Slc4a9 | solute carrier family 4, sodium bicarbonate cotransporter, member 9 |
| RV98 | ENSMUSG00000026610 | Esrrg | estrogen-related receptor gamma |
| RV99 | ENSMUSG00000024661 | Fth1 | ferritin heavy polypeptide 1 |
| RV100 | ENSMUSG00000027875 | Hmgcs2 | 3-hydroxy-3-methylglutaryl-Coenzyme A synthase 2 |
| RV101 | ENSMUSG00000109865 | Hspa14 | heat shock protein 14 |
| RV102 | ENSMUSG00000033327 | Tnxb | tenascin XB |
